# Dividing out quantification uncertainty allows efficient assessment of differential transcript expression with edgeR

**DOI:** 10.1101/2023.04.02.535231

**Authors:** Pedro L. Baldoni, Yunshun Chen, Soroor Hediyeh-zadeh, Yang Liao, Xueyi Dong, Matthew E. Ritchie, Wei Shi, Gordon K. Smyth

## Abstract

Differential expression analysis of RNA-seq is one of the most commonly performed bioinformatics analyses. Transcript-level quantifications are inherently more uncertain than gene-level read counts because of ambiguous assignment of sequence reads to transcripts. While sequence reads can usually be assigned unambiguously to a gene, reads are very often compatible with multiple transcripts for that gene, particularly for genes with many isoforms. Software tools designed for gene-level differential expression do not perform optimally on transcript counts because the read-to-transcript ambiguity (RTA) disrupts the mean-variance relationship normally observed for gene level RNA-seq data and interferes with the efficiency of the empirical Bayes dispersion estimation procedures. The pseudoaligners kallisto and Salmon provide bootstrap samples from which quantification uncertainty can be assessed. We show that the overdispersion arising from RTA can be elegantly estimated by fitting a quasi-Poisson model to the bootstrap counts for each transcript. The technical overdispersion arising from RTA can then be divided out of the transcript counts, leading to scaled counts that can be input for analysis by established gene-level software tools with full statistical efficiency. Comprehensive simulations and test data show that an edgeR analysis of the scaled counts is more powerful and efficient than previous differential transcript expression pipelines while providing correct control of the false discovery rate. Simulations explore a wide range of scenarios including the effects of paired vs single-end reads, different read lengths and different numbers of replicates.

## Introduction

Over the past fifteen years, RNA sequencing (RNA-seq) has proved to be a tremendously popular and powerful technology for profiling the transcriptomes of biological samples [1]. Analysis of RNA-seq data often focuses on the detection of differential expression between biological conditions such as treated vs control samples, tumor vs healthy tissues, distinct cell lines, or genetically modified vs wildtype organisms [2]. Differential expression analyses have most often been conducted to identify differentially expressed genes, but analyses may also be conducted to detect differential splicing between conditions or to identify the specific gene isoforms (transcripts) that are differentially expressed [3]. With reduced sequencing costs and the recent development of lightweight alignment algorithms [4, 5], there is growing interest in the assessment of differential expression at the transcript level.

Gene-level differential expression analyses have typically assumed a negative binomial (NB) distribution for the number of sequence reads assigned to each gene [6, 7, 8, 9]. The NB distribution can be viewed as a mathematical mixture of Poisson distributions, with a gamma distribution as the mixing distribution. In this representation, the gamma distribution represents the biological variation in expression levels between RNA samples and the Poisson distribution represents measurement error for individual samples [9]. Quasi-NB models have also been used in order to provide more rigorous false discovery rate control [10, 11].

Gene-level read counts analysed by software tools such as *edgeR* [12], *DESeq2* [13] or *voom* [14] may be integers from overlap counters like *featureCounts* [15] or *HTSeq* [16] or fractional estimated counts from transcript quantifiers such as *RSEM* [17], *kallisto* [4] or *Salmon* [5]. The *edgeR* package in particular implements a continuous generalization of the NB distribution so that fractional counts may be input without rounding or loss of information.

Transcript quantifications are inherently more uncertain than gene-level read counts because of ambiguous assignment of RNA fragments to isoforms [18]. Whereas different genes typically occupy non-overlapping regions of the genome, different transcripts for the same gene typically share one or more exons in common. While sequence reads can usually be assigned unambiguously to a gene, reads are very often compatible with multiple transcripts for that gene, particularly for genes with many isoforms. This phenomenon has variously been called assignment ambiguity, mapping ambiguity, quantification uncertainty or inferential uncertainty [19, 20, 21]. In this article we will call it *read to transcript ambiguity* (RTA) to avoid confusion with other sources of uncertainty.

Software tools designed for gene-level differential expression do not perform optimally on transcript counts because RTA disrupts the mean-variance relationship normally observed for gene level RNA-seq data and therefore interferes with the efficiency of the empirical Bayes dispersion estimation procedures. The purpose of the current article is to show that RTA can be elegantly modeled using an overdispersed Poisson distribution for each transcript, thus generalizing the measurement error part of the traditional NB model. The technical overdispersion arising from RTA can then be divided out of the transcript counts, leading to scaled counts that can be input for analysis by established gene-level software tools with full statistical efficiency.

The lightweight alignment tools *kallisto* [4] and *Salmon* [5] perform pseudo-alignment of RNA-seq reads to the transcriptome, classifying reads into equivalence classes according to compatibility with annotated transcripts followed by probabilistic assignment of sequence reads to transcripts. This strategy is computationally faster than any previous alignment or quantification approach for RNA-seq data [18]. The efficiency of the equivalence class representation allows technical bootstrap samples to be drawn rapidly for each RNA sample, providing a way to estimate the sampling distribution of the transcript-specific quantifications.

The statistical tools *sleuth* [20] and *Swish* [21] have been proposed to test for differential transcript expression (DTE) using output from *kallisto* or *Salmon* while leveraging the bootstrap samples to estimate the uncertainty associated with RTA. *Swish* uses Wilcoxon tests [22] computed on median-ratio scaled counts from each bootstrap sample and averaged over the bootstrap resamples. Bootstrap counts are scaled to adjust for sample-specific transcript length and sequencing depth via median-ratio size factors. *Swish* can test for DTE between two conditions of interest for single or multi-batch RNA-seq experiments via stratification. *Swish* can also perform DTE analyses for paired samples from a single group via signed-rank test statistics or for paired samples between two conditions via Wilcoxon tests computed on the log-fold change of pairs. *sleuth* on the other hand tests for DTE using an approach similar to *voom* with the addition of a measurement error model. Counts are normalized using sample-specific median-ratio size factors [7] followed by a started log-transformation to ensure positivity and normality. The measurement error model decomposes the total variance into a biological variance component, interpreted as arising from between-sample variation, and an inferential variance component resulting from sample-specific alignment and quantification uncertainty. The biological variance component is estimated using empirical Bayes moderation as for *voom* and the inferential variance component is estimated from *kallisto*’s bootstrap samples. DTE is assessed under a linear model framework using either likelihood ratio or Wald tests.

Our approach instead generalizes the successful NB model for gene-level read counts to account for RTA. The traditional Poisson model for sequencing variability is replaced by a quasi-Poisson model fitted to the bootstrap counts for each transcript. The quasi-Poisson dispersion estimates the variance-inflation induced by RTA and can be used to scale down the transcript counts so that the resulting library sizes reflect their true precision. The scaled counts can be shown to follow the traditional NB mean-variance relationship so that standard methods designed for the differential expression analyses at the gene-level can be applied without further modification.

Our DTE approach is implemented in the Bioconductor package *edgeR*. The *edgeR* functions *catchSalmon* and *catchKallisto* import transcript-counts and associated bootstrap resamples from *Salmon* and *kallisto*, respectively, and estimate the RTA-induced overdispersions. Downstream DTE analyses can then be conducted on scaled-transcript counts within the established *edgeR* framework exactly as for gene-level analyses. Our approach is shown to be both powerful and efficient while correctly controlling the false discovery rate using extensive simulations designed with the aid of the *Rsubread* Bioconductor package [23]. Our method is further demonstrated on a case study of human lung adenocarcinoma cell lines using both short-read Illumina and long-read Nanopore RNA-seq data.

## MATERIALS AND METHODS

### Simulated datasets

#### Reference RNA-seq dataset

A subset of genes and their transcripts, from which sequence reads were simulated, was selected from the mouse Gencode transcript annotation M27 using a real RNA-seq experiment as a reference (NCBI Gene Expression Omnibus series GSE60450). Specifically, we selected protein-coding and lncRNA transcripts from expressed protein-coding and lncRNA genes of the mouse autosome and sex chromosomes. Genes with expected counts-per-million greater than 1 in at least 6 of the 12 RNA-seq samples were considered to be expressed, which resulted in a reference list of 13,176 genes and 41,372 associated transcripts. See Supplementary Data Section 1.1.1 for further details.

#### Simulation of RNA-seq sequence reads

We used the Bioconductor package *Rsubread* and its function *simReads* to simulate sequence reads in FASTQ file format for the selected reference list of transcripts in a number of scenarios [23]. Scenarios varied with respect to the library size (either balanced with 50 Mi.reads, or unbalanced with alternating 25 Mi. and 100 Mi. reads over samples), the sequence read length and type (paired-end or single-end reads with 50, 75, 100, 125, or 150 base pairs (bp) long), the maximum number of expressed transcripts per gene (2, 3, 4, 5, or all reference transcripts), and the number of biological replicates per group (3, 5, or 10). For each scenario, 20 simulated experiments with RNA-seq libraries from 2 groups were generated.

The baseline relative expression level and the biological variation of selected transcripts were simulated under similar assumptions to the simulation study presented in [14]. Expected counts and associated dispersions were used to generate transcript-level expression following a gamma distribution, which in turn was transformed into transcripts-per-million (TPM) and used as input in *simReads*. The number of reads generated by *simReads* from each transcript varies according to a multinomial distribution with probability determined by the transcript TPM and effective length. These simulation steps ensured that the number of reads arising from each transcript follows a NB distribution across replicates with dispersion equal to the reciprocal of the gamma distribution shape parameter [9]. A random subset of 3,000 transcripts had their baseline relative expression adjusted with a 2-fold-change to establish differential expression between groups with up- and down-regulated transcripts. For every scenario, simulations without any real differential expression between groups (null simulations) were also generated to assess methods’ type I error rate control. See Supplementary Data Section 1.1.2 for more details.

#### Quantification of RNA-seq experiments

Simulated RNA-seq reads were quantified with *Salmon* and *kallisto* with index generated from the complete mouse Gencode transcriptome annotation M27. For *Salmon*, we used a decoy-aware mapping-based indexed transcriptome generated from the mouse mm39 reference genome with k-mers of length 31. The mean and the standard deviation of the fragment length distribution were given as input to *Salmon* and *kallisto* during quantification of single-end read experiments. A total of 100 bootstrap resamples were generated for every sample. To assess the performance of the *sleuth* method with *Salmon* quantification, we transformed *Salmon* output results to *abundance*.*h5* files using the R package *wasabi* (https://github.com/COMBINE-lab/wasabi), which was then used as input for *sleuth*. See Supplementary Data Section 1.1.3 for more details.

#### Assessment of differential transcript expression

We evaluated the performance of *edgeR* with count scaling (*edgeR-Scaled*) and other popular methods with respect to their power to detect DTE, false discovery rate (FDR) control, type I error rate control, and computational speed. Methods benchmarked in our study were *edgeR* with raw counts (*edgeR-Raw*), *sleuth* with likelihood ratio test (*sleuth-LRT*), *sleuth* with Wald test (*sleuth-Wald*), and *Swish* (implemented in the Bioconductor package *fishpond*). For *edgeR-Raw* and *edgeR-Scaled*, low-expression transcripts were filtered by *filterByExpr*, library sizes were normalized by *normLibSizes*, then differential expression was assessed by quasi-likelihood F-tests with default options. For *sleuth* and *Swish*, default filtering and pipeline options implemented in their respective packages were used throughout our simulations. In all analyses, transcripts were considered to be differentially expressed (DE) under an FDR control of 0.05. See Supplementary Data Section 1.1.4 for more details.

### Human lung adenocarcinoma cell lines

Illumina short-read paired-end RNA-seq libraries were obtained from NCBI Gene Expression Omnibus (GEO) series GSE172421. Three biological replicate samples were used to examine the transcriptomic profile of human adenocarcinoma cell lines NCI-H1975 and HCC827. Paired-end reads were quantified with *Salmon* with option *–validateMappings* turned on and using the decoy-aware transcriptome index generated from the human Gencode annotation version 33 and hg38 build of the human genome. A total of 100 bootstraps resamples were generated for every sample. The *edgeR* function *catchSalmon* was used to import *Salmon*’s quantification and estimate the RTA overdispersion parameter.

To explore how RTA depends on read length and read-pairing, we also obtained Oxford Nanopore Technologies (ONT) long-read data from GEO series GSE172421 and Illumina short-read single-end RNA-seq data from GEO series GSE86337 for the same adenocarcinoma cell lines. The short-read single-end libraries were quantified as for the paired-end libraries. The ONT libraries were aligned to the Gencode hg38 transcriptome version 33 using *minimap2* [24] with options *-ax map-ont –sam-hit-only* and secondary alignments excluded, then quantified with *Salmon* in alignment-based mode [25].

For all three technologies, transcript counts were scaled, non-expressed transcripts were filtered by *filterByExpr*, genes other than protein-coding and lncRNA were removed, library sizes were normalized by *normLibSizes* and differential expression was assessed with quasi-likelihood F-tests.

### Variance model for transcript counts

For an RNA-seq experiment consisting of *n* samples and transcriptome annotation containing *T* transcripts, let *y*_*ti*_ denote the fractional number of sequence reads probabilistically assigned to transcript *t* in sample *i*. Let *N*_*i*_ denote the total number of sequenced reads (library size) for sample *i* and *π*_*ti*_ denote the (un-observed) proportion of cDNA fragments originating from transcript *t* in sample *i*. Then, conditional on the true expression level that one would obtain if measuring the transcript expression unambiguously and exhaustively, we have

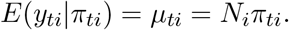

In gene-level expression estimation, it is reasonable to assume that the sampling process of cDNA fragments followed by sequencing, alignment, and counting of RNA-seq reads results in technical Poisson variation of gene-wise read counts over technical replicates [26]. However, at the transcript-level, RTA associated with assignment of reads to overlapping transcripts leads to extra-to-Poisson variation of transcript-wise read counts. We note that the extra-Poisson variation for transcript *i* depends on the degree of overlap that this transcript has with other transcripts, i.e., on annotation topology that is transcript-specific but sample-independent. We also note that, mathematically, the mean and variance of *y*_*ti*_ must both remain directly proportional to *N*_*i*_*π*_*ti*_ for genes that do not show differential transcript usage (i.e., if all transcripts have similar fold-changes between conditions). These considerations lead us to assume the quasi-Poisson variance model

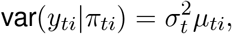

with 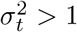. Extensive simulations show that this variance model remains broadly correct even in the presence of differential transcript usage (Supplementary Figure S121). Here 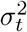 is the Poisson quasi-dispersion, which captures the technical overdispersion produced by RTA for this transcript. We will henceforth call 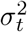 the RTA overdispersion parameter.

We account for the variation of expression levels of each transcript over biological replicates by assuming that the underlying proportions *π*_*ti*_ are random, have mean *π*_0,*ti*_, and have approximately constant coefficient of variation 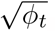, such that var 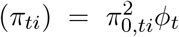. Using the law of total variance, we denote *E*(*y*_*ti*_) = *μ*_*ti*_ = *N*_*i*_*π*_0,*ti*_ and obtain the variance model for transcript counts

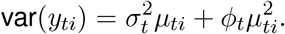

### Count scaling

From an inferential perspective, the proposed variance model is challenging to work with because the purely technical transcript-specific parameter 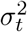 is neither additive with respect to the parameters that can be estimated from replicate samples (*μ*_*ti*_ and *ϕ*_*t*_) nor multiplicative to the variance function. Hence, the RTA overdispersion parameter cannot be directly incorporated into a linear model framework as model weights. Here, we make use of bootstrap samples to estimate the RTA overdispersion parameter 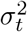 with high accuracy and adopt a count scaling approach to model transcript counts.

For transcript *t*, let *z*_*ti*_ denote the already fractional count *y*_*ti*_ scaled with respect to the RTA overdispersion 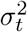, such that 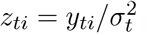. Then, we have that 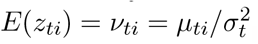, and

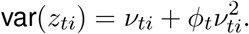

The scaling transformation preserves fold-changes and the variance of the resulting scaled count *z*_*ti*_ is a quadratic and strictly increasing function of the mean *ν*_*ti*_, with the same mean-variance relationship as for a NB model.

### Estimation of RTA overdispersion

The bootstrap resampling process, as performed by lightweight aligners *prior* to the probabilistic assignment of reads to transcripts, takes place at the level of equivalence classes, which by definition are sets of transcripts to which reads map unambiguously, and therefore should result in Poisson variation. We argue that transcript-level bootstrap counts can then be used to quantify the uncertainty associated with the subsequent probabilistic assignment of reads to transcript and estimate the RTA overdispersion parameters. Specifically, for a total of *B* bootstrap samples, any extra variability observed over bootstrap counts *u*_*ti*1_, …, *u*_*tiB*_ from transcript *t* and sample *i* must be due to the quantification uncertainty.

Under the quasi-Poisson model, we consider the Pearson residual statistic and propose the following moment estimator for the RTA overdispersion parameter

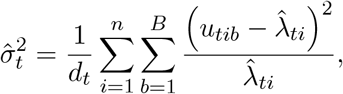

with *d*_*t*_ = *n*(*B −* 1) and 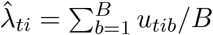.

We propose an empirical Bayes approach to moderate the 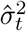 estimates. Specifically, let 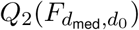 denote the median value of an *F* distribution with *d*_med_ and *d*_0_ degrees of freedom, with *d*_med_ denoting the observed median degree of freedom *d*_*t*_ from expressed transcripts, and *d*_0_ = 3 denoting a prior degree of freedom. In addition, let 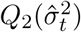 denote the observed median RTA overdispersion estimate of expressed transcripts. We assume a prior RTA overdispersion 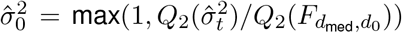 shared among all transcripts. Then, the transcript-specific empirical Bayes moderated estimator of the RTA overdispersion can be written as

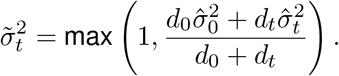

We note that the level of shrinkage applied on 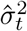 towards our proposed moderated statistic 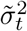 is minor, since most transcripts are expressed to a certain degree in most RNA-seq samples and the degrees of freedom *d*_*t*_ associated with the estimator 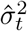 is often 2 to 3 orders of magnitude larger than *d*_0_.

### Usage and implementation

The degrees of freedom *d*_0_ +*d*_*t*_ used to estimate 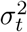 are typically large, for example, *d*_*t*_ = 990 for a transcript expressed in 10 RNA-seq samples with 100 bootstrap samples each. The estimator 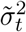 is therefore very precise and can be considered to be a known constant for most purposes. Our proposed method therefore is simply to scale the counts using the estimated dispersions, 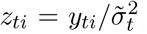, and to input the scaled counts to a standard differential expression pipeline designed for NB distributed counts such as *edgeR* or *limmavoom* [14]. *edgeR* implements a continuous generalization of the NB distribution, and *limma-voom* accepts continuous data, so that the scaled counts do not need to be rounded to integers.

The calculation of the empirical Bayes moderated RTA overdispersion statistic 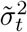 is implemented in the *edgeR* functions *catchSalmon* and *catchKallisto*. The functions return the matrix of transcript counts together with the associated RTA overdispersions.

## RESULTS

### RTA overdispersion increases with transcript overlap

We first explored RTA overdispersions for the paired-end lung adenocarcinoma cell line data. We observed a strong increasing trend with the number of transcripts per gene (Figure 1). For single-transcript genes, only 11% of their 813 expressed transcripts had overdispersion greater than 1.11, whereas for multi-transcript genes, this was so for 90% of their 26,553 expressed transcripts. For comparative purposes, we also estimated overdispersions from ONT long-read libraries of the same human adenocarcinoma cell lines. For long-reads, the overdispersions were close to zero regardless of the number of transcripts per gene (Figure 1), showing that sufficiently long reads can be assigned to transcripts uniquely, thus eliminating any ambiguity. We also confirmed that overdispersion is minimal for read counts summarized at the gene level (Supplementary Figure S122 and Table S11). Count scaling can be interpreted as reducing the effective number of reads for that transcript. For expressed genes with 10 or more annotated transcripts (4,687 genes), the average overdispersion of their expressed transcripts is 6.75, so the counts will be reduced nearly 7-fold to reflect the precision associated with their expression estimates. For such transcripts, one would need a 7 times higher sequencing depth for a transcript-level analysis with short-reads to achieve the same statistical power as for the corresponding gene-level analysis.

**Figure 1:**
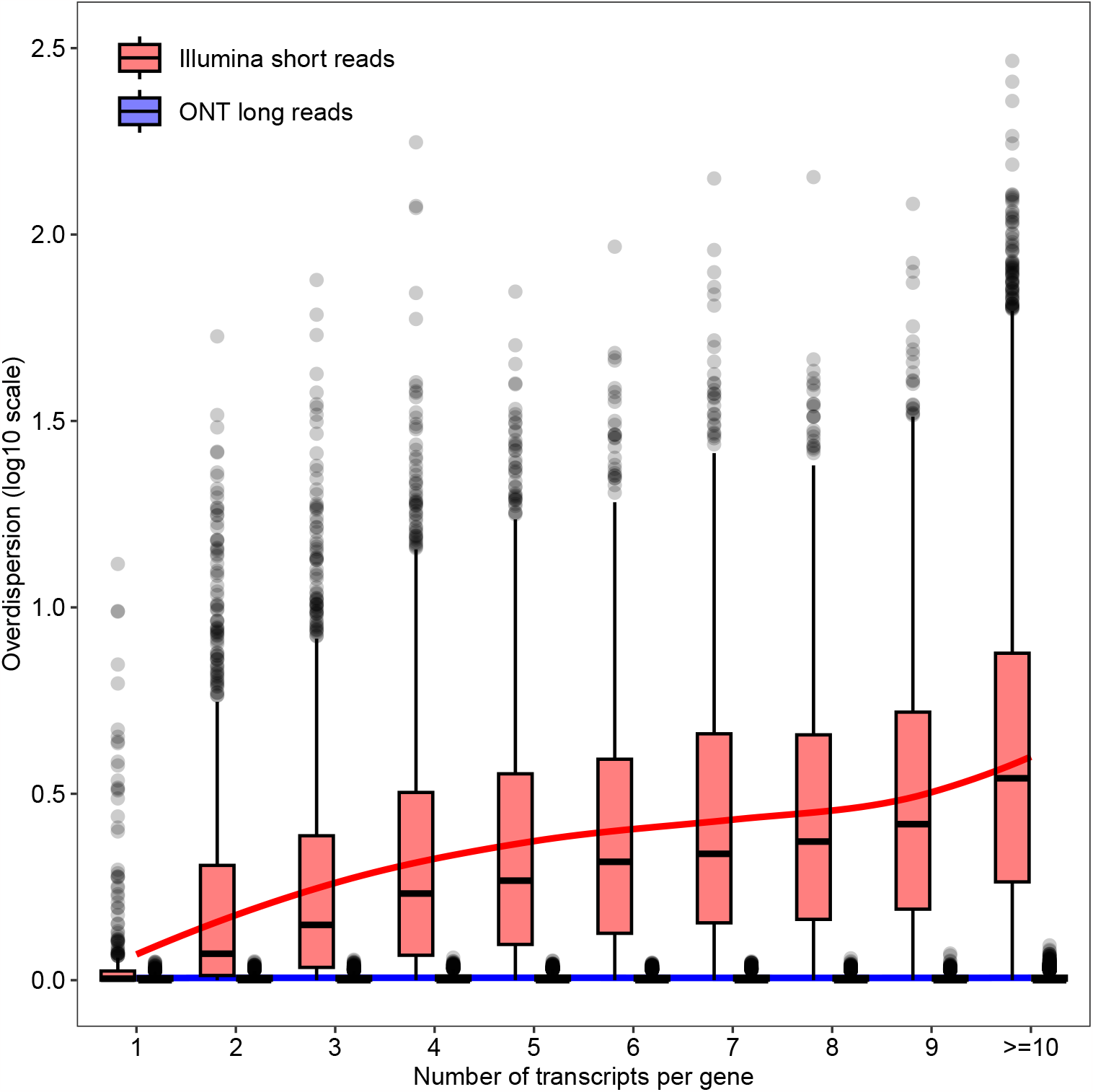
Transcript-level RTA overdispersion estimates from the lung adenocarcinoma cell line data with Illumina paired-end short-reads or ONT long-reads. Boxplots show overdispersion estimates (log10 scale) for expressed transcripts by the number of annotated transcripts for the corresponding gene. The red and blue lines show lowess trends for short-reads and long-reads respectively.

### Count scaling results in effective counts that better reflect their true precision

The effect of the count scaling approach can be visually appreciated via BCV plots. The square-root of the NB dispersion *ϕ*_*t*_ represents the coefficient of variation of the true expression values between biological replicates, termed the biological coefficient of variation (BCV) [9]. The BCV plot displays estimated BCVs against transcript abundances in log2 counts per million.

For RNA-seq data summarized at the gene-level, NB dispersions tend to be higher for genes with low counts with a trend that decreases smoothly with abundance while asymptoting to a constant value for genes with large counts [11]. At the transcript-level, the RTA is highly transcript-specific, and estimated dispersions based on raw counts may be artificially higher for transcripts associated with high RTA. As a result, the estimated dispersions of such ambiguous transcripts may deviate from the standard decreasing smooth dispersion trend. Under the presented quasi-Poisson model, effective counts computed via count scaling should lead to estimated dispersions and BCV plots that are free of RTA effects and, in principle, should resemble those from gene-level analyses.

Figure 2 presents BCV plots from the lung adenocarcinoma cell lines dataset, as well as from one of our mouse simulated datasets, generated before and after count scaling. BCV plots were generated with the *edgeR* function *plotBCV*. For both real and simulated datasets, the BCV plots computed with raw counts exhibit strong transcript-specific RTA effects, with artificially high estimated NB dispersions for ambiguous transcripts. Upon count scaling, effective counts better reflect their true precision with estimated dispersions that are truly representative of the biological variation of the RNA-seq experiment. Most importantly, BCV plots from scaled counts suggest that standard DE methods designed for NB distributed counts, such as *edgeR* and *limma-voom*, can be directly applied.

**Figure 2:**
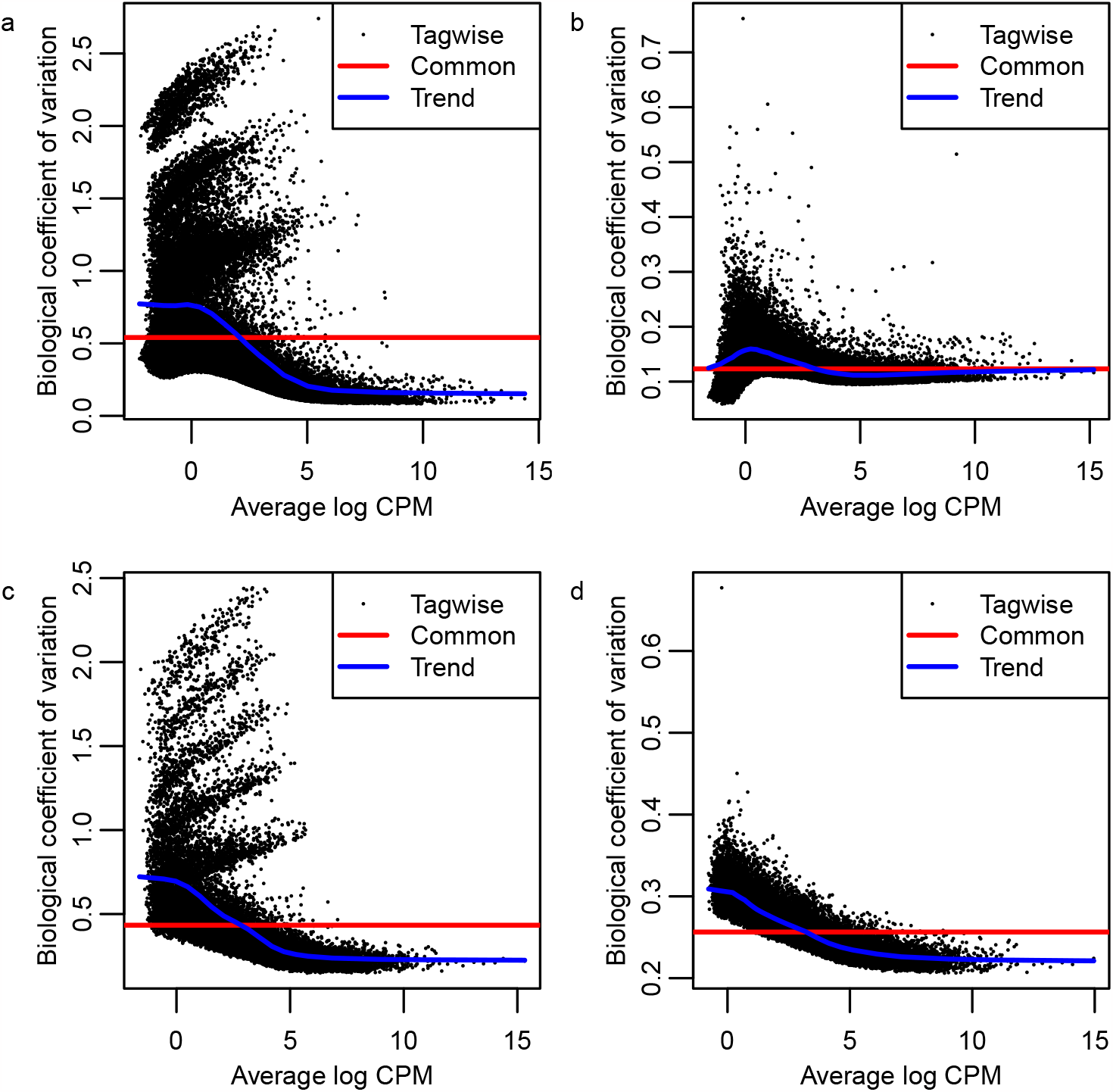
BCV plots of transcript-level counts from RNA-seq data. Panels (a) and (b) show results from the Illumina short paired-end reads RNA-seq experiment of the lung adenocarcinoma cell lines before and after counts scaling, respectively. Panels (c) and (d) show results from a simulated mouse RNA-seq experiment before and after counts scaling, respectively. Simulated data was generated with 100bp paired-end reads, unbalanced library sizes, 5 samples per group, and all reference transcripts expressed. For both real and simulated data, experiments were quantified with *Salmon*.

### Count scaling provides powerful and efficient differential transcript expression analyses

We assessed the performance of methods with respect to power to detect DTE between groups and ability to control the FDR. Figure 3 shows the observed number of true positive and false positive DE transcripts for all benchmarked methods under a nominal FDR control of 0.05, for 100 bp paired-end read simulations. We observed that *edgeR* with count scaling was able to detect the largest number of DE transcripts among all methods while controlling the FDR under the nominal value, regardless of the number of replicate samples per groups and library size. Under scenarios with 3 samples per group, *edgeR* with raw counts and *sleuth-LRT* provided only minimal power to detect DTE. In contrast, *Swish* exhibited reasonable power to detect DE transcripts in such scenarios but at the expense of an increased observed FDR. All methods showed a substantially increase in power to detect DE transcripts as more samples per group were considered. In scenarios with 10 samples per group, *Swish* outperformed both *sleuth-LRT* and *sleuth-Wald* in terms of statistical power (Supplementary Tables S1–S10). Yet, *edgeR* with count scaling still ranked as the most powerful DTE method in such scenarios.

**Figure 3:**
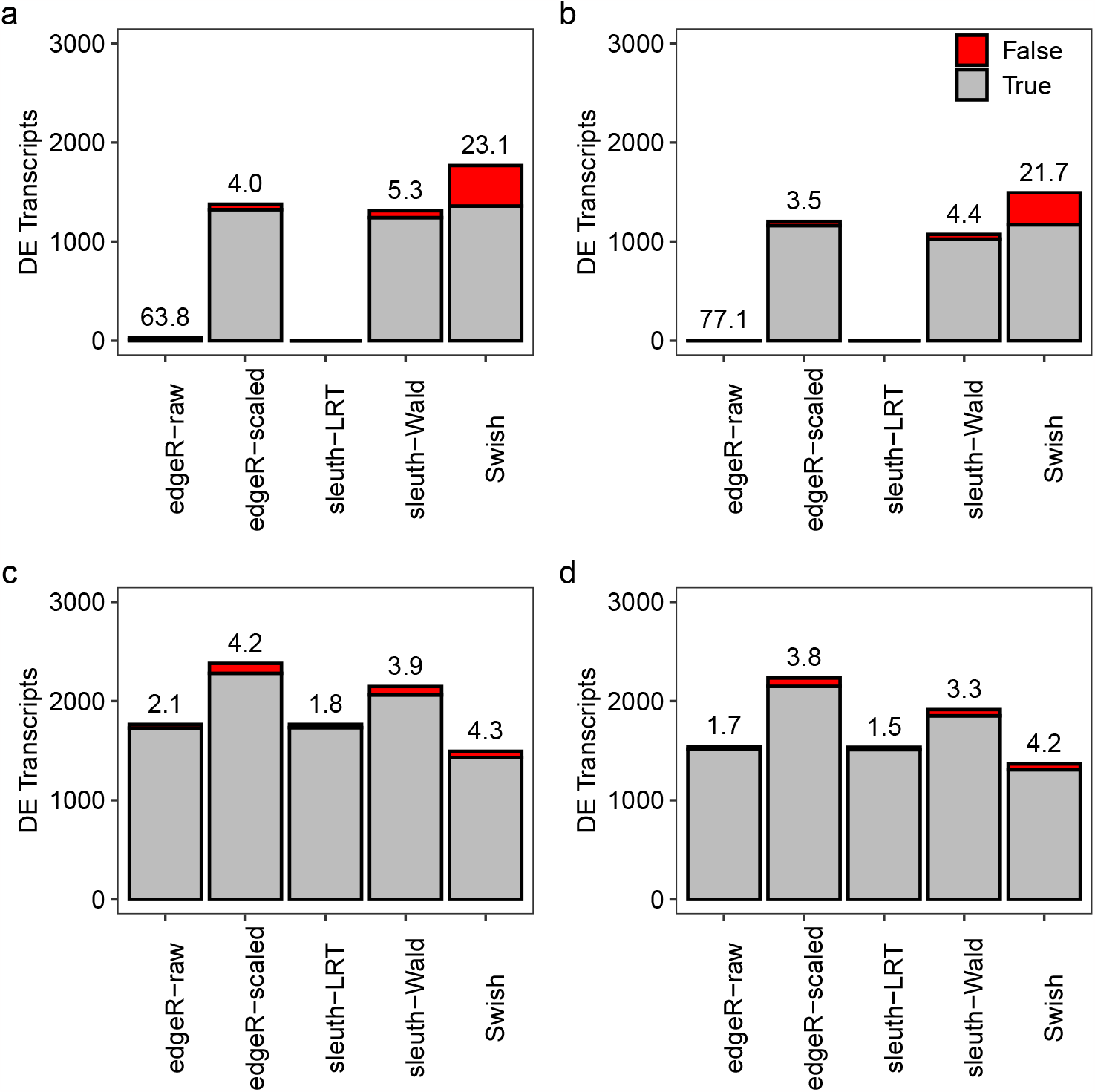
Stacked barplots showing the average number of true (gray) and false (red) positive DE transcripts at nominal 5% FDR in different simulation scenarios. The observed FDR is shown as a percentage over each bar. Panels (a) and (c) show results with balanced library sizes. Panels (b) and (d) show results with unbalanced library sizes. Panels (a) and (b) show results with 3 samples per group. Panels (c) and (d) show results with 5 samples per group. Results are averaged over 20 simulations with 100 bp paired-end read data quantified with Salmon with all reference transcripts expressed.

To further compare methods regarding FDR control, we assessed the number of false discoveries in the set of top-ranked most significant transcripts from each method (Figure 4). Overall, *edgeR* with count scaling provided the smallest number of false discoveries among all methods for any number of top-ranked transcripts. For all configurations of library sizes and number of samples per group, *Swish* consistently presented more false positive transcripts than any other method for any given number of top-ranked transcripts. Yet, with paired-end read experiments, we note that all methods successfully controlled the FDR under the nominal level in scenarios with 5 or 10 samples per group. In scenarios with single-end read experiments quantified with *kallisto* and high number of samples per group, we note that both *edgeR* with count scaling and *Swish*, the two most powerful DTE methods, presented FDR levels slightly over the nominal level 0.05, with values that ranged between 0.05 and 0.08, on average. We have not observed the same lack of FDR control in single-end experiments quantified with *Salmon* in equivalent scenarios (Supplementary Tables S1–S10).

**Figure 4:**
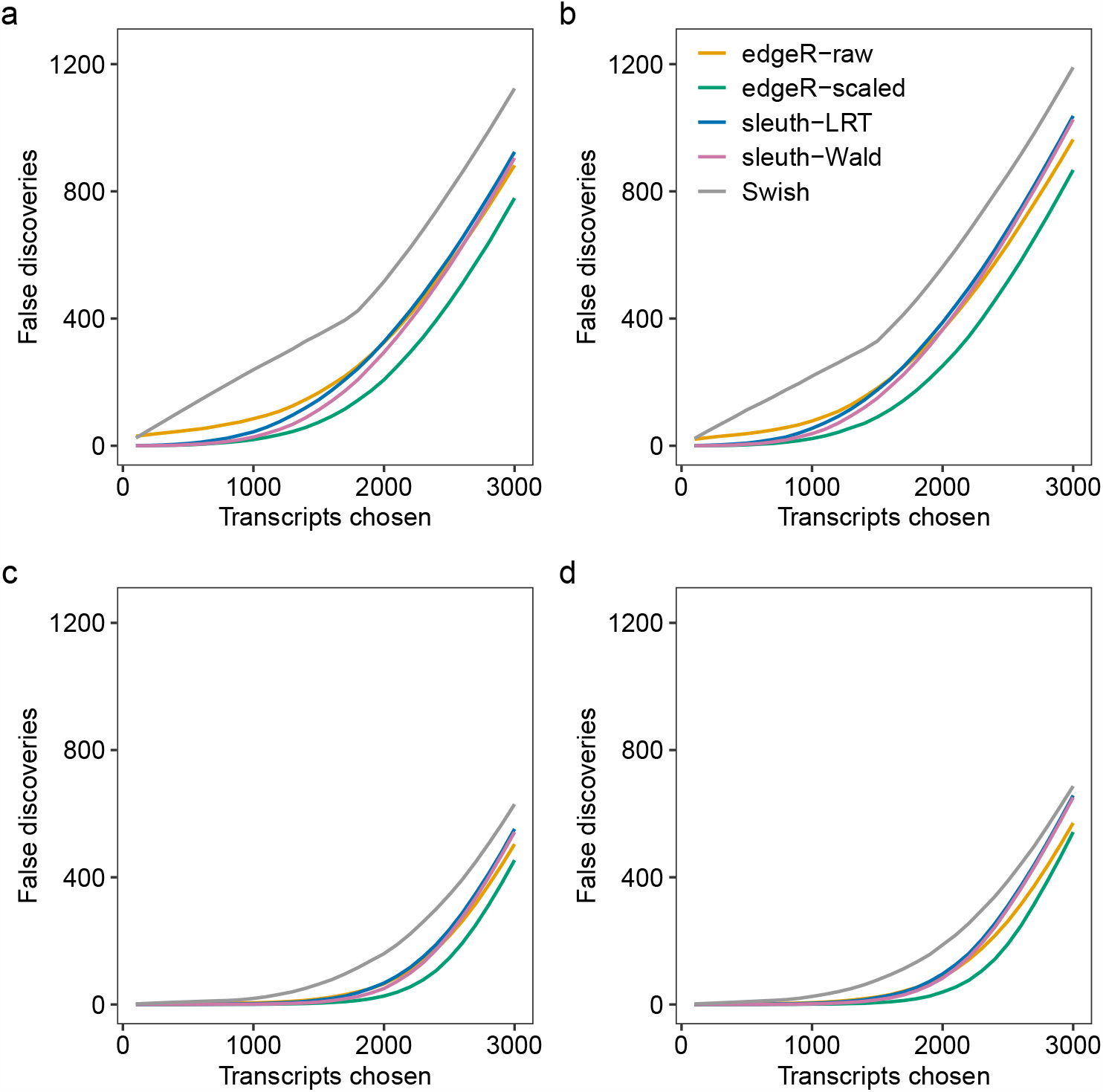
Panels (a)–(d) show the average number of false discoveries as a function of the number of chosen transcripts in different simulation scenarios. In (a) and (c), scenario with balanced library sizes. In (b) and (d), scenario with unbalanced library sizes. In (a) and (b), scenario with 3 samples per group. In (c) and (d), scenario with 5 samples per group. Results from the simulations with 100 bp paired-end read data quantified with Salmon with all reference transcripts expressed, averaged over 20 simulations.

For all benchmarked methods, we observed that experiments simulated with paired-end reads led to uniformly more powerful DTE analyses than single-end read experiments for any read length specification. Overall, methods exhibited an increase in power to detect DE transcripts as more transcripts from the reference set were left unexpressed. Starting from the quantification output of either *Salmon* or *kallisto, edgeR* with count scaling was the fastest method in comparison, while performing the entire DTE analysis pipeline of 10 sample RNA-seq experiments, with 100 bootstrap resamples each, in approximately 15 seconds, on average (Supplementary Figures S1–S40).

### Count scaling controls the type I error rate

We evaluated the ability of benchmarked methods to control the type I error rate in null simulations that were generated without any differential expression between groups. Under the hypothesis of no differential expression between groups, p-values are expected to be uniformly distributed between the 0 and 1. Figure 5 shows the proportion of significant p-values under a nominal significance level of 0.05 in various simulation scenarios. We observed that all methods exhibited control of the type I error rate with the proportion of false positive calls over all transcripts being below or near the nominal level.

**Figure 5:**
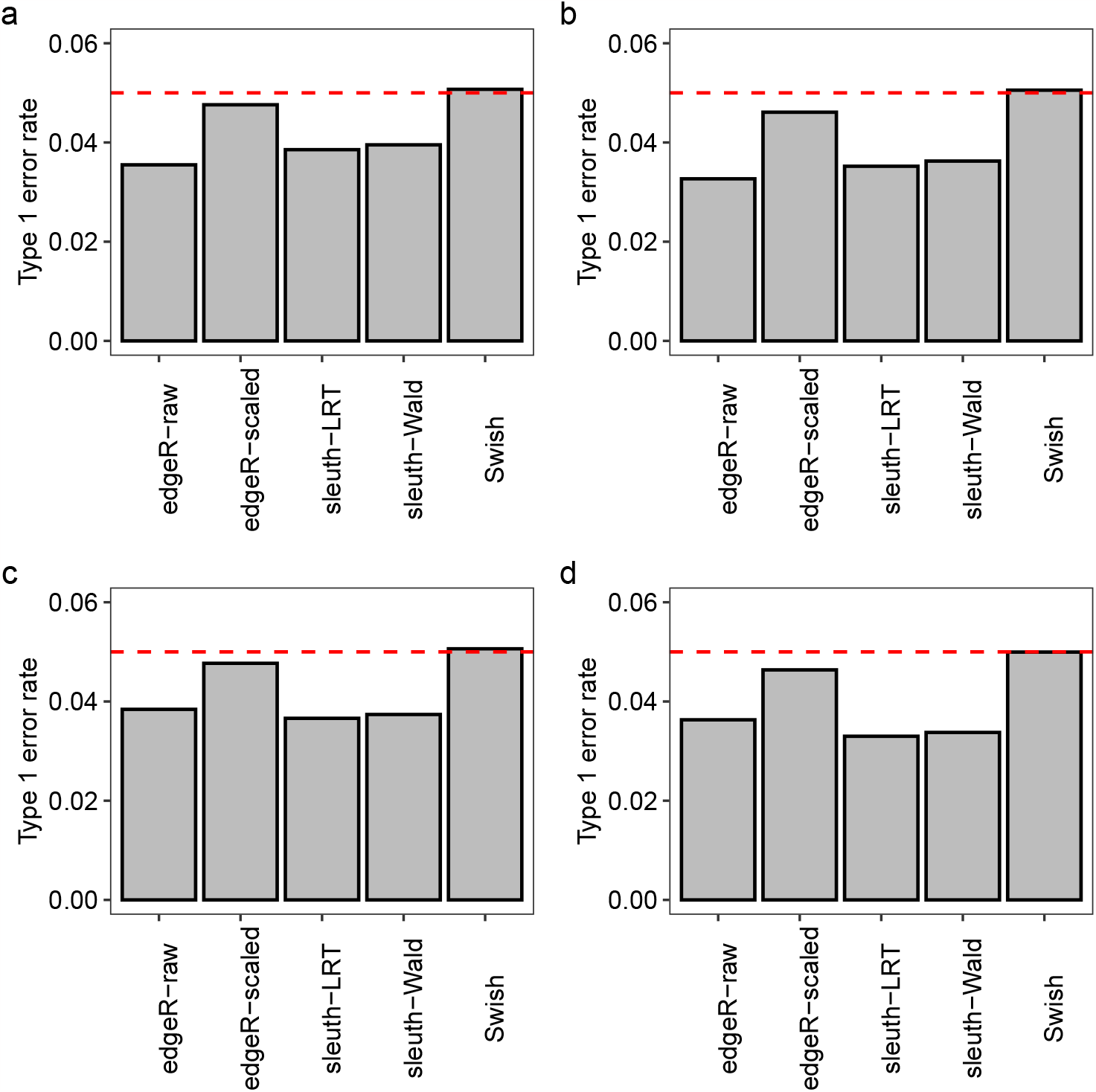
Panels (a)–(d) show the observed type 1 error rate calculated as the average proportion of transcripts with unadjusted p-values *<* 0.05 in different null simulation scenarios (without differential expression). Dashed line indicates the expected proportion of p-values *<* 0.05 under the null hypothesis of no differential expression. In (a) and (c), scenario with balanced library sizes. In (b) and (d), scenario with unbalanced library sizes. In (a) and (b), scenario with 3 samples per group. In (c) and (d), scenario with 5 samples per group. Results from the null simulations with 100 bp paired-end read data quantified with Salmon with all reference transcripts expressed, averaged over 20 simulations.

Among all methods and simulation scenarios, *Swish* exhibited the most uniform distribution of p-values with its observed type I error rate being the closest to the nominal level. Figure 6 presents density histograms of raw p-values from all methods for the null simulation scenario with unbalanced library sizes and 5 samples per group. Overall, *sleuth-LRT* and *sleuth-Wald* presented p-value distributions that were substantially skewed towards 1, on average. In comparison to raw counts, the presented count scaling approach for transcript-level analyses with *edgeR* led to p-value distributions that were approximately uniform throughout our simulations. Similar results were found in simulations generated with single-end reads, different read lengths, and fewer number of expressed transcripts per gene (Supplementary Figures S41–S120).

**Figure 6:**
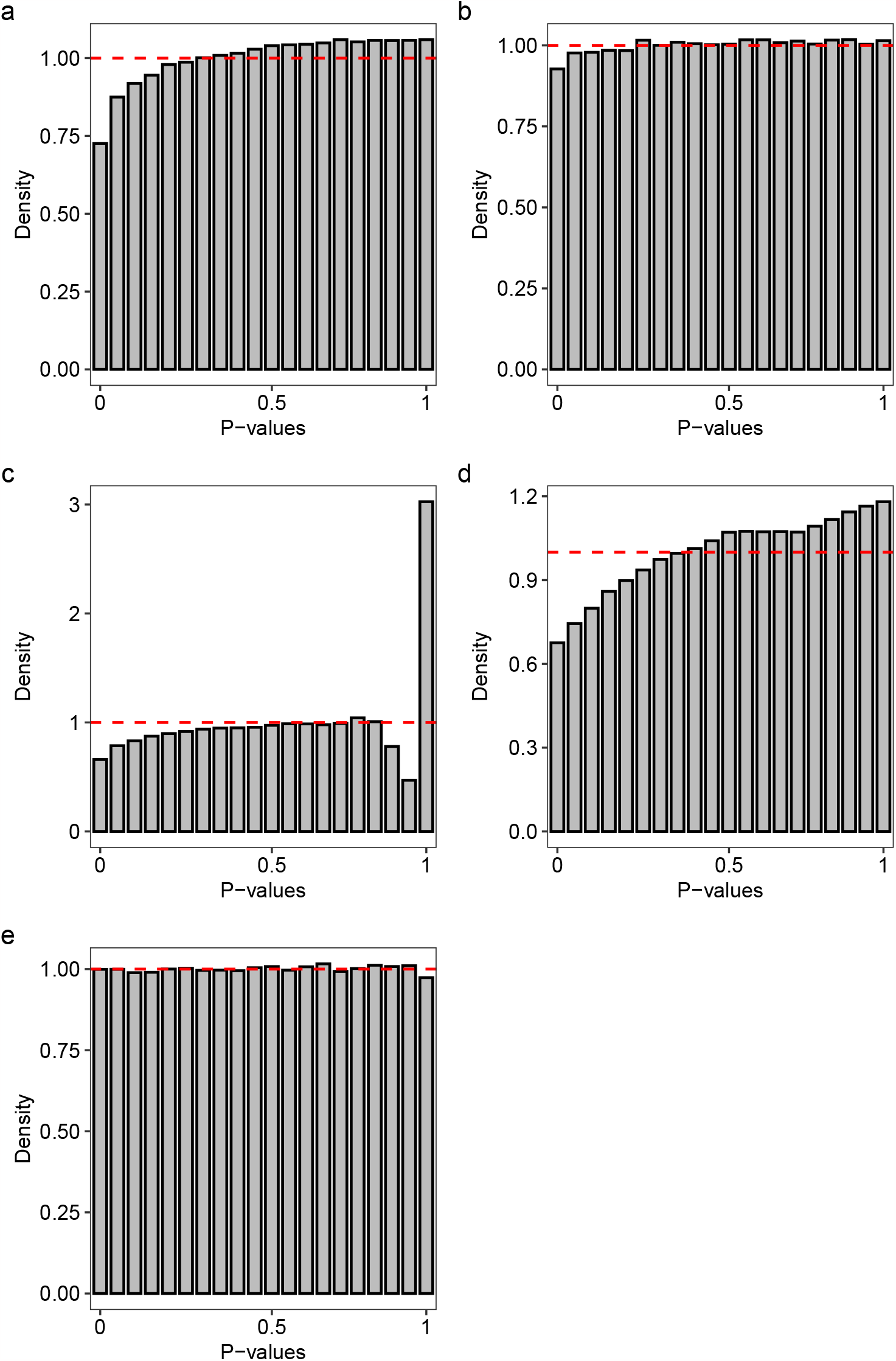
Panels (a)–(e) show density histograms of raw p-values from various methods in the null simulation scenario (without differential expression) with unbalanced library sizes and 5 samples per group. Dashed line indicates the expected distribution of p-values under the null hypothesis of no differential expression. In (a), results from *edgeR* with raw counts. In (b), results from *edgeR* with scaled counts. In (c), results from *sleuth* with LRT. In (d), results from *sleuth* with Wald test. In (e), results from *Swish*. Results from the null simulations with 100 bp paired-end read data quantified with Salmon with all reference transcripts expressed, averaged over 20 simulations.

### Longer paired-end reads decrease RTA and provide more powerful DTE analyses

In simulated RNA-seq experiments, we assessed the extent to which the sequence read specification influenced the RTA overdispersion of transcripts. To this end, we compared the average estimated RTA overdispersion parameter in single- and paired-end read RNA-seq experiments simulated under a range of different read lengths. We further evaluated the resulting power and FDR of *edgeR* with count scaling when detecting DE transcripts. Table 1 presents results from our analysis. Overall, we observed that longer paired-end read RNA-seq experiments led to lower RTA overdispersion. Experiments generated with 150 bp paired-end read data had an average RTA overdispersion nearly 40% smaller than 50 bp single-end read experiments (0.205 log10 fold-change). When comparing paired-end to single-end reads, we observed a decrease in RTA overdispersion that varied between 32% for 50 bp reads (0.205 and 0.038 log10 fold-changes) and 12% for 150 bp reads (0.057 log10 fold-change). Such an increase translates to a substantial loss of information in the analysis of single-end read RNA-seq experiments at the transcript-level, which is a result of the reduced precision associated with the estimation of transcript expression with single-end sequence reads. In DTE analyses, we observed a corresponding reduction in statistical power that varied between 5% and 2% for shorter and longer read lengths, respectively, with single-end read data.

**Table 1:** Fold-change of average RTA overdispersion estimates (log10-scale, log-FC), observed power and FDR from *edgeR* with count scaling for RNA-seq experiments simulated under different number of samples per group and sequence read specifications. For a given number of samples per group, 150 bp paired-end reads are used as reference when computing fold-changes. Results from the simulation scenario with unbalanced library sizes and quantified with Salmon with all reference transcripts expressed, averaged over 20 simulations.

| Samples per group | Read Length (bp) | Single-end Read |  |  | Paired-end Read |  |  |
| --- | --- | --- | --- | --- | --- | --- | --- |
|  |  | Mapping Ambiguity log-FC | Power | FDR | Mapping Ambiguity log-FC | Power | FDR |
| Three | 50 | 0.204 | 0.353 | 0.034 | 0.039 | 0.378 | 0.035 |
|  | 75 | 0.160 | 0.357 | 0.033 | 0.018 | 0.389 | 0.035 |
|  | 100 | 0.125 | 0.369 | 0.033 | 0.006 | 0.387 | 0.035 |
|  | 125 | 0.089 | 0.381 | 0.037 | 0.000 | 0.397 | 0.034 |
|  | 150 | 0.057 | 0.386 | 0.035 | 0.000 | 0.395 | 0.035 |
| Five | 50 | 0.205 | 0.660 | 0.038 | 0.038 | 0.708 | 0.039 |
|  | 75 | 0.160 | 0.670 | 0.036 | 0.019 | 0.719 | 0.039 |
|  | 100 | 0.126 | 0.680 | 0.038 | 0.007 | 0.716 | 0.038 |
|  | 125 | 0.089 | 0.699 | 0.040 | 0.000 | 0.726 | 0.039 |
|  | 150 | 0.057 | 0.707 | 0.039 | 0.000 | 0.725 | 0.037 |

In practice, we expect the performance gain for paired-end over single-end read data to be slightly greater than that shown in Table 1. In our simulations, the fragment length parameters for *Salmon* and *kallisto* were set to the true mean and standard deviation of the simulated fragment lengths. However, in practice, the fragment length distribution cannot be determined from single-end RNA-seq reads. To explore the sensitivity of the results to the fragment length settings, we repeated the quantification of the single-end read experiments with the fragment length mean and standard deviation set to the *Salmon* default values of 250 and 25, respectively. While preserving the same relative ranking of DTE methods, leaving the default values for the mean and standard deviation of the fragment length distribution in *Salmon* resulted in a slight decrease in performance for all methods, with a reduction of statistical power that varied between 1% and 3% for sequence reads of length 150bp and 50bp, respectively.

### Differential transcript expression in human adenocarcinoma cell lines

The transcriptomic profile of human lung cancers has been extensively discussed in the literature. In lung adenocarcinoma, it has been observed the existence of differential transcript expression for certain genes, such as the *KRAS* and the *CD274* genes for which the expression levels of a number of their splicing isoforms appeared to be associated with disease initiation and progression [27, 28]. Here, we performed a transcript-level analysis of the Illumina short paired-end read RNA-seq experiments from the human adenocarcinoma cell lines, which comprises 6 samples of NCI-H1975 and HCC827 cell lines with 3 biological replicate samples per cell line (Figure 7a). Libraries were sequenced with an Illumina NextSeq 500 sequencing system, producing 28–134 million 80 bp read-pairs per sample.

**Figure 7:**
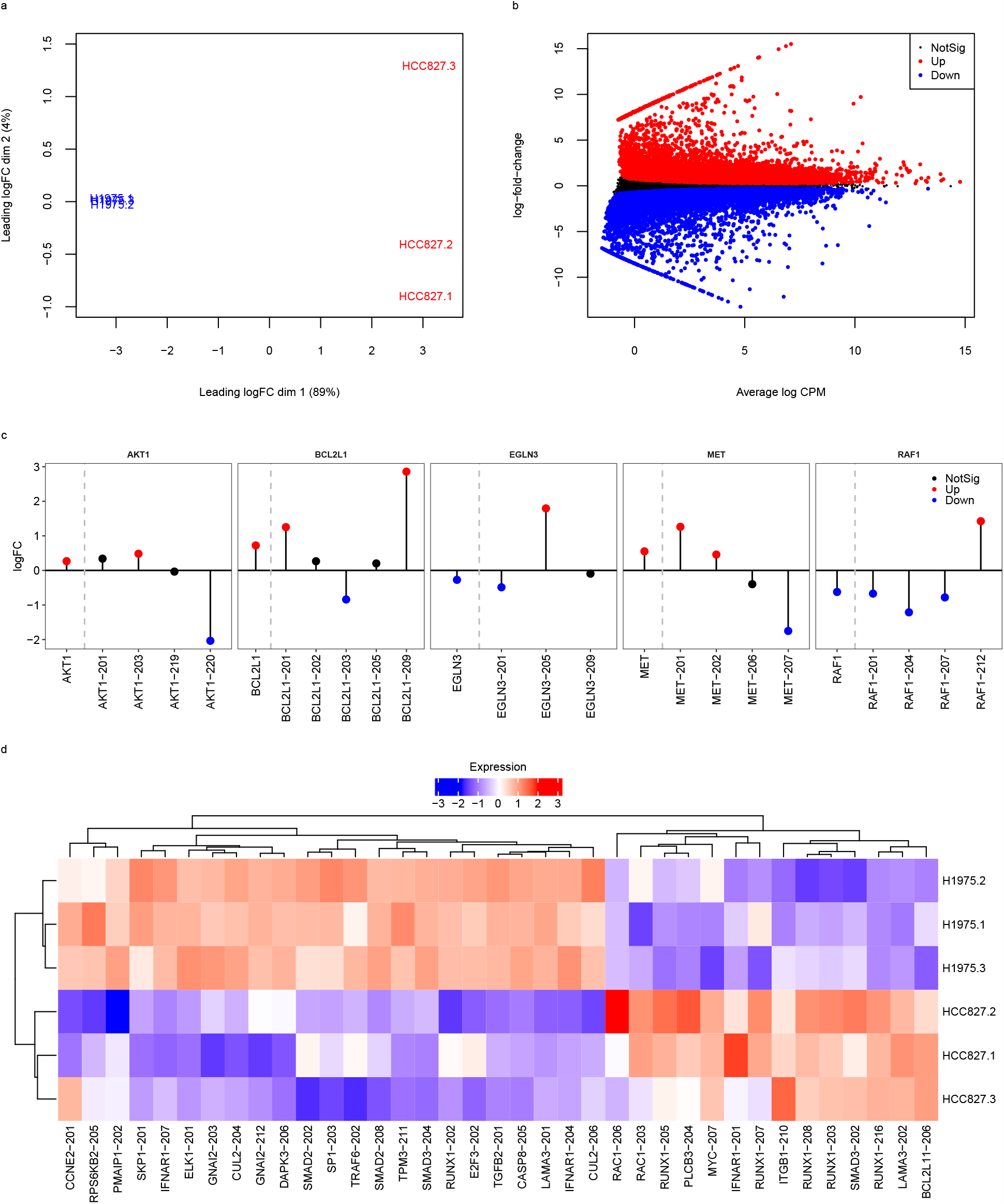
Panels (a)–(d) show the main results from the RNA-seq DTE analysis of the human lung adenocarcinoma cell lines (Illumina short-read paired-end data). In (a), multidimensional scaling plot of NCI-H1975 and HCC827 samples. In (b), mean-difference plot highlighting differentially expressed transcripts between NCI-H1975 and HCC827 cell lines. In (c), log-fold-change plot of a set of cancer-related DE genes (on the left of dashed lines) and their associated expressed transcripts (on the right of dashed lines). Genes and transcripts are highlighted in red, if differentially up-regulated, in blue, if differentially down-regulated, and in black, if non-significant. In (d), heatmap of DE transcripts between NCI-H1975 and HCC827 cell lines from non-significant genes associated with KEGG cancer and non-small cell lung cancer pathways. Scaled log2 counts per million are displayed as expression levels in the heatmap. Nominal FDR of 0.05 in both gene- and transcript-level analyses.

We applied *edgeR* with the presented count scaling approach and quasi-likelihood F-tests and detected a total of 18,699 DE transcripts between NCI-H1975 and HCC827 cell lines (9,079 up-regulated and 9,620 down-regulated transcripts in the HCC827 cell line; Figure 7b). Of particular interest was the detection of DE transcripts of the *KRAS* and *CD274* genes, namely the protein-coding transcripts *KRAS-201* (1.17 log fold-change, adjusted p-value 6.685 × 10^−4^) and *CD274-202* (-0.490 log fold-change, adjusted p-value 5.485 × 10^−3^). In addition, we performed a gene-level analysis of the same RNA-seq experiment and identified 403 DE genes between cell lines for which at least one of their transcripts was also DE but in the opposite direction (nominal FDR at 0.05 in both gene- and transcript-level analyses). This set of DE genes with contrasting transcript expression signatures included a total of 12 genes associated with the KEGG cancer and non-small cancer pathways, such as genes *AKT1, BCL2L1, EGLN3, MET*, and *RAF1* that have been extensively discussed and considered as potential therapy targets in lung carcinomas (Figure 7c, Supplementary Table S12) [29, 30, 31].

Our gene-level analysis also revealed a total of 841 non-significant DE genes for which at least one of their transcripts was significantly expressed between cell lines. Out of such non-significant genes, we have identified 24 genes associated with the KEGG cancer and non-small cell lung cancer pathways (Figure 7d). Such a list includes established prognostic markers for several types of cancers such as the proto-oncogene *MYC* and the pro-apoptotic gene *PMAIP1* [32]. Our analysis also revealed multiple DE transcripts of the *RUNX1* gene between cell lines, which its down-regulation has been associated with aggressive lung adenocarcinomas [33]. Moreover, we observed a case of isoform switching expression for gene *RUNX1* with a protein-coding transcript being expressed in the opposite direction between cell lines in contrast to its competing transcripts (*RUNX1-202*; Figure 7d). Other non-significant cancer-associated genes with significant isoform switch expression include *IFNAR1, SMAD3*, and *LAMA3*. The discovery of differential expression of transcripts associated with non-significant, albeit important cancer-related, genes between NCI-H1975 and HCC827 cell lines highlights the potential benefits of an analysis of RNA-seq data at the transcript-level.

Using ONT long-read data from the same RNA-seq experiment, we applied the standard *edgeR* pipeline at the transcript-level with quasi-likelihood F-tests on raw counts and detected a total of 27,817 DE transcripts between NCI-H1975 and HCC827 cell lines (14,146 up-regulated and 13,671 down-regulated transcripts in the HCC827 cell line). Given the almost negligible RTA associated with the quantification of ONT long-reads (Figure 1), the total number of DE transcripts found between cell lines using long-reads may serve as a benchmarking target for short-read DTE analyses. In fact, using Illumina short paired-end reads, *edgeR* with count scaling detected 42.3% of all DE transcripts found with long-reads, a percentage larger than that of *sleuth-LRT* (38.8%), *sleuth-Wald* (36.7%), and *Swish* (28.2%). Finally, to assess the benefits of performing a DTE analysis with paired-end over single-end read data, we performed an analysis at the transcript-level using single-end read libraries from the same cell lines (GSE86337). Using *edgeR* with count scaling, we found a total of 10,468 DE transcripts between NCI-H1975 and HCC827 cell lines (5,067 up-regulated and 5,401 down-regulated transcripts in HCC827), which suggests a slightly under powered analysis in comparison to paired-end read data (Supplementary Figures S123 and S124). Such results highlight the benefits of performing transcript-level analyses with paired-end over single-end read RNA-seq data and agree with the findings presented in our simulation study.

## DISCUSSION

Here, we present a simple, powerful and effective approach to account for the RTA overdispersion resulting from transcript quantification in differential analysis of RNA-seq data at the transcript-level. Our comprehensive simulation study demonstrates that the presented count scaling approach provides uniformly more powerful DTE analyses than current methods, while harnessing the flexible generalized linear model framework and the efficient implementation of statistical methods available in the *edgeR* Bioconductor package. We show that *edgeR* with count scaling properly controls the FDR in all evaluated scenarios, including those with small number of replicates, short single-end read data, and highly unbalanced library sizes. In null simulations, we further show that the presented approach provides proper type I error rate control. Our case study of the human adenocarcinoma cell lines uncovered several DE transcripts, including transcripts associated to key cancer-related genes that did not appear to be differentially expressed between cell lines in a gene-level analysis. *edgeR* implements a continuous generalization of the NB distribution, so the scaled counts can be used directly without rounding to integers, meaning that no information is lost for low counts.

Recommendations for transcript-level analyses of RNA-seq data are also presented. In contrast to gene-level analyses, for which single-end data may be sufficient, our simulation study shows that paired-end sequence reads lead to uniformly better power to assess DTE. When designing RNA-seq experiments for which the analysis is intended to be carried out at the transcript-level, we recommend paired-end sequence read libraries with 50 bp or greater read length and with at least 50 million read-pairs per sample. Our *edgeR-Scaled* method works for any number of replicates, but the improvement in statistical power from 3 replicates per group to 5 replicates per group was notable in our simulations.

We note that a single dispersion and fold-change setting was used for all sample size scenarios in order to emphasize the increase in power delivered by larger sample sizes. In practice, real datasets with large sample sizes are often observational human studies with a high level of biological variability. In addition, researchers might choose to generate fewer bootstrap resamples per sample in large sample size situations than in our simulation. We have not explored these possible settings but such factors might produce lower statistical power than in our simulation. Our simulations were set to be typical of designed experiments with model organisms such as genetically identical mice or cell lines. Studies with more variable units such as human subjects may exhibit higher BCVs and require large sample sizes.

Our simulations assumed the true transcript expression levels to be gamma distributed between replicates. To confirm that our conclusions are not sensitive to this distributional assumption, we repeated the simulations with log-normal distributed expression values instead of gamma and obtained similar results (data not shown).

Finally, we note that the accuracy of the RTA overdispersion estimates, as well as the accuracy of the estimated transcript-specific read counts as output by quantification tools *Salmon* and *kallisto*, heavily depends on the completeness assumption of the transcriptome annotation used during the RNA-seq quantification. The extent to which the presence of novel un-annotated transcripts in the sample affects the quantification of transcripts and, in turn, the estimation of the RTA overdispersion has not been explored in this work.

## CODE AVAILABILITY

The *catchSalmon* and *catchKallisto* functions are available in the *edgeR* Bioconductor [34] package at https://bioconductor.org/packages/edgeR. Both functions implement the methodology presented in this article and estimate the transcript-specific RTA overdispersion resulting from the transcript-level RNA-seq quantification step. When performing DTE analyses with *edgeR* with count scaling, users should divide transcript-level RNA-seq counts by the associated RTA overdispersion estimates. Data and code to reproduce the results presented in this article are available at https://github.com/plbaldoni/TranscriptDE-code.

The versions of software used in the paper are: *ComplexHeatmap*: 2.14.0 [35], *edgeR*: 3.40.2, *fishpond* (*Swish* method): 2.4.1, *kallisto*: 0.46.1, *minimap2*: 2.17, *R*: 4.2.1, *Rsubread* : 2.12.0, *sleuth*: 0.30.0, *Salmon*: 1.9.0, *wasabi* : 1.0.1, *tximeta*: 1.16.1.

## DATA AVAILABILITY

The RNA-seq experiments analyzed here are available from the NCBI Gene Expression Omnibus with the accession numbers GSE60450, GSE86337, and GSE172421.

## SUPPLEMENTARY DATA

Supplementary Data are available in the file supp.pdf.

## ACKNOWLEDGEMENTS

This work was supported by the Chan Zuckerberg Initiative (EOSS4 grant number 2021-237445), Australian National Health and Medical Research Council (NHMRC) IRIISS and Victorian State Government Operational Infrastructure Support. G.K.S. was supported by NHMRC Fellowship 1058892, Y.C. by Medical Research Future Fund Investigator Grant 1176199 and M.E.R. by NHMRC Investigator Grant 2017257.

