## Supplementary Material for "Dividing out quantification uncertainty allows efficient assessment of differential transcript expression with edgeR"

14 October 2023

**4 References** **153**

[1]Bioinformatics Division, WEHI, Parkville, VIC 3052, Australia
[2]ACRF Cancer Biology and Stem Cells Division, WEHI, Parkville, VIC 3052, Australia
[3]Epigenetics and Development Division, WEHI, Parkville, VIC 3052, Australia
[4]Olivia Newton-John Cancer Research Institute, Heidelberg, VIC 3084, Australia
[5]Department of Medical Biology, The University of Melbourne, Parkville, VIC 3010, Australia
[6]School of Mathematics and Statistics, The University of Melbourne, Parkville, VIC 3010, Australia
[7]School of Cancer Medicine, La Trobe University, Melbourne, VIC 3086, Australia
[†]These authors contributed equally to this work

### 1 Simulated datasets

In this section, we describe the creation of the simulated experiments and present the complete set of results from our simulation study.

#### 1.1 Creation of simulated datasets

Code necessary to replicate our results and generate the simulated datasets can be downloaded from https://github.com/plbaldoni/TranscriptDE-code. We simulated RNA-seq experiments in a variety of scenarios that are detailed in this section. Our simulation pipeline is organized in 4 main steps involving (1) the creation of a reference data set from real RNA-seq experiments, (2) the simulation of sequencing reads, (3) the quantification of simulated reads, and (4) differential transcript expression analysis. Below we describe each of these steps in detail.

##### 1.1.1 Reference dataset

A reference data set was generated from a real RNA-seq data from mouse experiment (NCBI Gene Expression Omnibus accession number GSE60450). For this reference dataset, a subset of relevant genes (protein-coding or lncRNA genes from reference chromosomes with expected CPM $>1$ in at least half of the samples) and their associated transcripts (protein-coding and lncRNA transcripts from relevant genes) was selected from the mouse transcriptome using the Gencode basic GTF annotation (version M27). Selected transcripts from the same gene were ranked (in decreasing order) according to their observed expression level (in TPM) averaged across all samples. Only transcripts with unique sequences from protein coding genes and long non-coding RNA (lncRNA) were considered.

More specifically, the selection of such a subset of relevant genes for which the expression of their transcripts would be simulated was done as follows. We summarized Salmon's quantification to the gene level using the function `tximeta::summarizeToGene`. Only protein-coding and lncRNA genes from chromosomes 1, ..., 22, X, and Y were considered. Next, we estimated baseline expression proportions using `edgeR::goodTuringProportions`. We selected relevant genes with an expected CPM>1 in at least 6 of the 12 libraries ($N_G = 13,176$). Only transcripts from relevant genes were considered in our simulation ($N_T = 41,372$). For each relevant gene, transcripts were ranked according to their sample-averaged TPM values obtained from Salmon's TPM quantifications. We used the baseline expression Good-Turing proportions of relevant genes to create a smoothing function (using `approxfun` function) to be used when simulating transcript-level expression, in a similar fashion to what was done in Law et al. (2014).

##### 1.1.2 Simulation of sequencing reads

Simulation scenarios varied according to the sequencing read length (50bp, 75bp, 100bp, 125bp, and 150bp), library size (either balanced with 50mi reads/sample or unbalanced with alternating 100mi and 25mi reads/sample), sequencing read type (either single-end or paired-end), maximum number of transcripts per gene considered (either 2, 3, 4, 5, or all transcripts available in the reference data set), the number of biological replicates per group (either 3, 5, or 10), and fold-change (either 2 or 1, in which the latter represents a null simulation without any differential expression). A total of 20 simulated experiments per scenario was generated. For each experiment, we simulated RNA-seq libraries for a total of 2 groups.

The relative expression levels of selected transcripts (the input for `Rsubread::simReads`) was simulated as follows. First, for a particular scenario, baseline expression proportions were generated for all selected transcripts using the smoothed Good-Turing proportions from our reference dataset. The maximum number of transcripts/gene considered in a given scenario as well as the ranking of each transcript (obtained from the reference dataset) dictated the set of selected transcripts in a simulation with only the most expressed ones (top ranked) being selected. For example, in a scenario with only 3 transcripts per gene being expressed, we simulate a positive expression level for all transcripts from genes that express at most 1 or 2 transcripts and, for genes that expresses 3 or more transcripts, only the top-ranked 3 transcripts had a positive expression. A subset of 3000 randomly selected transcripts had their baseline proportions adjusted with a 2 fold-change to create group-specific proportions with 1500 up-regulated and 1500 down-regulated transcripts. For each group, proportions were then transformed to sample-specific expected counts $\mu_{ts}$, for transcript $t$ and sample $s$, depending on the library size of each sample.

Biological variation was incorporated in the simulation with a trend on the expected count for each sample. This trend had the form $\mathrm{BCV}_{ts} = 0.2 + 1/\sqrt{\mu_{ts}}$. Dispersions $\phi_{ts}$ were generated with random shifts around the trend as $\phi_{ts} = \mathrm{BCV}_{ts}^2 \times \frac{df}{\chi^2_{df}}$ with $df = 40$. In this simulation, samples belonging to the same group share the random shift $\chi^2_{40}$. In other words, for each transcript and each group, a single random variable was drawn from $\chi^2_{40}$ and used to all biological replicates of that group. Note that (1) this approach is slightly different to the approach used in the `voom` paper, in which there were sample- and gene-specific random shifts around the trend to generate dispersions, and (2) this approach does not imply that there is no biological variability among samples from the same group (which will be introduced by the Gamma-Poisson model), but rather it just implies that transcript-wise expression levels from samples of the same group share the same mean and dispersion parameters (as they should). Apart from the differences in library size across replicates, the only variation among replicates should be a result of the variance model resulting from the Gamma-Poisson distribution. Since we generated differential expression states directly on the baseline proportions to define groups, it makes sense to have a single random shift around the dispersion trend per group, hence having a single dispersion shared among libraries of the same group.

Expected counts and dispersions were used to generate transcript-level expression following a Gamma distribution. Resulting transcript-wise expression levels were divided by the transcript length and scaled up to $1 \times 10^6$ to generate transcript-wise TPMs that were used as input in `Rsubread::simReads`. For read lengths other than 75 bp or 100bp, quality scores were samples from real data (ENCFF713MNU data for 50bp, ENCFF126GLV for 125bp, and ENCFF102BXZ for 150 bp experiments) and used as an input parameter in `Rsubread::simReads`. Note that quality scores are disregarded by Salmon and kallisto during quantification, and their choice is irrelevant to the overall results of this simulations study.

##### 1.1.3 Quantification

Sequencing reads in FASTQ format generated by `simReads` were quantified by Salmon (v. 1.9.0) and kallisto (v. 0.46.1). For both quantification algorithms, we used transcriptomic index from the complete Gencode annotation (version M27) and we generated a total of 100 bootstraps samples for each library. For `Salmon`, we used a decoy-aware mapping-based indexed transcriptome generated for the mouse mm39 reference genome with k-mers of length 31. For Salmon, the option `--validateMappings` was used as recommended in the software documentation. For single-end read libraries, we provided kallisto and salmon the true mean and standard deviation of the fragment length distribution (180 and 40, respectively) that is the default and used in `simReads`. To read Salmon quantification files in `sleuth`, Salmon quantification files `quant.sf` were transformed to `abundance.h5` files with the function `prepare_fish_for_sleuth` from the `wasabi` package (v. 1.0.1).

##### 1.1.4 Differential transcript expression

We compared differential transcript expression (DTE) among methods `edgeR-Raw` (edgeR using raw counts), `edgeR-Scaled` (edgeR using deflated counts), `sleuth-LRT` (with likelihood ratio test), `sleuth-Wald` (with Wald test), and `Swish`. In both `edgeR-Raw` and `edgeR-Scaled`, the QLF pipeline with default options in all functions was used. Transcript filtering in `edgeR` was performed with `filterByExpr` with default options. Default filtering functions were used in `sleuth` (transcripts with at least 5 counts in at least 47% of the samples) and `Swish` (transcripts with at least 10 counts in at least 3 samples). We acknowledge that using different filtering approach by each method introduce an extra, but nonetheless minimal, level variability that is separate from the statistical approach. Both `sleuth` and `Swish` were run with their default pipeline with default options. Unless otherwise noted, transcripts were claimed to be differentially expressed with a 0.05 FDR threshold.

#### 1.2 Simulation results

##### 1.2.1 Power, false discovery rate control, and speed

In this subsection, we present results from our simulations comparing methods with respect to power to detect DE transcripts, false discovery rate control, and computational speed.

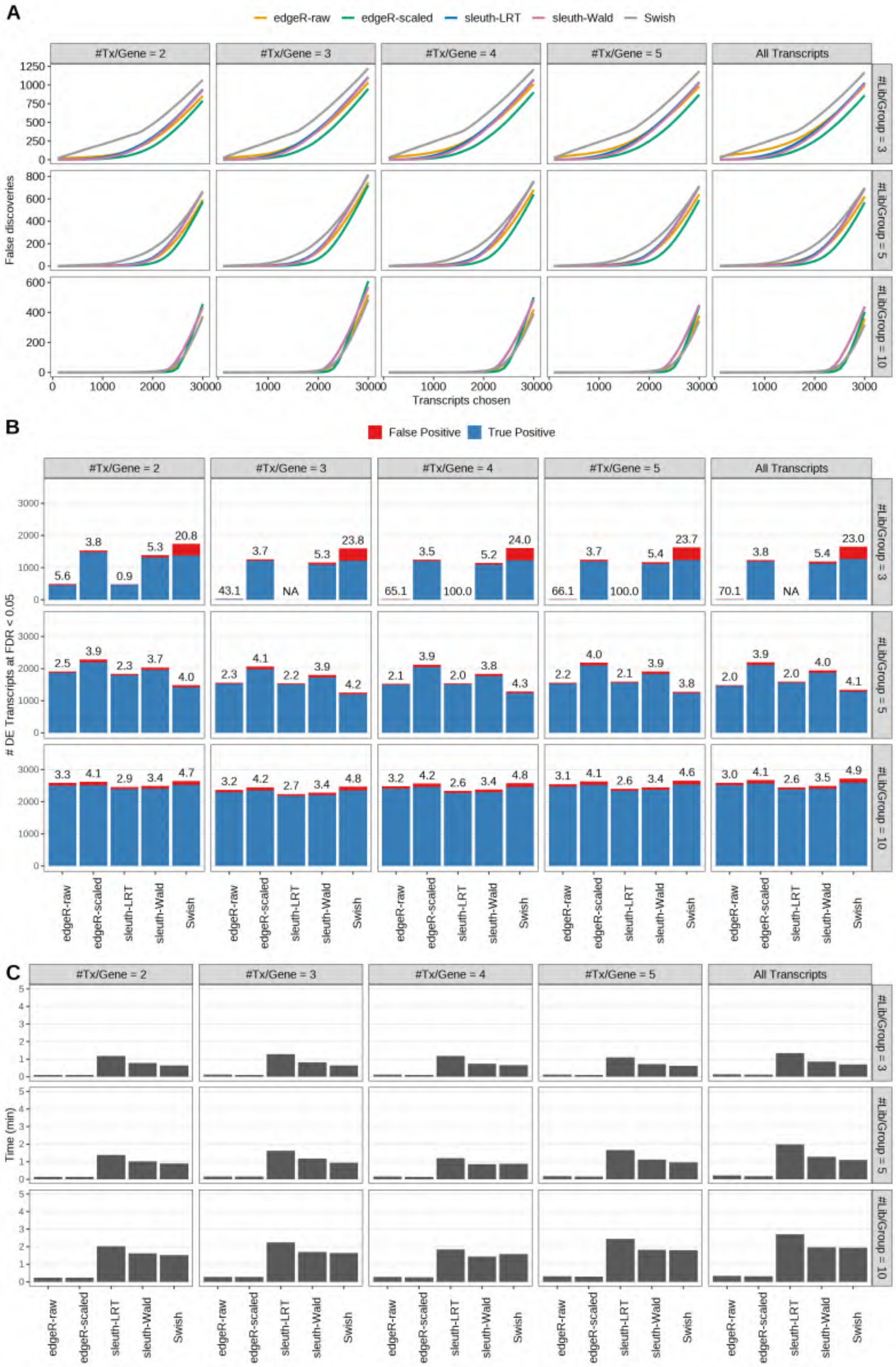

Figure S1: Simulation results. Scenario with mm39 genome, 50bp single-end reads quantified with Salmon, and balanced libraries. (A) Average number of false discoveries as a function of the number of chosen transcripts. (B) Average number of true (blue) and false (red) positive DE transcripts. Observed FDR annotated over bars. (C) Average computing time in minutes.

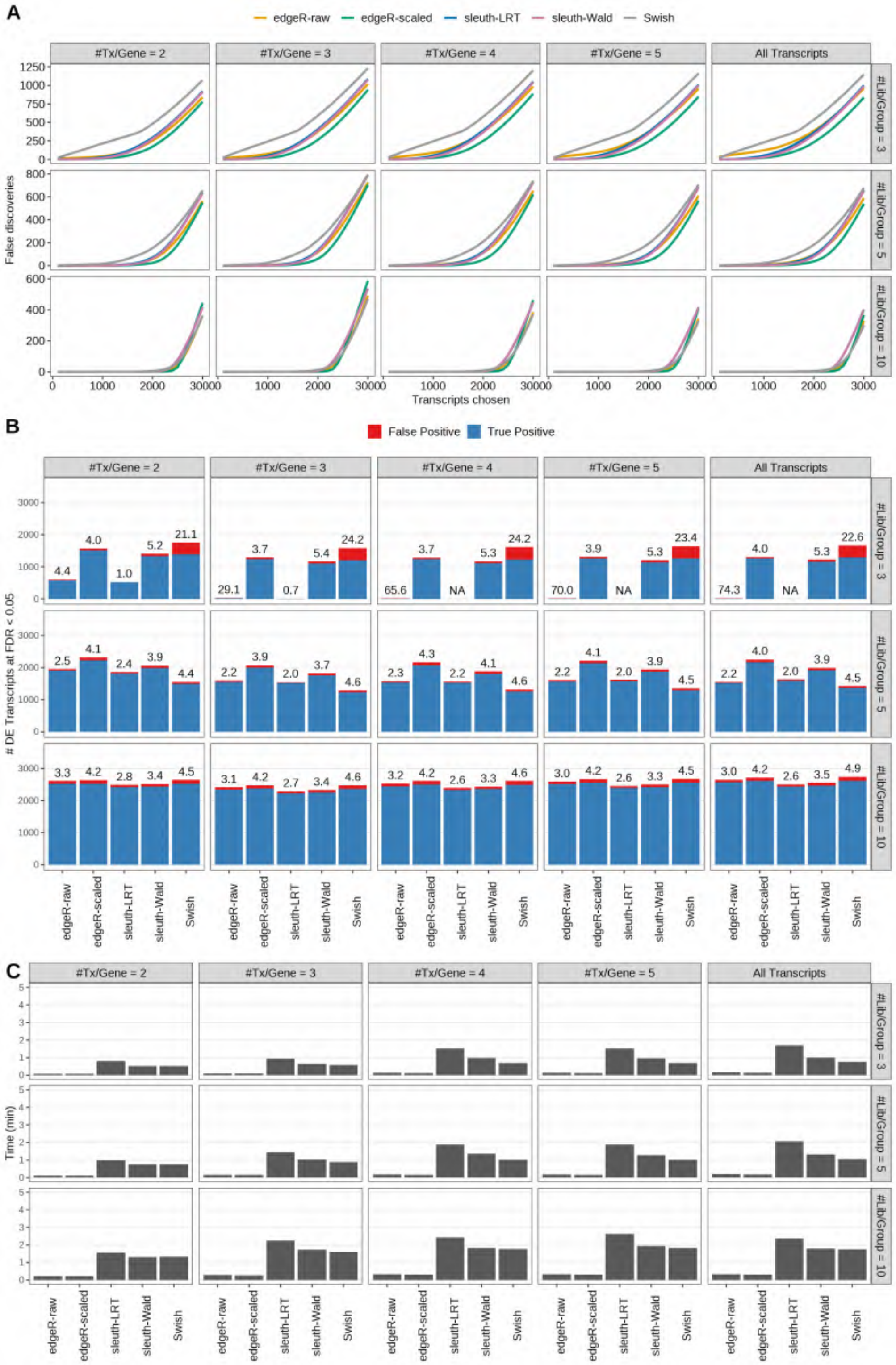

Figure S2: Simulation results. Scenario with mm39 genome, 75bp single-end reads quantified with Salmon, and balanced libraries. (A) Average number of false discoveries as a function of the number of chosen transcripts. (B) Average number of true (blue) and false (red) positive DE transcripts. Observed FDR annotated over bars. (C) Average computing time in minutes.

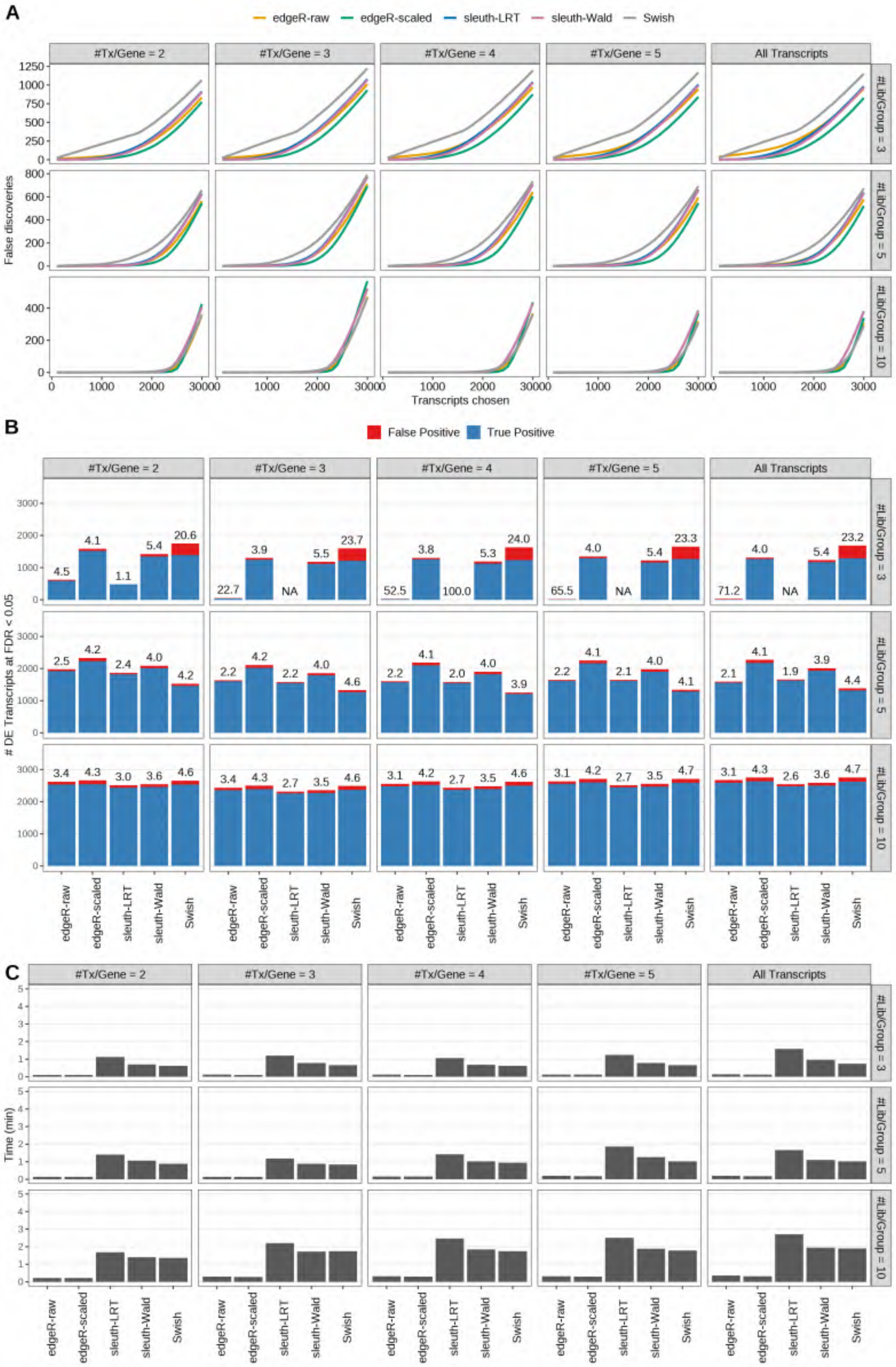

Figure S3: Simulation results. Scenario with mm39 genome, 100bp single-end reads quantified with Salmon, and balanced libraries. (A) Average number of false discoveries as a function of the number of chosen transcripts. (B) Average number of true (blue) and false (red) positive DE transcripts. Observed FDR annotated over bars. (C) Average computing time in minutes.

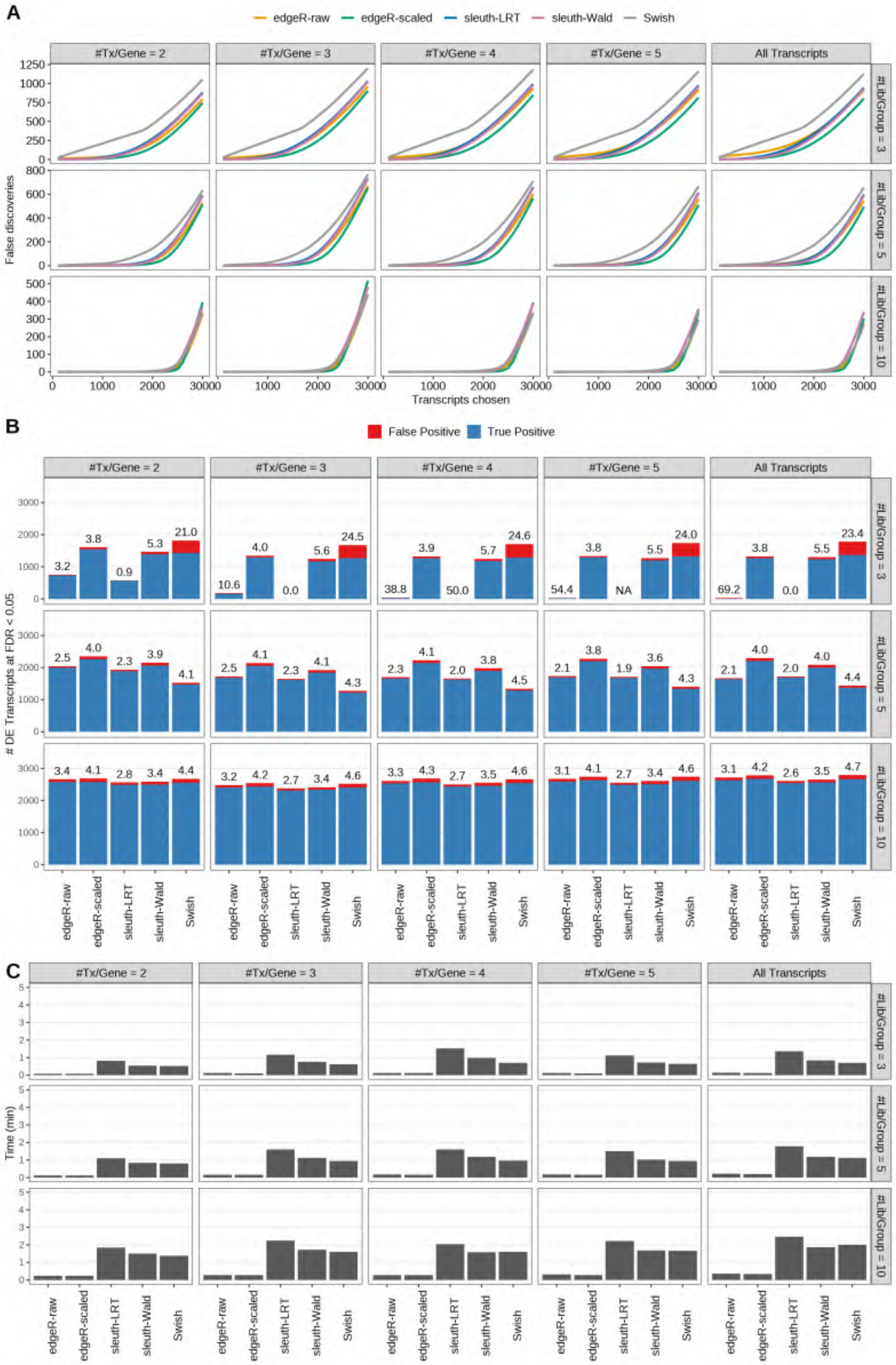

Figure S4: Simulation results. Scenario with mm39 genome, 125bp single-end reads quantified with Salmon, and balanced libraries. (A) Average number of false discoveries as a function of the number of chosen transcripts. (B) Average number of true (blue) and false (red) positive DE transcripts. Observed FDR annotated over bars. (C) Average computing time in minutes.

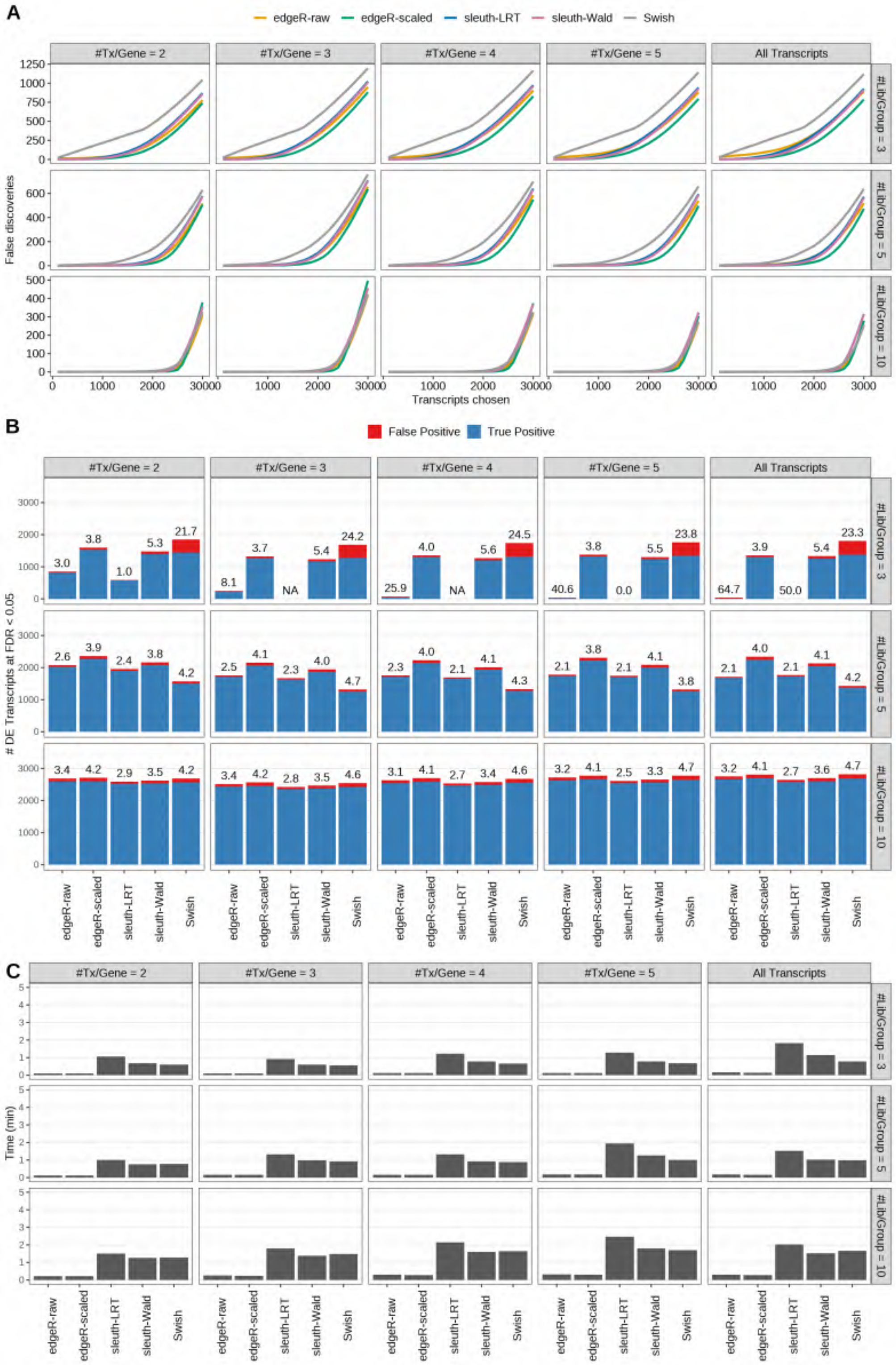

Figure S5: Simulation results. Scenario with mm39 genome, 150bp single-end reads quantified with Salmon, and balanced libraries. (A) Average number of false discoveries as a function of the number of chosen transcripts. (B) Average number of true (blue) and false (red) positive DE transcripts. Observed FDR annotated over bars. (C) Average computing time in minutes.

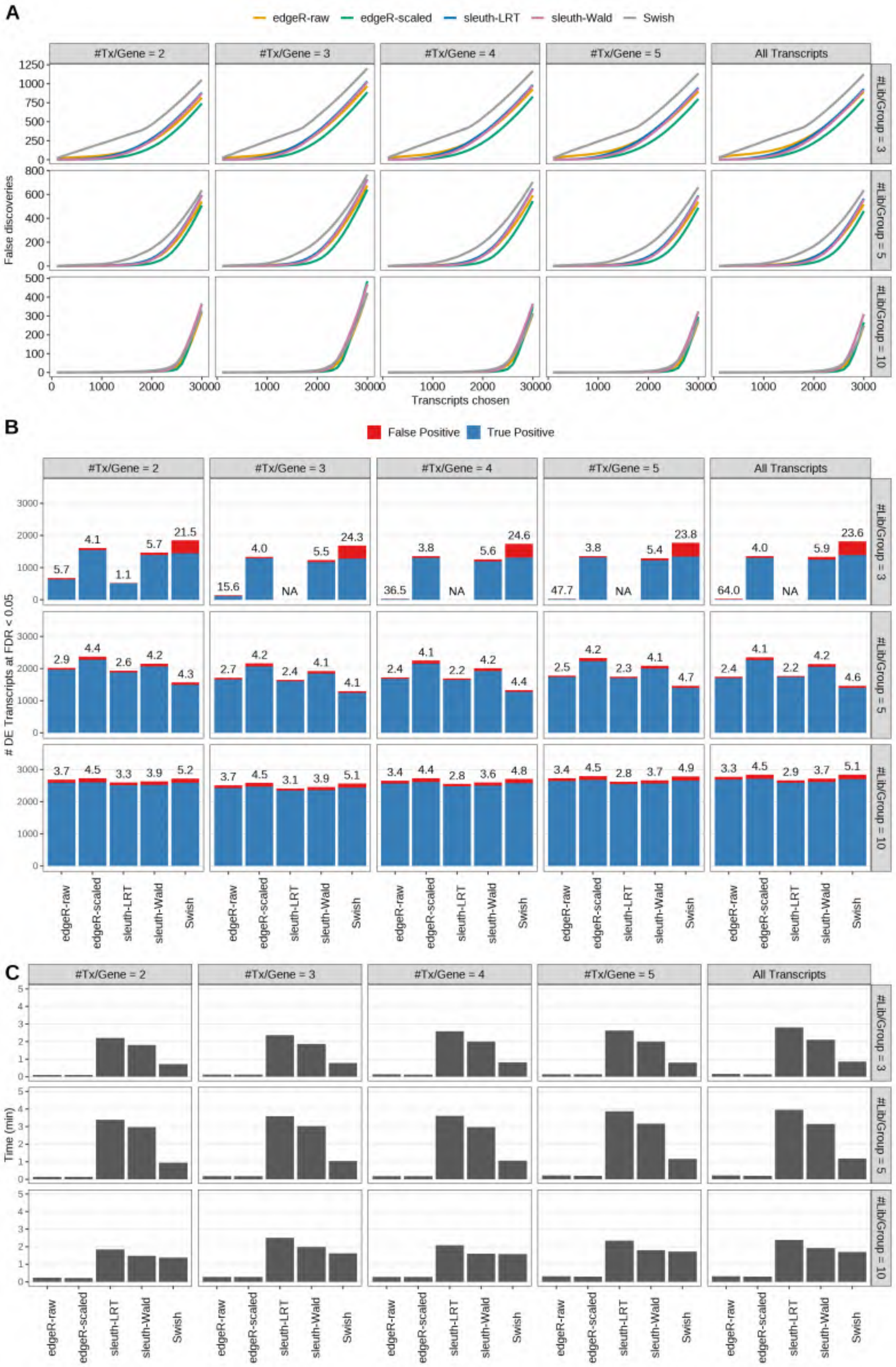

Figure S6: Simulation results. Scenario with mm39 genome, 50bp paired-end reads quantified with Salmon, and balanced libraries. (A) Average number of false discoveries as a function of the number of chosen transcripts. (B) Average number of true (blue) and false (red) positive DE transcripts. Observed FDR annotated over bars. (C) Average computing time in minutes.

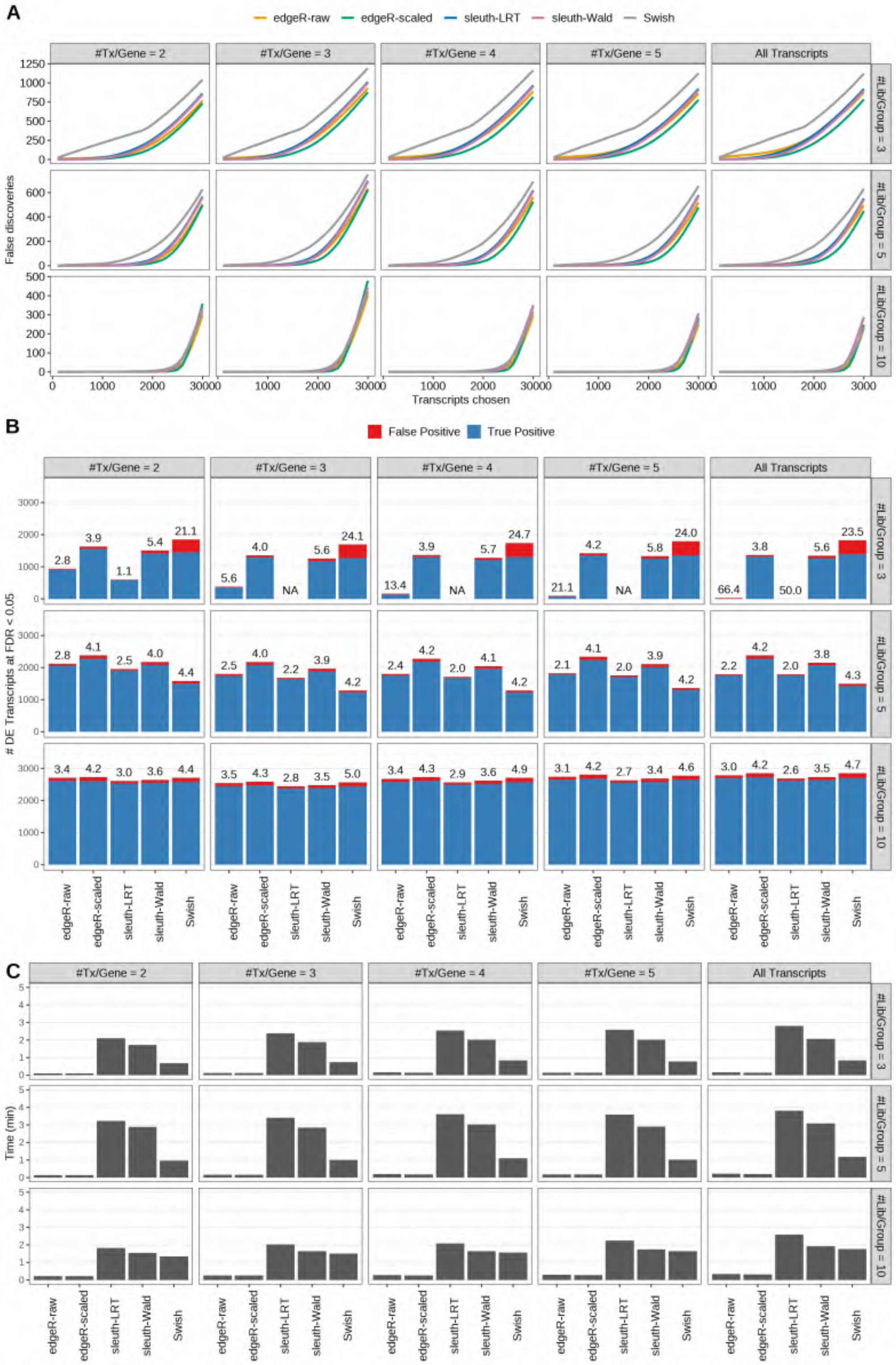

Figure S7: Simulation results. Scenario with mm39 genome, 75bp paired-end reads quantified with Salmon, and balanced libraries. (A) Average number of false discoveries as a function of the number of chosen transcripts. (B) Average number of true (blue) and false (red) positive DE transcripts. Observed FDR annotated over bars. (C) Average computing time in minutes.

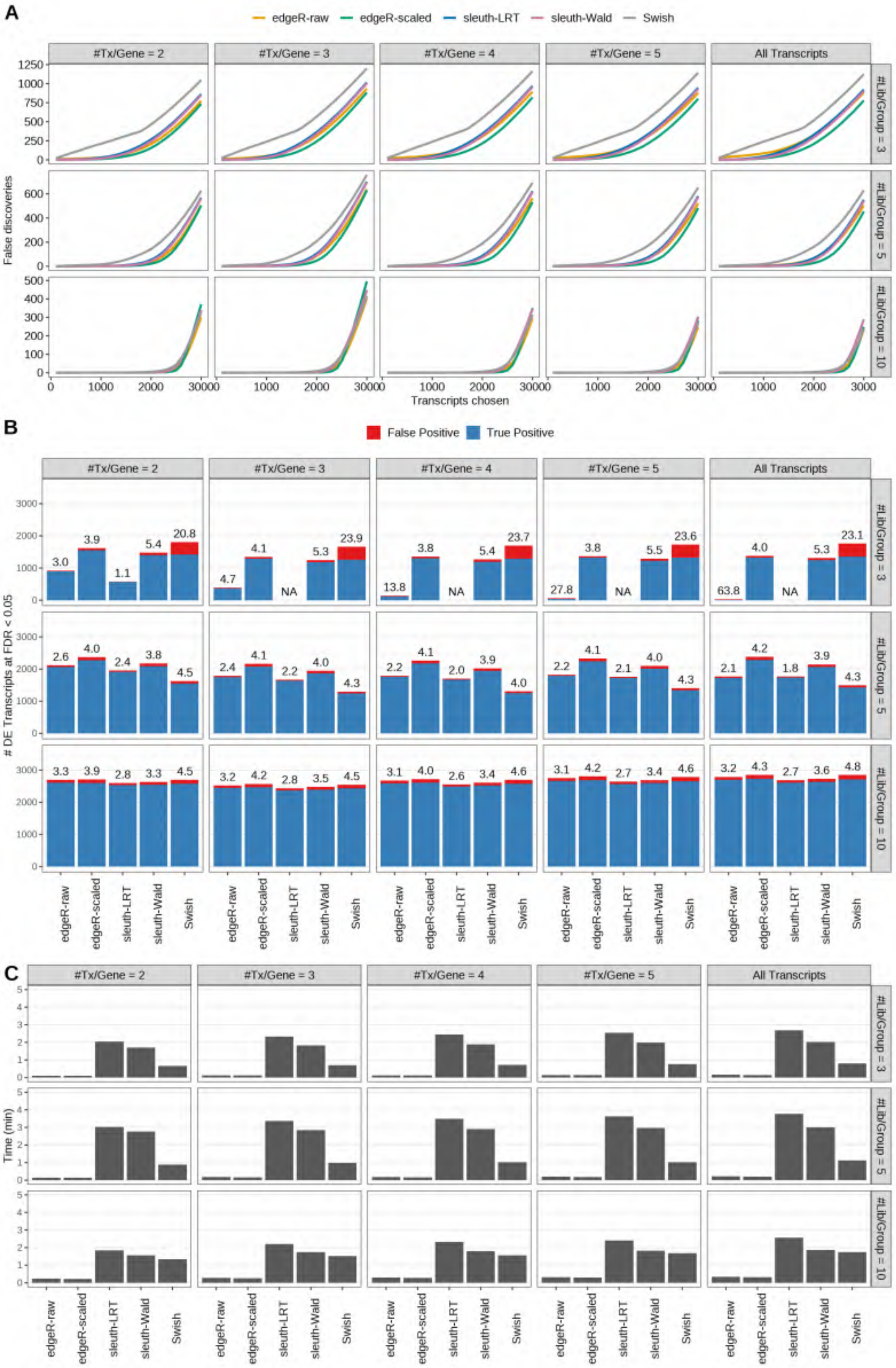

Figure S8: Simulation results. Scenario with mm39 genome, 100bp paired-end reads quantified with Salmon, and balanced libraries. (A) Average number of false discoveries as a function of the number of chosen transcripts. (B) Average number of true (blue) and false (red) positive DE transcripts. Observed FDR annotated over bars. (C) Average computing time in minutes.

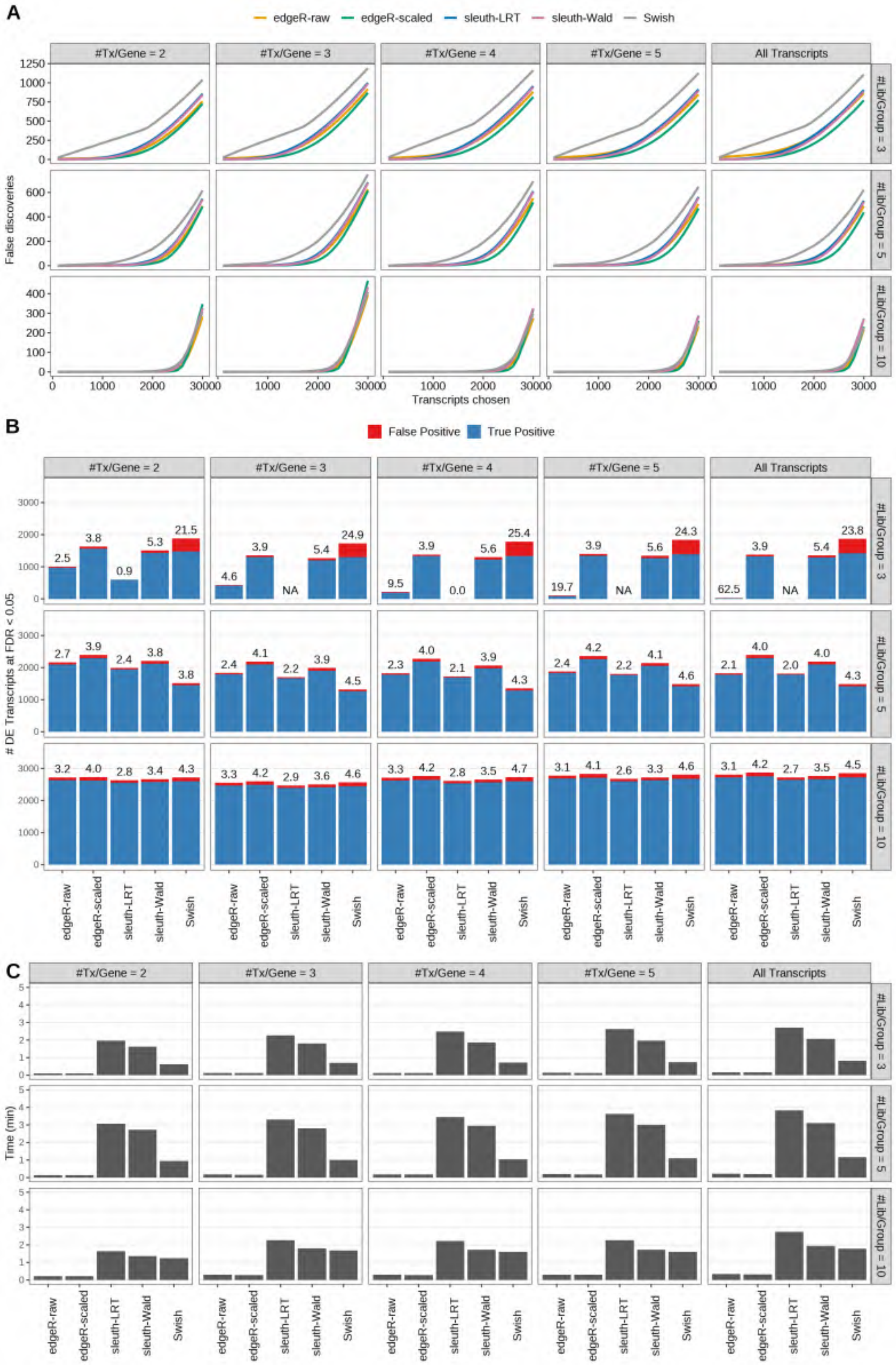

Figure S9: Simulation results. Scenario with mm39 genome, 125bp paired-end reads quantified with Salmon, and balanced libraries. (A) Average number of false discoveries as a function of the number of chosen transcripts. (B) Average number of true (blue) and false (red) positive DE transcripts. Observed FDR annotated over bars. (C) Average computing time in minutes.

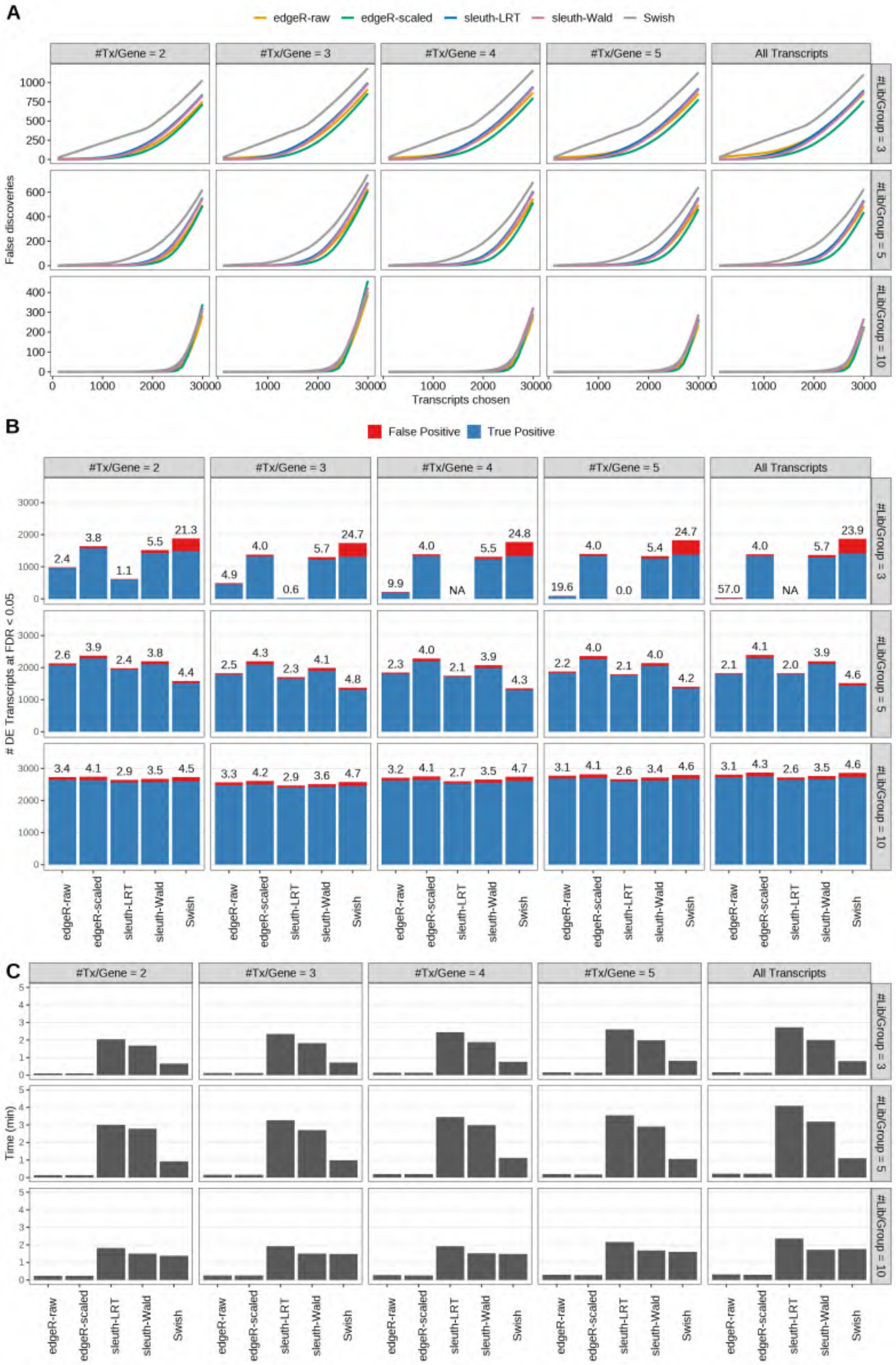

Figure S10: Simulation results. Scenario with mm39 genome, 150bp paired-end reads quantified with Salmon, and balanced libraries. (A) Average number of false discoveries as a function of the number of chosen transcripts. (B) Average number of true (blue) and false (red) positive DE transcripts. Observed FDR annotated over bars. (C) Average computing time in minutes.

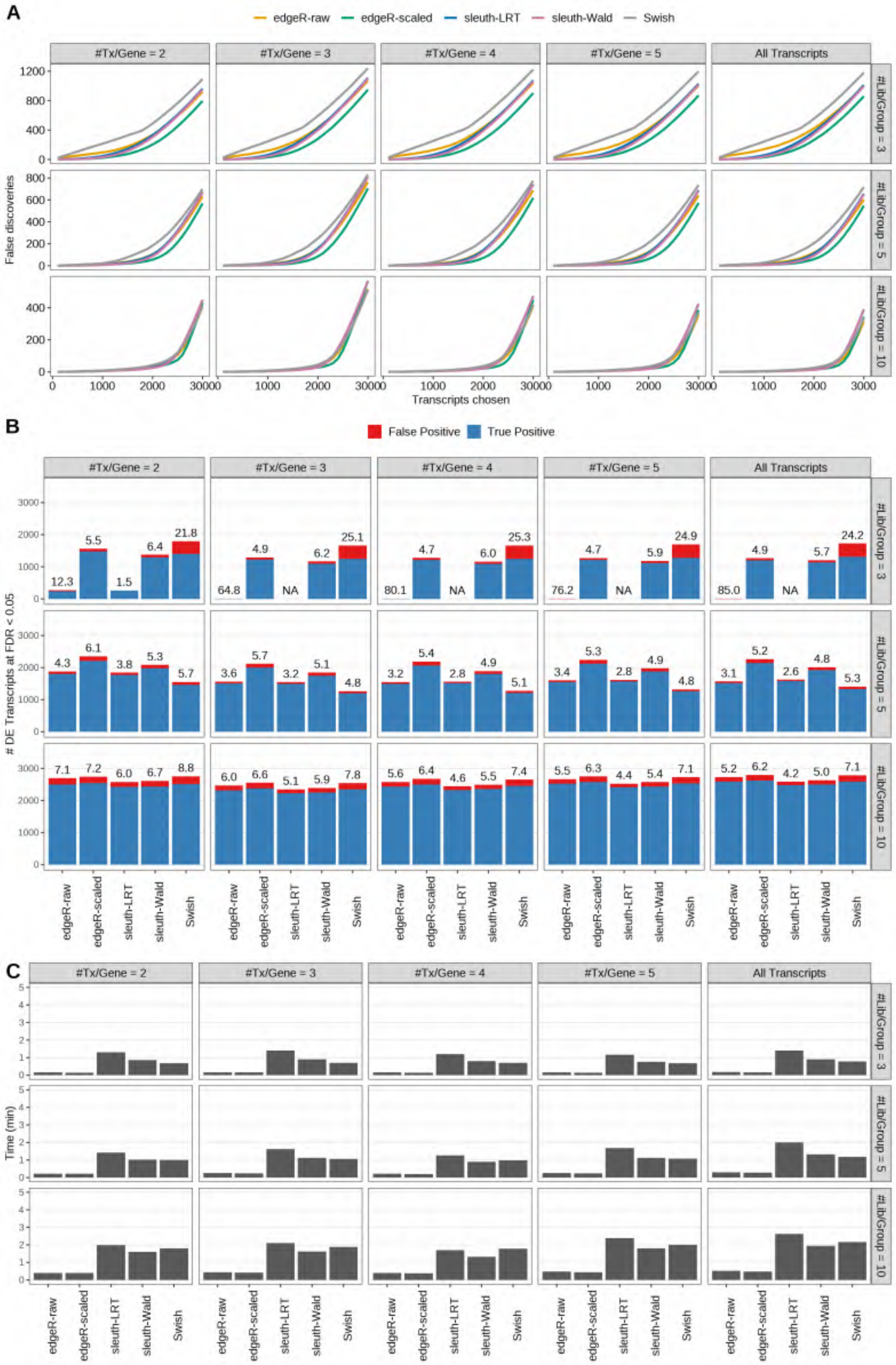

Figure S11: Simulation results. Scenario with mm39 genome, 50bp single-end reads quantified with kallisto, and balanced libraries. (A) Average number of false discoveries as a function of the number of chosen transcripts. (B) Average number of true (blue) and false (red) positive DE transcripts. Observed FDR annotated over bars. (C) Average computing time in minutes.

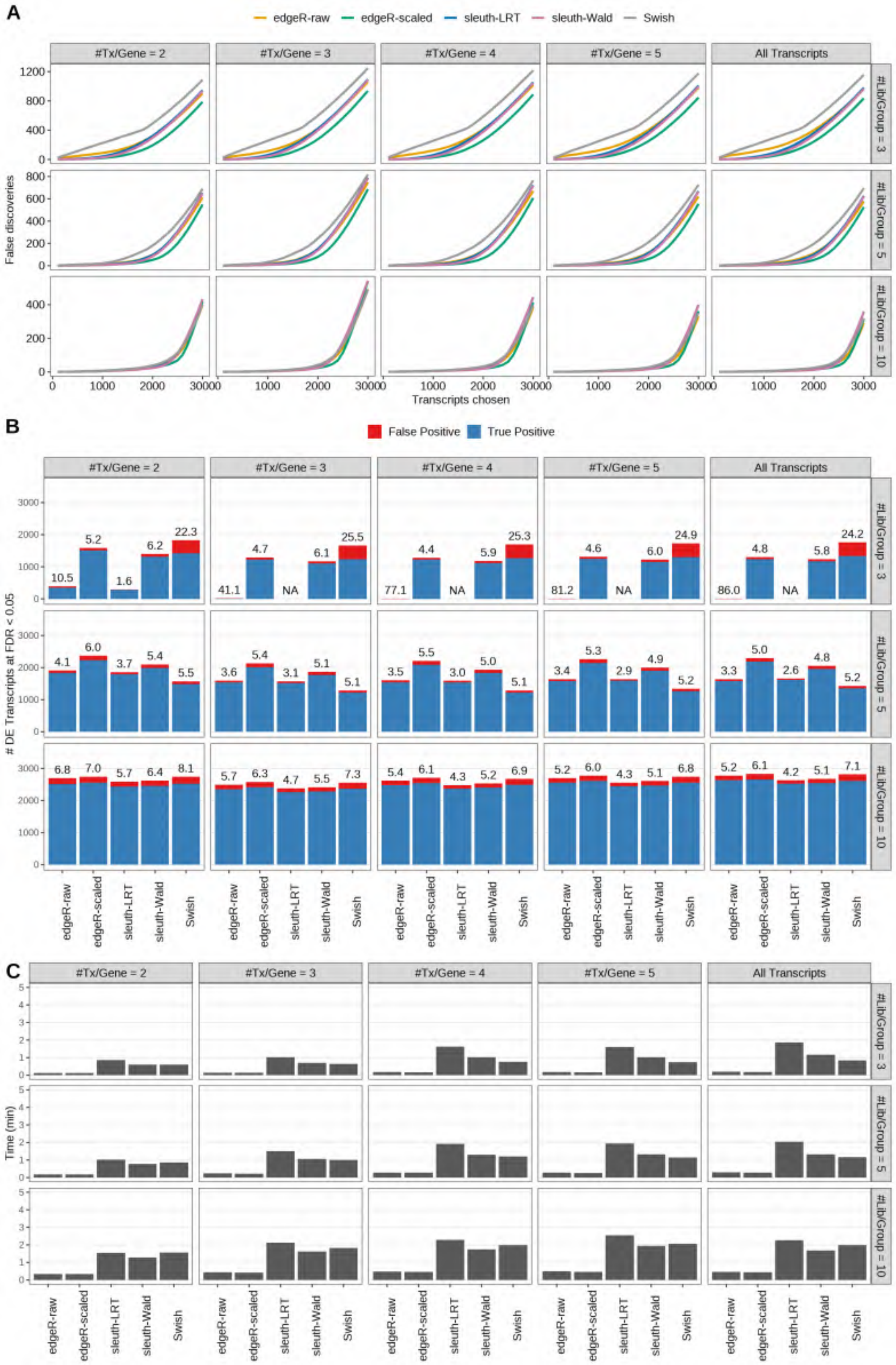

Figure S12: Simulation results. Scenario with mm39 genome, 75bp single-end reads quantified with kallisto, and balanced libraries. (A) Average number of false discoveries as a function of the number of chosen transcripts. (B) Average number of true (blue) and false (red) positive DE transcripts. Observed FDR annotated over bars. (C) Average computing time in minutes.

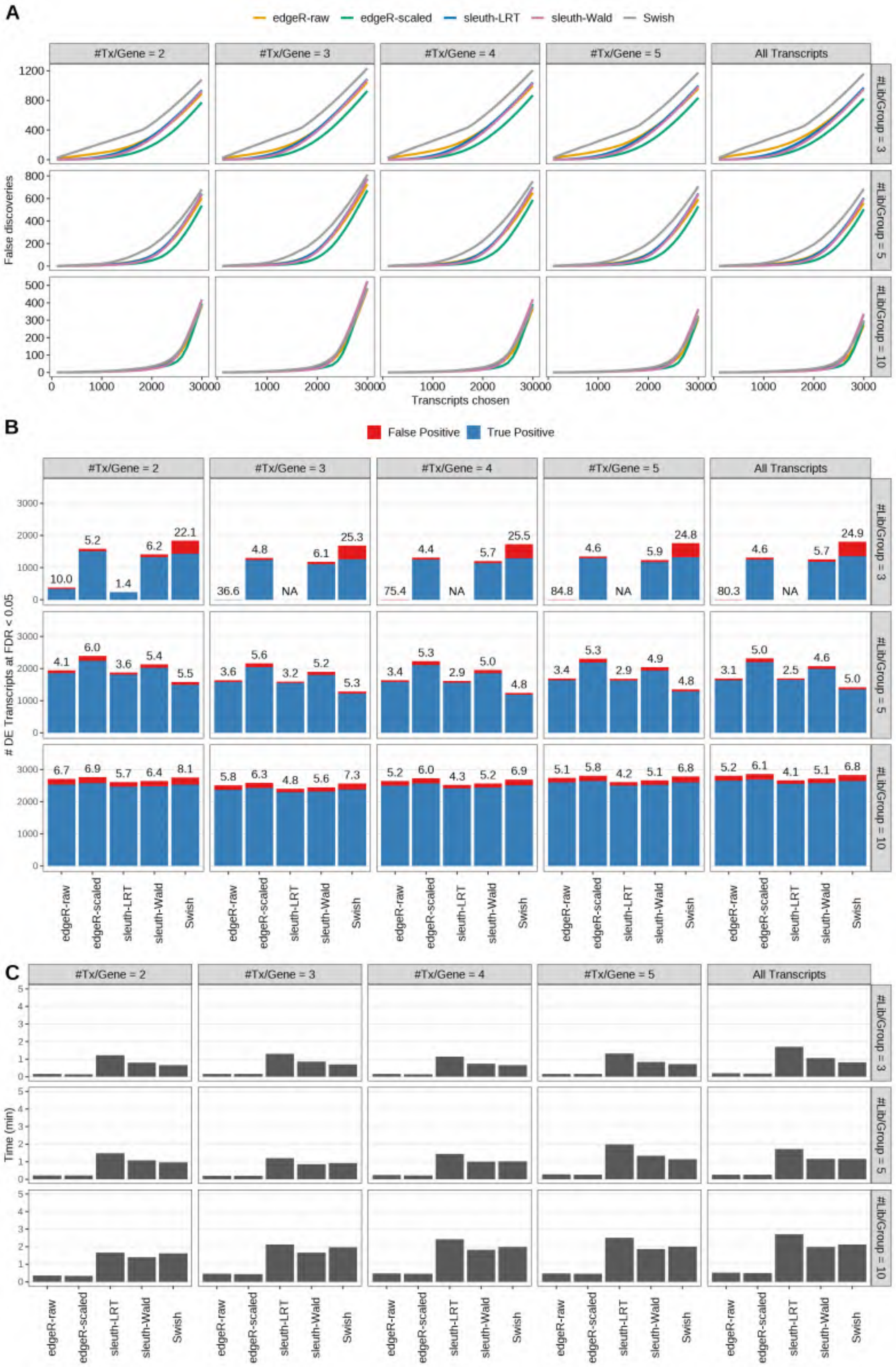

Figure S13: Simulation results. Scenario with mm39 genome, 100bp single-end reads quantified with kallisto, and balanced libraries. (A) Average number of false discoveries as a function of the number of chosen transcripts. (B) Average number of true (blue) and false (red) positive DE transcripts. Observed FDR annotated over bars. (C) Average computing time in minutes.

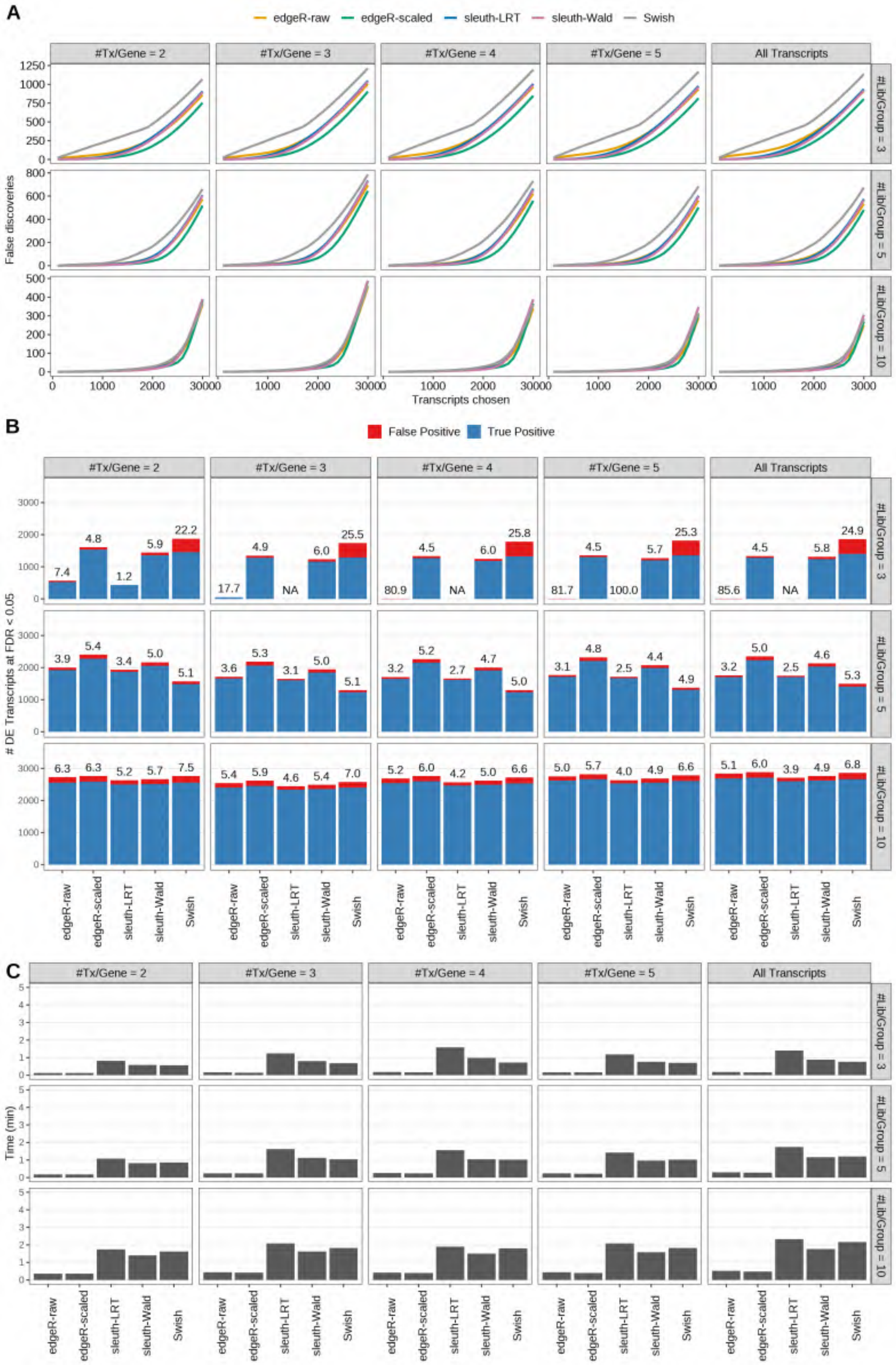

Figure S14: Simulation results. Scenario with mm39 genome, 125bp single-end reads quantified with kallisto, and balanced libraries. (A) Average number of false discoveries as a function of the number of chosen transcripts. (B) Average number of true (blue) and false (red) positive DE transcripts. Observed FDR annotated over bars. (C) Average computing time in minutes.

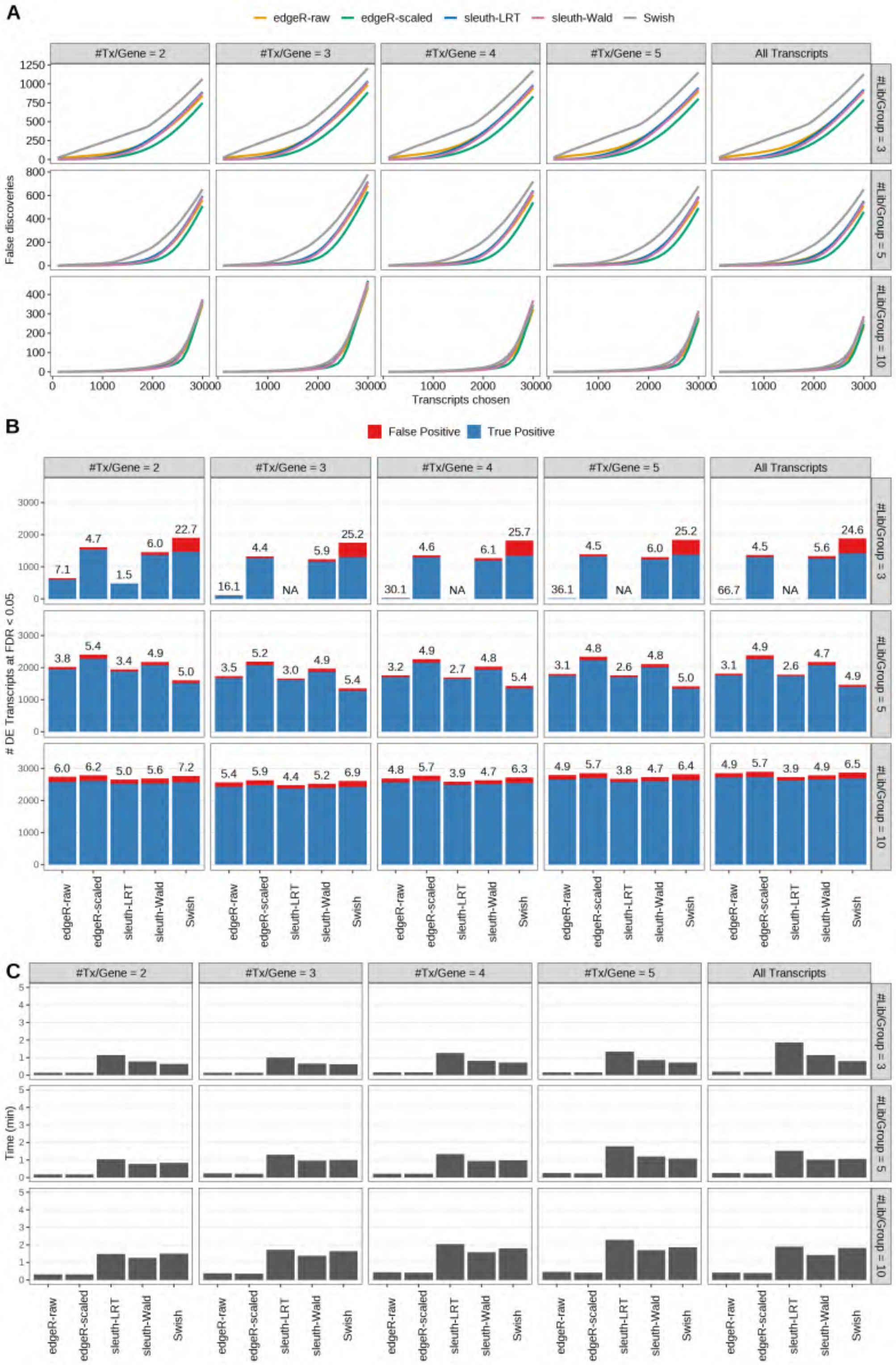

Figure S15: Simulation results. Scenario with mm39 genome, 150bp single-end reads quantified with kallisto, and balanced libraries. (A) Average number of false discoveries as a function of the number of chosen transcripts. (B) Average number of true (blue) and false (red) positive DE transcripts. Observed FDR annotated over bars. (C) Average computing time in minutes.

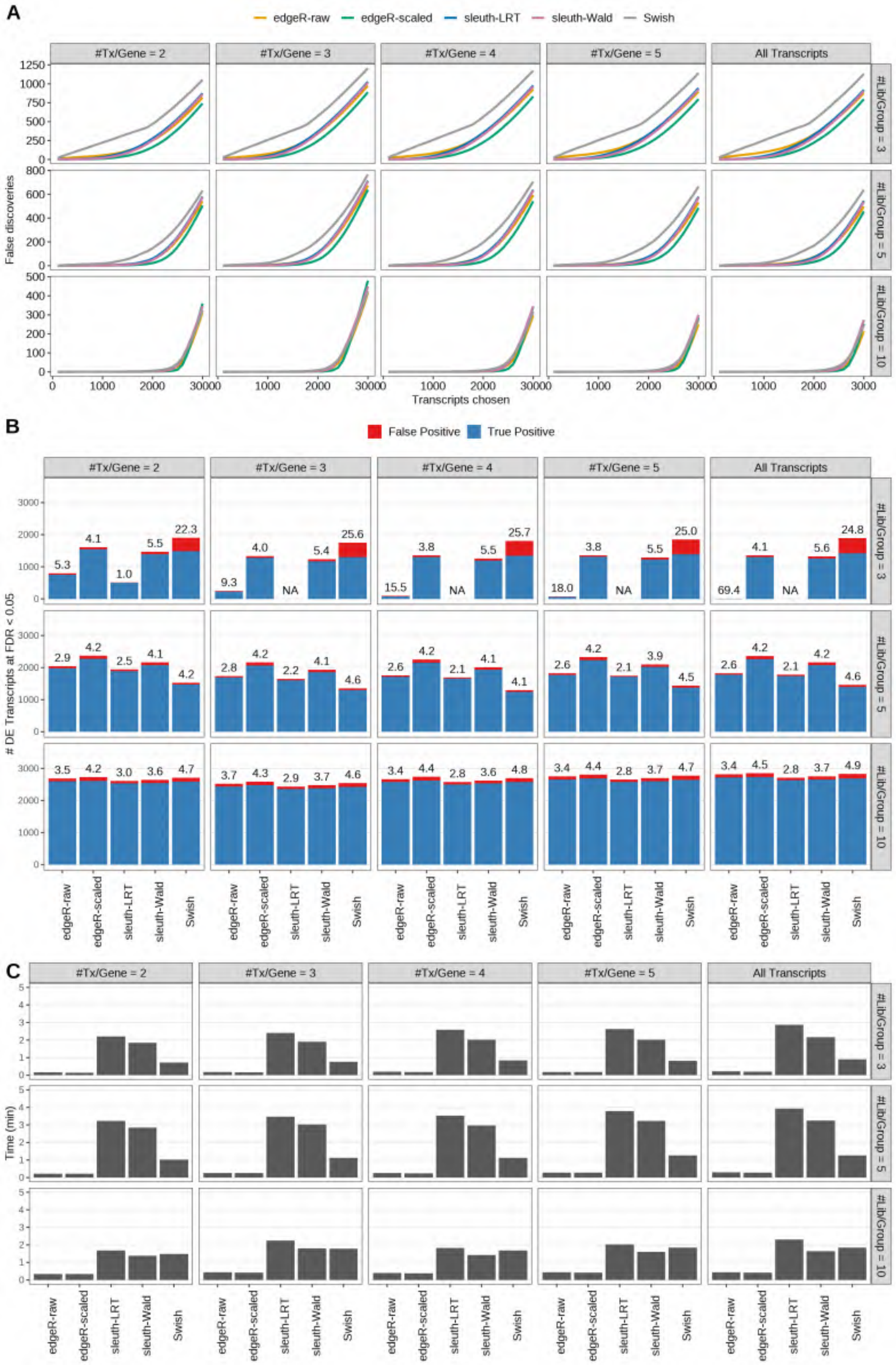

Figure S16: Simulation results. Scenario with mm39 genome, 50bp paired-end reads quantified with kallisto, and balanced libraries. (A) Average number of false discoveries as a function of the number of chosen transcripts. (B) Average number of true (blue) and false (red) positive DE transcripts. Observed FDR annotated over bars. (C) Average computing time in minutes.

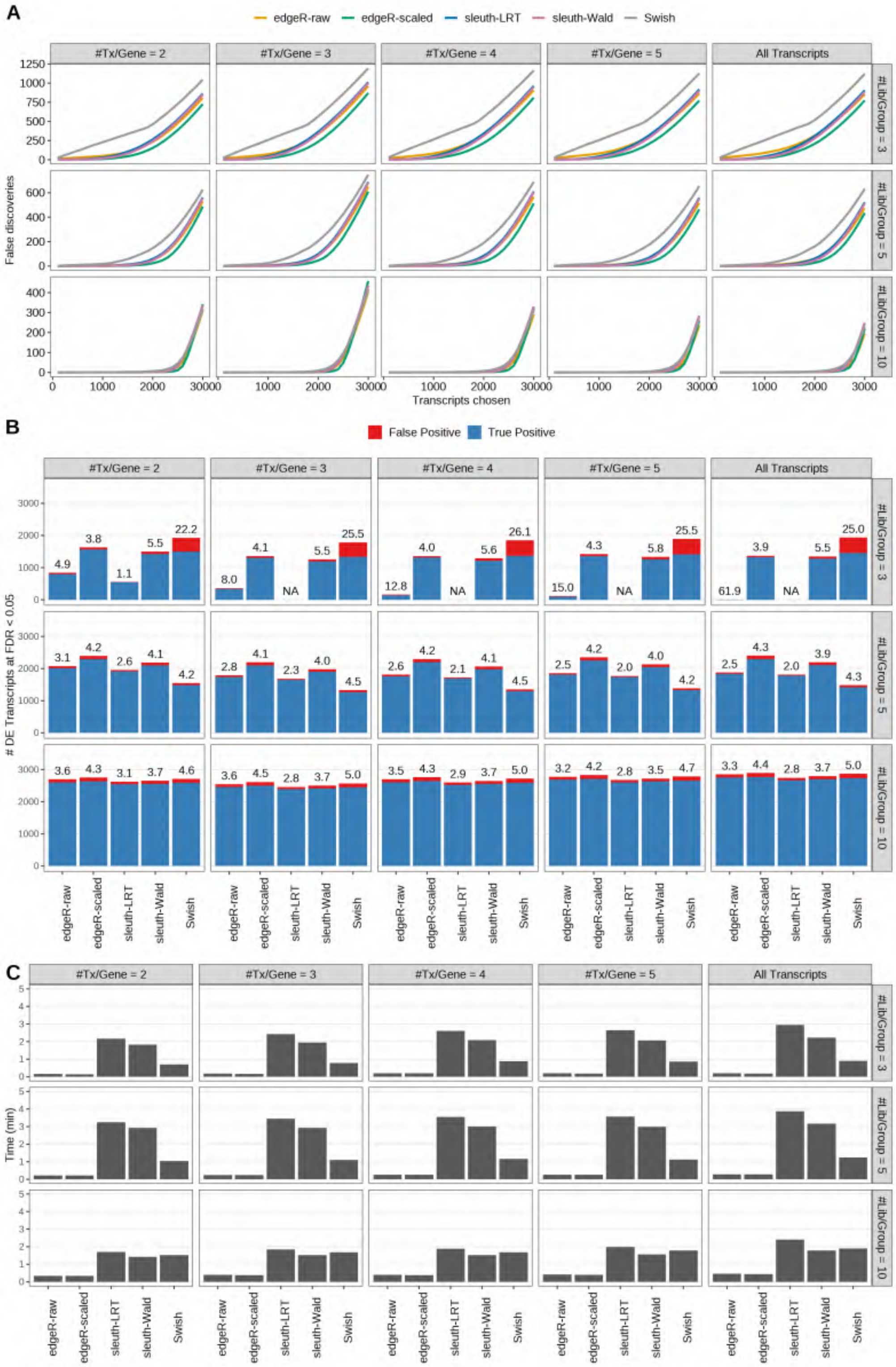

Figure S17: Simulation results. Scenario with mm39 genome, 75bp paired-end reads quantified with kallisto, and balanced libraries. (A) Average number of false discoveries as a function of the number of chosen transcripts. (B) Average number of true (blue) and false (red) positive DE transcripts. Observed FDR annotated over bars. (C) Average computing time in minutes.

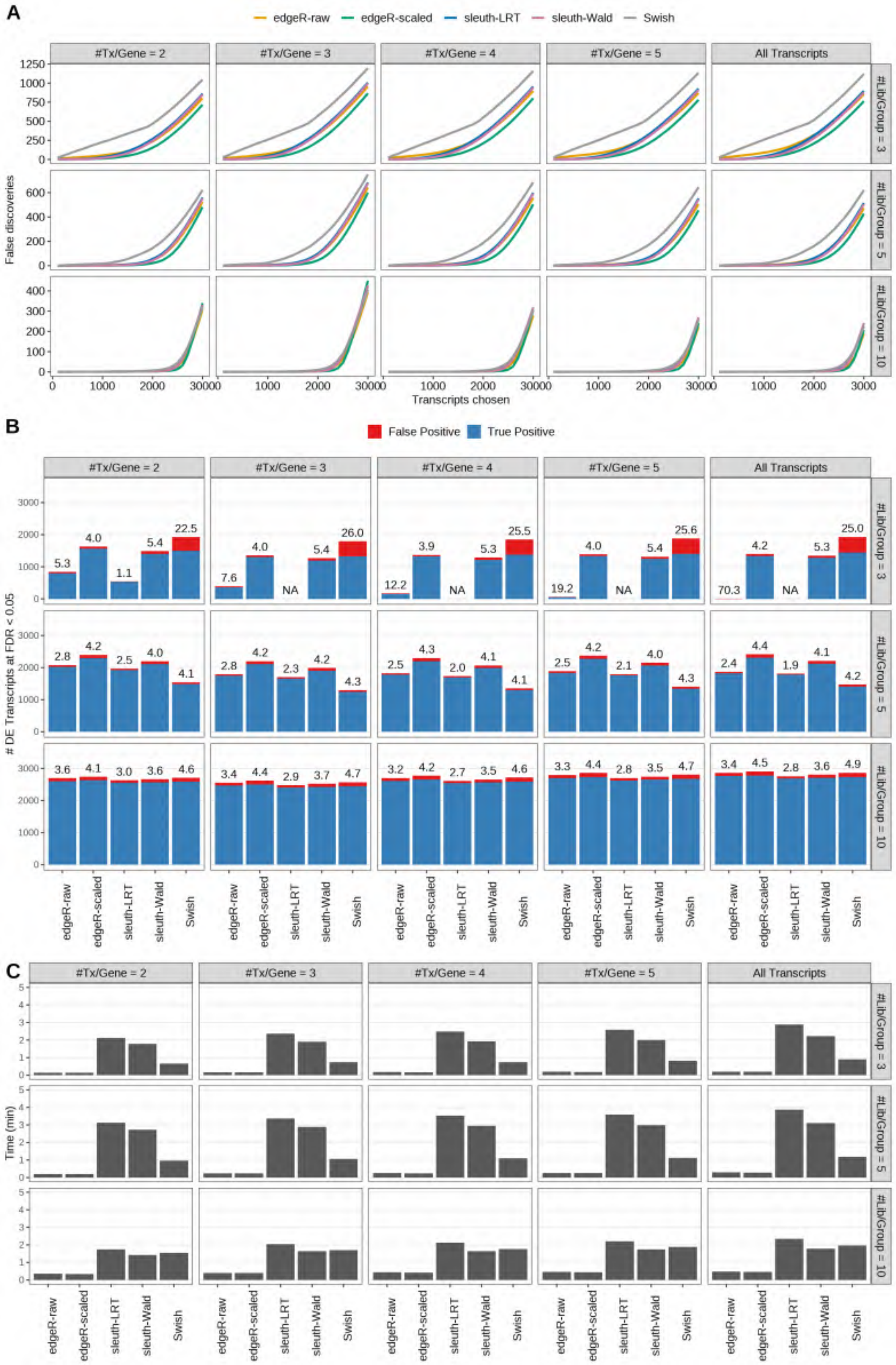

Figure S18: Simulation results. Scenario with mm39 genome, 100bp paired-end reads quantified with kallisto, and balanced libraries. (A) Average number of false discoveries as a function of the number of chosen transcripts. (B) Average number of true (blue) and false (red) positive DE transcripts. Observed FDR annotated over bars. (C) Average computing time in minutes.

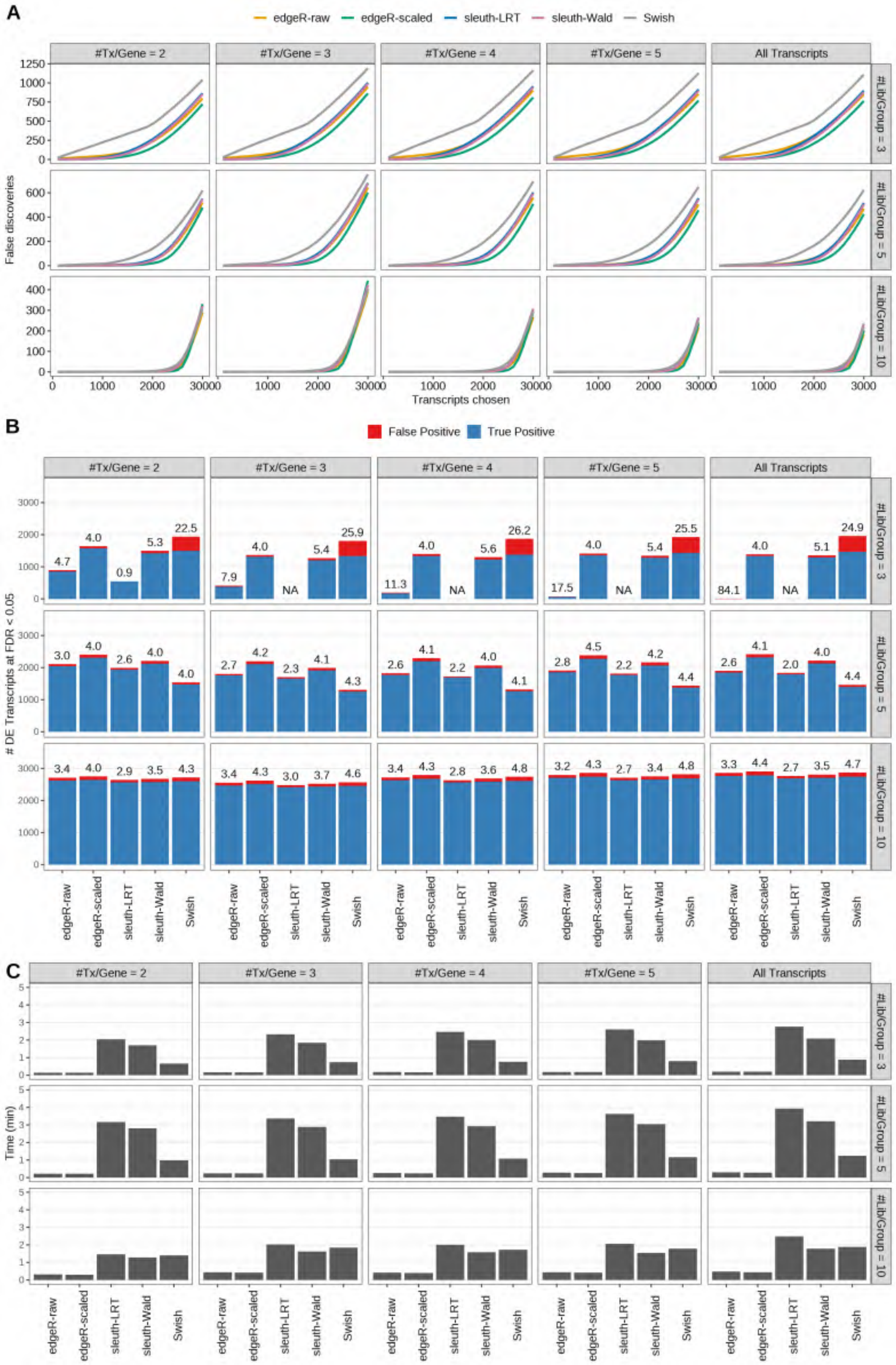

Figure S19: Simulation results. Scenario with mm39 genome, 125bp paired-end reads quantified with kallisto, and balanced libraries. (A) Average number of false discoveries as a function of the number of chosen transcripts. (B) Average number of true (blue) and false (red) positive DE transcripts. Observed FDR annotated over bars. (C) Average computing time in minutes.

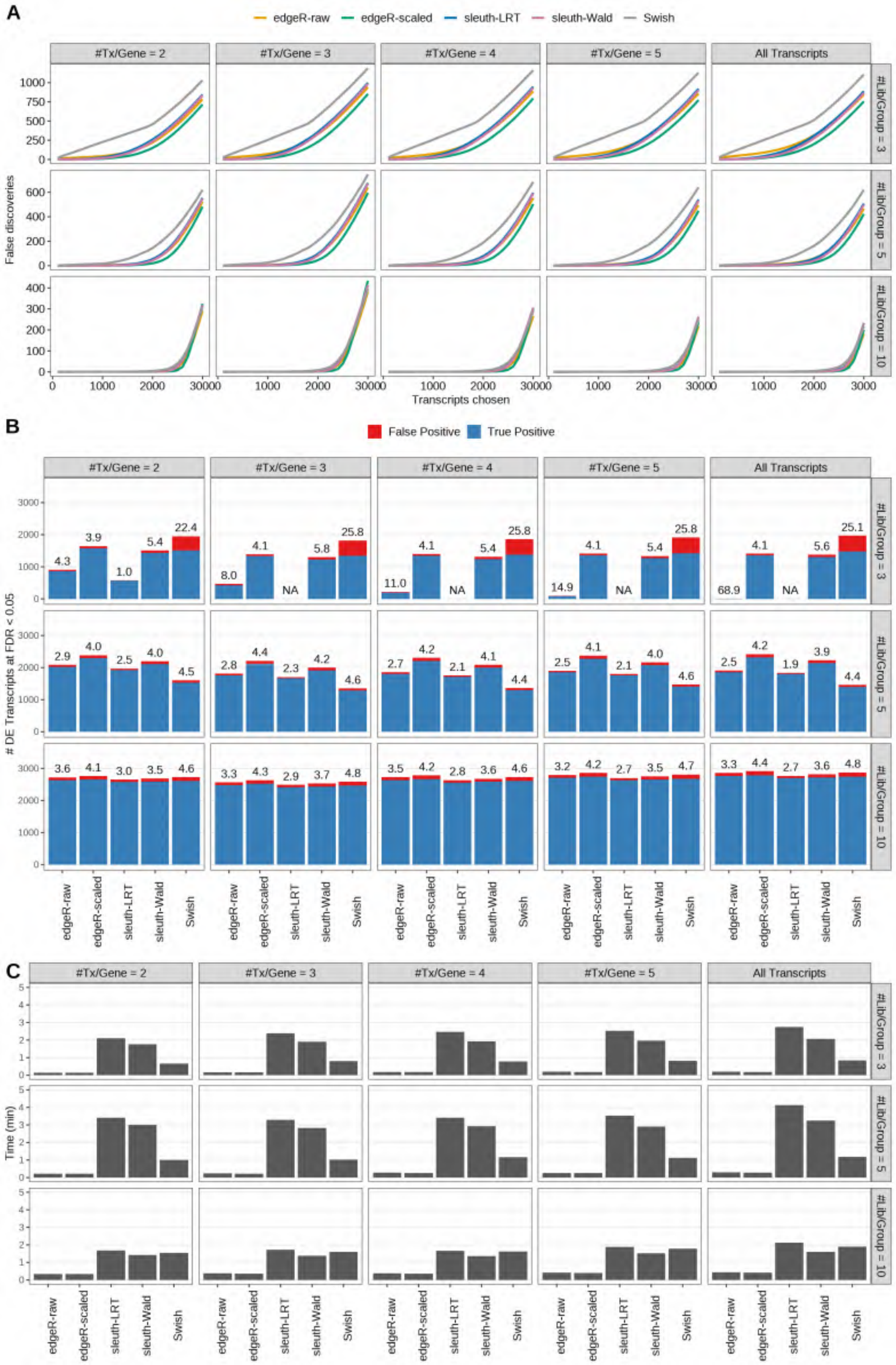

Figure S20: Simulation results. Scenario with mm39 genome, 150bp paired-end reads quantified with kallisto, and balanced libraries. (A) Average number of false discoveries as a function of the number of chosen transcripts. (B) Average number of true (blue) and false (red) positive DE transcripts. Observed FDR annotated over bars. (C) Average computing time in minutes.

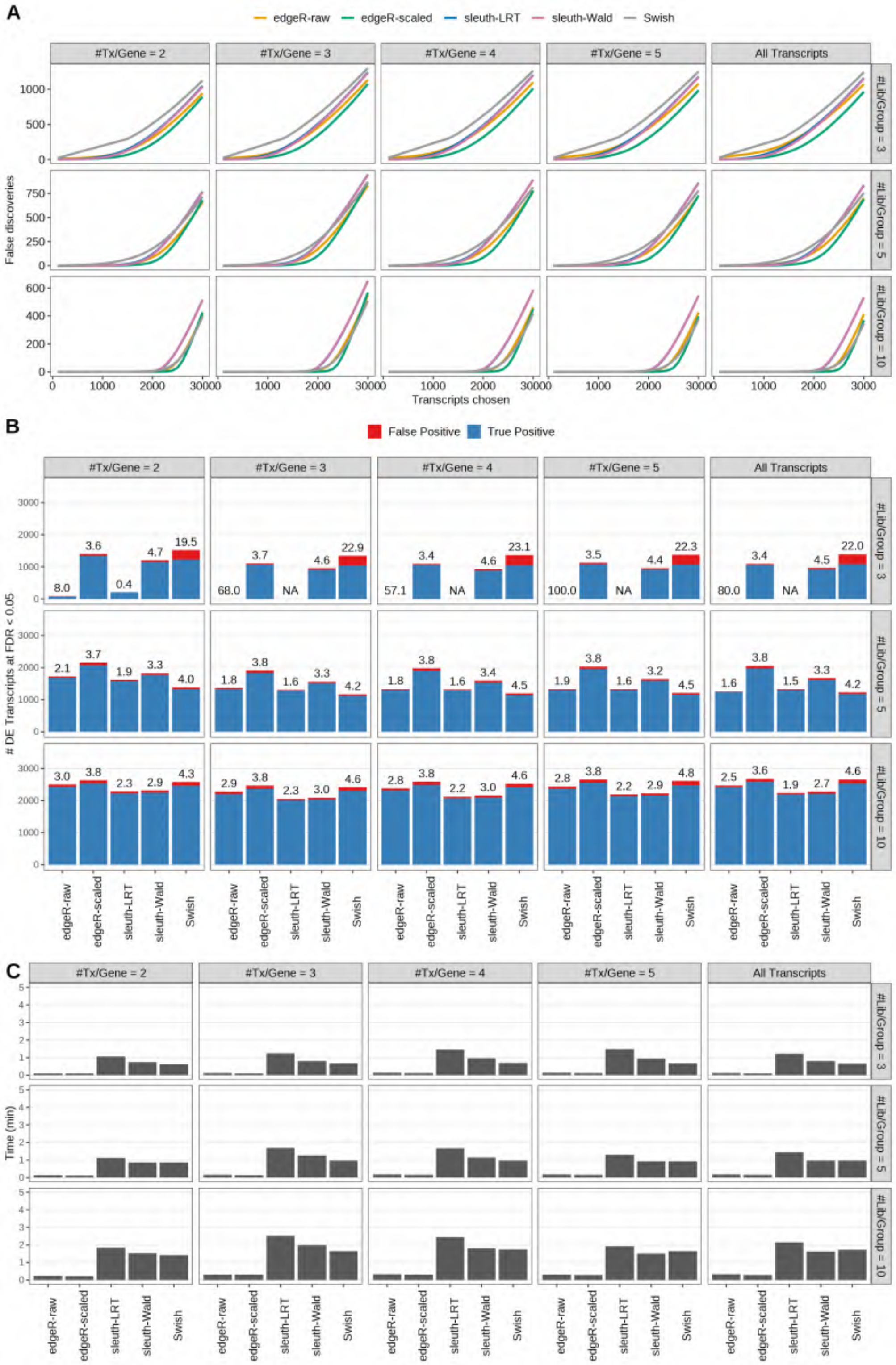

Figure S21: Simulation results. Scenario with mm39 genome, 50bp single-end reads quantified with Salmon, and unbalanced libraries. (A) Average number of false discoveries as a function of the number of chosen transcripts. (B) Average number of true (blue) and false (red) positive DE transcripts. Observed FDR annotated over bars. (C) Average computing time in minutes.

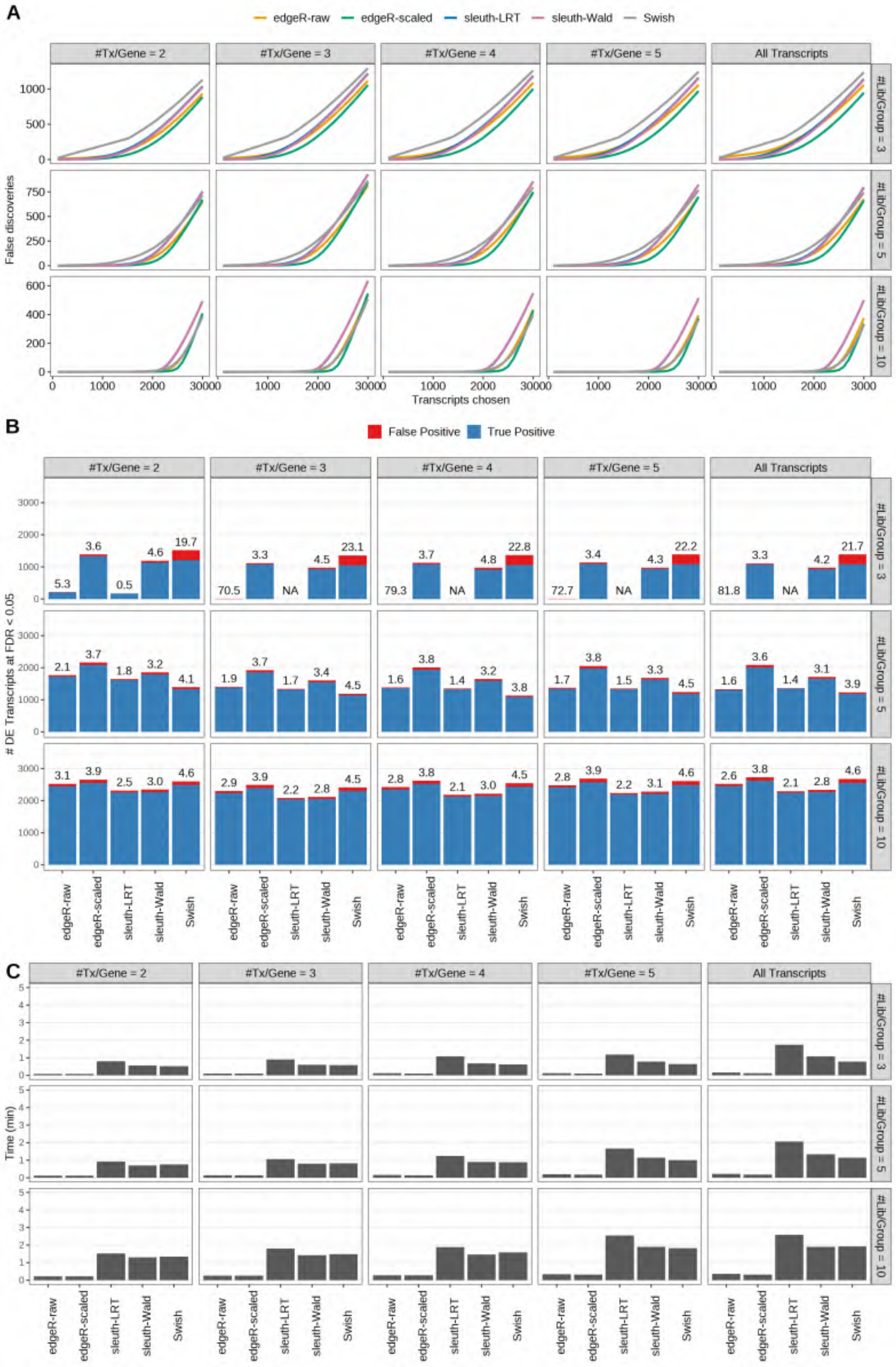

Figure S22: Simulation results. Scenario with mm39 genome, 75bp single-end reads quantified with Salmon, and unbalanced libraries. (A) Average number of false discoveries as a function of the number of chosen transcripts. (B) Average number of true (blue) and false (red) positive DE transcripts. Observed FDR annotated over bars. (C) Average computing time in minutes.

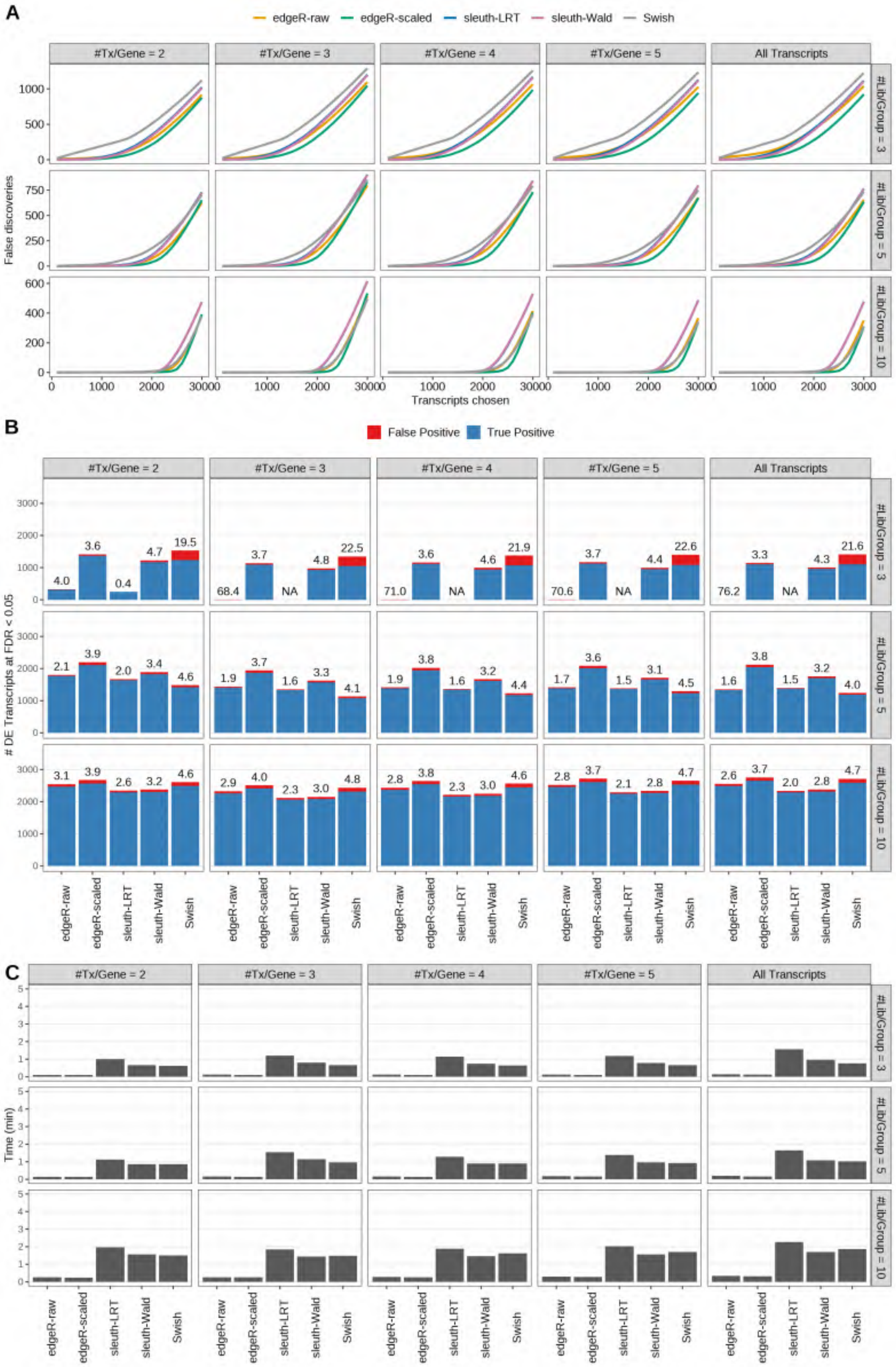

Figure S23: Simulation results. Scenario with mm39 genome, 100bp single-end reads quantified with Salmon, and unbalanced libraries. (A) Average number of false discoveries as a function of the number of chosen transcripts. (B) Average number of true (blue) and false (red) positive DE transcripts. Observed FDR annotated over bars. (C) Average computing time in minutes.

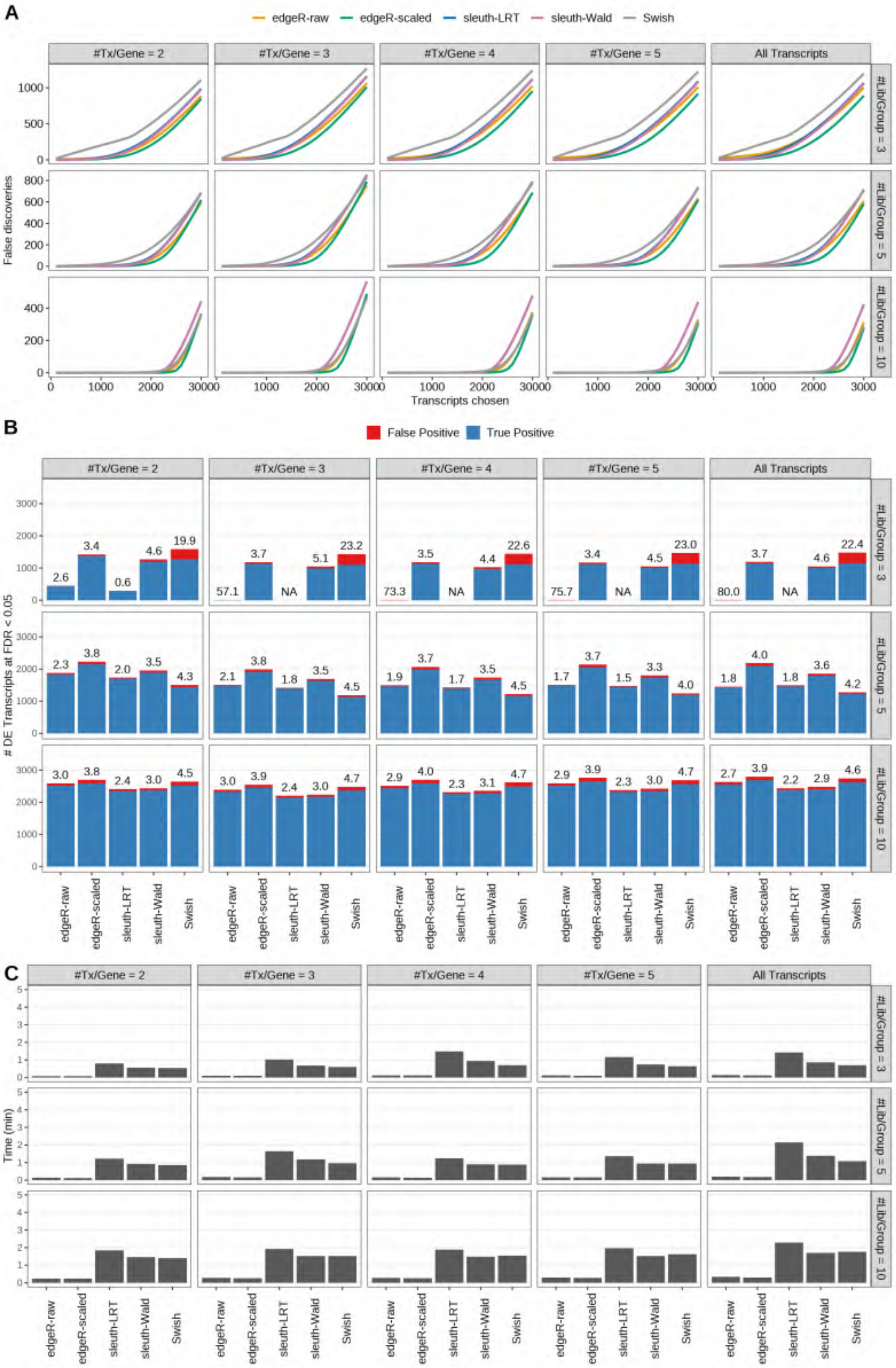

Figure S24: Simulation results. Scenario with mm39 genome, 125bp single-end reads quantified with Salmon, and unbalanced libraries. (A) Average number of false discoveries as a function of the number of chosen transcripts. (B) Average number of true (blue) and false (red) positive DE transcripts. Observed FDR annotated over bars. (C) Average computing time in minutes.

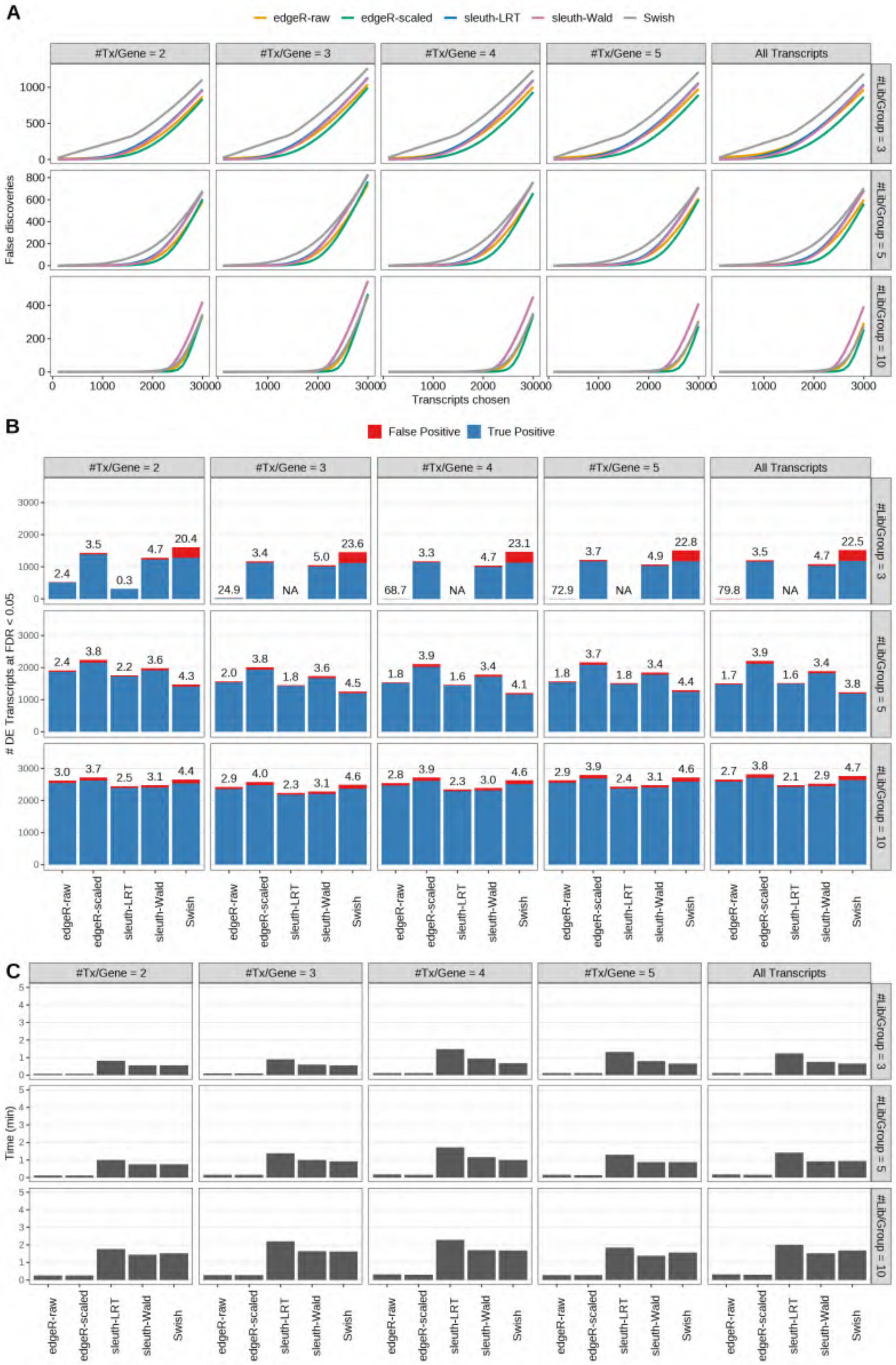

Figure S25: Simulation results. Scenario with mm39 genome, 150bp single-end reads quantified with Salmon, and unbalanced libraries. (A) Average number of false discoveries as a function of the number of chosen transcripts. (B) Average number of true (blue) and false (red) positive DE transcripts. Observed FDR annotated over bars. (C) Average computing time in minutes.

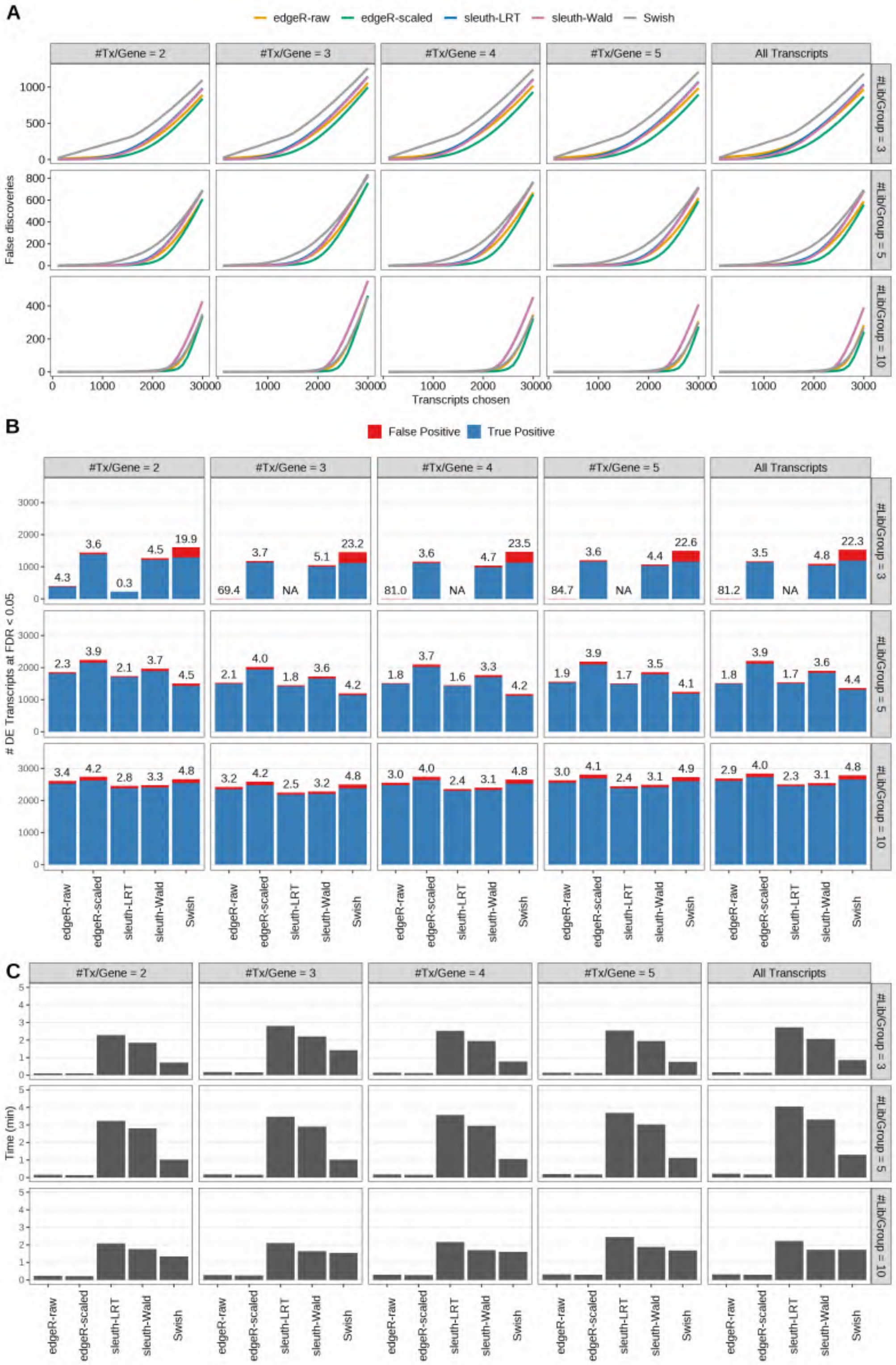

Figure S26: Simulation results. Scenario with mm39 genome, 50bp paired-end reads quantified with Salmon, and unbalanced libraries. (A) Average number of false discoveries as a function of the number of chosen transcripts. (B) Average number of true (blue) and false (red) positive DE transcripts. Observed FDR annotated over bars. (C) Average computing time in minutes.

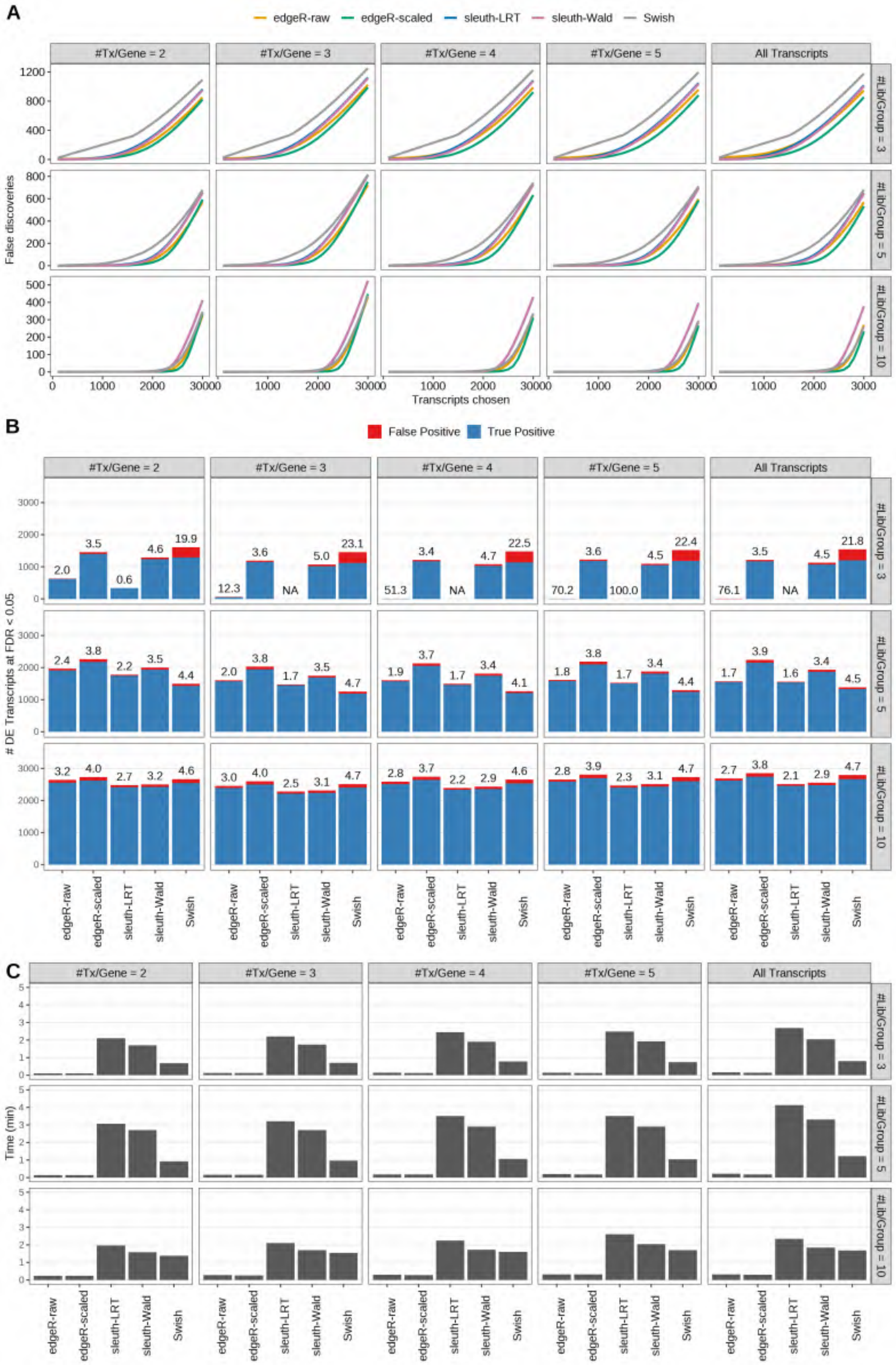

Figure S27: Simulation results. Scenario with mm39 genome, 75bp paired-end reads quantified with Salmon, and unbalanced libraries. (A) Average number of false discoveries as a function of the number of chosen transcripts. (B) Average number of true (blue) and false (red) positive DE transcripts. Observed FDR annotated over bars. (C) Average computing time in minutes.

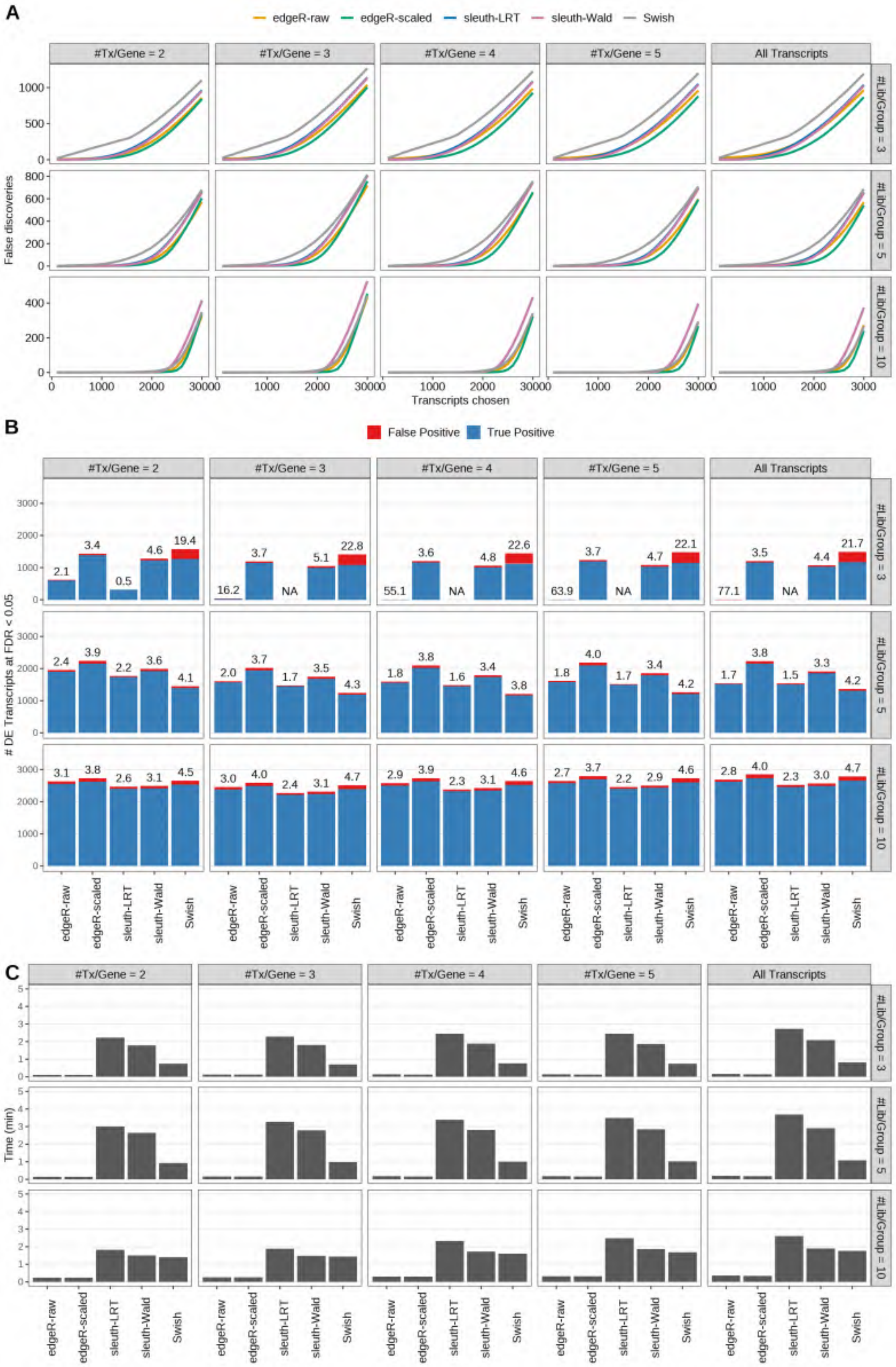

Figure S28: Simulation results. Scenario with mm39 genome, 100bp paired-end reads quantified with Salmon, and unbalanced libraries. (A) Average number of false discoveries as a function of the number of chosen transcripts. (B) Average number of true (blue) and false (red) positive DE transcripts. Observed FDR annotated over bars. (C) Average computing time in minutes.

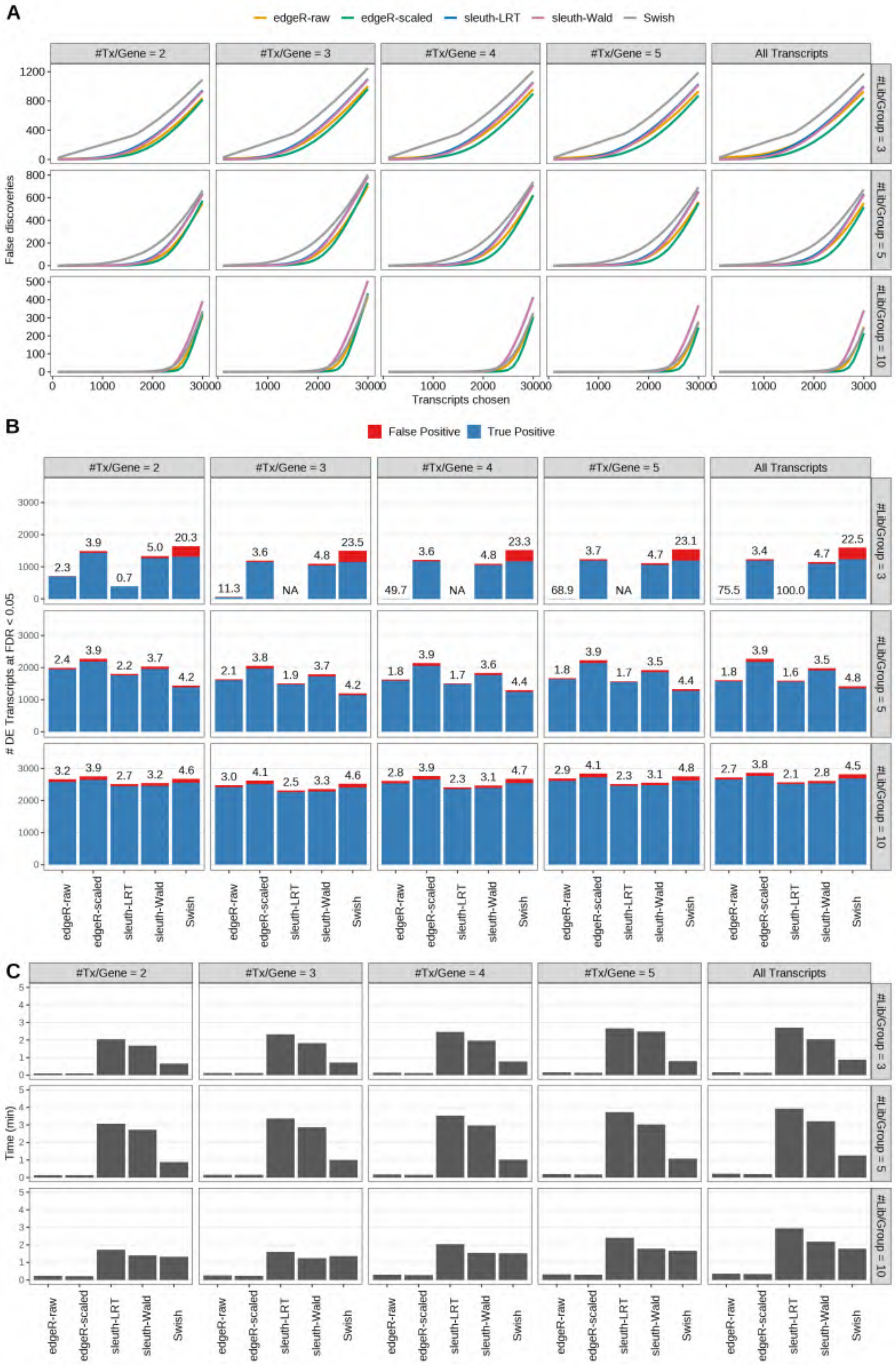

Figure S29: Simulation results. Scenario with mm39 genome, 125bp paired-end reads quantified with Salmon, and unbalanced libraries. (A) Average number of false discoveries as a function of the number of chosen transcripts. (B) Average number of true (blue) and false (red) positive DE transcripts. Observed FDR annotated over bars. (C) Average computing time in minutes.

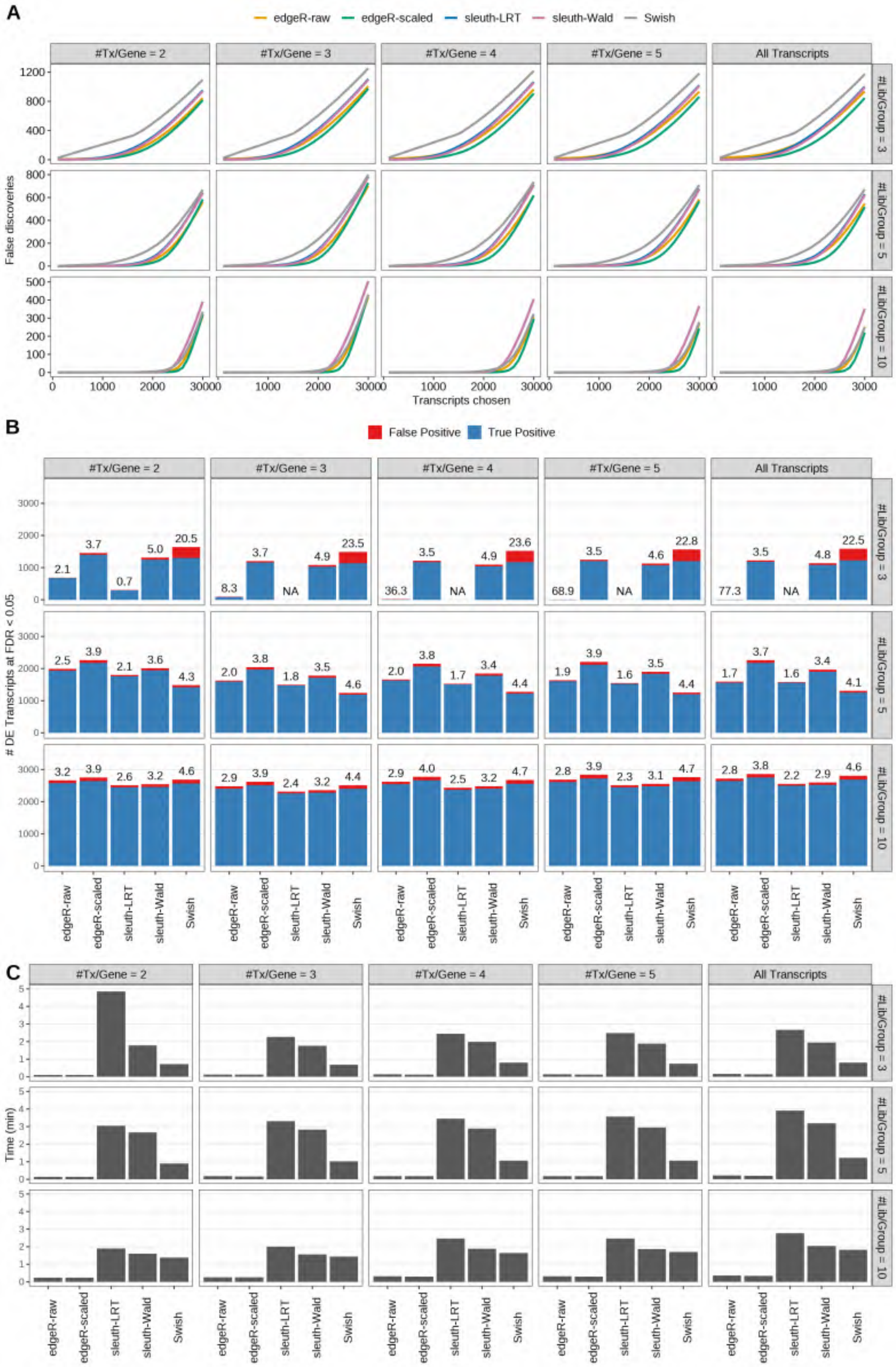

Figure S30: Simulation results. Scenario with mm39 genome, 150bp paired-end reads quantified with Salmon, and unbalanced libraries. (A) Average number of false discoveries as a function of the number of chosen transcripts. (B) Average number of true (blue) and false (red) positive DE transcripts. Observed FDR annotated over bars. (C) Average computing time in minutes.

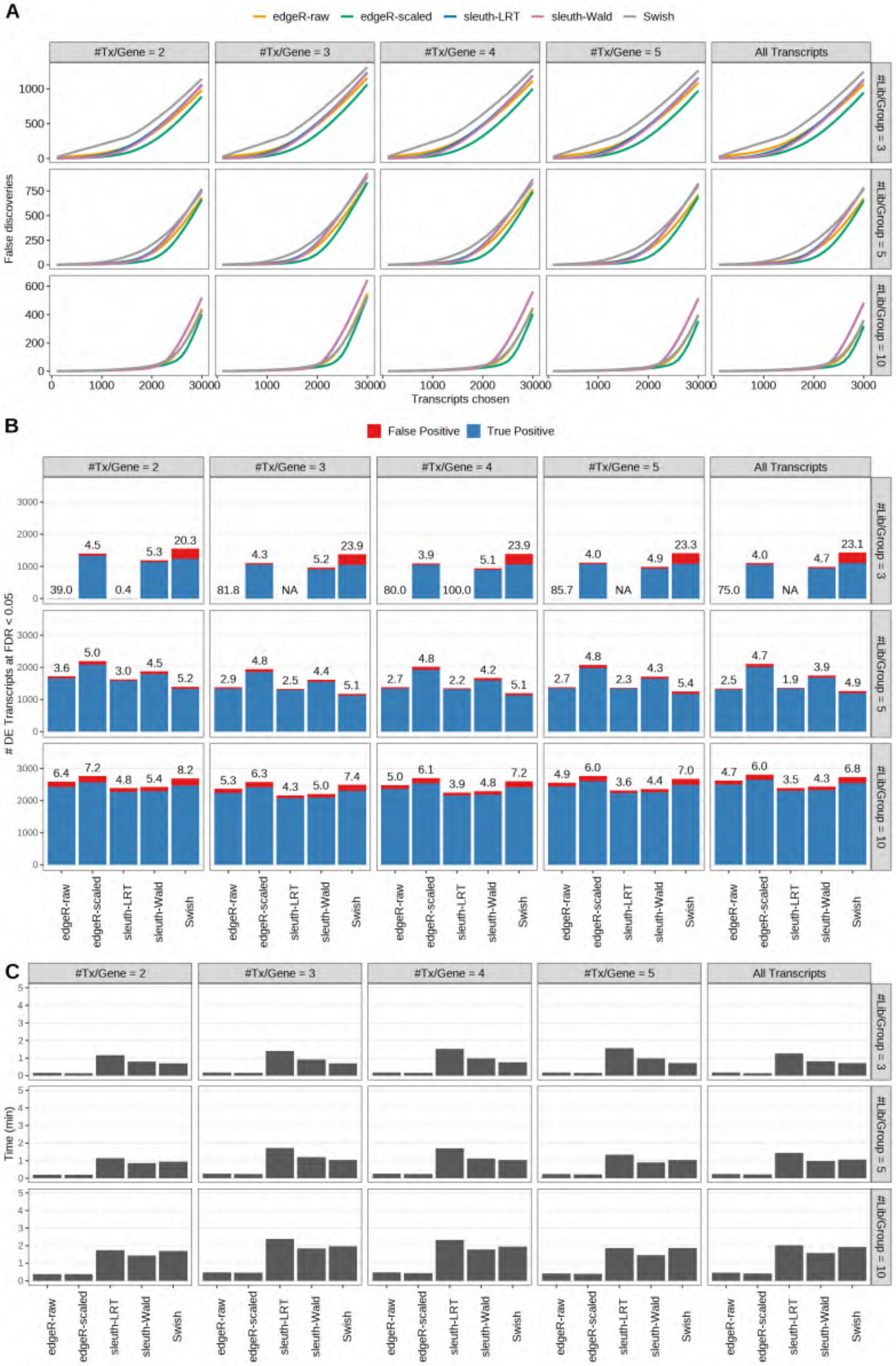

Figure S31: Simulation results. Scenario with mm39 genome, 50bp single-end reads quantified with kallisto, and unbalanced libraries. (A) Average number of false discoveries as a function of the number of chosen transcripts. (B) Average number of true (blue) and false (red) positive DE transcripts. Observed FDR annotated over bars. (C) Average computing time in minutes.

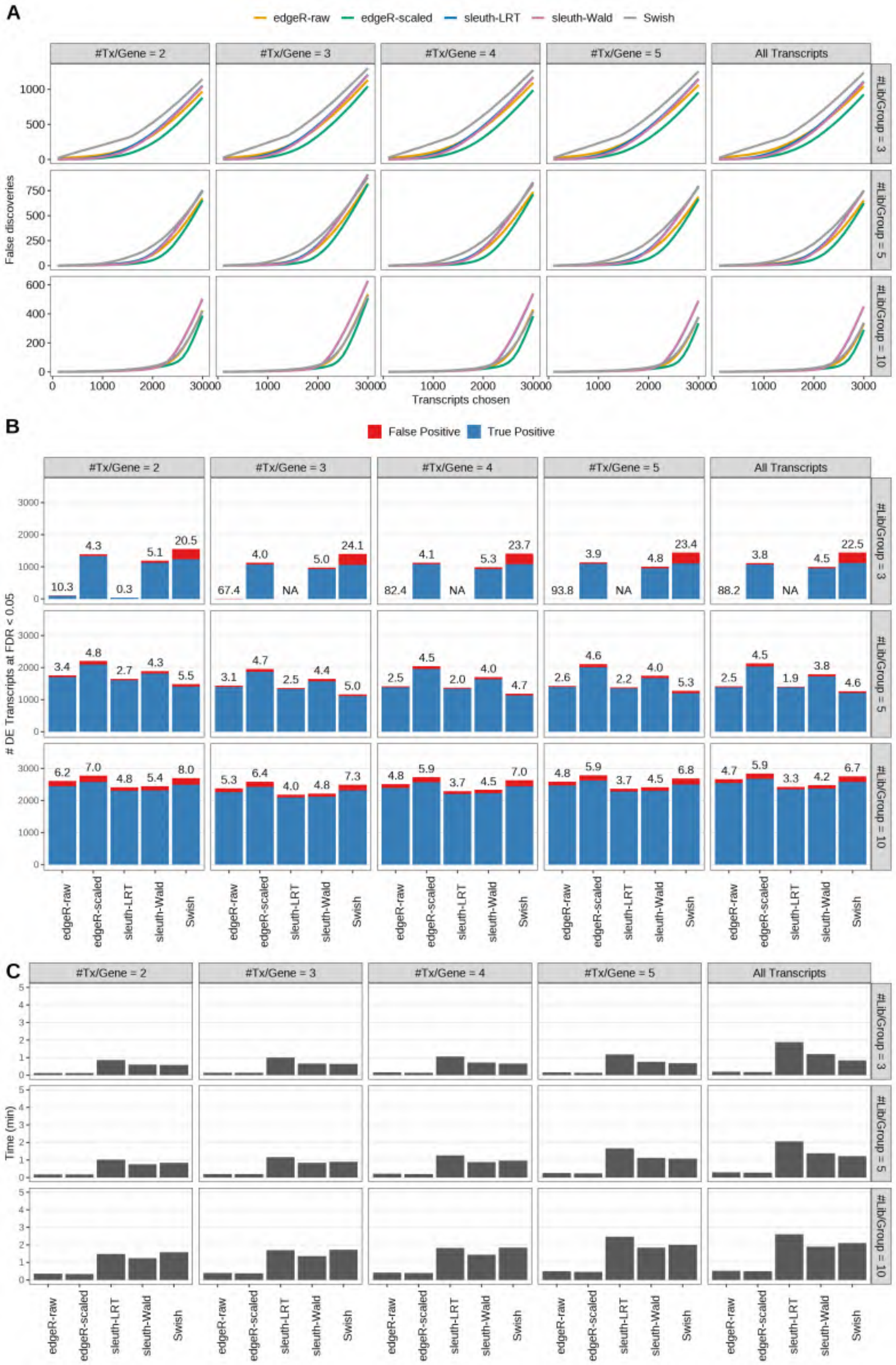

Figure S32: Simulation results. Scenario with mm39 genome, 75bp single-end reads quantified with kallisto, and unbalanced libraries. (A) Average number of false discoveries as a function of the number of chosen transcripts. (B) Average number of true (blue) and false (red) positive DE transcripts. Observed FDR annotated over bars. (C) Average computing time in minutes.

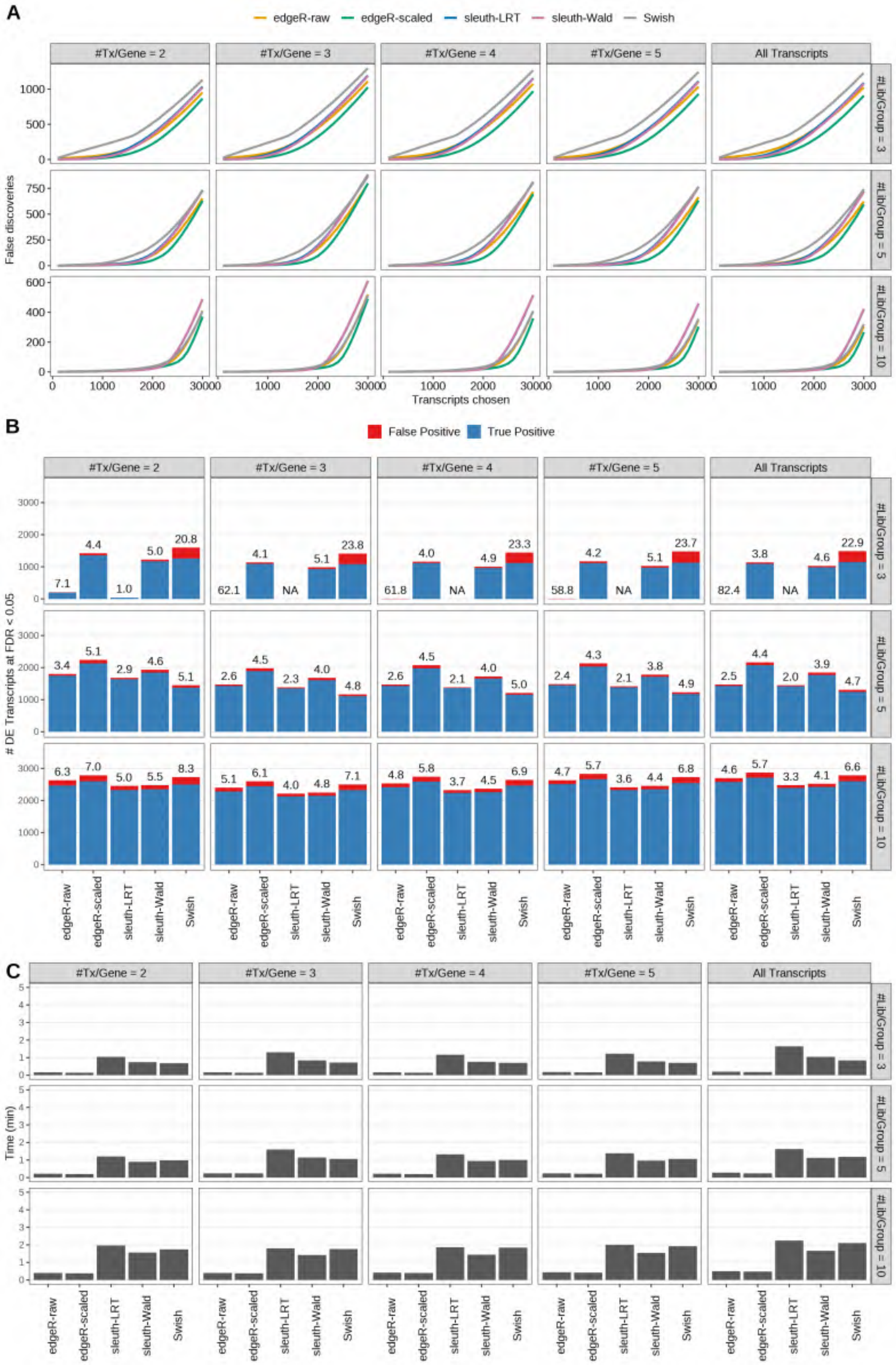

Figure S33: Simulation results. Scenario with mm39 genome, 100bp single-end reads quantified with kallisto, and unbalanced libraries. (A) Average number of false discoveries as a function of the number of chosen transcripts. (B) Average number of true (blue) and false (red) positive DE transcripts. Observed FDR annotated over bars. (C) Average computing time in minutes.

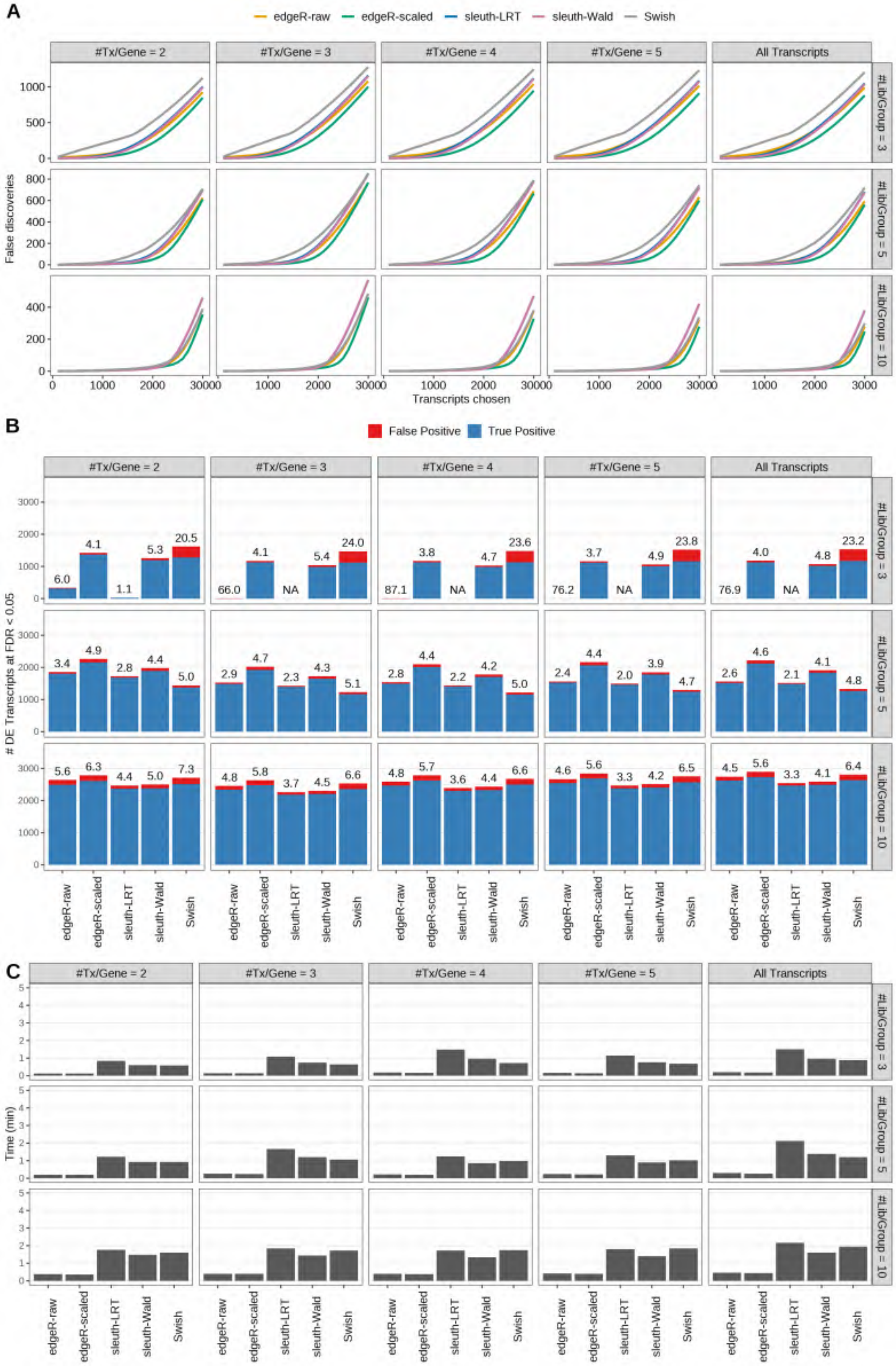

Figure S34: Simulation results. Scenario with mm39 genome, 125bp single-end reads quantified with kallisto, and unbalanced libraries. (A) Average number of false discoveries as a function of the number of chosen transcripts. (B) Average number of true (blue) and false (red) positive DE transcripts. Observed FDR annotated over bars. (C) Average computing time in minutes.

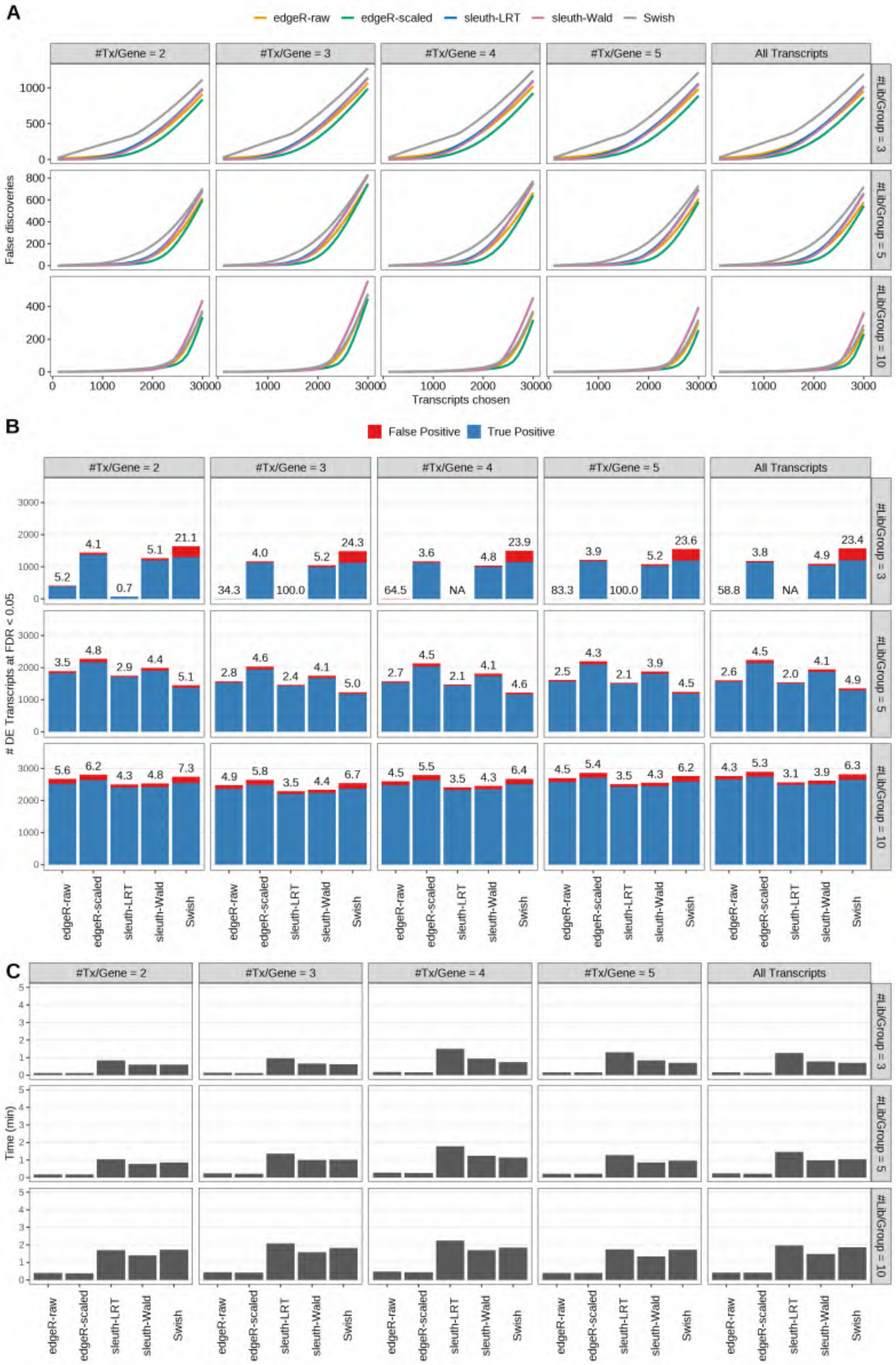

Figure S35: Simulation results. Scenario with mm39 genome, 150bp single-end reads quantified with kallisto, and unbalanced libraries. (A) Average number of false discoveries as a function of the number of chosen transcripts. (B) Average number of true (blue) and false (red) positive DE transcripts. Observed FDR annotated over bars. (C) Average computing time in minutes.

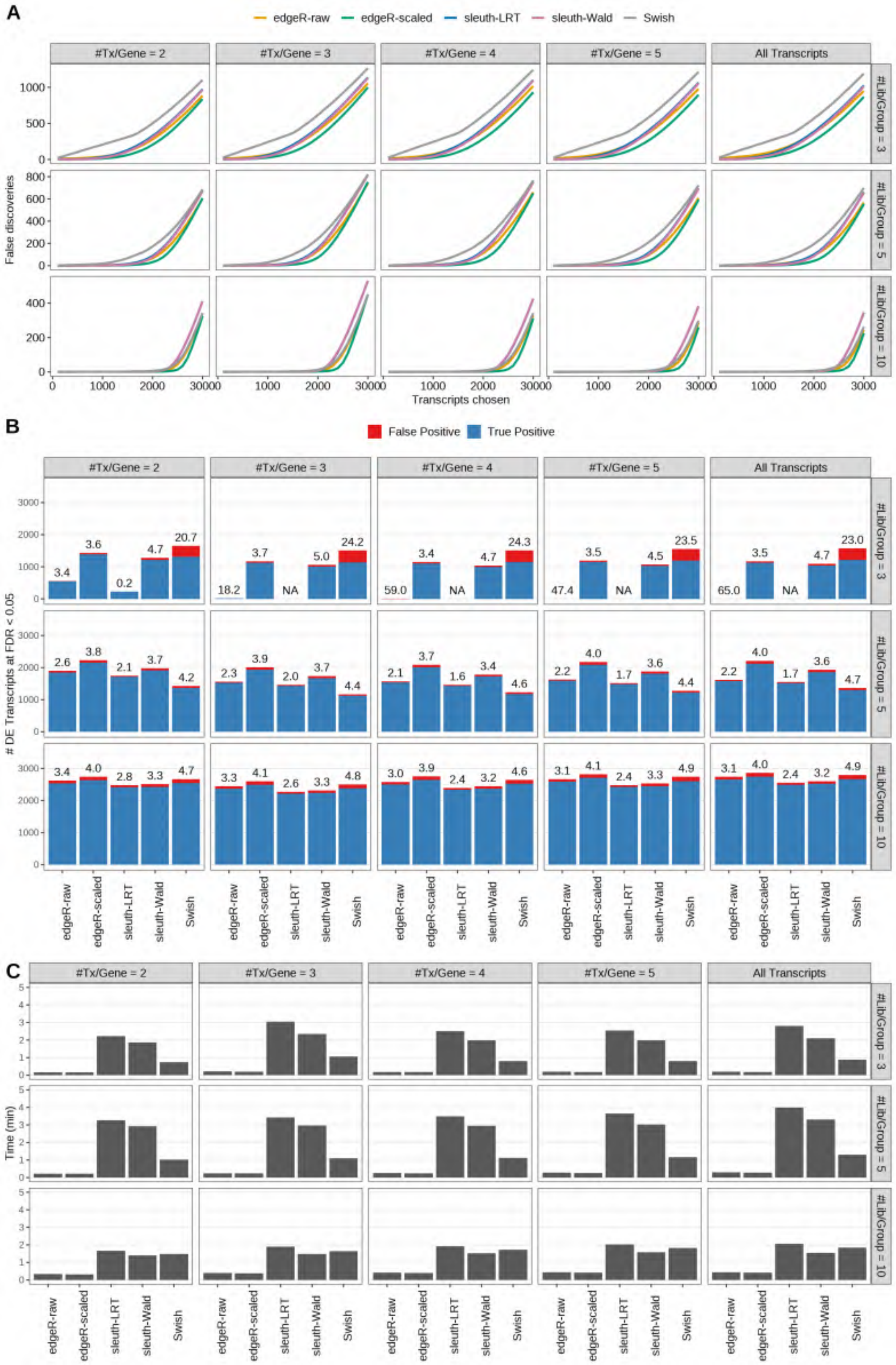

Figure S36: Simulation results. Scenario with mm39 genome, 50bp paired-end reads quantified with kallisto, and unbalanced libraries. (A) Average number of false discoveries as a function of the number of chosen transcripts. (B) Average number of true (blue) and false (red) positive DE transcripts. Observed FDR annotated over bars. (C) Average computing time in minutes.

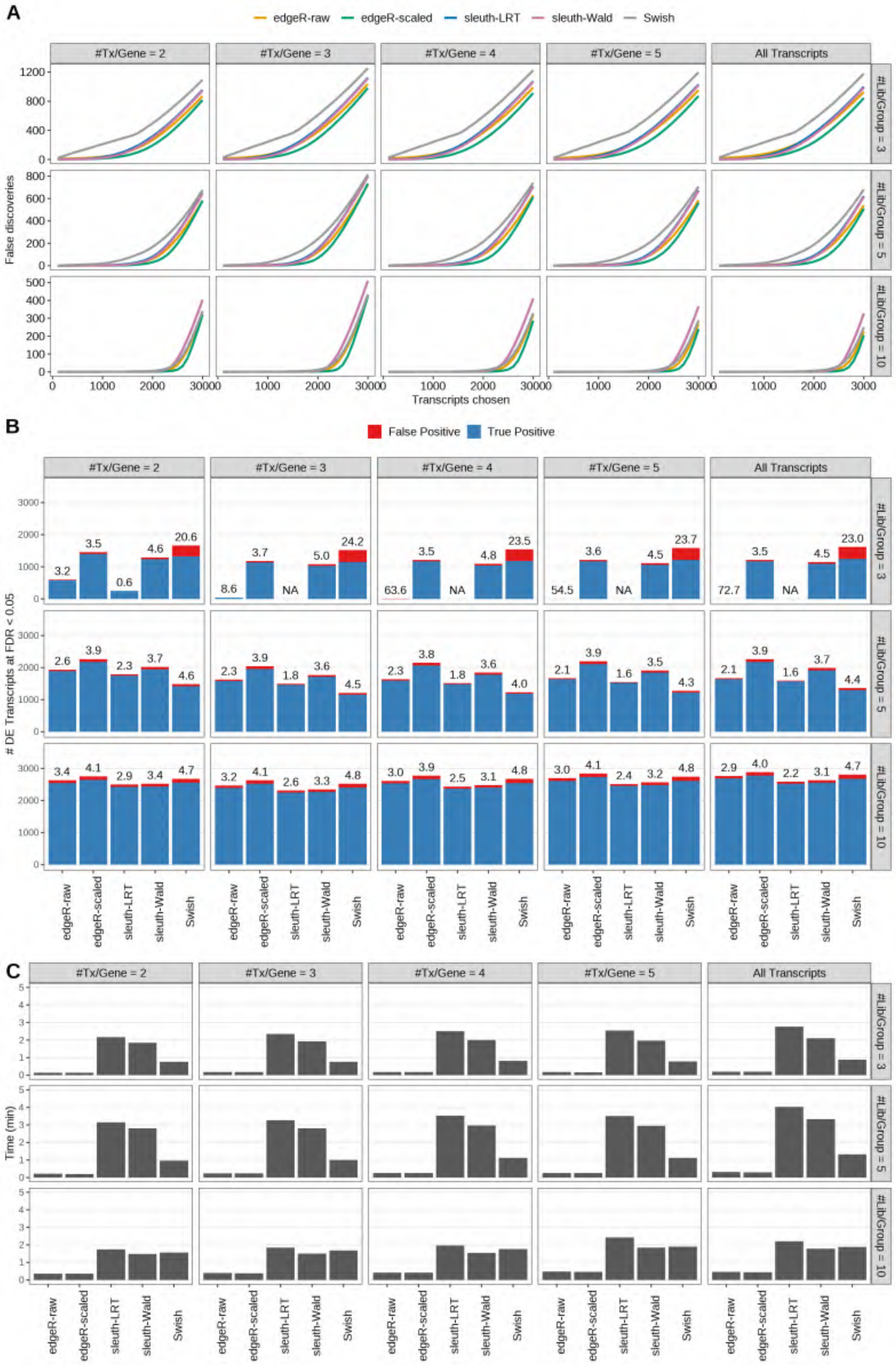

Figure S37: Simulation results. Scenario with mm39 genome, 75bp paired-end reads quantified with kallisto, and unbalanced libraries. (A) Average number of false discoveries as a function of the number of chosen transcripts. (B) Average number of true (blue) and false (red) positive DE transcripts. Observed FDR annotated over bars. (C) Average computing time in minutes.

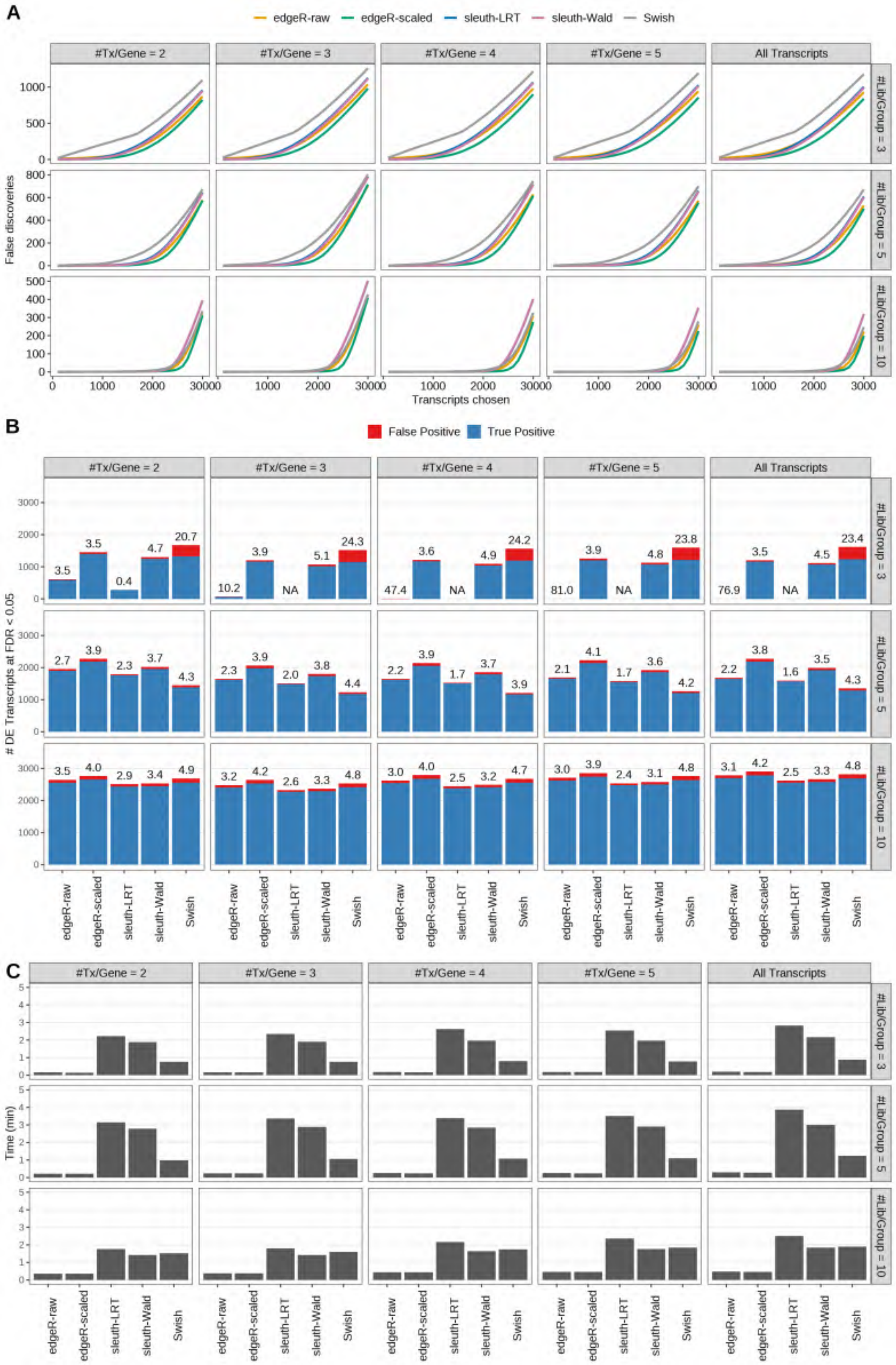

Figure S38: Simulation results. Scenario with mm39 genome, 100bp paired-end reads quantified with kallisto, and unbalanced libraries. (A) Average number of false discoveries as a function of the number of chosen transcripts. (B) Average number of true (blue) and false (red) positive DE transcripts. Observed FDR annotated over bars. (C) Average computing time in minutes.

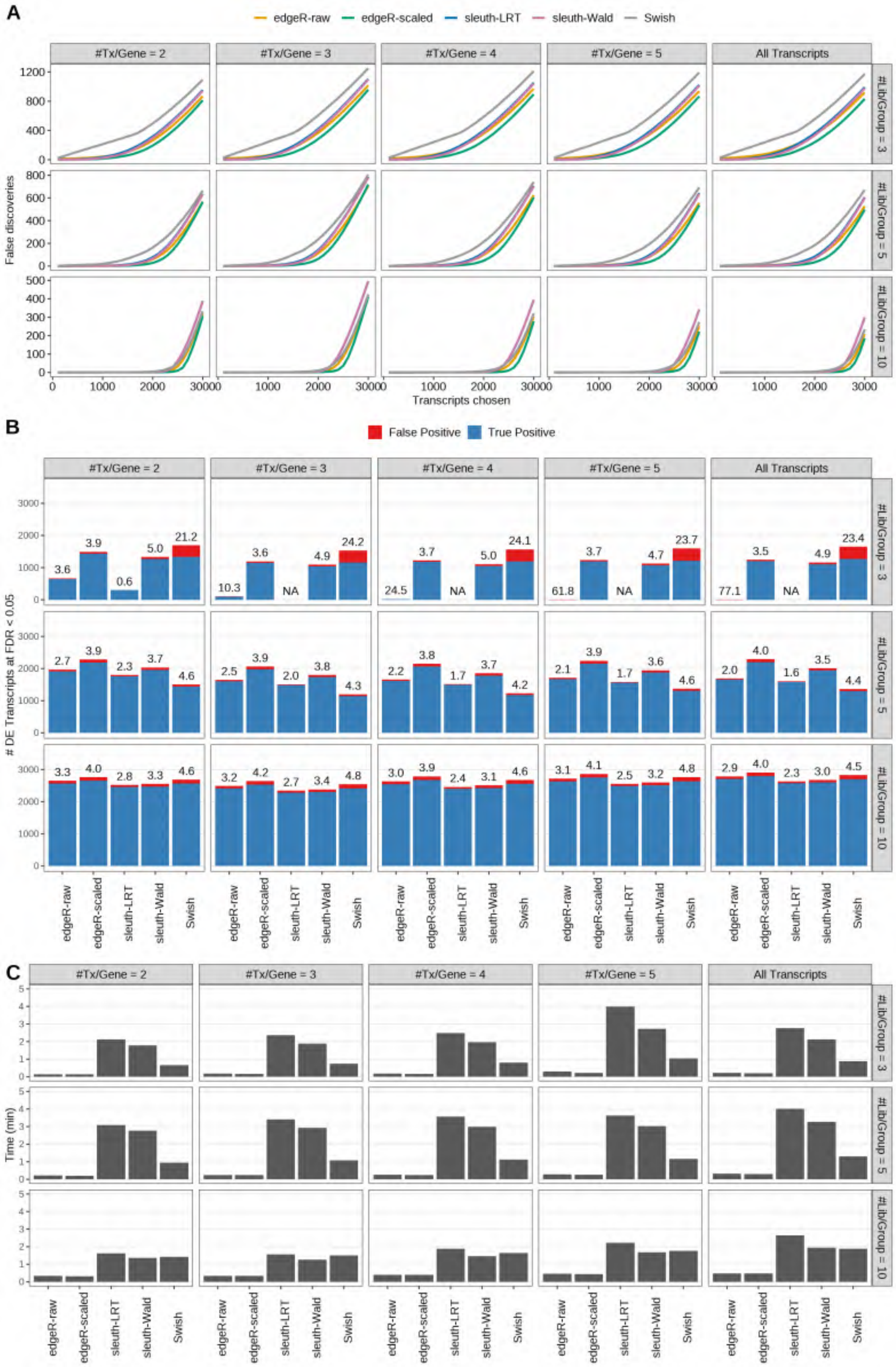

Figure S39: Simulation results. Scenario with mm39 genome, 125bp paired-end reads quantified with kallisto, and unbalanced libraries. (A) Average number of false discoveries as a function of the number of chosen transcripts. (B) Average number of true (blue) and false (red) positive DE transcripts. Observed FDR annotated over bars. (C) Average computing time in minutes.

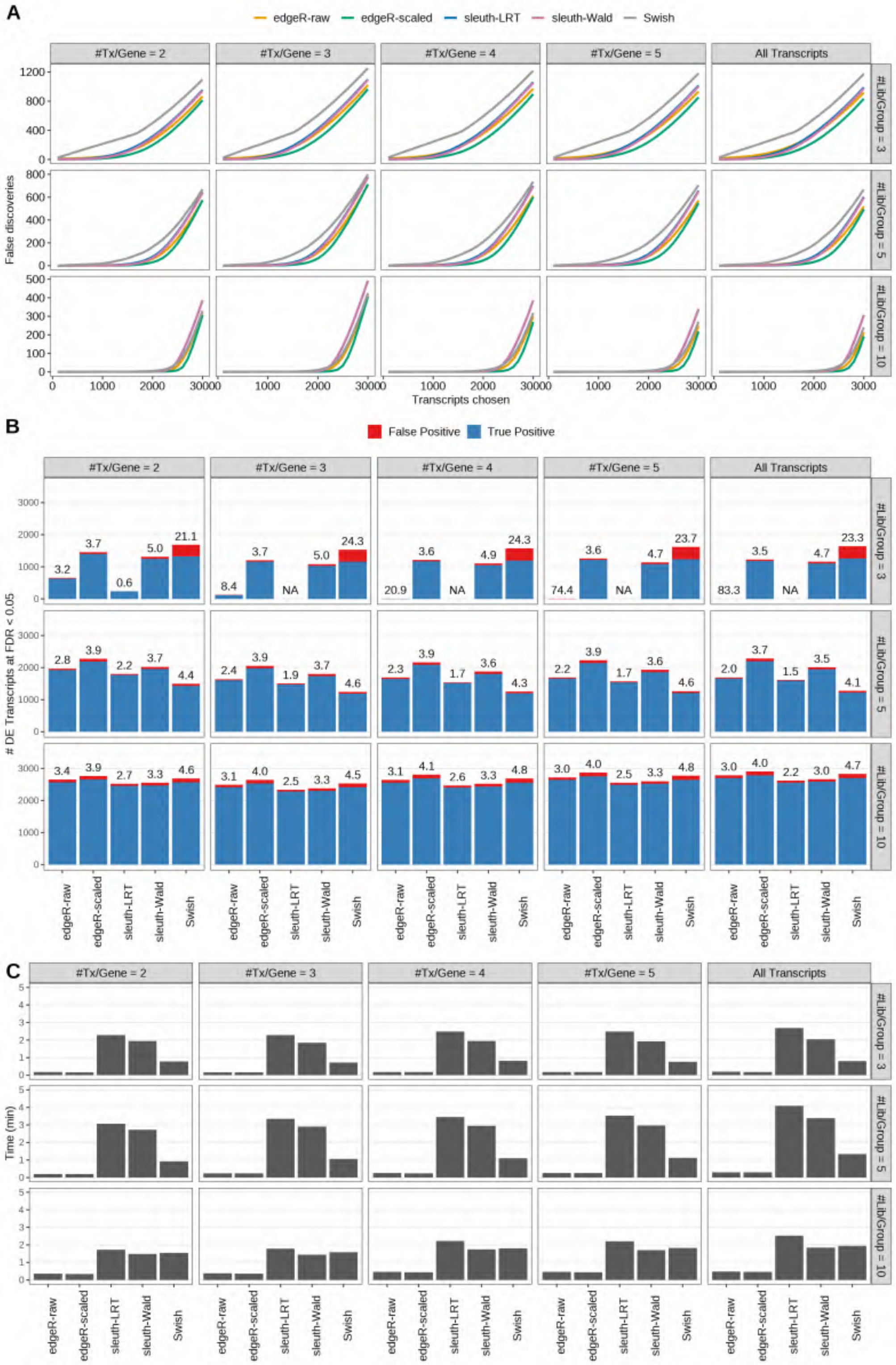

Figure S40: Simulation results. Scenario with mm39 genome, 150bp paired-end reads quantified with kallisto, and unbalanced libraries. (A) Average number of false discoveries as a function of the number of chosen transcripts. (B) Average number of true (blue) and false (red) positive DE transcripts. Observed FDR annotated over bars. (C) Average computing time in minutes.

Table S1: Simulation results - observed power and false discovery rate for different read types and read Lengths, averaged over 20 simulations. Scenario with mm39 genome, maximum of 2 transcripts/gene expressed, and reads quantified with Salmon. Library size shown in million reads (M) with 25/100 indicating library sizes alternating between 25M and 100M across replicates. Read lengths are shown in base pairs (bp). Red color indicates observed FDR values greater than the nominal 0.05. Blue color indicates the most powerful method that controls the FDR at the nominal level for a given scenario (row). Empty cells indicate cases in which a method failed to call any transcript as DE.

| | | | | Power | | | | | False Discovery Rate | | | | |
|---|---|---|---|---|---|---|---|---|---|---|---|---|---|
| Read | Samples/Group | Library Size | Read Length | edgeR-raw | edgeR-scaled | sleuth-LRT | sleuth-Wald | Swish | edgeR-raw | edgeR-scaled | sleuth-LRT | sleuth-Wald | Swish |
| paired-end | 3 | 50M | 50bp | 0.211 | 0.513 | 0.171 | 0.461 | 0.483 | 0.057 | 0.041 | 0.011 | 0.057 | 0.215 |
| | 3 | 50M | 75bp | 0.304 | 0.522 | 0.197 | 0.475 | 0.486 | 0.028 | 0.039 | 0.011 | 0.054 | 0.211 |
| | 3 | 50M | 100bp | 0.298 | 0.517 | 0.189 | 0.467 | 0.476 | 0.030 | 0.039 | 0.011 | 0.054 | 0.208 |
| | 3 | 50M | 125bp | 0.326 | 0.522 | 0.199 | 0.477 | 0.492 | 0.025 | 0.038 | 0.009 | 0.053 | 0.215 |
| | 3 | 50M | 150bp | 0.320 | 0.525 | 0.204 | 0.478 | 0.494 | 0.024 | 0.038 | 0.011 | 0.055 | 0.213 |
| | 3 | 25/100M | 50bp | 0.127 | 0.464 | 0.076 | 0.405 | 0.428 | 0.043 | 0.036 | 0.003 | 0.045 | 0.199 |
| | 3 | 25/100M | 75bp | 0.206 | 0.466 | 0.111 | 0.411 | 0.430 | 0.020 | 0.035 | 0.006 | 0.046 | 0.199 |
| | 3 | 25/100M | 100bp | 0.201 | 0.463 | 0.104 | 0.407 | 0.423 | 0.021 | 0.034 | 0.005 | 0.046 | 0.194 |
| | 3 | 25/100M | 125bp | 0.232 | 0.478 | 0.129 | 0.424 | 0.437 | 0.023 | 0.039 | 0.007 | 0.050 | 0.203 |
| | 3 | 25/100M | 150bp | 0.224 | 0.467 | 0.099 | 0.414 | 0.434 | 0.021 | 0.037 | 0.007 | 0.050 | 0.205 |
| | 5 | 50M | 50bp | 0.655 | 0.756 | 0.626 | 0.688 | 0.500 | 0.029 | 0.044 | 0.026 | 0.042 | 0.043 |
| | 5 | 50M | 75bp | 0.687 | 0.761 | 0.636 | 0.697 | 0.503 | 0.028 | 0.041 | 0.025 | 0.040 | 0.044 |
| | 5 | 50M | 100bp | 0.687 | 0.758 | 0.636 | 0.696 | 0.518 | 0.026 | 0.040 | 0.024 | 0.038 | 0.045 |
| | 5 | 50M | 125bp | 0.701 | 0.766 | 0.648 | 0.707 | 0.485 | 0.027 | 0.039 | 0.024 | 0.038 | 0.038 |
| | 5 | 50M | 150bp | 0.693 | 0.761 | 0.643 | 0.703 | 0.505 | 0.026 | 0.039 | 0.024 | 0.038 | 0.044 |
| | 5 | 25/100M | 50bp | 0.604 | 0.716 | 0.567 | 0.630 | 0.478 | 0.023 | 0.039 | 0.021 | 0.037 | 0.045 |
| | 5 | 25/100M | 75bp | 0.638 | 0.724 | 0.581 | 0.645 | 0.474 | 0.024 | 0.038 | 0.022 | 0.035 | 0.044 |
| | 5 | 25/100M | 100bp | 0.635 | 0.719 | 0.578 | 0.639 | 0.465 | 0.024 | 0.039 | 0.022 | 0.036 | 0.041 |
| | 5 | 25/100M | 125bp | 0.647 | 0.728 | 0.587 | 0.653 | 0.461 | 0.024 | 0.039 | 0.022 | 0.037 | 0.042 |
| | 5 | 25/100M | 150bp | 0.645 | 0.726 | 0.587 | 0.648 | 0.473 | 0.025 | 0.039 | 0.021 | 0.036 | 0.043 |
| | 10 | 50M | 50bp | 0.862 | 0.868 | 0.837 | 0.841 | 0.861 | 0.037 | 0.045 | 0.033 | 0.039 | 0.052 |
| | 10 | 50M | 75bp | 0.872 | 0.871 | 0.844 | 0.850 | 0.861 | 0.034 | 0.042 | 0.030 | 0.036 | 0.044 |
| | 10 | 50M | 100bp | 0.868 | 0.867 | 0.843 | 0.848 | 0.858 | 0.033 | 0.039 | 0.028 | 0.033 | 0.045 |
| | 10 | 50M | 125bp | 0.878 | 0.875 | 0.853 | 0.858 | 0.866 | 0.032 | 0.040 | 0.028 | 0.034 | 0.043 |
| | 10 | 50M | 150bp | 0.879 | 0.877 | 0.855 | 0.861 | 0.867 | 0.034 | 0.041 | 0.029 | 0.035 | 0.045 |
| | 10 | 25/100M | 50bp | 0.840 | 0.873 | 0.795 | 0.799 | 0.846 | 0.034 | 0.042 | 0.028 | 0.033 | 0.048 |
| | 10 | 25/100M | 75bp | 0.852 | 0.874 | 0.802 | 0.808 | 0.848 | 0.032 | 0.040 | 0.027 | 0.032 | 0.046 |
| | 10 | 25/100M | 100bp | 0.851 | 0.874 | 0.799 | 0.804 | 0.844 | 0.031 | 0.038 | 0.026 | 0.031 | 0.045 |
| | 10 | 25/100M | 125bp | 0.859 | 0.880 | 0.813 | 0.819 | 0.852 | 0.032 | 0.039 | 0.027 | 0.032 | 0.046 |
| | 10 | 25/100M | 150bp | 0.860 | 0.880 | 0.814 | 0.819 | 0.853 | 0.032 | 0.039 | 0.026 | 0.032 | 0.046 |

Table S1: Simulation results - observed power and false discovery rate for different read types and read Lengths, averaged over 20 simulations. Scenario with mm39 genome, maximum of 2 transcripts/gene expressed, and reads quantified with Salmon. Library size shown in million reads (M) with 25/100 indicating library sizes alternating between 25M and 100M across replicates. Read lengths are shown in base pairs (bp). Red color indicates observed FDR values greater than the nominal 0.05. Blue color indicates the most powerful method that controls the FDR at the nominal level for a given scenario (row). Empty cells indicate cases in which a method failed to call any transcript as DE. *(continued)*

| | | | | Power | | | | | False Discovery Rate | | | | |
|---|---|---|---|---|---|---|---|---|---|---|---|---|---|
| Read | Samples/Group | Library Size | Read Length | edgeR-raw | edgeR-scaled | sleuth-LRT | sleuth-Wald | Swish | edgeR-raw | edgeR-scaled | sleuth-LRT | sleuth-Wald | Swish |
| single-end | 3 | 50M | 50bp | 0.153 | 0.492 | 0.158 | 0.438 | 0.458 | 0.056 | 0.038 | 0.009 | 0.053 | 0.208 |
| | 3 | 50M | 75bp | 0.191 | 0.503 | 0.172 | 0.445 | 0.462 | 0.044 | 0.040 | 0.010 | 0.052 | 0.211 |
| | 3 | 50M | 100bp | 0.196 | 0.506 | 0.158 | 0.449 | 0.463 | 0.045 | 0.041 | 0.011 | 0.054 | 0.206 |
| | 3 | 50M | 125bp | 0.243 | 0.515 | 0.190 | 0.461 | 0.479 | 0.032 | 0.038 | 0.009 | 0.053 | 0.210 |
| | 3 | 50M | 150bp | 0.275 | 0.512 | 0.194 | 0.468 | 0.482 | 0.030 | 0.038 | 0.010 | 0.053 | 0.217 |
| | 3 | 25/100M | 50bp | 0.025 | 0.449 | 0.067 | 0.382 | 0.407 | 0.080 | 0.036 | 0.004 | 0.047 | 0.195 |
| | 3 | 25/100M | 75bp | 0.067 | 0.446 | 0.058 | 0.380 | 0.406 | 0.053 | 0.036 | 0.005 | 0.046 | 0.197 |
| | 3 | 25/100M | 100bp | 0.104 | 0.455 | 0.081 | 0.391 | 0.410 | 0.040 | 0.036 | 0.004 | 0.047 | 0.195 |
| | 3 | 25/100M | 125bp | 0.146 | 0.459 | 0.097 | 0.403 | 0.424 | 0.026 | 0.034 | 0.006 | 0.046 | 0.199 |
| | 3 | 25/100M | 150bp | 0.171 | 0.463 | 0.105 | 0.408 | 0.428 | 0.024 | 0.035 | 0.003 | 0.047 | 0.204 |
| | 5 | 50M | 50bp | 0.621 | 0.731 | 0.596 | 0.654 | 0.473 | 0.025 | 0.039 | 0.023 | 0.037 | 0.040 |
| | 5 | 50M | 75bp | 0.634 | 0.742 | 0.604 | 0.663 | 0.497 | 0.025 | 0.041 | 0.024 | 0.039 | 0.044 |
| | 5 | 50M | 100bp | 0.641 | 0.743 | 0.609 | 0.669 | 0.488 | 0.025 | 0.042 | 0.024 | 0.040 | 0.042 |
| | 5 | 50M | 125bp | 0.666 | 0.753 | 0.630 | 0.689 | 0.489 | 0.025 | 0.040 | 0.023 | 0.039 | 0.041 |
| | 5 | 50M | 150bp | 0.675 | 0.757 | 0.635 | 0.694 | 0.501 | 0.026 | 0.039 | 0.024 | 0.038 | 0.042 |
| | 5 | 25/100M | 50bp | 0.562 | 0.692 | 0.528 | 0.590 | 0.444 | 0.021 | 0.037 | 0.019 | 0.033 | 0.040 |
| | 5 | 25/100M | 75bp | 0.577 | 0.694 | 0.539 | 0.598 | 0.445 | 0.021 | 0.037 | 0.018 | 0.032 | 0.041 |
| | 5 | 25/100M | 100bp | 0.589 | 0.704 | 0.547 | 0.608 | 0.472 | 0.021 | 0.039 | 0.020 | 0.034 | 0.046 |
| | 5 | 25/100M | 125bp | 0.611 | 0.716 | 0.566 | 0.630 | 0.480 | 0.023 | 0.038 | 0.020 | 0.035 | 0.043 |
| | 5 | 25/100M | 150bp | 0.623 | 0.719 | 0.574 | 0.636 | 0.469 | 0.024 | 0.038 | 0.022 | 0.036 | 0.043 |
| | 10 | 50M | 50bp | 0.833 | 0.838 | 0.796 | 0.800 | 0.839 | 0.033 | 0.041 | 0.029 | 0.034 | 0.047 |
| | 10 | 50M | 75bp | 0.840 | 0.842 | 0.805 | 0.810 | 0.840 | 0.033 | 0.042 | 0.028 | 0.034 | 0.045 |
| | 10 | 50M | 100bp | 0.845 | 0.849 | 0.813 | 0.817 | 0.844 | 0.034 | 0.043 | 0.030 | 0.036 | 0.046 |
| | 10 | 50M | 125bp | 0.857 | 0.859 | 0.830 | 0.834 | 0.852 | 0.034 | 0.041 | 0.028 | 0.034 | 0.044 |
| | 10 | 50M | 150bp | 0.864 | 0.864 | 0.839 | 0.844 | 0.856 | 0.034 | 0.042 | 0.029 | 0.035 | 0.042 |
| | 10 | 25/100M | 50bp | 0.807 | 0.844 | 0.743 | 0.747 | 0.822 | 0.030 | 0.038 | 0.023 | 0.029 | 0.043 |
| | 10 | 25/100M | 75bp | 0.816 | 0.850 | 0.753 | 0.758 | 0.828 | 0.031 | 0.039 | 0.025 | 0.030 | 0.046 |
| | 10 | 25/100M | 100bp | 0.823 | 0.856 | 0.762 | 0.767 | 0.829 | 0.031 | 0.039 | 0.026 | 0.032 | 0.046 |
| | 10 | 25/100M | 125bp | 0.837 | 0.864 | 0.783 | 0.788 | 0.840 | 0.030 | 0.038 | 0.024 | 0.030 | 0.045 |
| | 10 | 25/100M | 150bp | 0.847 | 0.872 | 0.796 | 0.800 | 0.846 | 0.030 | 0.037 | 0.025 | 0.031 | 0.044 |

Table S2: Simulation results - observed power and false discovery rate for different read types and read Lengths, averaged over 20 simulations. Scenario with mm39 genome, maximum of 2 transcripts/gene expressed, and reads quantified with kallisto. Library size shown in million reads (M) with 25/100 indicating library sizes alternating between 25M and 100M across replicates. Read lengths are shown in base pairs (bp). Red color indicates observed FDR values greater than the nominal 0.05. Blue color indicates the most powerful method that controls the FDR at the nominal level for a given scenario (row). Empty cells indicate cases in which a method failed to call any transcript as DE.

| | | | | Power | | | | | False Discovery Rate | | | | |
|---|---|---|---|---|---|---|---|---|---|---|---|---|---|
| Read | Samples/Group | Library Size | Read Length | edgeR-raw | edgeR-scaled | sleuth-LRT | sleuth-Wald | Swish | edgeR-raw | edgeR-scaled | sleuth-LRT | sleuth-Wald | Swish |
| paired-end | 3 | 50M | 50bp | 0.251 | 0.515 | 0.168 | 0.463 | 0.495 | 0.053 | 0.041 | 0.010 | 0.055 | 0.223 |
| | 3 | 50M | 75bp | 0.266 | 0.521 | 0.181 | 0.473 | 0.500 | 0.049 | 0.038 | 0.011 | 0.055 | 0.222 |
| | 3 | 50M | 100bp | 0.264 | 0.521 | 0.178 | 0.468 | 0.498 | 0.053 | 0.040 | 0.011 | 0.054 | 0.225 |
| | 3 | 50M | 125bp | 0.284 | 0.524 | 0.181 | 0.473 | 0.501 | 0.047 | 0.040 | 0.009 | 0.053 | 0.225 |
| | 3 | 50M | 150bp | 0.289 | 0.527 | 0.190 | 0.476 | 0.505 | 0.043 | 0.039 | 0.010 | 0.054 | 0.224 |
| | 3 | 25/100M | 50bp | 0.180 | 0.462 | 0.075 | 0.406 | 0.436 | 0.034 | 0.036 | 0.002 | 0.047 | 0.207 |
| | 3 | 25/100M | 75bp | 0.193 | 0.468 | 0.083 | 0.412 | 0.441 | 0.032 | 0.035 | 0.006 | 0.046 | 0.206 |
| | 3 | 25/100M | 100bp | 0.198 | 0.467 | 0.094 | 0.415 | 0.442 | 0.035 | 0.035 | 0.004 | 0.047 | 0.207 |
| | 3 | 25/100M | 125bp | 0.215 | 0.478 | 0.100 | 0.421 | 0.444 | 0.036 | 0.039 | 0.006 | 0.050 | 0.212 |
| | 3 | 25/100M | 150bp | 0.211 | 0.467 | 0.078 | 0.414 | 0.442 | 0.032 | 0.037 | 0.006 | 0.050 | 0.211 |
| | 5 | 50M | 50bp | 0.661 | 0.757 | 0.631 | 0.692 | 0.489 | 0.029 | 0.042 | 0.025 | 0.041 | 0.042 |
| | 5 | 50M | 75bp | 0.670 | 0.764 | 0.635 | 0.699 | 0.495 | 0.031 | 0.042 | 0.026 | 0.041 | 0.042 |
| | 5 | 50M | 100bp | 0.674 | 0.766 | 0.640 | 0.702 | 0.493 | 0.028 | 0.042 | 0.025 | 0.040 | 0.041 |
| | 5 | 50M | 125bp | 0.681 | 0.769 | 0.646 | 0.707 | 0.492 | 0.030 | 0.040 | 0.026 | 0.040 | 0.040 |
| | 5 | 50M | 150bp | 0.675 | 0.764 | 0.640 | 0.702 | 0.511 | 0.029 | 0.040 | 0.025 | 0.040 | 0.045 |
| | 5 | 25/100M | 50bp | 0.617 | 0.716 | 0.571 | 0.636 | 0.455 | 0.026 | 0.038 | 0.021 | 0.037 | 0.042 |
| | 5 | 25/100M | 75bp | 0.629 | 0.726 | 0.583 | 0.649 | 0.473 | 0.026 | 0.039 | 0.023 | 0.037 | 0.046 |
| | 5 | 25/100M | 100bp | 0.633 | 0.727 | 0.584 | 0.650 | 0.463 | 0.027 | 0.039 | 0.023 | 0.037 | 0.043 |
| | 5 | 25/100M | 125bp | 0.639 | 0.730 | 0.586 | 0.653 | 0.479 | 0.027 | 0.039 | 0.023 | 0.037 | 0.046 |
| | 5 | 25/100M | 150bp | 0.639 | 0.728 | 0.588 | 0.650 | 0.476 | 0.028 | 0.039 | 0.022 | 0.037 | 0.044 |
| | 10 | 50M | 50bp | 0.864 | 0.870 | 0.843 | 0.848 | 0.861 | 0.035 | 0.042 | 0.030 | 0.036 | 0.047 |
| | 10 | 50M | 75bp | 0.865 | 0.876 | 0.847 | 0.852 | 0.862 | 0.036 | 0.043 | 0.031 | 0.037 | 0.046 |
| | 10 | 50M | 100bp | 0.867 | 0.877 | 0.850 | 0.855 | 0.861 | 0.036 | 0.041 | 0.030 | 0.036 | 0.046 |
| | 10 | 50M | 125bp | 0.873 | 0.880 | 0.855 | 0.860 | 0.867 | 0.034 | 0.040 | 0.029 | 0.035 | 0.043 |
| | 10 | 50M | 150bp | 0.875 | 0.883 | 0.858 | 0.863 | 0.869 | 0.036 | 0.041 | 0.030 | 0.035 | 0.046 |
| | 10 | 25/100M | 50bp | 0.845 | 0.878 | 0.803 | 0.808 | 0.848 | 0.034 | 0.040 | 0.028 | 0.033 | 0.047 |
| | 10 | 25/100M | 75bp | 0.848 | 0.880 | 0.808 | 0.813 | 0.850 | 0.034 | 0.041 | 0.029 | 0.034 | 0.047 |
| | 10 | 25/100M | 100bp | 0.851 | 0.884 | 0.812 | 0.817 | 0.852 | 0.035 | 0.040 | 0.029 | 0.034 | 0.049 |
| | 10 | 25/100M | 125bp | 0.854 | 0.885 | 0.817 | 0.822 | 0.854 | 0.033 | 0.040 | 0.028 | 0.033 | 0.046 |
| | 10 | 25/100M | 150bp | 0.855 | 0.884 | 0.818 | 0.823 | 0.855 | 0.034 | 0.039 | 0.027 | 0.033 | 0.046 |

Table S2: Simulation results - observed power and false discovery rate for different read types and read Lengths, averaged over 20 simulations. Scenario with mm39 genome, maximum of 2 transcripts/gene expressed, and reads quantified with kallisto. Library size shown in million reads (M) with 25/100 indicating library sizes alternating between 25M and 100M across replicates. Read lengths are shown in base pairs (bp). Red color indicates observed FDR values greater than the nominal 0.05. Blue color indicates the most powerful method that controls the FDR at the nominal level for a given scenario (row). Empty cells indicate cases in which a method failed to call any transcript as DE. *(continued)*

| | | | | Power | | | | | False Discovery Rate | | | | |
|---|---|---|---|---|---|---|---|---|---|---|---|---|---|
| Read | Samples/Group | Library Size | Read Length | edgeR-raw | edgeR-scaled | sleuth-LRT | sleuth-Wald | Swish | edgeR-raw | edgeR-scaled | sleuth-LRT | sleuth-Wald | Swish |
| single-end | 3 | 50M | 50bp | 0.080 | 0.492 | 0.086 | 0.429 | 0.467 | 0.123 | 0.055 | 0.015 | 0.064 | 0.218 |
| | 3 | 50M | 75bp | 0.116 | 0.502 | 0.095 | 0.437 | 0.473 | 0.105 | 0.052 | 0.016 | 0.062 | 0.223 |
| | 3 | 50M | 100bp | 0.114 | 0.502 | 0.080 | 0.441 | 0.477 | 0.100 | 0.052 | 0.014 | 0.062 | 0.221 |
| | 3 | 50M | 125bp | 0.174 | 0.512 | 0.144 | 0.452 | 0.487 | 0.074 | 0.048 | 0.012 | 0.059 | 0.222 |
| | 3 | 50M | 150bp | 0.200 | 0.510 | 0.159 | 0.458 | 0.490 | 0.071 | 0.047 | 0.015 | 0.060 | 0.227 |
| | 3 | 25/100M | 50bp | 0.001 | 0.445 | 0.004 | 0.378 | 0.412 | 0.390 | 0.045 | 0.004 | 0.053 | 0.203 |
| | 3 | 25/100M | 75bp | 0.030 | 0.444 | 0.012 | 0.376 | 0.412 | 0.103 | 0.043 | 0.003 | 0.051 | 0.205 |
| | 3 | 25/100M | 100bp | 0.065 | 0.453 | 0.015 | 0.388 | 0.421 | 0.071 | 0.044 | 0.010 | 0.050 | 0.208 |
| | 3 | 25/100M | 125bp | 0.105 | 0.454 | 0.010 | 0.396 | 0.428 | 0.060 | 0.041 | 0.011 | 0.053 | 0.205 |
| | 3 | 25/100M | 150bp | 0.131 | 0.460 | 0.023 | 0.401 | 0.433 | 0.052 | 0.041 | 0.007 | 0.051 | 0.211 |
| | 5 | 50M | 50bp | 0.601 | 0.735 | 0.591 | 0.658 | 0.489 | 0.043 | 0.061 | 0.038 | 0.053 | 0.057 |
| | 5 | 50M | 75bp | 0.610 | 0.744 | 0.596 | 0.663 | 0.495 | 0.041 | 0.060 | 0.037 | 0.054 | 0.055 |
| | 5 | 50M | 100bp | 0.620 | 0.748 | 0.603 | 0.672 | 0.499 | 0.041 | 0.060 | 0.036 | 0.054 | 0.055 |
| | 5 | 50M | 125bp | 0.642 | 0.757 | 0.621 | 0.686 | 0.496 | 0.039 | 0.054 | 0.034 | 0.050 | 0.051 |
| | 5 | 50M | 150bp | 0.648 | 0.759 | 0.626 | 0.691 | 0.507 | 0.038 | 0.054 | 0.034 | 0.049 | 0.050 |
| | 5 | 25/100M | 50bp | 0.553 | 0.696 | 0.526 | 0.596 | 0.443 | 0.036 | 0.050 | 0.030 | 0.045 | 0.052 |
| | 5 | 25/100M | 75bp | 0.568 | 0.699 | 0.536 | 0.603 | 0.468 | 0.034 | 0.048 | 0.027 | 0.043 | 0.055 |
| | 5 | 25/100M | 100bp | 0.582 | 0.710 | 0.545 | 0.615 | 0.458 | 0.034 | 0.051 | 0.029 | 0.046 | 0.051 |
| | 5 | 25/100M | 125bp | 0.599 | 0.718 | 0.560 | 0.630 | 0.458 | 0.034 | 0.049 | 0.028 | 0.044 | 0.050 |
| | 5 | 25/100M | 150bp | 0.609 | 0.720 | 0.566 | 0.634 | 0.459 | 0.035 | 0.048 | 0.029 | 0.044 | 0.051 |
| | 10 | 50M | 50bp | 0.834 | 0.847 | 0.807 | 0.812 | 0.838 | 0.071 | 0.072 | 0.060 | 0.067 | 0.088 |
| | 10 | 50M | 75bp | 0.836 | 0.851 | 0.813 | 0.817 | 0.839 | 0.068 | 0.070 | 0.057 | 0.064 | 0.081 |
| | 10 | 50M | 100bp | 0.843 | 0.858 | 0.820 | 0.825 | 0.845 | 0.067 | 0.069 | 0.057 | 0.064 | 0.081 |
| | 10 | 50M | 125bp | 0.852 | 0.864 | 0.833 | 0.838 | 0.851 | 0.063 | 0.063 | 0.052 | 0.057 | 0.075 |
| | 10 | 50M | 150bp | 0.858 | 0.869 | 0.839 | 0.844 | 0.856 | 0.060 | 0.062 | 0.050 | 0.056 | 0.072 |
| | 10 | 25/100M | 50bp | 0.807 | 0.853 | 0.757 | 0.762 | 0.822 | 0.064 | 0.072 | 0.048 | 0.054 | 0.082 |
| | 10 | 25/100M | 75bp | 0.815 | 0.858 | 0.766 | 0.771 | 0.828 | 0.062 | 0.070 | 0.048 | 0.054 | 0.080 |
| | 10 | 25/100M | 100bp | 0.822 | 0.865 | 0.776 | 0.781 | 0.834 | 0.063 | 0.070 | 0.050 | 0.055 | 0.083 |
| | 10 | 25/100M | 125bp | 0.831 | 0.869 | 0.787 | 0.792 | 0.838 | 0.056 | 0.063 | 0.044 | 0.050 | 0.073 |
| | 10 | 25/100M | 150bp | 0.841 | 0.877 | 0.799 | 0.803 | 0.846 | 0.056 | 0.062 | 0.043 | 0.048 | 0.073 |

Table S3: Simulation results - observed power and false discovery rate for different read types and read Lengths, averaged over 20 simulations. Scenario with mm39 genome, maximum of 3 transcripts/gene expressed, and reads quantified with Salmon. Library size shown in million reads (M) with 25/100 indicating library sizes alternating between 25M and 100M across replicates. Read lengths are shown in base pairs (bp). Red color indicates observed FDR values greater than the nominal 0.05. Blue color indicates the most powerful method that controls the FDR at the nominal level for a given scenario (row). Empty cells indicate cases in which a method failed to call any transcript as DE.

| | | | | Power | | | | | False Discovery Rate | | | | |
|---|---|---|---|---|---|---|---|---|---|---|---|---|---|
| Read | Samples/Group | Library Size | Read Length | edgeR-raw | edgeR-scaled | sleuth-LRT | sleuth-Wald | Swish | edgeR-raw | edgeR-scaled | sleuth-LRT | sleuth-Wald | Swish |
| paired-end | 3 | 50M | 50bp | 0.040 | 0.428 | 0.000 | 0.390 | 0.426 | 0.156 | 0.040 | - | 0.055 | 0.243 |
| | 3 | 50M | 75bp | 0.120 | 0.435 | 0.000 | 0.395 | 0.428 | 0.056 | 0.040 | - | 0.056 | 0.241 |
| | 3 | 50M | 100bp | 0.124 | 0.431 | 0.000 | 0.392 | 0.420 | 0.047 | 0.041 | - | 0.053 | 0.239 |
| | 3 | 50M | 125bp | 0.139 | 0.433 | 0.000 | 0.400 | 0.434 | 0.046 | 0.039 | - | 0.054 | 0.249 |
| | 3 | 50M | 150bp | 0.154 | 0.442 | 0.008 | 0.409 | 0.437 | 0.049 | 0.040 | 0.006 | 0.057 | 0.247 |
| | 3 | 25/100M | 50bp | 0.000 | 0.379 | 0.000 | 0.334 | 0.373 | 0.694 | 0.037 | - | 0.051 | 0.232 |
| | 3 | 25/100M | 75bp | 0.018 | 0.382 | 0.000 | 0.338 | 0.373 | 0.123 | 0.036 | - | 0.050 | 0.231 |
| | 3 | 25/100M | 100bp | 0.011 | 0.382 | 0.000 | 0.331 | 0.362 | 0.162 | 0.037 | - | 0.051 | 0.228 |
| | 3 | 25/100M | 125bp | 0.018 | 0.384 | 0.000 | 0.346 | 0.381 | 0.113 | 0.036 | - | 0.048 | 0.235 |
| | 3 | 25/100M | 150bp | 0.028 | 0.385 | 0.000 | 0.342 | 0.380 | 0.083 | 0.037 | - | 0.049 | 0.235 |
| | 5 | 50M | 50bp | 0.555 | 0.690 | 0.535 | 0.616 | 0.415 | 0.027 | 0.042 | 0.024 | 0.041 | 0.041 |
| | 5 | 50M | 75bp | 0.584 | 0.697 | 0.548 | 0.629 | 0.409 | 0.025 | 0.040 | 0.022 | 0.039 | 0.042 |
| | 5 | 50M | 100bp | 0.582 | 0.693 | 0.545 | 0.623 | 0.415 | 0.024 | 0.041 | 0.022 | 0.040 | 0.043 |
| | 5 | 50M | 125bp | 0.595 | 0.699 | 0.555 | 0.635 | 0.420 | 0.024 | 0.041 | 0.022 | 0.039 | 0.045 |
| | 5 | 50M | 150bp | 0.592 | 0.701 | 0.553 | 0.634 | 0.437 | 0.025 | 0.043 | 0.023 | 0.041 | 0.048 |
| | 5 | 25/100M | 50bp | 0.497 | 0.646 | 0.474 | 0.554 | 0.382 | 0.021 | 0.040 | 0.018 | 0.036 | 0.042 |
| | 5 | 25/100M | 75bp | 0.526 | 0.650 | 0.483 | 0.563 | 0.397 | 0.020 | 0.038 | 0.017 | 0.035 | 0.047 |
| | 5 | 25/100M | 100bp | 0.525 | 0.649 | 0.482 | 0.562 | 0.396 | 0.020 | 0.037 | 0.017 | 0.035 | 0.043 |
| | 5 | 25/100M | 125bp | 0.535 | 0.657 | 0.493 | 0.576 | 0.381 | 0.021 | 0.038 | 0.019 | 0.037 | 0.042 |
| | 5 | 25/100M | 150bp | 0.532 | 0.656 | 0.489 | 0.573 | 0.394 | 0.020 | 0.038 | 0.018 | 0.035 | 0.046 |
| | 10 | 50M | 50bp | 0.807 | 0.822 | 0.780 | 0.787 | 0.810 | 0.037 | 0.045 | 0.031 | 0.039 | 0.051 |
| | 10 | 50M | 75bp | 0.818 | 0.826 | 0.791 | 0.798 | 0.813 | 0.035 | 0.043 | 0.028 | 0.035 | 0.050 |
| | 10 | 50M | 100bp | 0.815 | 0.820 | 0.788 | 0.796 | 0.809 | 0.032 | 0.042 | 0.028 | 0.035 | 0.045 |
| | 10 | 50M | 125bp | 0.822 | 0.830 | 0.797 | 0.805 | 0.816 | 0.033 | 0.042 | 0.029 | 0.036 | 0.046 |
| | 10 | 50M | 150bp | 0.826 | 0.832 | 0.799 | 0.807 | 0.820 | 0.033 | 0.042 | 0.029 | 0.036 | 0.047 |
| | 10 | 25/100M | 50bp | 0.782 | 0.826 | 0.729 | 0.736 | 0.794 | 0.032 | 0.042 | 0.025 | 0.032 | 0.048 |
| | 10 | 25/100M | 75bp | 0.795 | 0.832 | 0.739 | 0.747 | 0.798 | 0.030 | 0.040 | 0.025 | 0.031 | 0.047 |
| | 10 | 25/100M | 100bp | 0.793 | 0.828 | 0.738 | 0.746 | 0.796 | 0.030 | 0.040 | 0.024 | 0.031 | 0.047 |
| | 10 | 25/100M | 125bp | 0.802 | 0.836 | 0.751 | 0.760 | 0.801 | 0.030 | 0.041 | 0.025 | 0.033 | 0.046 |
| | 10 | 25/100M | 150bp | 0.802 | 0.838 | 0.751 | 0.759 | 0.801 | 0.029 | 0.039 | 0.024 | 0.032 | 0.044 |

Table S3: Simulation results - observed power and false discovery rate for different read types and read Lengths, averaged over 20 simulations. Scenario with mm39 genome, maximum of 3 transcripts/gene expressed, and reads quantified with Salmon. Library size shown in million reads (M) with 25/100 indicating library sizes alternating between 25M and 100M across replicates. Read lengths are shown in base pairs (bp). Red color indicates observed FDR values greater than the nominal 0.05. Blue color indicates the most powerful method that controls the FDR at the nominal level for a given scenario (row). Empty cells indicate cases in which a method failed to call any transcript as DE. *(continued)*

| | | | | Power | | | | | False Discovery Rate | | | | |
|---|---|---|---|---|---|---|---|---|---|---|---|---|---|
| Read | Samples/Group | Library Size | Read Length | edgeR-raw | edgeR-scaled | sleuth-LRT | sleuth-Wald | Swish | edgeR-raw | edgeR-scaled | sleuth-LRT | sleuth-Wald | Swish |
| single-end | 3 | 50M | 50bp | 0.005 | 0.405 | 0.000 | 0.366 | 0.405 | 0.431 | 0.037 | - | 0.053 | 0.238 |
| | 3 | 50M | 75bp | 0.008 | 0.413 | 0.003 | 0.367 | 0.401 | 0.291 | 0.037 | 0.007 | 0.054 | 0.242 |
| | 3 | 50M | 100bp | 0.014 | 0.416 | 0.000 | 0.371 | 0.406 | 0.227 | 0.039 | - | 0.055 | 0.237 |
| | 3 | 50M | 125bp | 0.053 | 0.430 | 0.000 | 0.390 | 0.421 | 0.106 | 0.040 | 0.000 | 0.056 | 0.245 |
| | 3 | 50M | 150bp | 0.077 | 0.424 | 0.000 | 0.391 | 0.427 | 0.081 | 0.037 | - | 0.054 | 0.242 |
| | 3 | 25/100M | 50bp | 0.000 | 0.355 | 0.000 | 0.302 | 0.346 | 0.680 | 0.037 | - | 0.046 | 0.229 |
| | 3 | 25/100M | 75bp | 0.000 | 0.360 | 0.000 | 0.309 | 0.349 | 0.705 | 0.033 | - | 0.045 | 0.231 |
| | 3 | 25/100M | 100bp | 0.000 | 0.366 | 0.000 | 0.309 | 0.349 | 0.684 | 0.037 | - | 0.048 | 0.225 |
| | 3 | 25/100M | 125bp | 0.001 | 0.379 | 0.000 | 0.331 | 0.367 | 0.571 | 0.037 | - | 0.051 | 0.232 |
| | 3 | 25/100M | 150bp | 0.006 | 0.377 | 0.000 | 0.333 | 0.370 | 0.249 | 0.034 | - | 0.050 | 0.236 |
| | 5 | 50M | 50bp | 0.509 | 0.660 | 0.501 | 0.576 | 0.399 | 0.023 | 0.041 | 0.022 | 0.039 | 0.042 |
| | 5 | 50M | 75bp | 0.521 | 0.666 | 0.508 | 0.585 | 0.411 | 0.022 | 0.039 | 0.020 | 0.037 | 0.046 |
| | 5 | 50M | 100bp | 0.534 | 0.673 | 0.516 | 0.595 | 0.422 | 0.022 | 0.042 | 0.022 | 0.040 | 0.046 |
| | 5 | 50M | 125bp | 0.561 | 0.685 | 0.538 | 0.615 | 0.406 | 0.025 | 0.041 | 0.023 | 0.041 | 0.043 |
| | 5 | 50M | 150bp | 0.571 | 0.690 | 0.543 | 0.622 | 0.418 | 0.025 | 0.041 | 0.023 | 0.040 | 0.047 |
| | 5 | 25/100M | 50bp | 0.444 | 0.612 | 0.429 | 0.504 | 0.374 | 0.018 | 0.038 | 0.016 | 0.033 | 0.042 |
| | 5 | 25/100M | 75bp | 0.460 | 0.617 | 0.440 | 0.515 | 0.379 | 0.019 | 0.037 | 0.017 | 0.034 | 0.045 |
| | 5 | 25/100M | 100bp | 0.471 | 0.623 | 0.445 | 0.523 | 0.360 | 0.019 | 0.037 | 0.016 | 0.033 | 0.041 |
| | 5 | 25/100M | 125bp | 0.492 | 0.638 | 0.464 | 0.544 | 0.378 | 0.021 | 0.038 | 0.018 | 0.035 | 0.045 |
| | 5 | 25/100M | 150bp | 0.512 | 0.644 | 0.477 | 0.557 | 0.399 | 0.020 | 0.038 | 0.018 | 0.036 | 0.045 |
| | 10 | 50M | 50bp | 0.765 | 0.782 | 0.726 | 0.733 | 0.782 | 0.032 | 0.042 | 0.027 | 0.034 | 0.048 |
| | 10 | 50M | 75bp | 0.777 | 0.790 | 0.741 | 0.749 | 0.787 | 0.031 | 0.042 | 0.027 | 0.034 | 0.046 |
| | 10 | 50M | 100bp | 0.784 | 0.796 | 0.751 | 0.758 | 0.790 | 0.034 | 0.043 | 0.027 | 0.035 | 0.046 |
| | 10 | 50M | 125bp | 0.800 | 0.812 | 0.770 | 0.778 | 0.800 | 0.032 | 0.042 | 0.027 | 0.034 | 0.046 |
| | 10 | 50M | 150bp | 0.809 | 0.820 | 0.784 | 0.792 | 0.809 | 0.034 | 0.042 | 0.028 | 0.035 | 0.046 |
| | 10 | 25/100M | 50bp | 0.736 | 0.791 | 0.669 | 0.675 | 0.766 | 0.029 | 0.038 | 0.023 | 0.030 | 0.046 |
| | 10 | 25/100M | 75bp | 0.744 | 0.797 | 0.678 | 0.684 | 0.767 | 0.029 | 0.039 | 0.022 | 0.028 | 0.045 |
| | 10 | 25/100M | 100bp | 0.753 | 0.802 | 0.689 | 0.695 | 0.773 | 0.029 | 0.040 | 0.023 | 0.030 | 0.048 |
| | 10 | 25/100M | 125bp | 0.774 | 0.817 | 0.716 | 0.724 | 0.788 | 0.030 | 0.039 | 0.024 | 0.030 | 0.047 |
| | 10 | 25/100M | 150bp | 0.785 | 0.824 | 0.728 | 0.736 | 0.790 | 0.029 | 0.040 | 0.023 | 0.031 | 0.046 |

Table S4: Simulation results - observed power and false discovery rate for different read types and read Lengths, averaged over 20 simulations. Scenario with mm39 genome, maximum of 3 transcripts/gene expressed, and reads quantified with kallisto. Library size shown in million reads (M) with 25/100 indicating library sizes alternating between 25M and 100M across replicates. Read lengths are shown in base pairs (bp). Red color indicates observed FDR values greater than the nominal 0.05. Blue color indicates the most powerful method that controls the FDR at the nominal level for a given scenario (row). Empty cells indicate cases in which a method failed to call any transcript as DE.

| | | | | Power | | | | | False Discovery Rate | | | | |
|---|---|---|---|---|---|---|---|---|---|---|---|---|---|
| Read | Samples/Group | Library Size | Read Length | edgeR-raw | edgeR-scaled | sleuth-LRT | sleuth-Wald | Swish | edgeR-raw | edgeR-scaled | sleuth-LRT | sleuth-Wald | Swish |
| paired-end | 3 | 50M | 50bp | 0.075 | 0.428 | 0.000 | 0.391 | 0.435 | 0.093 | 0.040 | - | 0.054 | 0.256 |
| | 3 | 50M | 75bp | 0.111 | 0.435 | 0.000 | 0.396 | 0.443 | 0.080 | 0.041 | - | 0.055 | 0.255 |
| | 3 | 50M | 100bp | 0.121 | 0.435 | 0.000 | 0.399 | 0.442 | 0.076 | 0.040 | - | 0.054 | 0.260 |
| | 3 | 50M | 125bp | 0.127 | 0.437 | 0.000 | 0.399 | 0.445 | 0.079 | 0.040 | - | 0.054 | 0.259 |
| | 3 | 50M | 150bp | 0.142 | 0.445 | 0.000 | 0.408 | 0.450 | 0.080 | 0.041 | - | 0.058 | 0.258 |
| | 3 | 25/100M | 50bp | 0.005 | 0.375 | 0.000 | 0.334 | 0.381 | 0.182 | 0.037 | - | 0.050 | 0.242 |
| | 3 | 25/100M | 75bp | 0.014 | 0.379 | 0.000 | 0.341 | 0.384 | 0.086 | 0.037 | - | 0.050 | 0.242 |
| | 3 | 25/100M | 100bp | 0.020 | 0.385 | 0.000 | 0.340 | 0.383 | 0.102 | 0.039 | - | 0.051 | 0.243 |
| | 3 | 25/100M | 125bp | 0.032 | 0.384 | 0.000 | 0.346 | 0.388 | 0.103 | 0.036 | - | 0.049 | 0.242 |
| | 3 | 25/100M | 150bp | 0.038 | 0.385 | 0.000 | 0.344 | 0.386 | 0.084 | 0.037 | - | 0.050 | 0.243 |
| | 5 | 50M | 50bp | 0.563 | 0.689 | 0.537 | 0.619 | 0.431 | 0.028 | 0.042 | 0.022 | 0.041 | 0.046 |
| | 5 | 50M | 75bp | 0.579 | 0.701 | 0.549 | 0.633 | 0.423 | 0.028 | 0.041 | 0.023 | 0.040 | 0.045 |
| | 5 | 50M | 100bp | 0.581 | 0.703 | 0.553 | 0.635 | 0.414 | 0.028 | 0.042 | 0.023 | 0.042 | 0.043 |
| | 5 | 50M | 125bp | 0.585 | 0.702 | 0.554 | 0.636 | 0.417 | 0.027 | 0.042 | 0.023 | 0.041 | 0.043 |
| | 5 | 50M | 150bp | 0.586 | 0.705 | 0.555 | 0.638 | 0.429 | 0.028 | 0.044 | 0.023 | 0.042 | 0.046 |
| | 5 | 25/100M | 50bp | 0.510 | 0.645 | 0.477 | 0.558 | 0.372 | 0.023 | 0.039 | 0.020 | 0.037 | 0.044 |
| | 5 | 25/100M | 75bp | 0.529 | 0.653 | 0.488 | 0.570 | 0.386 | 0.023 | 0.039 | 0.018 | 0.036 | 0.045 |
| | 5 | 25/100M | 100bp | 0.537 | 0.660 | 0.494 | 0.577 | 0.394 | 0.023 | 0.039 | 0.020 | 0.038 | 0.044 |
| | 5 | 25/100M | 125bp | 0.536 | 0.660 | 0.493 | 0.578 | 0.382 | 0.025 | 0.039 | 0.020 | 0.038 | 0.043 |
| | 5 | 25/100M | 150bp | 0.535 | 0.658 | 0.491 | 0.578 | 0.395 | 0.024 | 0.039 | 0.019 | 0.037 | 0.046 |
| | 10 | 50M | 50bp | 0.810 | 0.824 | 0.787 | 0.794 | 0.808 | 0.037 | 0.043 | 0.029 | 0.037 | 0.046 |
| | 10 | 50M | 75bp | 0.817 | 0.832 | 0.797 | 0.803 | 0.814 | 0.036 | 0.045 | 0.028 | 0.037 | 0.050 |
| | 10 | 50M | 100bp | 0.822 | 0.835 | 0.800 | 0.808 | 0.816 | 0.034 | 0.044 | 0.029 | 0.037 | 0.047 |
| | 10 | 50M | 125bp | 0.824 | 0.837 | 0.803 | 0.810 | 0.817 | 0.034 | 0.043 | 0.030 | 0.037 | 0.046 |
| | 10 | 50M | 150bp | 0.826 | 0.840 | 0.804 | 0.812 | 0.822 | 0.033 | 0.043 | 0.029 | 0.037 | 0.048 |
| | 10 | 25/100M | 50bp | 0.788 | 0.830 | 0.737 | 0.745 | 0.794 | 0.033 | 0.041 | 0.026 | 0.033 | 0.048 |
| | 10 | 25/100M | 75bp | 0.796 | 0.840 | 0.750 | 0.757 | 0.801 | 0.032 | 0.041 | 0.026 | 0.033 | 0.048 |
| | 10 | 25/100M | 100bp | 0.800 | 0.844 | 0.755 | 0.762 | 0.803 | 0.032 | 0.042 | 0.026 | 0.033 | 0.048 |
| | 10 | 25/100M | 125bp | 0.803 | 0.844 | 0.759 | 0.767 | 0.806 | 0.032 | 0.042 | 0.027 | 0.034 | 0.048 |
| | 10 | 25/100M | 150bp | 0.803 | 0.846 | 0.759 | 0.767 | 0.805 | 0.031 | 0.040 | 0.025 | 0.033 | 0.045 |

Table S4: Simulation results - observed power and false discovery rate for different read types and read Lengths, averaged over 20 simulations. Scenario with mm39 genome, maximum of 3 transcripts/gene expressed, and reads quantified with kallisto. Library size shown in million reads (M) with 25/100 indicating library sizes alternating between 25M and 100M across replicates. Read lengths are shown in base pairs (bp). Red color indicates observed FDR values greater than the nominal 0.05. Blue color indicates the most powerful method that controls the FDR at the nominal level for a given scenario (row). Empty cells indicate cases in which a method failed to call any transcript as DE. *(continued)*

| | | | | Power | | | | | False Discovery Rate | | | | |
|---|---|---|---|---|---|---|---|---|---|---|---|---|---|
| Read | Samples/Group | Library Size | Read Length | edgeR-raw | edgeR-scaled | sleuth-LRT | sleuth-Wald | Swish | edgeR-raw | edgeR-scaled | sleuth-LRT | sleuth-Wald | Swish |
| single-end | 3 | 50M | 50bp | 0.001 | 0.409 | 0.000 | 0.365 | 0.415 | 0.648 | 0.049 | - | 0.062 | 0.251 |
| | 3 | 50M | 75bp | 0.003 | 0.409 | 0.000 | 0.366 | 0.413 | 0.411 | 0.047 | - | 0.061 | 0.255 |
| | 3 | 50M | 100bp | 0.002 | 0.413 | 0.000 | 0.369 | 0.420 | 0.366 | 0.048 | - | 0.061 | 0.253 |
| | 3 | 50M | 125bp | 0.015 | 0.428 | 0.000 | 0.386 | 0.431 | 0.177 | 0.049 | - | 0.060 | 0.255 |
| | 3 | 50M | 150bp | 0.034 | 0.423 | 0.000 | 0.386 | 0.435 | 0.161 | 0.044 | - | 0.059 | 0.252 |
| | 3 | 25/100M | 50bp | 0.000 | 0.352 | 0.000 | 0.303 | 0.350 | 0.818 | 0.043 | - | 0.052 | 0.239 |
| | 3 | 25/100M | 75bp | 0.000 | 0.359 | 0.000 | 0.309 | 0.355 | 0.674 | 0.040 | - | 0.050 | 0.241 |
| | 3 | 25/100M | 100bp | 0.000 | 0.363 | 0.000 | 0.310 | 0.359 | 0.621 | 0.041 | - | 0.051 | 0.238 |
| | 3 | 25/100M | 125bp | 0.000 | 0.375 | 0.000 | 0.329 | 0.372 | 0.660 | 0.041 | - | 0.054 | 0.240 |
| | 3 | 25/100M | 150bp | 0.001 | 0.374 | 0.000 | 0.331 | 0.376 | 0.343 | 0.040 | 1.000 | 0.052 | 0.243 |
| | 5 | 50M | 50bp | 0.501 | 0.665 | 0.499 | 0.583 | 0.400 | 0.036 | 0.057 | 0.032 | 0.051 | 0.048 |
| | 5 | 50M | 75bp | 0.513 | 0.671 | 0.506 | 0.591 | 0.406 | 0.036 | 0.054 | 0.031 | 0.051 | 0.051 |
| | 5 | 50M | 100bp | 0.527 | 0.679 | 0.516 | 0.602 | 0.407 | 0.036 | 0.056 | 0.032 | 0.052 | 0.053 |
| | 5 | 50M | 125bp | 0.552 | 0.690 | 0.533 | 0.616 | 0.410 | 0.036 | 0.053 | 0.031 | 0.050 | 0.051 |
| | 5 | 50M | 150bp | 0.558 | 0.692 | 0.537 | 0.622 | 0.426 | 0.035 | 0.052 | 0.030 | 0.049 | 0.054 |
| | 5 | 25/100M | 50bp | 0.447 | 0.617 | 0.432 | 0.516 | 0.374 | 0.029 | 0.048 | 0.025 | 0.044 | 0.051 |
| | 5 | 25/100M | 75bp | 0.464 | 0.622 | 0.443 | 0.525 | 0.369 | 0.031 | 0.047 | 0.025 | 0.044 | 0.050 |
| | 5 | 25/100M | 100bp | 0.477 | 0.630 | 0.450 | 0.536 | 0.371 | 0.026 | 0.045 | 0.023 | 0.040 | 0.048 |
| | 5 | 25/100M | 125bp | 0.495 | 0.642 | 0.466 | 0.549 | 0.390 | 0.029 | 0.047 | 0.023 | 0.043 | 0.051 |
| | 5 | 25/100M | 150bp | 0.511 | 0.648 | 0.475 | 0.558 | 0.391 | 0.028 | 0.046 | 0.024 | 0.041 | 0.050 |
| | 10 | 50M | 50bp | 0.773 | 0.794 | 0.742 | 0.749 | 0.781 | 0.060 | 0.066 | 0.051 | 0.059 | 0.078 |
| | 10 | 50M | 75bp | 0.783 | 0.804 | 0.754 | 0.762 | 0.788 | 0.057 | 0.063 | 0.047 | 0.055 | 0.073 |
| | 10 | 50M | 100bp | 0.790 | 0.809 | 0.763 | 0.770 | 0.791 | 0.058 | 0.063 | 0.048 | 0.056 | 0.073 |
| | 10 | 50M | 125bp | 0.800 | 0.820 | 0.777 | 0.785 | 0.799 | 0.054 | 0.059 | 0.046 | 0.054 | 0.070 |
| | 10 | 50M | 150bp | 0.809 | 0.827 | 0.788 | 0.796 | 0.808 | 0.054 | 0.059 | 0.044 | 0.052 | 0.069 |
| | 10 | 25/100M | 50bp | 0.746 | 0.804 | 0.691 | 0.697 | 0.767 | 0.053 | 0.063 | 0.043 | 0.050 | 0.074 |
| | 10 | 25/100M | 75bp | 0.752 | 0.808 | 0.697 | 0.705 | 0.770 | 0.053 | 0.064 | 0.040 | 0.048 | 0.073 |
| | 10 | 25/100M | 100bp | 0.761 | 0.814 | 0.708 | 0.715 | 0.775 | 0.051 | 0.061 | 0.040 | 0.048 | 0.071 |
| | 10 | 25/100M | 125bp | 0.777 | 0.826 | 0.726 | 0.734 | 0.786 | 0.048 | 0.058 | 0.037 | 0.045 | 0.066 |
| | 10 | 25/100M | 150bp | 0.784 | 0.831 | 0.736 | 0.744 | 0.790 | 0.049 | 0.058 | 0.035 | 0.044 | 0.067 |

Table S5: Simulation results - observed power and false discovery rate for different read types and read Lengths, averaged over 20 simulations. Scenario with mm39 genome, maximum of 4 transcripts/gene expressed, and reads quantified with Salmon. Library size shown in million reads (M) with 25/100 indicating library sizes alternating between 25M and 100M across replicates. Read lengths are shown in base pairs (bp). Red color indicates observed FDR values greater than the nominal 0.05. Blue color indicates the most powerful method that controls the FDR at the nominal level for a given scenario (row). Empty cells indicate cases in which a method failed to call any transcript as DE.

| | | | | Power | | | | | False Discovery Rate | | | | |
|---|---|---|---|---|---|---|---|---|---|---|---|---|---|
| Read | Samples/Group | Library Size | Read Length | edgeR-raw | edgeR-scaled | sleuth-LRT | sleuth-Wald | Swish | edgeR-raw | edgeR-scaled | sleuth-LRT | sleuth-Wald | Swish |
| paired-end | 3 | 50M | 50bp | 0.007 | 0.433 | 0.000 | 0.397 | 0.436 | 0.365 | 0.038 | - | 0.056 | 0.246 |
| | 3 | 50M | 75bp | 0.047 | 0.437 | 0.000 | 0.403 | 0.438 | 0.134 | 0.039 | - | 0.057 | 0.247 |
| | 3 | 50M | 100bp | 0.040 | 0.435 | 0.000 | 0.399 | 0.431 | 0.138 | 0.038 | - | 0.054 | 0.237 |
| | 3 | 50M | 125bp | 0.065 | 0.443 | 0.000 | 0.409 | 0.444 | 0.095 | 0.039 | 0.000 | 0.056 | 0.254 |
| | 3 | 50M | 150bp | 0.064 | 0.444 | 0.000 | 0.412 | 0.444 | 0.099 | 0.040 | - | 0.055 | 0.248 |
| | 3 | 25/100M | 50bp | 0.000 | 0.372 | 0.000 | 0.329 | 0.374 | 0.810 | 0.036 | - | 0.047 | 0.235 |
| | 3 | 25/100M | 75bp | 0.002 | 0.390 | 0.000 | 0.342 | 0.380 | 0.513 | 0.034 | - | 0.047 | 0.225 |
| | 3 | 25/100M | 100bp | 0.001 | 0.387 | 0.000 | 0.335 | 0.373 | 0.551 | 0.036 | - | 0.048 | 0.226 |
| | 3 | 25/100M | 125bp | 0.001 | 0.391 | 0.000 | 0.348 | 0.388 | 0.497 | 0.036 | - | 0.048 | 0.233 |
| | 3 | 25/100M | 150bp | 0.003 | 0.390 | 0.000 | 0.348 | 0.388 | 0.363 | 0.035 | - | 0.049 | 0.236 |
| | 5 | 50M | 50bp | 0.561 | 0.718 | 0.550 | 0.641 | 0.424 | 0.024 | 0.041 | 0.022 | 0.042 | 0.044 |
| | 5 | 50M | 75bp | 0.587 | 0.727 | 0.561 | 0.654 | 0.411 | 0.024 | 0.042 | 0.020 | 0.041 | 0.042 |
| | 5 | 50M | 100bp | 0.584 | 0.724 | 0.557 | 0.648 | 0.417 | 0.022 | 0.041 | 0.020 | 0.039 | 0.040 |
| | 5 | 50M | 125bp | 0.593 | 0.728 | 0.563 | 0.660 | 0.430 | 0.023 | 0.040 | 0.021 | 0.039 | 0.043 |
| | 5 | 50M | 150bp | 0.600 | 0.731 | 0.571 | 0.663 | 0.431 | 0.023 | 0.040 | 0.021 | 0.039 | 0.043 |
| | 5 | 25/100M | 50bp | 0.496 | 0.672 | 0.475 | 0.569 | 0.375 | 0.018 | 0.037 | 0.016 | 0.033 | 0.042 |
| | 5 | 25/100M | 75bp | 0.524 | 0.684 | 0.489 | 0.583 | 0.404 | 0.019 | 0.037 | 0.017 | 0.034 | 0.041 |
| | 5 | 25/100M | 100bp | 0.521 | 0.674 | 0.485 | 0.578 | 0.388 | 0.018 | 0.038 | 0.016 | 0.034 | 0.038 |
| | 5 | 25/100M | 125bp | 0.532 | 0.686 | 0.494 | 0.589 | 0.413 | 0.018 | 0.039 | 0.017 | 0.036 | 0.044 |
| | 5 | 25/100M | 150bp | 0.541 | 0.689 | 0.502 | 0.595 | 0.406 | 0.020 | 0.038 | 0.017 | 0.034 | 0.044 |
| | 10 | 50M | 50bp | 0.854 | 0.871 | 0.826 | 0.834 | 0.858 | 0.034 | 0.044 | 0.028 | 0.036 | 0.048 |
| | 10 | 50M | 75bp | 0.862 | 0.872 | 0.832 | 0.840 | 0.860 | 0.034 | 0.043 | 0.029 | 0.036 | 0.049 |
| | 10 | 50M | 100bp | 0.863 | 0.870 | 0.830 | 0.839 | 0.857 | 0.031 | 0.040 | 0.026 | 0.034 | 0.046 |
| | 10 | 50M | 125bp | 0.872 | 0.882 | 0.846 | 0.854 | 0.867 | 0.033 | 0.042 | 0.028 | 0.035 | 0.047 |
| | 10 | 50M | 150bp | 0.873 | 0.881 | 0.844 | 0.853 | 0.870 | 0.032 | 0.041 | 0.027 | 0.035 | 0.047 |
| | 10 | 25/100M | 50bp | 0.826 | 0.876 | 0.766 | 0.775 | 0.842 | 0.030 | 0.040 | 0.024 | 0.031 | 0.048 |
| | 10 | 25/100M | 75bp | 0.837 | 0.880 | 0.778 | 0.786 | 0.844 | 0.028 | 0.037 | 0.022 | 0.029 | 0.046 |
| | 10 | 25/100M | 100bp | 0.833 | 0.876 | 0.774 | 0.782 | 0.840 | 0.029 | 0.039 | 0.023 | 0.031 | 0.046 |
| | 10 | 25/100M | 125bp | 0.843 | 0.884 | 0.787 | 0.797 | 0.849 | 0.028 | 0.039 | 0.023 | 0.031 | 0.047 |
| | 10 | 25/100M | 150bp | 0.847 | 0.887 | 0.790 | 0.799 | 0.851 | 0.029 | 0.040 | 0.025 | 0.032 | 0.047 |

Table S5: Simulation results - observed power and false discovery rate for different read types and read Lengths, averaged over 20 simulations. Scenario with mm39 genome, maximum of 4 transcripts/gene expressed, and reads quantified with Salmon. Library size shown in million reads (M) with 25/100 indicating library sizes alternating between 25M and 100M across replicates. Read lengths are shown in base pairs (bp). Red color indicates observed FDR values greater than the nominal 0.05. Blue color indicates the most powerful method that controls the FDR at the nominal level for a given scenario (row). Empty cells indicate cases in which a method failed to call any transcript as DE. *(continued)*

| | | | | Power | | | | | False Discovery Rate | | | | |
|---|---|---|---|---|---|---|---|---|---|---|---|---|---|
| Read | Samples/Group | Library Size | Read Length | edgeR-raw | edgeR-scaled | sleuth-LRT | sleuth-Wald | Swish | edgeR-raw | edgeR-scaled | sleuth-LRT | sleuth-Wald | Swish |
| single-end | 3 | 50M | 50bp | 0.002 | 0.399 | 0.000 | 0.363 | 0.407 | 0.651 | 0.035 | 1.000 | 0.052 | 0.240 |
| | 3 | 50M | 75bp | 0.002 | 0.411 | 0.000 | 0.370 | 0.408 | 0.656 | 0.037 | - | 0.053 | 0.242 |
| | 3 | 50M | 100bp | 0.004 | 0.419 | 0.000 | 0.376 | 0.413 | 0.525 | 0.038 | 1.000 | 0.053 | 0.240 |
| | 3 | 50M | 125bp | 0.009 | 0.423 | 0.000 | 0.394 | 0.430 | 0.388 | 0.039 | 0.500 | 0.057 | 0.246 |
| | 3 | 50M | 150bp | 0.017 | 0.434 | 0.000 | 0.400 | 0.438 | 0.259 | 0.040 | - | 0.056 | 0.245 |
| | 3 | 25/100M | 50bp | 0.000 | 0.352 | 0.000 | 0.295 | 0.350 | 0.571 | 0.034 | - | 0.046 | 0.231 |
| | 3 | 25/100M | 75bp | 0.000 | 0.361 | 0.000 | 0.307 | 0.353 | 0.793 | 0.037 | - | 0.048 | 0.228 |
| | 3 | 25/100M | 100bp | 0.000 | 0.373 | 0.000 | 0.314 | 0.358 | 0.710 | 0.036 | - | 0.046 | 0.219 |
| | 3 | 25/100M | 125bp | 0.000 | 0.379 | 0.000 | 0.326 | 0.372 | 0.733 | 0.035 | - | 0.044 | 0.226 |
| | 3 | 25/100M | 150bp | 0.001 | 0.379 | 0.000 | 0.331 | 0.376 | 0.687 | 0.033 | - | 0.047 | 0.231 |
| | 5 | 50M | 50bp | 0.498 | 0.680 | 0.502 | 0.588 | 0.409 | 0.021 | 0.039 | 0.020 | 0.038 | 0.043 |
| | 5 | 50M | 75bp | 0.515 | 0.691 | 0.511 | 0.601 | 0.419 | 0.023 | 0.043 | 0.022 | 0.041 | 0.046 |
| | 5 | 50M | 100bp | 0.522 | 0.698 | 0.518 | 0.609 | 0.402 | 0.022 | 0.041 | 0.020 | 0.040 | 0.039 |
| | 5 | 50M | 125bp | 0.553 | 0.712 | 0.541 | 0.633 | 0.428 | 0.023 | 0.041 | 0.020 | 0.038 | 0.045 |
| | 5 | 50M | 150bp | 0.571 | 0.714 | 0.553 | 0.643 | 0.424 | 0.023 | 0.040 | 0.021 | 0.041 | 0.043 |
| | 5 | 25/100M | 50bp | 0.437 | 0.633 | 0.432 | 0.514 | 0.382 | 0.018 | 0.038 | 0.016 | 0.034 | 0.045 |
| | 5 | 25/100M | 75bp | 0.454 | 0.644 | 0.443 | 0.530 | 0.362 | 0.016 | 0.038 | 0.014 | 0.032 | 0.038 |
| | 5 | 25/100M | 100bp | 0.463 | 0.650 | 0.447 | 0.538 | 0.391 | 0.019 | 0.038 | 0.016 | 0.032 | 0.044 |
| | 5 | 25/100M | 125bp | 0.488 | 0.663 | 0.467 | 0.558 | 0.388 | 0.019 | 0.037 | 0.017 | 0.035 | 0.045 |
| | 5 | 25/100M | 150bp | 0.505 | 0.674 | 0.479 | 0.572 | 0.388 | 0.018 | 0.039 | 0.016 | 0.034 | 0.041 |
| | 10 | 50M | 50bp | 0.799 | 0.818 | 0.757 | 0.765 | 0.818 | 0.032 | 0.042 | 0.026 | 0.034 | 0.048 |
| | 10 | 50M | 75bp | 0.816 | 0.832 | 0.776 | 0.784 | 0.830 | 0.032 | 0.042 | 0.026 | 0.033 | 0.046 |
| | 10 | 50M | 100bp | 0.826 | 0.841 | 0.788 | 0.797 | 0.834 | 0.031 | 0.042 | 0.027 | 0.035 | 0.046 |
| | 10 | 50M | 125bp | 0.843 | 0.856 | 0.811 | 0.820 | 0.847 | 0.033 | 0.043 | 0.027 | 0.035 | 0.046 |
| | 10 | 50M | 150bp | 0.849 | 0.862 | 0.821 | 0.830 | 0.852 | 0.031 | 0.041 | 0.027 | 0.034 | 0.046 |
| | 10 | 25/100M | 50bp | 0.769 | 0.830 | 0.691 | 0.698 | 0.803 | 0.028 | 0.038 | 0.022 | 0.030 | 0.046 |
| | 10 | 25/100M | 75bp | 0.783 | 0.839 | 0.710 | 0.717 | 0.810 | 0.028 | 0.038 | 0.021 | 0.030 | 0.045 |
| | 10 | 25/100M | 100bp | 0.789 | 0.847 | 0.720 | 0.728 | 0.816 | 0.028 | 0.038 | 0.023 | 0.030 | 0.046 |
| | 10 | 25/100M | 125bp | 0.813 | 0.864 | 0.751 | 0.760 | 0.831 | 0.029 | 0.040 | 0.023 | 0.031 | 0.047 |
| | 10 | 25/100M | 150bp | 0.823 | 0.870 | 0.763 | 0.772 | 0.835 | 0.028 | 0.039 | 0.023 | 0.030 | 0.046 |

Table S6: Simulation results - observed power and false discovery rate for different read types and read Lengths, averaged over 20 simulations. Scenario with mm39 genome, maximum of 4 transcripts/gene expressed, and reads quantified with kallisto. Library size shown in million reads (M) with 25/100 indicating library sizes alternating between 25M and 100M across replicates. Read lengths are shown in base pairs (bp). Red color indicates observed FDR values greater than the nominal 0.05. Blue color indicates the most powerful method that controls the FDR at the nominal level for a given scenario (row). Empty cells indicate cases in which a method failed to call any transcript as DE.

| | | | | Power | | | | | False Discovery Rate | | | | |
|---|---|---|---|---|---|---|---|---|---|---|---|---|---|
| Read | Samples/Group | Library Size | Read Length | edgeR-raw | edgeR-scaled | sleuth-LRT | sleuth-Wald | Swish | edgeR-raw | edgeR-scaled | sleuth-LRT | sleuth-Wald | Swish |
| paired-end | 3 | 50M | 50bp | 0.026 | 0.435 | 0.000 | 0.396 | 0.448 | 0.155 | 0.038 | - | 0.055 | 0.257 |
| | 3 | 50M | 75bp | 0.047 | 0.436 | 0.000 | 0.405 | 0.455 | 0.128 | 0.040 | - | 0.056 | 0.261 |
| | 3 | 50M | 100bp | 0.049 | 0.440 | 0.000 | 0.406 | 0.458 | 0.122 | 0.039 | - | 0.053 | 0.255 |
| | 3 | 50M | 125bp | 0.057 | 0.447 | 0.000 | 0.408 | 0.459 | 0.113 | 0.040 | - | 0.056 | 0.262 |
| | 3 | 50M | 150bp | 0.065 | 0.447 | 0.000 | 0.413 | 0.459 | 0.110 | 0.041 | - | 0.054 | 0.258 |
| | 3 | 25/100M | 50bp | 0.000 | 0.369 | 0.000 | 0.330 | 0.382 | 0.590 | 0.034 | - | 0.047 | 0.243 |
| | 3 | 25/100M | 75bp | 0.000 | 0.390 | 0.000 | 0.345 | 0.394 | 0.636 | 0.035 | - | 0.048 | 0.235 |
| | 3 | 25/100M | 100bp | 0.000 | 0.392 | 0.000 | 0.347 | 0.395 | 0.474 | 0.036 | - | 0.049 | 0.242 |
| | 3 | 25/100M | 125bp | 0.005 | 0.392 | 0.000 | 0.351 | 0.397 | 0.245 | 0.037 | - | 0.050 | 0.241 |
| | 3 | 25/100M | 150bp | 0.002 | 0.390 | 0.000 | 0.350 | 0.396 | 0.209 | 0.036 | - | 0.049 | 0.243 |
| | 5 | 50M | 50bp | 0.572 | 0.719 | 0.551 | 0.644 | 0.414 | 0.026 | 0.042 | 0.021 | 0.041 | 0.041 |
| | 5 | 50M | 75bp | 0.588 | 0.731 | 0.563 | 0.658 | 0.431 | 0.026 | 0.042 | 0.021 | 0.041 | 0.045 |
| | 5 | 50M | 100bp | 0.593 | 0.734 | 0.565 | 0.662 | 0.431 | 0.025 | 0.043 | 0.020 | 0.041 | 0.041 |
| | 5 | 50M | 125bp | 0.591 | 0.732 | 0.563 | 0.662 | 0.422 | 0.026 | 0.041 | 0.022 | 0.040 | 0.041 |
| | 5 | 50M | 150bp | 0.600 | 0.736 | 0.572 | 0.668 | 0.434 | 0.027 | 0.042 | 0.021 | 0.041 | 0.044 |
| | 5 | 25/100M | 50bp | 0.512 | 0.671 | 0.478 | 0.574 | 0.393 | 0.021 | 0.037 | 0.016 | 0.034 | 0.046 |
| | 5 | 25/100M | 75bp | 0.533 | 0.689 | 0.495 | 0.593 | 0.395 | 0.023 | 0.038 | 0.018 | 0.036 | 0.040 |
| | 5 | 25/100M | 100bp | 0.538 | 0.686 | 0.501 | 0.597 | 0.387 | 0.022 | 0.039 | 0.017 | 0.037 | 0.039 |
| | 5 | 25/100M | 125bp | 0.540 | 0.690 | 0.497 | 0.595 | 0.392 | 0.022 | 0.038 | 0.017 | 0.037 | 0.042 |
| | 5 | 25/100M | 150bp | 0.551 | 0.694 | 0.505 | 0.602 | 0.398 | 0.023 | 0.039 | 0.017 | 0.036 | 0.043 |
| | 10 | 50M | 50bp | 0.859 | 0.873 | 0.834 | 0.843 | 0.857 | 0.034 | 0.044 | 0.028 | 0.036 | 0.048 |
| | 10 | 50M | 75bp | 0.866 | 0.880 | 0.841 | 0.850 | 0.861 | 0.035 | 0.043 | 0.029 | 0.037 | 0.050 |
| | 10 | 50M | 100bp | 0.871 | 0.886 | 0.847 | 0.855 | 0.864 | 0.032 | 0.042 | 0.027 | 0.035 | 0.046 |
| | 10 | 50M | 125bp | 0.877 | 0.891 | 0.854 | 0.862 | 0.869 | 0.034 | 0.043 | 0.028 | 0.036 | 0.048 |
| | 10 | 50M | 150bp | 0.877 | 0.890 | 0.853 | 0.862 | 0.870 | 0.035 | 0.042 | 0.028 | 0.036 | 0.046 |
| | 10 | 25/100M | 50bp | 0.834 | 0.880 | 0.778 | 0.788 | 0.840 | 0.030 | 0.039 | 0.024 | 0.032 | 0.046 |
| | 10 | 25/100M | 75bp | 0.844 | 0.889 | 0.791 | 0.800 | 0.849 | 0.030 | 0.039 | 0.025 | 0.031 | 0.048 |
| | 10 | 25/100M | 100bp | 0.847 | 0.893 | 0.795 | 0.803 | 0.850 | 0.030 | 0.040 | 0.025 | 0.032 | 0.047 |
| | 10 | 25/100M | 125bp | 0.849 | 0.893 | 0.800 | 0.809 | 0.851 | 0.030 | 0.039 | 0.024 | 0.031 | 0.046 |
| | 10 | 25/100M | 150bp | 0.852 | 0.896 | 0.802 | 0.811 | 0.853 | 0.031 | 0.041 | 0.026 | 0.033 | 0.048 |

Table S6: Simulation results - observed power and false discovery rate for different read types and read Lengths, averaged over 20 simulations. Scenario with mm39 genome, maximum of 4 transcripts/gene expressed, and reads quantified with kallisto. Library size shown in million reads (M) with 25/100 indicating library sizes alternating between 25M and 100M across replicates. Read lengths are shown in base pairs (bp). Red color indicates observed FDR values greater than the nominal 0.05. Blue color indicates the most powerful method that controls the FDR at the nominal level for a given scenario (row). Empty cells indicate cases in which a method failed to call any transcript as DE. *(continued)*

| | | | | Power | | | | | False Discovery Rate | | | | |
|---|---|---|---|---|---|---|---|---|---|---|---|---|---|
| Read | Samples/Group | Library Size | Read Length | edgeR-raw | edgeR-scaled | sleuth-LRT | sleuth-Wald | Swish | edgeR-raw | edgeR-scaled | sleuth-LRT | sleuth-Wald | Swish |
| single-end | 3 | 50M | 50bp | 0.001 | 0.406 | 0.000 | 0.365 | 0.415 | 0.801 | 0.047 | - | 0.060 | 0.253 |
| | 3 | 50M | 75bp | 0.000 | 0.406 | 0.000 | 0.371 | 0.421 | 0.771 | 0.044 | - | 0.059 | 0.253 |
| | 3 | 50M | 100bp | 0.000 | 0.416 | 0.000 | 0.377 | 0.429 | 0.754 | 0.044 | - | 0.057 | 0.255 |
| | 3 | 50M | 125bp | 0.000 | 0.423 | 0.000 | 0.392 | 0.442 | 0.809 | 0.045 | - | 0.060 | 0.258 |
| | 3 | 50M | 150bp | 0.007 | 0.431 | 0.000 | 0.398 | 0.449 | 0.301 | 0.046 | - | 0.061 | 0.257 |
| | 3 | 25/100M | 50bp | 0.000 | 0.350 | 0.000 | 0.298 | 0.354 | 0.800 | 0.039 | 1.000 | 0.051 | 0.239 |
| | 3 | 25/100M | 75bp | 0.000 | 0.359 | 0.000 | 0.310 | 0.360 | 0.824 | 0.041 | - | 0.053 | 0.237 |
| | 3 | 25/100M | 100bp | 0.000 | 0.371 | 0.000 | 0.320 | 0.370 | 0.618 | 0.040 | - | 0.049 | 0.233 |
| | 3 | 25/100M | 125bp | 0.000 | 0.375 | 0.000 | 0.327 | 0.377 | 0.871 | 0.038 | - | 0.047 | 0.236 |
| | 3 | 25/100M | 150bp | 0.000 | 0.375 | 0.000 | 0.331 | 0.381 | 0.645 | 0.036 | - | 0.048 | 0.239 |
| | 5 | 50M | 50bp | 0.499 | 0.689 | 0.505 | 0.600 | 0.404 | 0.032 | 0.054 | 0.028 | 0.049 | 0.051 |
| | 5 | 50M | 75bp | 0.515 | 0.696 | 0.513 | 0.611 | 0.408 | 0.035 | 0.055 | 0.030 | 0.050 | 0.051 |
| | 5 | 50M | 100bp | 0.527 | 0.702 | 0.522 | 0.618 | 0.396 | 0.034 | 0.053 | 0.029 | 0.050 | 0.048 |
| | 5 | 50M | 125bp | 0.551 | 0.716 | 0.539 | 0.637 | 0.412 | 0.032 | 0.052 | 0.027 | 0.047 | 0.050 |
| | 5 | 50M | 150bp | 0.567 | 0.718 | 0.548 | 0.645 | 0.452 | 0.032 | 0.049 | 0.027 | 0.048 | 0.054 |
| | 5 | 25/100M | 50bp | 0.447 | 0.640 | 0.439 | 0.532 | 0.378 | 0.027 | 0.048 | 0.022 | 0.042 | 0.051 |
| | 5 | 25/100M | 75bp | 0.462 | 0.651 | 0.448 | 0.544 | 0.376 | 0.025 | 0.045 | 0.020 | 0.040 | 0.047 |
| | 5 | 25/100M | 100bp | 0.478 | 0.659 | 0.453 | 0.553 | 0.384 | 0.026 | 0.045 | 0.021 | 0.040 | 0.050 |
| | 5 | 25/100M | 125bp | 0.498 | 0.668 | 0.470 | 0.567 | 0.386 | 0.028 | 0.044 | 0.022 | 0.042 | 0.050 |
| | 5 | 25/100M | 150bp | 0.511 | 0.678 | 0.481 | 0.578 | 0.389 | 0.027 | 0.045 | 0.021 | 0.041 | 0.046 |
| | 10 | 50M | 50bp | 0.812 | 0.834 | 0.776 | 0.785 | 0.818 | 0.056 | 0.064 | 0.046 | 0.055 | 0.074 |
| | 10 | 50M | 75bp | 0.825 | 0.847 | 0.791 | 0.800 | 0.829 | 0.054 | 0.061 | 0.043 | 0.052 | 0.069 |
| | 10 | 50M | 100bp | 0.834 | 0.855 | 0.803 | 0.812 | 0.836 | 0.052 | 0.060 | 0.043 | 0.052 | 0.069 |
| | 10 | 50M | 125bp | 0.849 | 0.866 | 0.821 | 0.829 | 0.846 | 0.052 | 0.060 | 0.042 | 0.050 | 0.066 |
| | 10 | 50M | 150bp | 0.854 | 0.871 | 0.828 | 0.837 | 0.849 | 0.048 | 0.057 | 0.039 | 0.047 | 0.063 |
| | 10 | 25/100M | 50bp | 0.785 | 0.846 | 0.720 | 0.727 | 0.805 | 0.050 | 0.061 | 0.039 | 0.048 | 0.072 |
| | 10 | 25/100M | 75bp | 0.796 | 0.854 | 0.734 | 0.742 | 0.814 | 0.048 | 0.059 | 0.037 | 0.045 | 0.070 |
| | 10 | 25/100M | 100bp | 0.805 | 0.862 | 0.746 | 0.754 | 0.821 | 0.048 | 0.058 | 0.037 | 0.045 | 0.069 |
| | 10 | 25/100M | 125bp | 0.821 | 0.875 | 0.767 | 0.775 | 0.832 | 0.048 | 0.057 | 0.036 | 0.044 | 0.066 |
| | 10 | 25/100M | 150bp | 0.827 | 0.878 | 0.775 | 0.783 | 0.836 | 0.045 | 0.055 | 0.035 | 0.043 | 0.064 |

Table S7: Simulation results - observed power and false discovery rate for different read types and read Lengths, averaged over 20 simulations. Scenario with mm39 genome, maximum of 5 transcripts/gene expressed, and reads quantified with Salmon. Library size shown in million reads (M) with 25/100 indicating library sizes alternating between 25M and 100M across replicates. Read lengths are shown in base pairs (bp). Red color indicates observed FDR values greater than the nominal 0.05. Blue color indicates the most powerful method that controls the FDR at the nominal level for a given scenario (row). Empty cells indicate cases in which a method failed to call any transcript as DE.

| | | | | Power | | | | | False Discovery Rate | | | | |
|---|---|---|---|---|---|---|---|---|---|---|---|---|---|
| Read | Samples/Group | Library Size | Read Length | edgeR-raw | edgeR-scaled | sleuth-LRT | sleuth-Wald | Swish | edgeR-raw | edgeR-scaled | sleuth-LRT | sleuth-Wald | Swish |
| paired-end | 3 | 50M | 50bp | 0.005 | 0.437 | 0.000 | 0.407 | 0.449 | 0.477 | 0.038 | - | 0.054 | 0.238 |
| | 3 | 50M | 75bp | 0.025 | 0.453 | 0.000 | 0.418 | 0.454 | 0.211 | 0.042 | - | 0.058 | 0.240 |
| | 3 | 50M | 100bp | 0.014 | 0.440 | 0.000 | 0.406 | 0.440 | 0.278 | 0.038 | - | 0.055 | 0.236 |
| | 3 | 50M | 125bp | 0.027 | 0.450 | 0.000 | 0.425 | 0.464 | 0.197 | 0.039 | - | 0.056 | 0.243 |
| | 3 | 50M | 150bp | 0.027 | 0.448 | 0.000 | 0.420 | 0.458 | 0.196 | 0.040 | 0.000 | 0.054 | 0.247 |
| | 3 | 25/100M | 50bp | 0.000 | 0.386 | 0.000 | 0.341 | 0.387 | 0.847 | 0.036 | - | 0.044 | 0.226 |
| | 3 | 25/100M | 75bp | 0.001 | 0.392 | 0.000 | 0.349 | 0.393 | 0.702 | 0.036 | 1.000 | 0.045 | 0.224 |
| | 3 | 25/100M | 100bp | 0.001 | 0.396 | 0.000 | 0.344 | 0.383 | 0.639 | 0.037 | - | 0.047 | 0.221 |
| | 3 | 25/100M | 125bp | 0.001 | 0.396 | 0.000 | 0.354 | 0.397 | 0.689 | 0.037 | - | 0.047 | 0.231 |
| | 3 | 25/100M | 150bp | 0.001 | 0.400 | 0.000 | 0.359 | 0.402 | 0.689 | 0.035 | - | 0.046 | 0.228 |
| | 5 | 50M | 50bp | 0.578 | 0.743 | 0.569 | 0.667 | 0.464 | 0.025 | 0.042 | 0.023 | 0.041 | 0.047 |
| | 5 | 50M | 75bp | 0.595 | 0.746 | 0.573 | 0.674 | 0.436 | 0.021 | 0.041 | 0.020 | 0.039 | 0.042 |
| | 5 | 50M | 100bp | 0.596 | 0.746 | 0.573 | 0.672 | 0.448 | 0.022 | 0.041 | 0.021 | 0.040 | 0.043 |
| | 5 | 50M | 125bp | 0.612 | 0.754 | 0.589 | 0.684 | 0.473 | 0.024 | 0.042 | 0.022 | 0.041 | 0.046 |
| | 5 | 50M | 150bp | 0.612 | 0.754 | 0.584 | 0.686 | 0.450 | 0.022 | 0.040 | 0.021 | 0.040 | 0.042 |
| | 5 | 25/100M | 50bp | 0.511 | 0.700 | 0.495 | 0.597 | 0.396 | 0.019 | 0.039 | 0.017 | 0.035 | 0.041 |
| | 5 | 25/100M | 75bp | 0.527 | 0.699 | 0.501 | 0.603 | 0.414 | 0.018 | 0.038 | 0.017 | 0.034 | 0.044 |
| | 5 | 25/100M | 100bp | 0.527 | 0.699 | 0.497 | 0.597 | 0.403 | 0.018 | 0.040 | 0.017 | 0.034 | 0.042 |
| | 5 | 25/100M | 125bp | 0.547 | 0.714 | 0.517 | 0.620 | 0.425 | 0.018 | 0.039 | 0.017 | 0.035 | 0.044 |
| | 5 | 25/100M | 150bp | 0.536 | 0.707 | 0.508 | 0.611 | 0.399 | 0.019 | 0.039 | 0.016 | 0.035 | 0.044 |
| | 10 | 50M | 50bp | 0.879 | 0.891 | 0.848 | 0.856 | 0.883 | 0.034 | 0.045 | 0.028 | 0.037 | 0.049 |
| | 10 | 50M | 75bp | 0.886 | 0.896 | 0.855 | 0.863 | 0.884 | 0.031 | 0.042 | 0.027 | 0.034 | 0.046 |
| | 10 | 50M | 100bp | 0.887 | 0.896 | 0.856 | 0.865 | 0.885 | 0.031 | 0.042 | 0.027 | 0.034 | 0.046 |
| | 10 | 50M | 125bp | 0.896 | 0.904 | 0.868 | 0.876 | 0.892 | 0.031 | 0.041 | 0.026 | 0.033 | 0.046 |
| | 10 | 50M | 150bp | 0.894 | 0.902 | 0.865 | 0.874 | 0.890 | 0.031 | 0.041 | 0.026 | 0.034 | 0.046 |
| | 10 | 25/100M | 50bp | 0.851 | 0.896 | 0.795 | 0.802 | 0.866 | 0.030 | 0.041 | 0.024 | 0.031 | 0.049 |
| | 10 | 25/100M | 75bp | 0.861 | 0.899 | 0.801 | 0.810 | 0.867 | 0.028 | 0.039 | 0.023 | 0.031 | 0.047 |
| | 10 | 25/100M | 100bp | 0.857 | 0.899 | 0.801 | 0.810 | 0.867 | 0.027 | 0.037 | 0.022 | 0.029 | 0.046 |
| | 10 | 25/100M | 125bp | 0.869 | 0.906 | 0.817 | 0.825 | 0.874 | 0.029 | 0.041 | 0.023 | 0.031 | 0.048 |
| | 10 | 25/100M | 150bp | 0.870 | 0.908 | 0.817 | 0.825 | 0.876 | 0.028 | 0.039 | 0.023 | 0.031 | 0.047 |

Table S7: Simulation results - observed power and false discovery rate for different read types and read Lengths, averaged over 20 simulations. Scenario with mm39 genome, maximum of 5 transcripts/gene expressed, and reads quantified with Salmon. Library size shown in million reads (M) with 25/100 indicating library sizes alternating between 25M and 100M across replicates. Read lengths are shown in base pairs (bp). Red color indicates observed FDR values greater than the nominal 0.05. Blue color indicates the most powerful method that controls the FDR at the nominal level for a given scenario (row). Empty cells indicate cases in which a method failed to call any transcript as DE. *(continued)*

| | | | | Power | | | | | False Discovery Rate | | | | |
|---|---|---|---|---|---|---|---|---|---|---|---|---|---|
| Read | Samples/Group | Library Size | Read Length | edgeR-raw | edgeR-scaled | sleuth-LRT | sleuth-Wald | Swish | edgeR-raw | edgeR-scaled | sleuth-LRT | sleuth-Wald | Swish |
| single-end | 3 | 50M | 50bp | 0.002 | 0.399 | 0.000 | 0.370 | 0.414 | 0.661 | 0.037 | 1.000 | 0.054 | 0.237 |
| | 3 | 50M | 75bp | 0.002 | 0.420 | 0.000 | 0.381 | 0.421 | 0.700 | 0.039 | - | 0.053 | 0.234 |
| | 3 | 50M | 100bp | 0.002 | 0.432 | 0.000 | 0.385 | 0.423 | 0.655 | 0.040 | - | 0.054 | 0.233 |
| | 3 | 50M | 125bp | 0.004 | 0.429 | 0.000 | 0.401 | 0.441 | 0.544 | 0.038 | - | 0.055 | 0.240 |
| | 3 | 50M | 150bp | 0.008 | 0.441 | 0.000 | 0.411 | 0.448 | 0.406 | 0.038 | 0.000 | 0.055 | 0.238 |
| | 3 | 25/100M | 50bp | 0.000 | 0.362 | 0.000 | 0.305 | 0.357 | 1.000 | 0.035 | - | 0.044 | 0.223 |
| | 3 | 25/100M | 75bp | 0.000 | 0.366 | 0.000 | 0.312 | 0.360 | 0.727 | 0.034 | - | 0.043 | 0.222 |
| | 3 | 25/100M | 100bp | 0.000 | 0.377 | 0.000 | 0.318 | 0.362 | 0.706 | 0.037 | - | 0.044 | 0.226 |
| | 3 | 25/100M | 125bp | 0.000 | 0.376 | 0.000 | 0.333 | 0.377 | 0.757 | 0.034 | - | 0.045 | 0.230 |
| | 3 | 25/100M | 150bp | 0.001 | 0.391 | 0.000 | 0.341 | 0.389 | 0.729 | 0.037 | - | 0.049 | 0.228 |
| | 5 | 50M | 50bp | 0.510 | 0.698 | 0.522 | 0.612 | 0.411 | 0.022 | 0.040 | 0.021 | 0.039 | 0.038 |
| | 5 | 50M | 75bp | 0.525 | 0.709 | 0.527 | 0.622 | 0.432 | 0.022 | 0.041 | 0.020 | 0.039 | 0.045 |
| | 5 | 50M | 100bp | 0.538 | 0.720 | 0.539 | 0.633 | 0.429 | 0.022 | 0.041 | 0.021 | 0.040 | 0.041 |
| | 5 | 50M | 125bp | 0.565 | 0.731 | 0.559 | 0.655 | 0.448 | 0.021 | 0.038 | 0.019 | 0.036 | 0.043 |
| | 5 | 50M | 150bp | 0.580 | 0.739 | 0.569 | 0.667 | 0.422 | 0.021 | 0.038 | 0.021 | 0.041 | 0.038 |
| | 5 | 25/100M | 50bp | 0.436 | 0.650 | 0.437 | 0.529 | 0.385 | 0.019 | 0.038 | 0.016 | 0.032 | 0.045 |
| | 5 | 25/100M | 75bp | 0.449 | 0.660 | 0.443 | 0.541 | 0.396 | 0.017 | 0.038 | 0.015 | 0.033 | 0.045 |
| | 5 | 25/100M | 100bp | 0.463 | 0.669 | 0.454 | 0.553 | 0.411 | 0.017 | 0.036 | 0.015 | 0.031 | 0.045 |
| | 5 | 25/100M | 125bp | 0.495 | 0.686 | 0.482 | 0.580 | 0.398 | 0.017 | 0.037 | 0.015 | 0.033 | 0.040 |
| | 5 | 25/100M | 150bp | 0.513 | 0.694 | 0.495 | 0.594 | 0.413 | 0.018 | 0.037 | 0.018 | 0.034 | 0.044 |
| | 10 | 50M | 50bp | 0.822 | 0.840 | 0.779 | 0.788 | 0.844 | 0.031 | 0.041 | 0.026 | 0.034 | 0.046 |
| | 10 | 50M | 75bp | 0.837 | 0.852 | 0.796 | 0.804 | 0.850 | 0.030 | 0.042 | 0.026 | 0.033 | 0.045 |
| | 10 | 50M | 100bp | 0.851 | 0.866 | 0.813 | 0.821 | 0.862 | 0.031 | 0.042 | 0.027 | 0.035 | 0.047 |
| | 10 | 50M | 125bp | 0.863 | 0.876 | 0.830 | 0.838 | 0.870 | 0.031 | 0.041 | 0.027 | 0.034 | 0.046 |
| | 10 | 50M | 150bp | 0.878 | 0.888 | 0.847 | 0.856 | 0.881 | 0.032 | 0.041 | 0.025 | 0.033 | 0.047 |
| | 10 | 25/100M | 50bp | 0.790 | 0.850 | 0.714 | 0.721 | 0.827 | 0.028 | 0.038 | 0.022 | 0.029 | 0.048 |
| | 10 | 25/100M | 75bp | 0.802 | 0.860 | 0.730 | 0.738 | 0.831 | 0.028 | 0.039 | 0.022 | 0.031 | 0.046 |
| | 10 | 25/100M | 100bp | 0.818 | 0.871 | 0.748 | 0.756 | 0.843 | 0.028 | 0.037 | 0.021 | 0.028 | 0.047 |
| | 10 | 25/100M | 125bp | 0.837 | 0.884 | 0.775 | 0.783 | 0.855 | 0.029 | 0.039 | 0.023 | 0.030 | 0.047 |
| | 10 | 25/100M | 150bp | 0.850 | 0.895 | 0.790 | 0.799 | 0.864 | 0.029 | 0.039 | 0.024 | 0.031 | 0.046 |

Table S8: Simulation results - observed power and false discovery rate for different read types and read Lengths, averaged over 20 simulations. Scenario with mm39 genome, maximum of 5 transcripts/gene expressed, and reads quantified with kallisto. Library size shown in million reads (M) with 25/100 indicating library sizes alternating between 25M and 100M across replicates. Read lengths are shown in base pairs (bp). Red color indicates observed FDR values greater than the nominal 0.05. Blue color indicates the most powerful method that controls the FDR at the nominal level for a given scenario (row). Empty cells indicate cases in which a method failed to call any transcript as DE.

| | | | | Power | | | | | False Discovery Rate | | | | |
|---|---|---|---|---|---|---|---|---|---|---|---|---|---|
| Read | Samples/Group | Library Size | Read Length | edgeR-raw | edgeR-scaled | sleuth-LRT | sleuth-Wald | Swish | edgeR-raw | edgeR-scaled | sleuth-LRT | sleuth-Wald | Swish |
| paired-end | 3 | 50M | 50bp | 0.019 | 0.436 | 0.000 | 0.407 | 0.461 | 0.180 | 0.038 | - | 0.055 | 0.250 |
| | 3 | 50M | 75bp | 0.033 | 0.454 | 0.000 | 0.420 | 0.471 | 0.150 | 0.043 | - | 0.058 | 0.255 |
| | 3 | 50M | 100bp | 0.016 | 0.443 | 0.000 | 0.415 | 0.467 | 0.192 | 0.040 | - | 0.054 | 0.256 |
| | 3 | 50M | 125bp | 0.021 | 0.453 | 0.000 | 0.425 | 0.477 | 0.175 | 0.040 | - | 0.054 | 0.255 |
| | 3 | 50M | 150bp | 0.027 | 0.451 | 0.000 | 0.422 | 0.473 | 0.149 | 0.041 | - | 0.054 | 0.258 |
| | 3 | 25/100M | 50bp | 0.000 | 0.384 | 0.000 | 0.341 | 0.396 | 0.474 | 0.035 | - | 0.045 | 0.235 |
| | 3 | 25/100M | 75bp | 0.000 | 0.391 | 0.000 | 0.353 | 0.403 | 0.545 | 0.036 | - | 0.045 | 0.237 |
| | 3 | 25/100M | 100bp | 0.000 | 0.402 | 0.000 | 0.358 | 0.405 | 0.810 | 0.039 | - | 0.048 | 0.238 |
| | 3 | 25/100M | 125bp | 0.000 | 0.397 | 0.000 | 0.356 | 0.406 | 0.618 | 0.037 | - | 0.047 | 0.237 |
| | 3 | 25/100M | 150bp | 0.000 | 0.403 | 0.000 | 0.361 | 0.411 | 0.744 | 0.036 | - | 0.047 | 0.237 |
| | 5 | 50M | 50bp | 0.591 | 0.743 | 0.571 | 0.672 | 0.458 | 0.026 | 0.042 | 0.021 | 0.039 | 0.045 |
| | 5 | 50M | 75bp | 0.603 | 0.750 | 0.579 | 0.682 | 0.442 | 0.025 | 0.042 | 0.020 | 0.040 | 0.042 |
| | 5 | 50M | 100bp | 0.613 | 0.757 | 0.586 | 0.688 | 0.448 | 0.025 | 0.042 | 0.021 | 0.040 | 0.043 |
| | 5 | 50M | 125bp | 0.618 | 0.759 | 0.591 | 0.690 | 0.459 | 0.028 | 0.045 | 0.022 | 0.042 | 0.044 |
| | 5 | 50M | 150bp | 0.617 | 0.759 | 0.587 | 0.693 | 0.468 | 0.025 | 0.041 | 0.021 | 0.040 | 0.046 |
| | 5 | 25/100M | 50bp | 0.530 | 0.696 | 0.496 | 0.603 | 0.407 | 0.022 | 0.040 | 0.017 | 0.036 | 0.044 |
| | 5 | 25/100M | 75bp | 0.544 | 0.705 | 0.509 | 0.614 | 0.406 | 0.021 | 0.039 | 0.016 | 0.035 | 0.043 |
| | 5 | 25/100M | 100bp | 0.552 | 0.712 | 0.517 | 0.618 | 0.403 | 0.021 | 0.041 | 0.017 | 0.036 | 0.042 |
| | 5 | 25/100M | 125bp | 0.558 | 0.719 | 0.519 | 0.626 | 0.435 | 0.021 | 0.039 | 0.017 | 0.036 | 0.046 |
| | 5 | 25/100M | 150bp | 0.552 | 0.712 | 0.514 | 0.621 | 0.403 | 0.022 | 0.039 | 0.017 | 0.036 | 0.046 |
| | 10 | 50M | 50bp | 0.886 | 0.894 | 0.858 | 0.867 | 0.881 | 0.034 | 0.044 | 0.028 | 0.037 | 0.047 |
| | 10 | 50M | 75bp | 0.894 | 0.904 | 0.867 | 0.875 | 0.885 | 0.032 | 0.042 | 0.028 | 0.035 | 0.047 |
| | 10 | 50M | 100bp | 0.899 | 0.911 | 0.875 | 0.883 | 0.891 | 0.033 | 0.044 | 0.028 | 0.035 | 0.047 |
| | 10 | 50M | 125bp | 0.903 | 0.913 | 0.878 | 0.886 | 0.895 | 0.032 | 0.043 | 0.027 | 0.034 | 0.048 |
| | 10 | 50M | 150bp | 0.902 | 0.912 | 0.877 | 0.885 | 0.893 | 0.032 | 0.042 | 0.027 | 0.035 | 0.047 |
| | 10 | 25/100M | 50bp | 0.862 | 0.902 | 0.807 | 0.815 | 0.867 | 0.031 | 0.041 | 0.024 | 0.033 | 0.049 |
| | 10 | 25/100M | 75bp | 0.871 | 0.908 | 0.818 | 0.826 | 0.871 | 0.030 | 0.041 | 0.024 | 0.032 | 0.048 |
| | 10 | 25/100M | 100bp | 0.875 | 0.914 | 0.825 | 0.833 | 0.877 | 0.030 | 0.039 | 0.024 | 0.031 | 0.048 |
| | 10 | 25/100M | 125bp | 0.878 | 0.916 | 0.830 | 0.838 | 0.878 | 0.031 | 0.041 | 0.025 | 0.032 | 0.048 |
| | 10 | 25/100M | 150bp | 0.880 | 0.917 | 0.831 | 0.839 | 0.880 | 0.030 | 0.040 | 0.025 | 0.033 | 0.048 |

Table S8: Simulation results - observed power and false discovery rate for different read types and read Lengths, averaged over 20 simulations. Scenario with mm39 genome, maximum of 5 transcripts/gene expressed, and reads quantified with kallisto. Library size shown in million reads (M) with 25/100 indicating library sizes alternating between 25M and 100M across replicates. Read lengths are shown in base pairs (bp). Red color indicates observed FDR values greater than the nominal 0.05. Blue color indicates the most powerful method that controls the FDR at the nominal level for a given scenario (row). Empty cells indicate cases in which a method failed to call any transcript as DE. *(continued)*

| | | | | Power | | | | | False Discovery Rate | | | | |
|---|---|---|---|---|---|---|---|---|---|---|---|---|---|
| Read | Samples/Group | Library Size | Read Length | edgeR-raw | edgeR-scaled | sleuth-LRT | sleuth-Wald | Swish | edgeR-raw | edgeR-scaled | sleuth-LRT | sleuth-Wald | Swish |
| single-end | 3 | 50M | 50bp | 0.000 | 0.404 | 0.000 | 0.372 | 0.426 | 0.762 | 0.047 | - | 0.059 | 0.249 |
| | 3 | 50M | 75bp | 0.000 | 0.417 | 0.000 | 0.383 | 0.432 | 0.812 | 0.046 | - | 0.060 | 0.249 |
| | 3 | 50M | 100bp | 0.000 | 0.429 | 0.000 | 0.389 | 0.441 | 0.848 | 0.046 | - | 0.059 | 0.248 |
| | 3 | 50M | 125bp | 0.000 | 0.432 | 0.000 | 0.400 | 0.452 | 0.817 | 0.045 | 1.000 | 0.057 | 0.253 |
| | 3 | 50M | 150bp | 0.005 | 0.441 | 0.000 | 0.409 | 0.460 | 0.361 | 0.045 | - | 0.060 | 0.252 |
| | 3 | 25/100M | 50bp | 0.000 | 0.358 | 0.000 | 0.311 | 0.361 | 0.857 | 0.040 | - | 0.049 | 0.233 |
| | 3 | 25/100M | 75bp | 0.000 | 0.365 | 0.000 | 0.317 | 0.368 | 0.938 | 0.039 | - | 0.048 | 0.234 |
| | 3 | 25/100M | 100bp | 0.000 | 0.373 | 0.000 | 0.324 | 0.375 | 0.588 | 0.042 | - | 0.051 | 0.237 |
| | 3 | 25/100M | 125bp | 0.000 | 0.373 | 0.000 | 0.336 | 0.385 | 0.762 | 0.037 | - | 0.049 | 0.238 |
| | 3 | 25/100M | 150bp | 0.000 | 0.389 | 0.000 | 0.342 | 0.396 | 0.833 | 0.039 | 1.000 | 0.052 | 0.236 |
| | 5 | 50M | 50bp | 0.515 | 0.708 | 0.525 | 0.625 | 0.420 | 0.034 | 0.053 | 0.028 | 0.049 | 0.048 |
| | 5 | 50M | 75bp | 0.528 | 0.713 | 0.530 | 0.633 | 0.423 | 0.034 | 0.053 | 0.029 | 0.049 | 0.052 |
| | 5 | 50M | 100bp | 0.546 | 0.727 | 0.543 | 0.646 | 0.429 | 0.034 | 0.053 | 0.029 | 0.049 | 0.048 |
| | 5 | 50M | 125bp | 0.570 | 0.736 | 0.558 | 0.662 | 0.434 | 0.031 | 0.048 | 0.025 | 0.044 | 0.049 |
| | 5 | 50M | 150bp | 0.583 | 0.742 | 0.569 | 0.671 | 0.448 | 0.031 | 0.048 | 0.026 | 0.048 | 0.050 |
| | 5 | 25/100M | 50bp | 0.450 | 0.659 | 0.445 | 0.548 | 0.394 | 0.027 | 0.048 | 0.023 | 0.043 | 0.054 |
| | 5 | 25/100M | 75bp | 0.464 | 0.669 | 0.452 | 0.557 | 0.401 | 0.026 | 0.046 | 0.022 | 0.040 | 0.053 |
| | 5 | 25/100M | 100bp | 0.484 | 0.679 | 0.463 | 0.570 | 0.392 | 0.024 | 0.043 | 0.021 | 0.038 | 0.049 |
| | 5 | 25/100M | 125bp | 0.509 | 0.690 | 0.487 | 0.592 | 0.413 | 0.024 | 0.044 | 0.020 | 0.039 | 0.047 |
| | 5 | 25/100M | 150bp | 0.525 | 0.699 | 0.498 | 0.602 | 0.397 | 0.025 | 0.043 | 0.021 | 0.039 | 0.045 |
| | 10 | 50M | 50bp | 0.839 | 0.858 | 0.803 | 0.811 | 0.844 | 0.055 | 0.063 | 0.044 | 0.054 | 0.071 |
| | 10 | 50M | 75bp | 0.851 | 0.868 | 0.815 | 0.824 | 0.851 | 0.052 | 0.060 | 0.043 | 0.051 | 0.068 |
| | 10 | 50M | 100bp | 0.866 | 0.882 | 0.833 | 0.841 | 0.865 | 0.051 | 0.058 | 0.042 | 0.051 | 0.068 |
| | 10 | 50M | 125bp | 0.872 | 0.887 | 0.843 | 0.851 | 0.868 | 0.050 | 0.057 | 0.040 | 0.049 | 0.066 |
| | 10 | 50M | 150bp | 0.885 | 0.898 | 0.858 | 0.866 | 0.878 | 0.049 | 0.057 | 0.038 | 0.047 | 0.064 |
| | 10 | 25/100M | 50bp | 0.810 | 0.866 | 0.744 | 0.751 | 0.829 | 0.049 | 0.060 | 0.036 | 0.044 | 0.070 |
| | 10 | 25/100M | 75bp | 0.820 | 0.875 | 0.760 | 0.767 | 0.834 | 0.048 | 0.059 | 0.037 | 0.045 | 0.068 |
| | 10 | 25/100M | 100bp | 0.835 | 0.887 | 0.776 | 0.784 | 0.849 | 0.047 | 0.057 | 0.036 | 0.044 | 0.068 |
| | 10 | 25/100M | 125bp | 0.849 | 0.895 | 0.794 | 0.802 | 0.856 | 0.046 | 0.056 | 0.033 | 0.042 | 0.065 |
| | 10 | 25/100M | 150bp | 0.860 | 0.904 | 0.807 | 0.815 | 0.863 | 0.045 | 0.054 | 0.035 | 0.043 | 0.062 |

Table S9: Simulation results - observed power and false discovery rate for different read types and read Lengths, averaged over 20 simulations. Scenario with mm39 genome, all transcripts expressed, and reads quantified with Salmon. Library size shown in million reads (M) with 25/100 indicating library sizes alternating between 25M and 100M across replicates. Read lengths are shown in base pairs (bp). Red color indicates observed FDR values greater than the nominal 0.05. Blue color indicates the most powerful method that controls the FDR at the nominal level for a given scenario (row). Empty cells indicate cases in which a method failed to call any transcript as DE.

| | | | | Power | | | | | False Discovery Rate | | | | |
|---|---|---|---|---|---|---|---|---|---|---|---|---|---|
| Read | Samples/Group | Library Size | Read Length | edgeR-raw | edgeR-scaled | sleuth-LRT | sleuth-Wald | Swish | edgeR-raw | edgeR-scaled | sleuth-LRT | sleuth-Wald | Swish |
| paired-end | 3 | 50M | 50bp | 0.004 | 0.433 | 0.000 | 0.417 | 0.462 | 0.640 | 0.040 | - | 0.059 | 0.236 |
| | 3 | 50M | 75bp | 0.003 | 0.440 | 0.000 | 0.422 | 0.466 | 0.664 | 0.038 | 0.500 | 0.056 | 0.235 |
| | 3 | 50M | 100bp | 0.004 | 0.441 | 0.000 | 0.414 | 0.453 | 0.638 | 0.040 | - | 0.053 | 0.231 |
| | 3 | 50M | 125bp | 0.004 | 0.440 | 0.000 | 0.428 | 0.475 | 0.625 | 0.039 | - | 0.054 | 0.238 |
| | 3 | 50M | 150bp | 0.006 | 0.444 | 0.000 | 0.430 | 0.475 | 0.570 | 0.040 | - | 0.057 | 0.239 |
| | 3 | 25/100M | 50bp | 0.000 | 0.378 | 0.000 | 0.346 | 0.398 | 0.812 | 0.035 | - | 0.048 | 0.223 |
| | 3 | 25/100M | 75bp | 0.000 | 0.389 | 0.000 | 0.357 | 0.402 | 0.761 | 0.035 | - | 0.045 | 0.218 |
| | 3 | 25/100M | 100bp | 0.000 | 0.387 | 0.000 | 0.342 | 0.390 | 0.771 | 0.035 | - | 0.044 | 0.217 |
| | 3 | 25/100M | 125bp | 0.001 | 0.397 | 0.000 | 0.364 | 0.412 | 0.755 | 0.034 | 1.000 | 0.047 | 0.225 |
| | 3 | 25/100M | 150bp | 0.001 | 0.395 | 0.000 | 0.362 | 0.410 | 0.773 | 0.035 | - | 0.048 | 0.225 |
| | 5 | 50M | 50bp | 0.567 | 0.752 | 0.577 | 0.685 | 0.466 | 0.024 | 0.041 | 0.022 | 0.042 | 0.046 |
| | 5 | 50M | 75bp | 0.584 | 0.761 | 0.586 | 0.692 | 0.475 | 0.022 | 0.042 | 0.020 | 0.038 | 0.043 |
| | 5 | 50M | 100bp | 0.576 | 0.760 | 0.577 | 0.687 | 0.477 | 0.021 | 0.042 | 0.018 | 0.039 | 0.043 |
| | 5 | 50M | 125bp | 0.594 | 0.764 | 0.593 | 0.700 | 0.473 | 0.021 | 0.040 | 0.020 | 0.040 | 0.043 |
| | 5 | 50M | 150bp | 0.596 | 0.767 | 0.596 | 0.705 | 0.481 | 0.021 | 0.041 | 0.020 | 0.039 | 0.046 |
| | 5 | 25/100M | 50bp | 0.498 | 0.708 | 0.503 | 0.612 | 0.436 | 0.018 | 0.039 | 0.017 | 0.036 | 0.044 |
| | 5 | 25/100M | 75bp | 0.516 | 0.719 | 0.513 | 0.623 | 0.442 | 0.017 | 0.039 | 0.016 | 0.034 | 0.045 |
| | 5 | 25/100M | 100bp | 0.506 | 0.716 | 0.503 | 0.617 | 0.436 | 0.017 | 0.038 | 0.015 | 0.033 | 0.042 |
| | 5 | 25/100M | 125bp | 0.525 | 0.726 | 0.521 | 0.636 | 0.451 | 0.018 | 0.039 | 0.016 | 0.035 | 0.048 |
| | 5 | 25/100M | 150bp | 0.522 | 0.725 | 0.520 | 0.635 | 0.417 | 0.017 | 0.037 | 0.016 | 0.034 | 0.041 |
| | 10 | 50M | 50bp | 0.894 | 0.904 | 0.863 | 0.871 | 0.899 | 0.033 | 0.045 | 0.029 | 0.037 | 0.051 |
| | 10 | 50M | 75bp | 0.902 | 0.911 | 0.873 | 0.880 | 0.904 | 0.030 | 0.042 | 0.026 | 0.035 | 0.047 |
| | 10 | 50M | 100bp | 0.900 | 0.910 | 0.871 | 0.878 | 0.903 | 0.032 | 0.043 | 0.027 | 0.036 | 0.048 |
| | 10 | 50M | 125bp | 0.908 | 0.916 | 0.880 | 0.888 | 0.907 | 0.031 | 0.042 | 0.027 | 0.035 | 0.045 |
| | 10 | 50M | 150bp | 0.908 | 0.917 | 0.881 | 0.889 | 0.909 | 0.031 | 0.043 | 0.026 | 0.035 | 0.046 |
| | 10 | 25/100M | 50bp | 0.870 | 0.909 | 0.814 | 0.822 | 0.884 | 0.029 | 0.040 | 0.023 | 0.031 | 0.048 |
| | 10 | 25/100M | 75bp | 0.873 | 0.913 | 0.819 | 0.827 | 0.889 | 0.027 | 0.038 | 0.021 | 0.029 | 0.047 |
| | 10 | 25/100M | 100bp | 0.872 | 0.910 | 0.820 | 0.828 | 0.885 | 0.028 | 0.040 | 0.023 | 0.030 | 0.047 |
| | 10 | 25/100M | 125bp | 0.883 | 0.919 | 0.837 | 0.846 | 0.896 | 0.027 | 0.038 | 0.021 | 0.028 | 0.045 |
| | 10 | 25/100M | 150bp | 0.882 | 0.918 | 0.834 | 0.842 | 0.894 | 0.028 | 0.038 | 0.022 | 0.029 | 0.046 |

Table S9: Simulation results - observed power and false discovery rate for different read types and read Lengths, averaged over 20 simulations. Scenario with mm39 genome, all transcripts expressed, and reads quantified with Salmon. Library size shown in million reads (M) with 25/100 indicating library sizes alternating between 25M and 100M across replicates. Read lengths are shown in base pairs (bp). Red color indicates observed FDR values greater than the nominal 0.05. Blue color indicates the most powerful method that controls the FDR at the nominal level for a given scenario (row). Empty cells indicate cases in which a method failed to call any transcript as DE. *(continued)*

| | | | | Power | | | | | False Discovery Rate | | | | |
|---|---|---|---|---|---|---|---|---|---|---|---|---|---|
| Read | Samples/Group | Library Size | Read Length | edgeR-raw | edgeR-scaled | sleuth-LRT | sleuth-Wald | Swish | edgeR-raw | edgeR-scaled | sleuth-LRT | sleuth-Wald | Swish |
| single-end | 3 | 50M | 50bp | 0.001 | 0.396 | 0.000 | 0.375 | 0.424 | 0.701 | 0.038 | - | 0.054 | 0.230 |
| | 3 | 50M | 75bp | 0.002 | 0.418 | 0.000 | 0.386 | 0.430 | 0.743 | 0.040 | - | 0.053 | 0.226 |
| | 3 | 50M | 100bp | 0.003 | 0.421 | 0.000 | 0.388 | 0.431 | 0.712 | 0.040 | - | 0.054 | 0.232 |
| | 3 | 50M | 125bp | 0.003 | 0.424 | 0.000 | 0.411 | 0.455 | 0.692 | 0.038 | 0.000 | 0.055 | 0.234 |
| | 3 | 50M | 150bp | 0.004 | 0.433 | 0.000 | 0.419 | 0.461 | 0.647 | 0.039 | 0.500 | 0.054 | 0.233 |
| | 3 | 25/100M | 50bp | 0.000 | 0.353 | 0.000 | 0.305 | 0.362 | 0.800 | 0.034 | - | 0.045 | 0.220 |
| | 3 | 25/100M | 75bp | 0.000 | 0.357 | 0.000 | 0.309 | 0.362 | 0.818 | 0.033 | - | 0.042 | 0.217 |
| | 3 | 25/100M | 100bp | 0.000 | 0.369 | 0.000 | 0.320 | 0.369 | 0.762 | 0.033 | - | 0.043 | 0.216 |
| | 3 | 25/100M | 125bp | 0.000 | 0.381 | 0.000 | 0.335 | 0.383 | 0.800 | 0.037 | - | 0.046 | 0.224 |
| | 3 | 25/100M | 150bp | 0.000 | 0.386 | 0.000 | 0.343 | 0.394 | 0.798 | 0.035 | - | 0.047 | 0.225 |
| | 5 | 50M | 50bp | 0.486 | 0.702 | 0.520 | 0.622 | 0.430 | 0.020 | 0.039 | 0.020 | 0.040 | 0.041 |
| | 5 | 50M | 75bp | 0.508 | 0.722 | 0.532 | 0.639 | 0.453 | 0.022 | 0.040 | 0.020 | 0.039 | 0.045 |
| | 5 | 50M | 100bp | 0.520 | 0.726 | 0.542 | 0.645 | 0.439 | 0.021 | 0.041 | 0.019 | 0.039 | 0.044 |
| | 5 | 50M | 125bp | 0.545 | 0.736 | 0.562 | 0.667 | 0.459 | 0.021 | 0.040 | 0.020 | 0.040 | 0.044 |
| | 5 | 50M | 150bp | 0.560 | 0.748 | 0.576 | 0.682 | 0.458 | 0.021 | 0.040 | 0.021 | 0.041 | 0.042 |
| | 5 | 25/100M | 50bp | 0.413 | 0.660 | 0.436 | 0.539 | 0.392 | 0.016 | 0.038 | 0.015 | 0.033 | 0.042 |
| | 5 | 25/100M | 75bp | 0.437 | 0.670 | 0.449 | 0.553 | 0.396 | 0.016 | 0.036 | 0.014 | 0.031 | 0.039 |
| | 5 | 25/100M | 100bp | 0.445 | 0.680 | 0.460 | 0.567 | 0.397 | 0.016 | 0.038 | 0.015 | 0.032 | 0.040 |
| | 5 | 25/100M | 125bp | 0.476 | 0.699 | 0.488 | 0.597 | 0.408 | 0.018 | 0.040 | 0.018 | 0.036 | 0.042 |
| | 5 | 25/100M | 150bp | 0.492 | 0.707 | 0.498 | 0.608 | 0.395 | 0.017 | 0.039 | 0.016 | 0.034 | 0.038 |
| | 10 | 50M | 50bp | 0.838 | 0.854 | 0.794 | 0.802 | 0.861 | 0.030 | 0.041 | 0.026 | 0.035 | 0.049 |
| | 10 | 50M | 75bp | 0.855 | 0.868 | 0.812 | 0.820 | 0.869 | 0.030 | 0.042 | 0.026 | 0.035 | 0.049 |
| | 10 | 50M | 100bp | 0.864 | 0.877 | 0.824 | 0.832 | 0.875 | 0.031 | 0.043 | 0.026 | 0.036 | 0.047 |
| | 10 | 50M | 125bp | 0.878 | 0.890 | 0.846 | 0.854 | 0.887 | 0.031 | 0.042 | 0.026 | 0.035 | 0.047 |
| | 10 | 50M | 150bp | 0.887 | 0.898 | 0.858 | 0.867 | 0.895 | 0.032 | 0.041 | 0.027 | 0.036 | 0.047 |
| | 10 | 25/100M | 50bp | 0.803 | 0.861 | 0.729 | 0.737 | 0.845 | 0.025 | 0.036 | 0.019 | 0.027 | 0.046 |
| | 10 | 25/100M | 75bp | 0.818 | 0.874 | 0.749 | 0.757 | 0.852 | 0.026 | 0.038 | 0.021 | 0.028 | 0.046 |
| | 10 | 25/100M | 100bp | 0.830 | 0.883 | 0.763 | 0.771 | 0.861 | 0.026 | 0.037 | 0.020 | 0.028 | 0.047 |
| | 10 | 25/100M | 125bp | 0.852 | 0.896 | 0.794 | 0.801 | 0.873 | 0.027 | 0.039 | 0.022 | 0.029 | 0.046 |
| | 10 | 25/100M | 150bp | 0.862 | 0.904 | 0.808 | 0.816 | 0.879 | 0.027 | 0.038 | 0.021 | 0.029 | 0.047 |

Table S10: Simulation results - observed power and false discovery rate for different read types and read Lengths, averaged over 20 simulations. Scenario with mm39 genome, all transcripts expressed, and reads quantified with kallisto. Library size shown in million reads (M) with 25/100 indicating library sizes alternating between 25M and 100M across replicates. Read lengths are shown in base pairs (bp). Red color indicates observed FDR values greater than the nominal 0.05. Blue color indicates the most powerful method that controls the FDR at the nominal level for a given scenario (row). Empty cells indicate cases in which a method failed to call any transcript as DE.

| | | | | Power | | | | | False Discovery Rate | | | | |
|---|---|---|---|---|---|---|---|---|---|---|---|---|---|
| Read | Samples/Group | Library Size | Read Length | edgeR-raw | edgeR-scaled | sleuth-LRT | sleuth-Wald | Swish | edgeR-raw | edgeR-scaled | sleuth-LRT | sleuth-Wald | Swish |
| paired-end | 3 | 50M | 50bp | 0.001 | 0.434 | 0.000 | 0.418 | 0.474 | 0.694 | 0.041 | - | 0.056 | 0.248 |
| | 3 | 50M | 75bp | 0.001 | 0.440 | 0.000 | 0.425 | 0.484 | 0.619 | 0.039 | - | 0.055 | 0.250 |
| | 3 | 50M | 100bp | 0.000 | 0.445 | 0.000 | 0.425 | 0.482 | 0.703 | 0.042 | - | 0.053 | 0.250 |
| | 3 | 50M | 125bp | 0.000 | 0.445 | 0.000 | 0.429 | 0.490 | 0.841 | 0.040 | - | 0.051 | 0.249 |
| | 3 | 50M | 150bp | 0.001 | 0.451 | 0.000 | 0.433 | 0.491 | 0.689 | 0.041 | - | 0.056 | 0.251 |
| | 3 | 25/100M | 50bp | 0.000 | 0.376 | 0.000 | 0.347 | 0.404 | 0.650 | 0.035 | - | 0.047 | 0.230 |
| | 3 | 25/100M | 75bp | 0.000 | 0.390 | 0.000 | 0.364 | 0.415 | 0.727 | 0.035 | - | 0.045 | 0.230 |
| | 3 | 25/100M | 100bp | 0.000 | 0.387 | 0.000 | 0.356 | 0.412 | 0.769 | 0.035 | - | 0.045 | 0.234 |
| | 3 | 25/100M | 125bp | 0.000 | 0.399 | 0.000 | 0.369 | 0.422 | 0.771 | 0.035 | - | 0.049 | 0.234 |
| | 3 | 25/100M | 150bp | 0.000 | 0.395 | 0.000 | 0.367 | 0.420 | 0.833 | 0.035 | - | 0.047 | 0.233 |
| | 5 | 50M | 50bp | 0.592 | 0.754 | 0.579 | 0.692 | 0.465 | 0.026 | 0.042 | 0.021 | 0.042 | 0.046 |
| | 5 | 50M | 75bp | 0.610 | 0.766 | 0.593 | 0.703 | 0.473 | 0.025 | 0.043 | 0.020 | 0.039 | 0.043 |
| | 5 | 50M | 100bp | 0.609 | 0.769 | 0.592 | 0.707 | 0.470 | 0.024 | 0.044 | 0.019 | 0.041 | 0.042 |
| | 5 | 50M | 125bp | 0.616 | 0.771 | 0.598 | 0.709 | 0.467 | 0.026 | 0.041 | 0.020 | 0.040 | 0.044 |
| | 5 | 50M | 150bp | 0.621 | 0.773 | 0.601 | 0.714 | 0.465 | 0.025 | 0.042 | 0.019 | 0.039 | 0.044 |
| | 5 | 25/100M | 50bp | 0.527 | 0.708 | 0.507 | 0.620 | 0.434 | 0.022 | 0.040 | 0.017 | 0.036 | 0.047 |
| | 5 | 25/100M | 75bp | 0.547 | 0.725 | 0.523 | 0.637 | 0.433 | 0.021 | 0.039 | 0.016 | 0.037 | 0.044 |
| | 5 | 25/100M | 100bp | 0.550 | 0.730 | 0.524 | 0.641 | 0.430 | 0.022 | 0.038 | 0.016 | 0.035 | 0.043 |
| | 5 | 25/100M | 125bp | 0.552 | 0.734 | 0.527 | 0.647 | 0.434 | 0.020 | 0.040 | 0.016 | 0.035 | 0.044 |
| | 5 | 25/100M | 150bp | 0.554 | 0.733 | 0.529 | 0.647 | 0.407 | 0.020 | 0.037 | 0.015 | 0.035 | 0.041 |
| | 10 | 50M | 50bp | 0.908 | 0.908 | 0.877 | 0.885 | 0.897 | 0.034 | 0.045 | 0.028 | 0.037 | 0.049 |
| | 10 | 50M | 75bp | 0.917 | 0.921 | 0.890 | 0.898 | 0.908 | 0.033 | 0.044 | 0.028 | 0.037 | 0.050 |
| | 10 | 50M | 100bp | 0.921 | 0.925 | 0.894 | 0.901 | 0.909 | 0.034 | 0.045 | 0.028 | 0.036 | 0.049 |
| | 10 | 50M | 125bp | 0.922 | 0.927 | 0.896 | 0.903 | 0.913 | 0.033 | 0.044 | 0.027 | 0.035 | 0.047 |
| | 10 | 50M | 150bp | 0.922 | 0.928 | 0.898 | 0.905 | 0.912 | 0.033 | 0.044 | 0.027 | 0.036 | 0.048 |
| | 10 | 25/100M | 50bp | 0.885 | 0.915 | 0.831 | 0.839 | 0.887 | 0.031 | 0.040 | 0.024 | 0.032 | 0.049 |
| | 10 | 25/100M | 75bp | 0.894 | 0.924 | 0.842 | 0.850 | 0.892 | 0.029 | 0.040 | 0.022 | 0.031 | 0.047 |
| | 10 | 25/100M | 100bp | 0.898 | 0.927 | 0.850 | 0.858 | 0.895 | 0.031 | 0.042 | 0.025 | 0.033 | 0.048 |
| | 10 | 25/100M | 125bp | 0.902 | 0.931 | 0.857 | 0.866 | 0.900 | 0.029 | 0.040 | 0.023 | 0.030 | 0.045 |
| | 10 | 25/100M | 150bp | 0.901 | 0.929 | 0.853 | 0.862 | 0.898 | 0.030 | 0.040 | 0.022 | 0.030 | 0.047 |

Table S10: Simulation results - observed power and false discovery rate for different read types and read Lengths, averaged over 20 simulations. Scenario with mm39 genome, all transcripts expressed, and reads quantified with kallisto. Library size shown in million reads (M) with 25/100 indicating library sizes alternating between 25M and 100M across replicates. Read lengths are shown in base pairs (bp). Red color indicates observed FDR values greater than the nominal 0.05. Blue color indicates the most powerful method that controls the FDR at the nominal level for a given scenario (row). Empty cells indicate cases in which a method failed to call any transcript as DE. *(continued)*

| | | | | Power | | | | | False Discovery Rate | | | | |
|---|---|---|---|---|---|---|---|---|---|---|---|---|---|
| Read | Samples/Group | Library Size | Read Length | edgeR-raw | edgeR-scaled | sleuth-LRT | sleuth-Wald | Swish | edgeR-raw | edgeR-scaled | sleuth-LRT | sleuth-Wald | Swish |
| single-end | 3 | 50M | 50bp | 0.000 | 0.401 | 0.000 | 0.378 | 0.436 | 0.850 | 0.049 | - | 0.057 | 0.242 |
| | 3 | 50M | 75bp | 0.000 | 0.413 | 0.000 | 0.389 | 0.444 | 0.860 | 0.048 | - | 0.058 | 0.242 |
| | 3 | 50M | 100bp | 0.000 | 0.416 | 0.000 | 0.394 | 0.451 | 0.803 | 0.046 | - | 0.057 | 0.249 |
| | 3 | 50M | 125bp | 0.000 | 0.426 | 0.000 | 0.413 | 0.467 | 0.856 | 0.045 | - | 0.058 | 0.249 |
| | 3 | 50M | 150bp | 0.001 | 0.435 | 0.000 | 0.420 | 0.473 | 0.667 | 0.045 | - | 0.056 | 0.246 |
| | 3 | 25/100M | 50bp | 0.000 | 0.351 | 0.000 | 0.312 | 0.368 | 0.750 | 0.040 | - | 0.047 | 0.231 |
| | 3 | 25/100M | 75bp | 0.000 | 0.356 | 0.000 | 0.316 | 0.372 | 0.882 | 0.038 | - | 0.045 | 0.225 |
| | 3 | 25/100M | 100bp | 0.000 | 0.365 | 0.000 | 0.328 | 0.382 | 0.824 | 0.038 | - | 0.046 | 0.229 |
| | 3 | 25/100M | 125bp | 0.000 | 0.377 | 0.000 | 0.339 | 0.392 | 0.769 | 0.040 | - | 0.048 | 0.232 |
| | 3 | 25/100M | 150bp | 0.000 | 0.380 | 0.000 | 0.347 | 0.401 | 0.588 | 0.038 | - | 0.049 | 0.234 |
| | 5 | 50M | 50bp | 0.506 | 0.715 | 0.526 | 0.639 | 0.445 | 0.031 | 0.052 | 0.026 | 0.048 | 0.053 |
| | 5 | 50M | 75bp | 0.528 | 0.728 | 0.539 | 0.653 | 0.452 | 0.033 | 0.050 | 0.026 | 0.048 | 0.052 |
| | 5 | 50M | 100bp | 0.545 | 0.732 | 0.548 | 0.662 | 0.450 | 0.031 | 0.050 | 0.025 | 0.046 | 0.050 |
| | 5 | 50M | 125bp | 0.569 | 0.743 | 0.566 | 0.678 | 0.471 | 0.032 | 0.050 | 0.025 | 0.046 | 0.053 |
| | 5 | 50M | 150bp | 0.586 | 0.754 | 0.579 | 0.691 | 0.465 | 0.031 | 0.049 | 0.026 | 0.047 | 0.049 |
| | 5 | 25/100M | 50bp | 0.436 | 0.669 | 0.447 | 0.560 | 0.400 | 0.025 | 0.047 | 0.019 | 0.039 | 0.049 |
| | 5 | 25/100M | 75bp | 0.462 | 0.679 | 0.461 | 0.575 | 0.401 | 0.025 | 0.045 | 0.019 | 0.038 | 0.046 |
| | 5 | 25/100M | 100bp | 0.477 | 0.691 | 0.472 | 0.590 | 0.416 | 0.025 | 0.044 | 0.020 | 0.039 | 0.047 |
| | 5 | 25/100M | 125bp | 0.505 | 0.705 | 0.496 | 0.612 | 0.422 | 0.026 | 0.046 | 0.021 | 0.041 | 0.048 |
| | 5 | 25/100M | 150bp | 0.522 | 0.712 | 0.503 | 0.621 | 0.428 | 0.026 | 0.045 | 0.020 | 0.041 | 0.049 |
| | 10 | 50M | 50bp | 0.863 | 0.875 | 0.825 | 0.833 | 0.864 | 0.052 | 0.062 | 0.042 | 0.050 | 0.071 |
| | 10 | 50M | 75bp | 0.875 | 0.886 | 0.839 | 0.847 | 0.871 | 0.052 | 0.061 | 0.042 | 0.051 | 0.071 |
| | 10 | 50M | 100bp | 0.886 | 0.894 | 0.851 | 0.860 | 0.879 | 0.052 | 0.061 | 0.041 | 0.051 | 0.068 |
| | 10 | 50M | 125bp | 0.897 | 0.904 | 0.866 | 0.874 | 0.888 | 0.051 | 0.060 | 0.039 | 0.049 | 0.068 |
| | 10 | 50M | 150bp | 0.905 | 0.911 | 0.875 | 0.883 | 0.895 | 0.049 | 0.057 | 0.039 | 0.049 | 0.065 |
| | 10 | 25/100M | 50bp | 0.833 | 0.880 | 0.768 | 0.775 | 0.848 | 0.047 | 0.060 | 0.035 | 0.043 | 0.068 |
| | 10 | 25/100M | 75bp | 0.846 | 0.891 | 0.782 | 0.791 | 0.857 | 0.047 | 0.059 | 0.033 | 0.042 | 0.067 |
| | 10 | 25/100M | 100bp | 0.859 | 0.902 | 0.798 | 0.806 | 0.866 | 0.046 | 0.057 | 0.033 | 0.041 | 0.066 |
| | 10 | 25/100M | 125bp | 0.874 | 0.909 | 0.820 | 0.828 | 0.876 | 0.045 | 0.056 | 0.033 | 0.041 | 0.064 |
| | 10 | 25/100M | 150bp | 0.883 | 0.915 | 0.830 | 0.838 | 0.882 | 0.043 | 0.053 | 0.031 | 0.039 | 0.063 |

##### 1.2.2 Type 1 error rate control

In this subsection, we present results from our null simulations comparing methods with respect to the control of the type 1 error rate. Simulations considered in this subsection were absent of any truly differential transcript expression between groups.

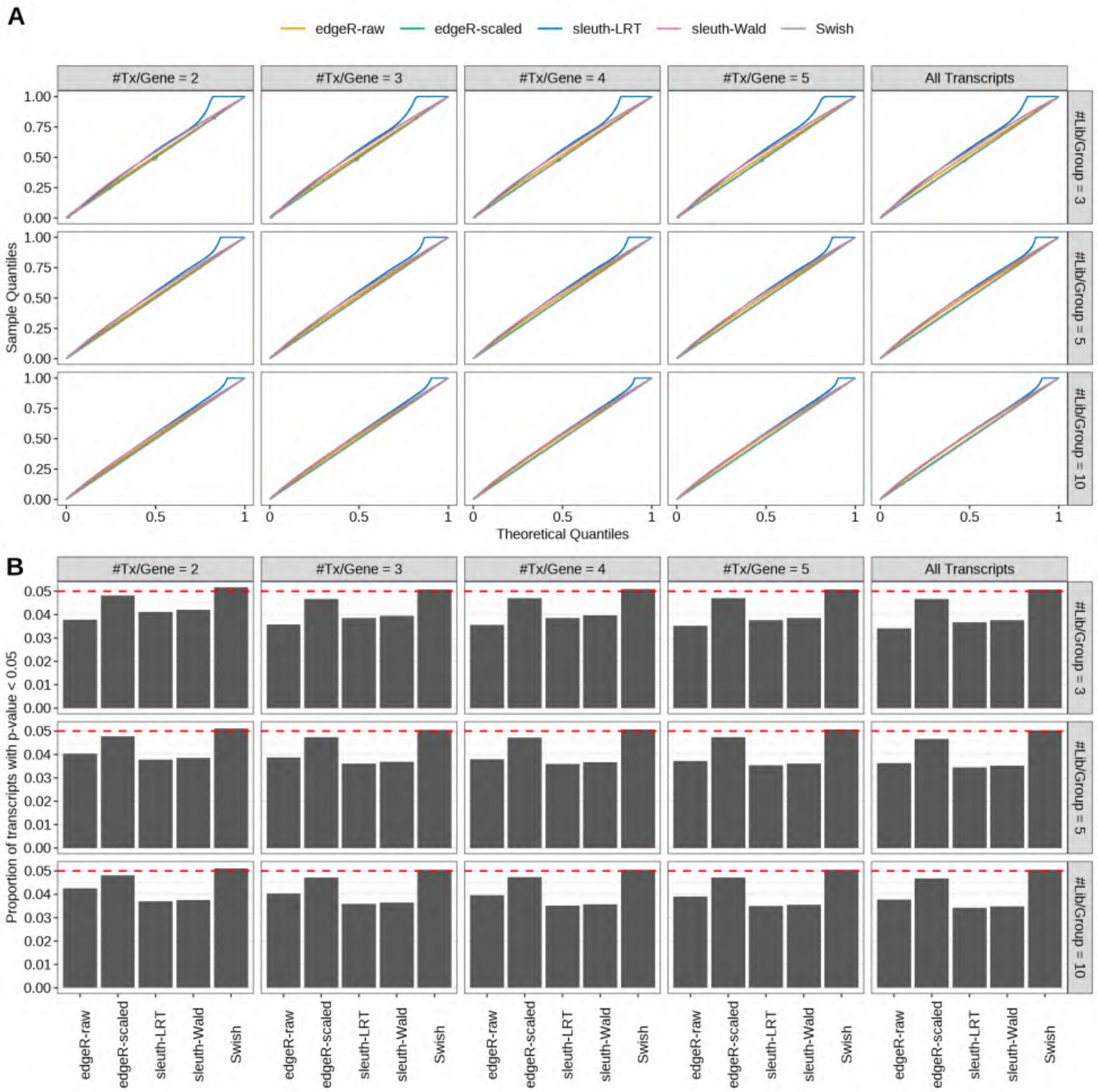

Figure S41: Simulation results. Scenario with mm39 genome, 50bp single-end reads quantified with Salmon, and balanced libraries. (A) QQ plots of p-values for simulations without any differential expression (averaged over 20 simulations). (B) Proportion of transcripts with unadjusted p-values less than 0.05 for simulations without any differential expression (averaged over 20 simulations)

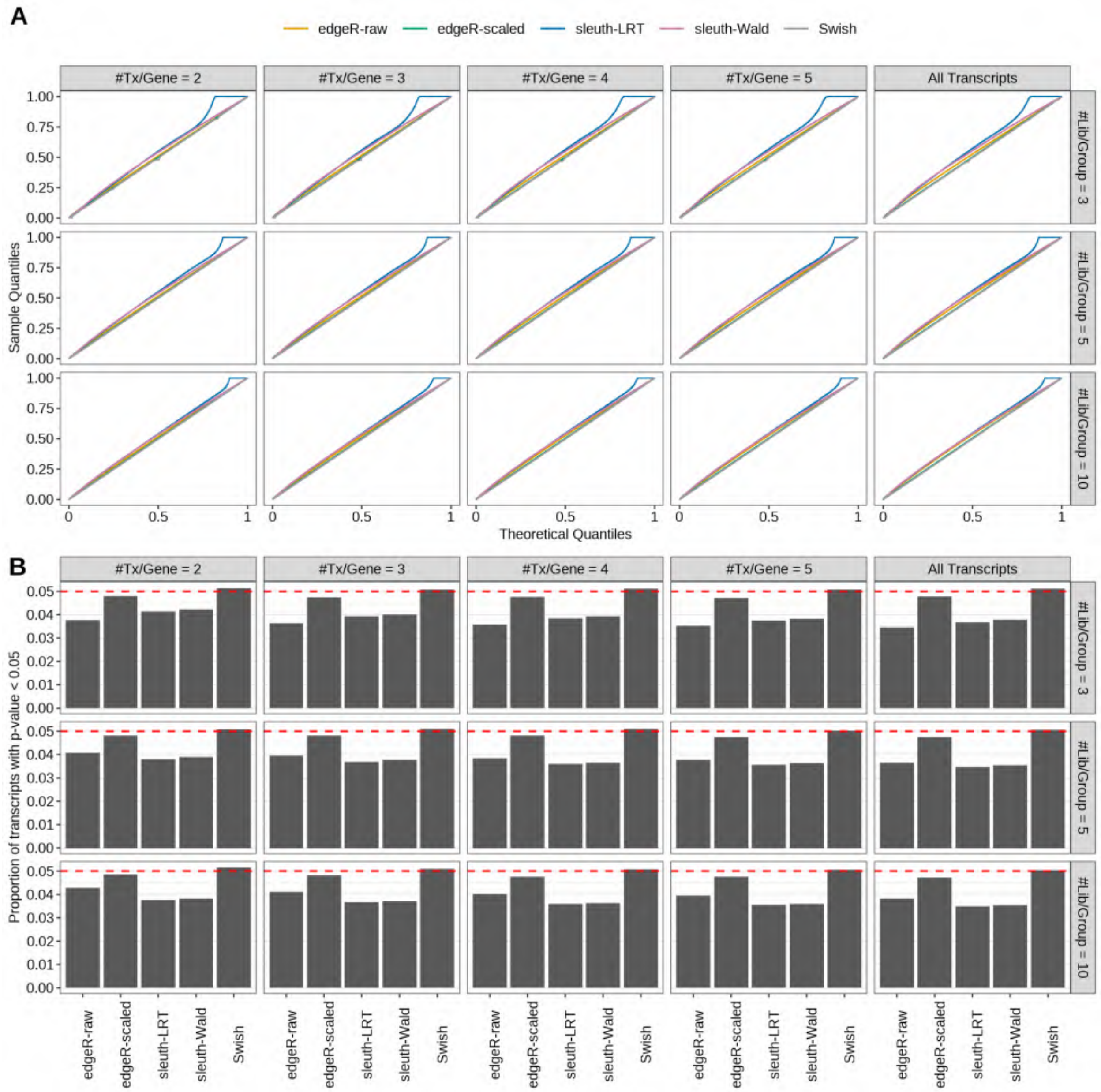

Figure S42: Simulation results. Scenario with mm39 genome, 75bp single-end reads quantified with Salmon, and balanced libraries. (A) QQ plots of p-values for simulations without any differential expression (averaged over 20 simulations). (B) Proportion of transcripts with unadjusted p-values less than 0.05 for simulations without any differential expression (averaged over 20 simulations)

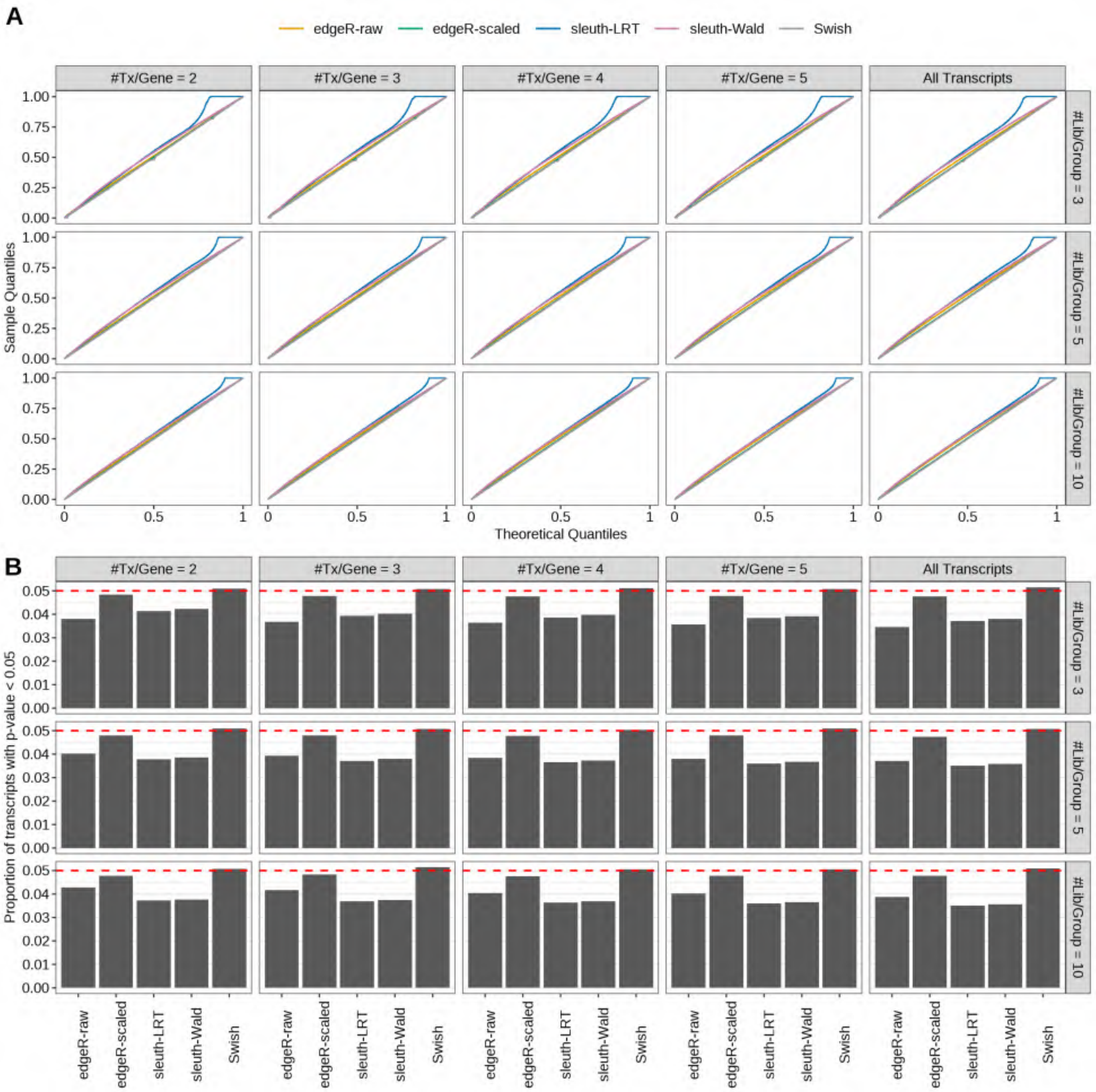

Figure S43: Simulation results. Scenario with mm39 genome, 100bp single-end reads quantified with Salmon, and balanced libraries. (A) QQ plots of p-values for simulations without any differential expression (averaged over 20 simulations). (B) Proportion of transcripts with unadjusted p-values less than 0.05 for simulations without any differential expression (averaged over 20 simulations)

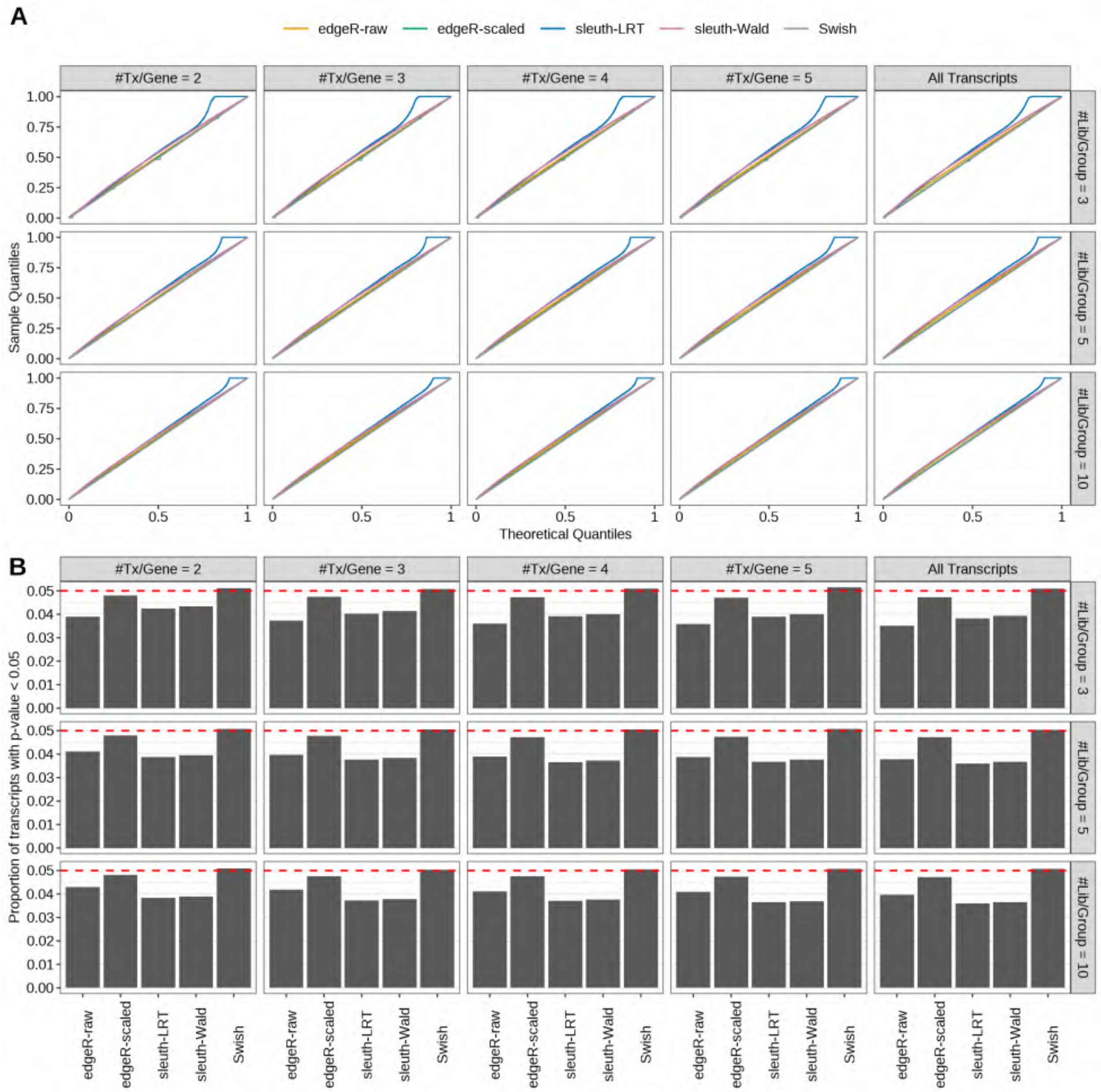

Figure S44: Simulation results. Scenario with mm39 genome, 125bp single-end reads quantified with Salmon, and balanced libraries. (A) QQ plots of p-values for simulations without any differential expression (averaged over 20 simulations). (B) Proportion of transcripts with unadjusted p-values less than 0.05 for simulations without any differential expression (averaged over 20 simulations)

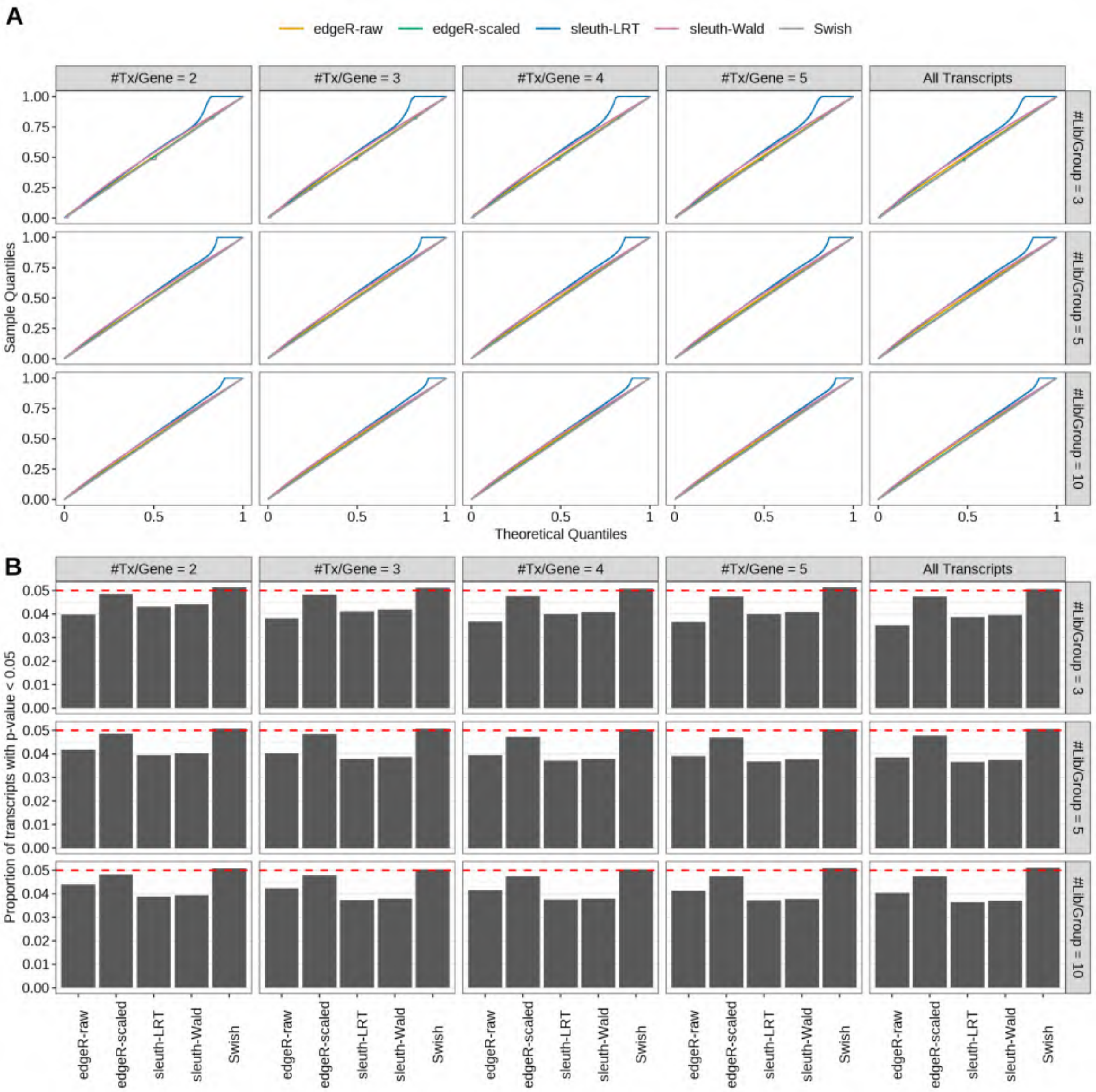

Figure S45: Simulation results. Scenario with mm39 genome, 150bp single-end reads quantified with Salmon, and balanced libraries. (A) QQ plots of p-values for simulations without any differential expression (averaged over 20 simulations). (B) Proportion of transcripts with unadjusted p-values less than 0.05 for simulations without any differential expression (averaged over 20 simulations)

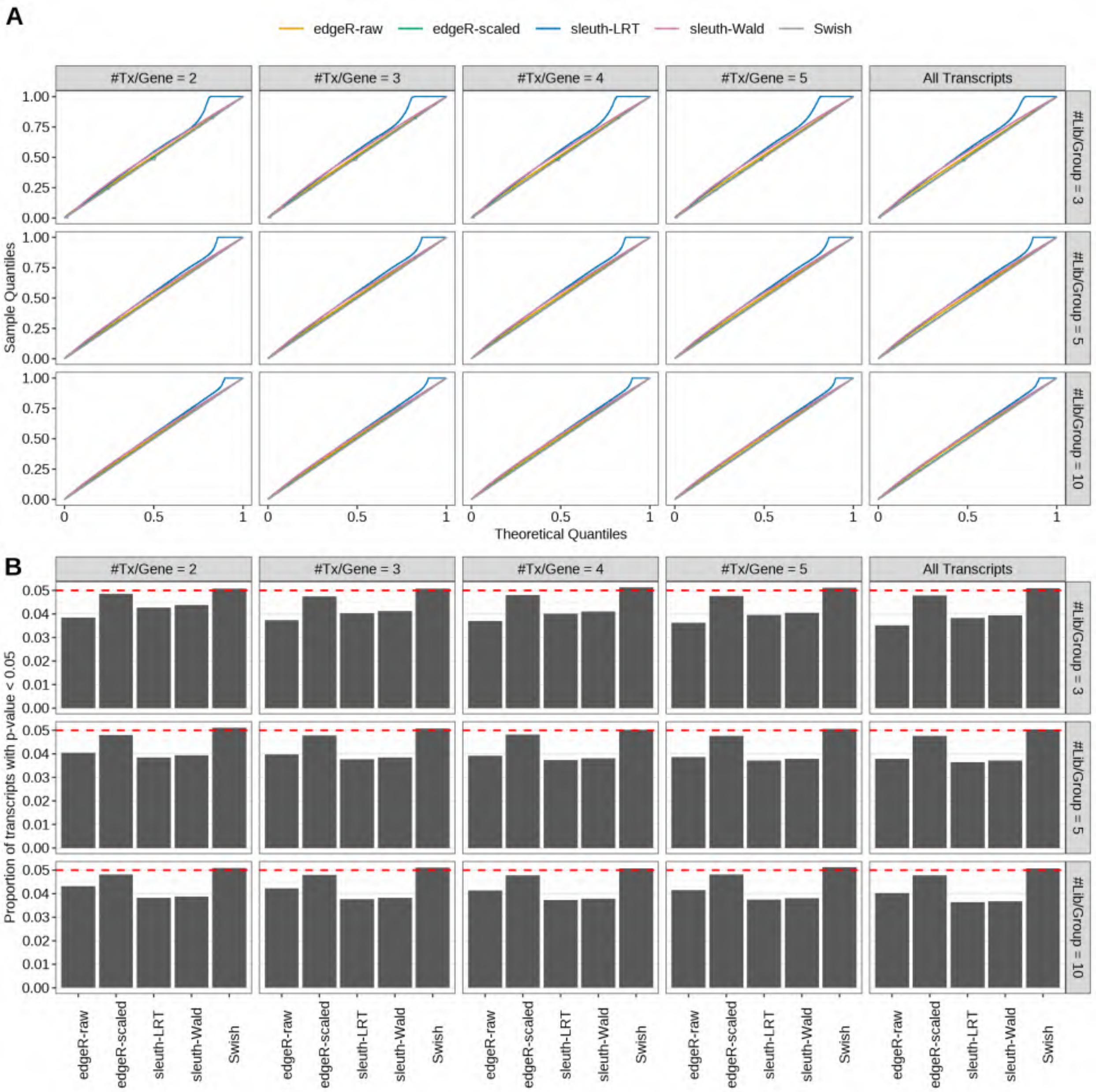

Figure S46: Simulation results. Scenario with mm39 genome, 50bp paired-end reads quantified with Salmon, and balanced libraries. (A) QQ plots of p-values for simulations without any differential expression (averaged over 20 simulations). (B) Proportion of transcripts with unadjusted p-values less than 0.05 for simulations without any differential expression (averaged over 20 simulations)

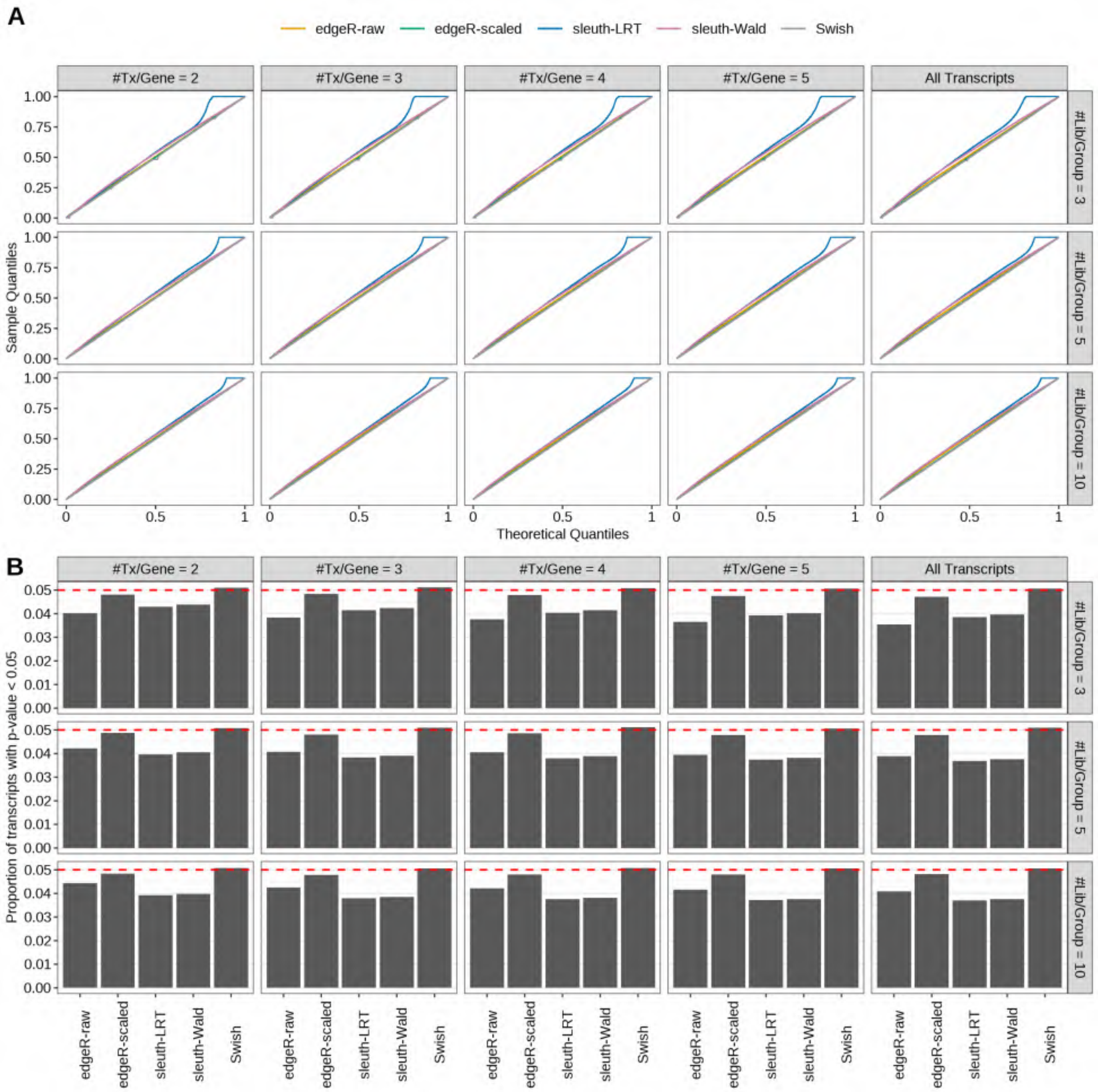

Figure S47: Simulation results. Scenario with mm39 genome, 75bp paired-end reads quantified with Salmon, and balanced libraries. (A) QQ plots of p-values for simulations without any differential expression (averaged over 20 simulations). (B) Proportion of transcripts with unadjusted p-values less than 0.05 for simulations without any differential expression (averaged over 20 simulations)

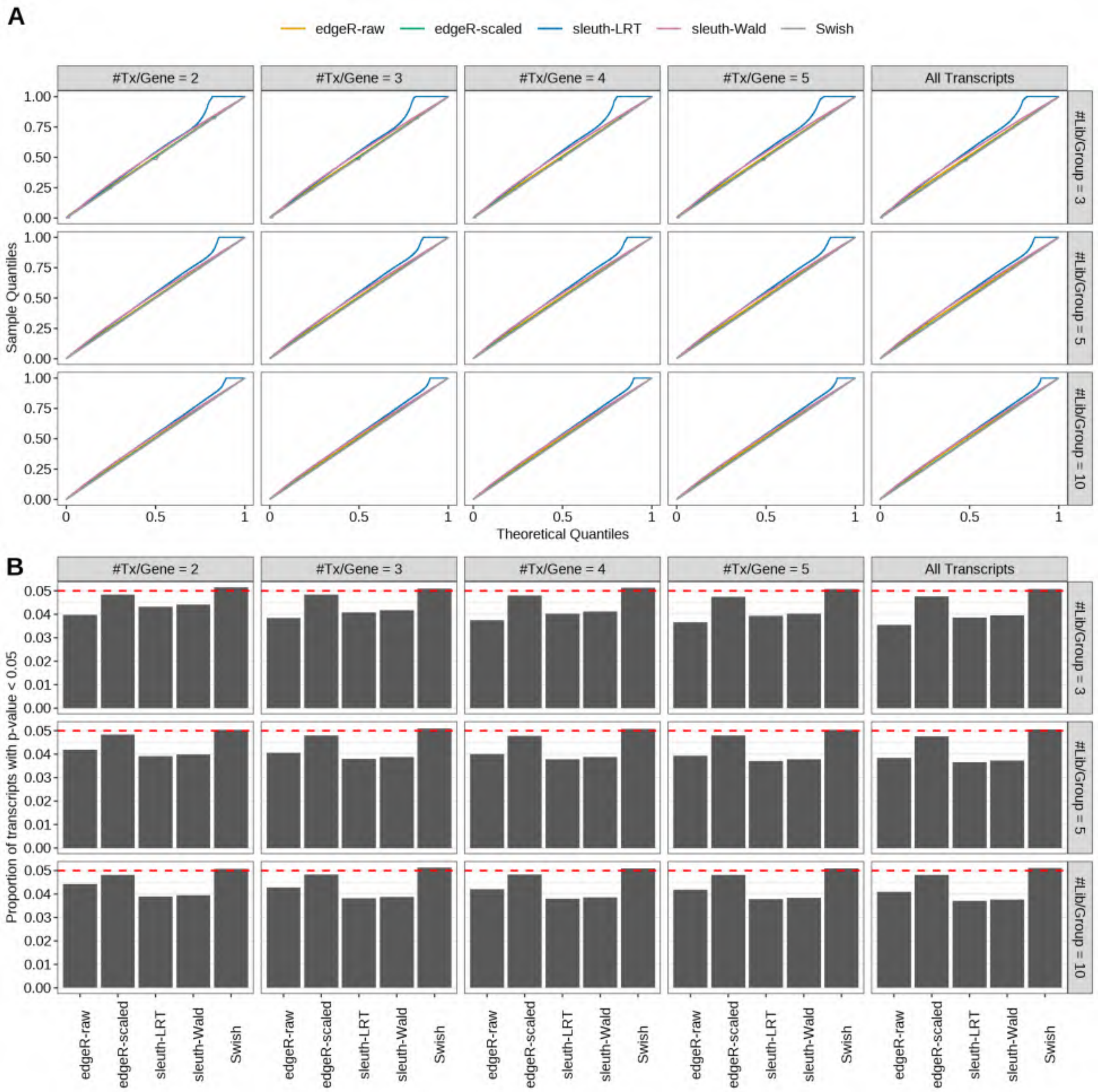

Figure S48: Simulation results. Scenario with mm39 genome, 100bp paired-end reads quantified with Salmon, and balanced libraries. (A) QQ plots of p-values for simulations without any differential expression (averaged over 20 simulations). (B) Proportion of transcripts with unadjusted p-values less than 0.05 for simulations without any differential expression (averaged over 20 simulations)

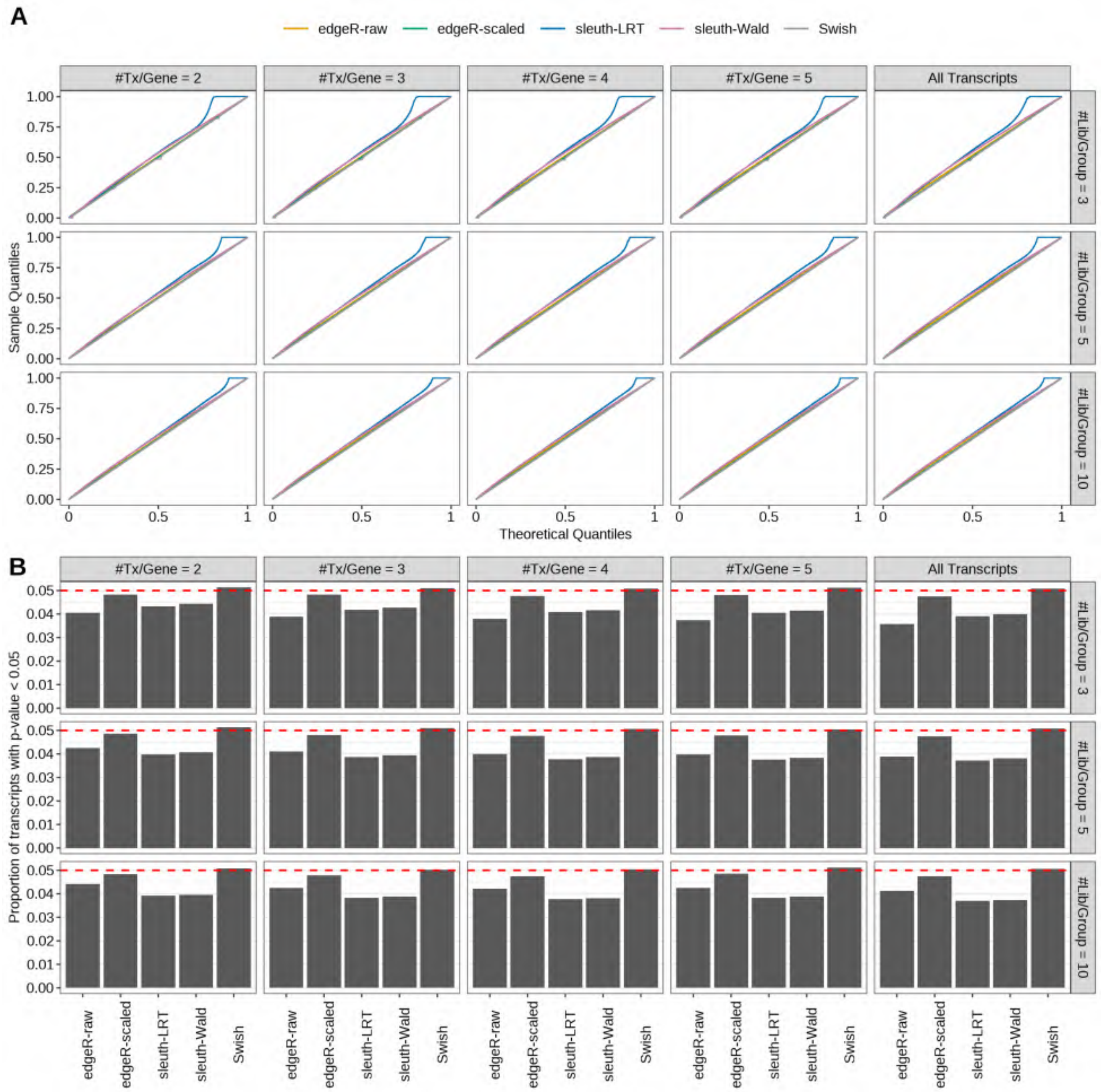

Figure S49: Simulation results. Scenario with mm39 genome, 125bp paired-end reads quantified with Salmon, and balanced libraries. (A) QQ plots of p-values for simulations without any differential expression (averaged over 20 simulations). (B) Proportion of transcripts with unadjusted p-values less than 0.05 for simulations without any differential expression (averaged over 20 simulations)

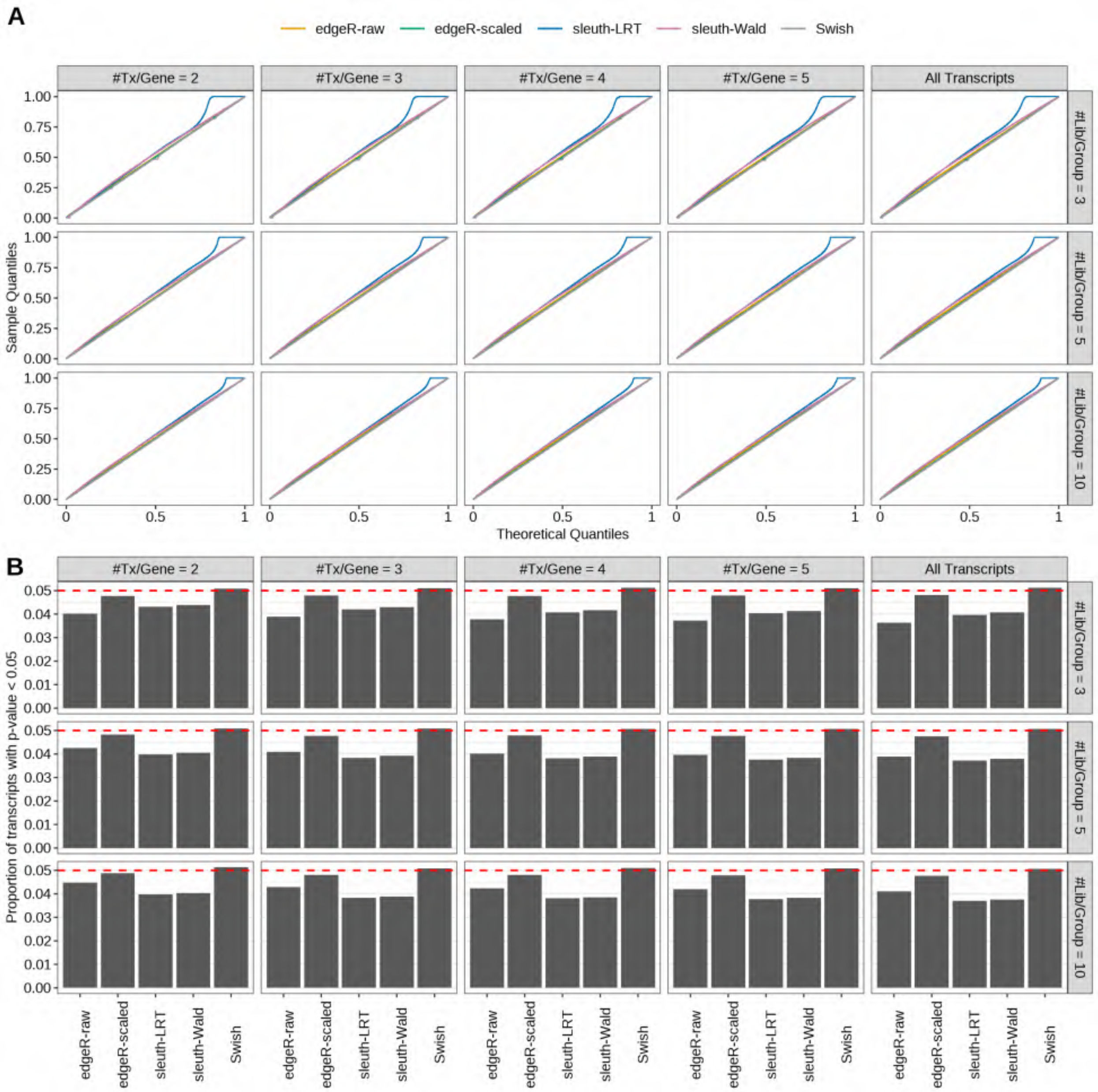

Figure S50: Simulation results. Scenario with mm39 genome, 150bp paired-end reads quantified with Salmon, and balanced libraries. (A) QQ plots of p-values for simulations without any differential expression (averaged over 20 simulations). (B) Proportion of transcripts with unadjusted p-values less than 0.05 for simulations without any differential expression (averaged over 20 simulations)

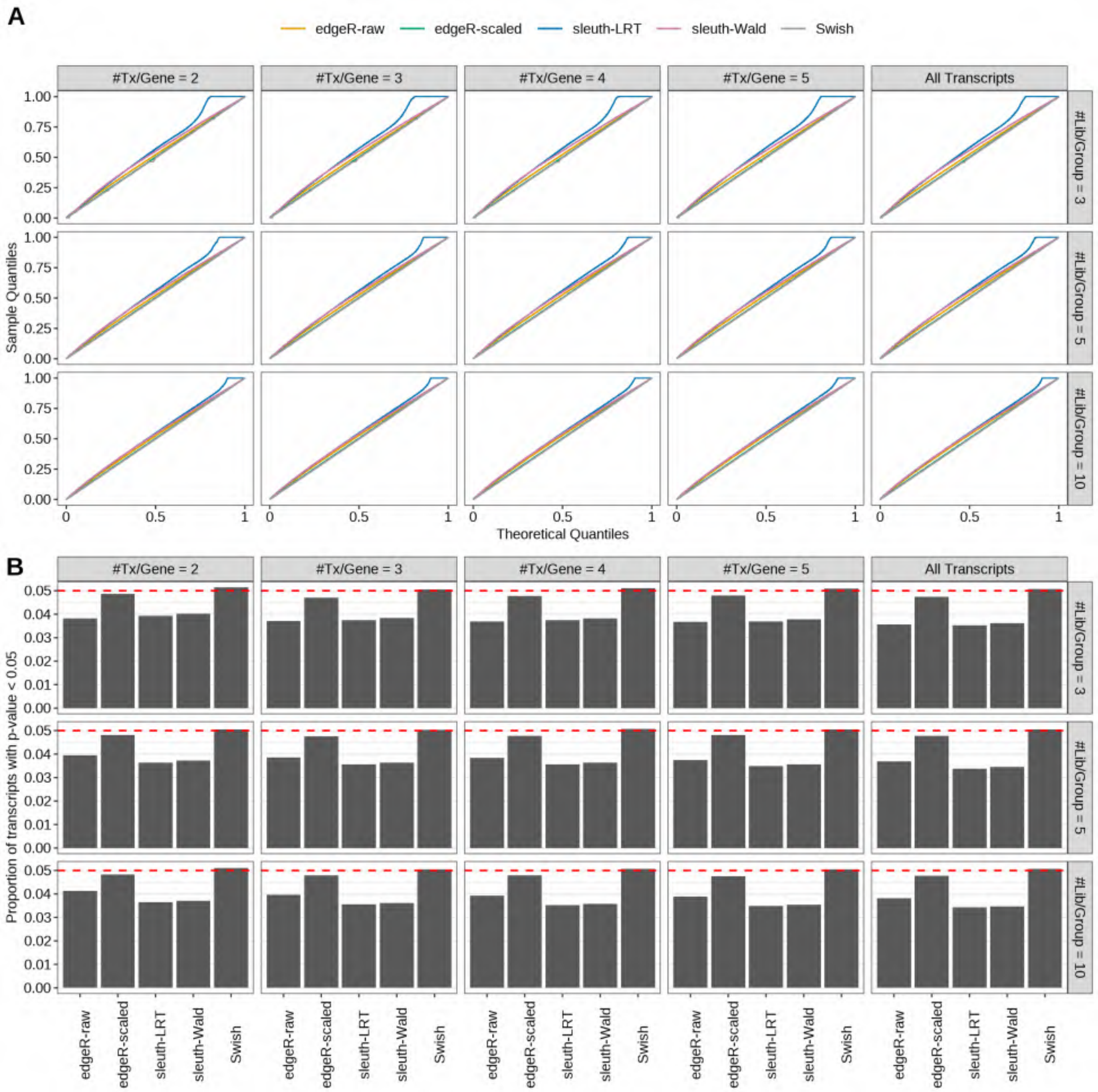

Figure S51: Simulation results. Scenario with mm39 genome, 50bp single-end reads quantified with kallisto, and balanced libraries. (A) QQ plots of p-values for simulations without any differential expression (averaged over 20 simulations). (B) Proportion of transcripts with unadjusted p-values less than 0.05 for simulations without any differential expression (averaged over 20 simulations)

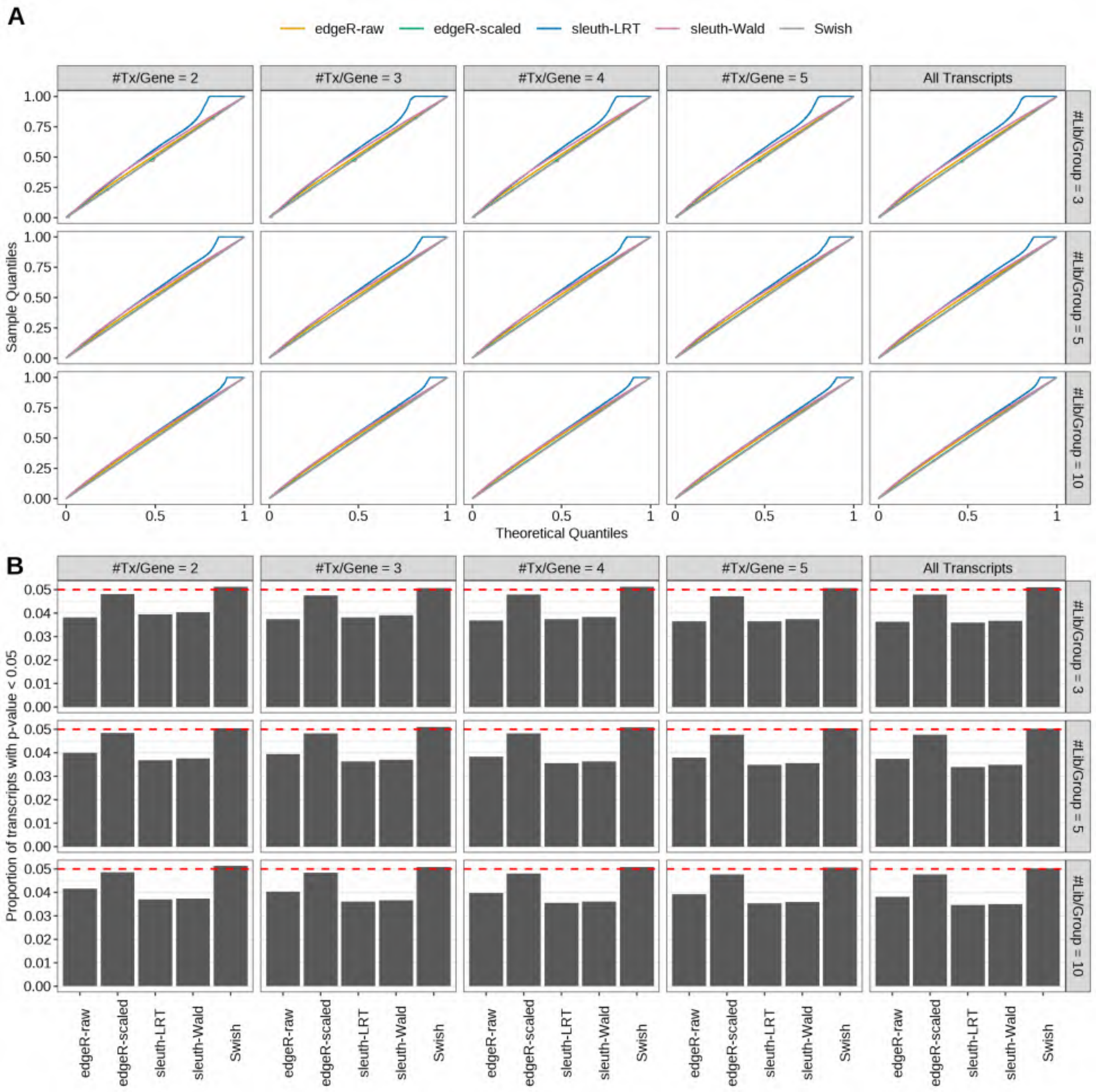

Figure S52: Simulation results. Scenario with mm39 genome, 75bp single-end reads quantified with kallisto, and balanced libraries. (A) QQ plots of p-values for simulations without any differential expression (averaged over 20 simulations). (B) Proportion of transcripts with unadjusted p-values less than 0.05 for simulations without any differential expression (averaged over 20 simulations)

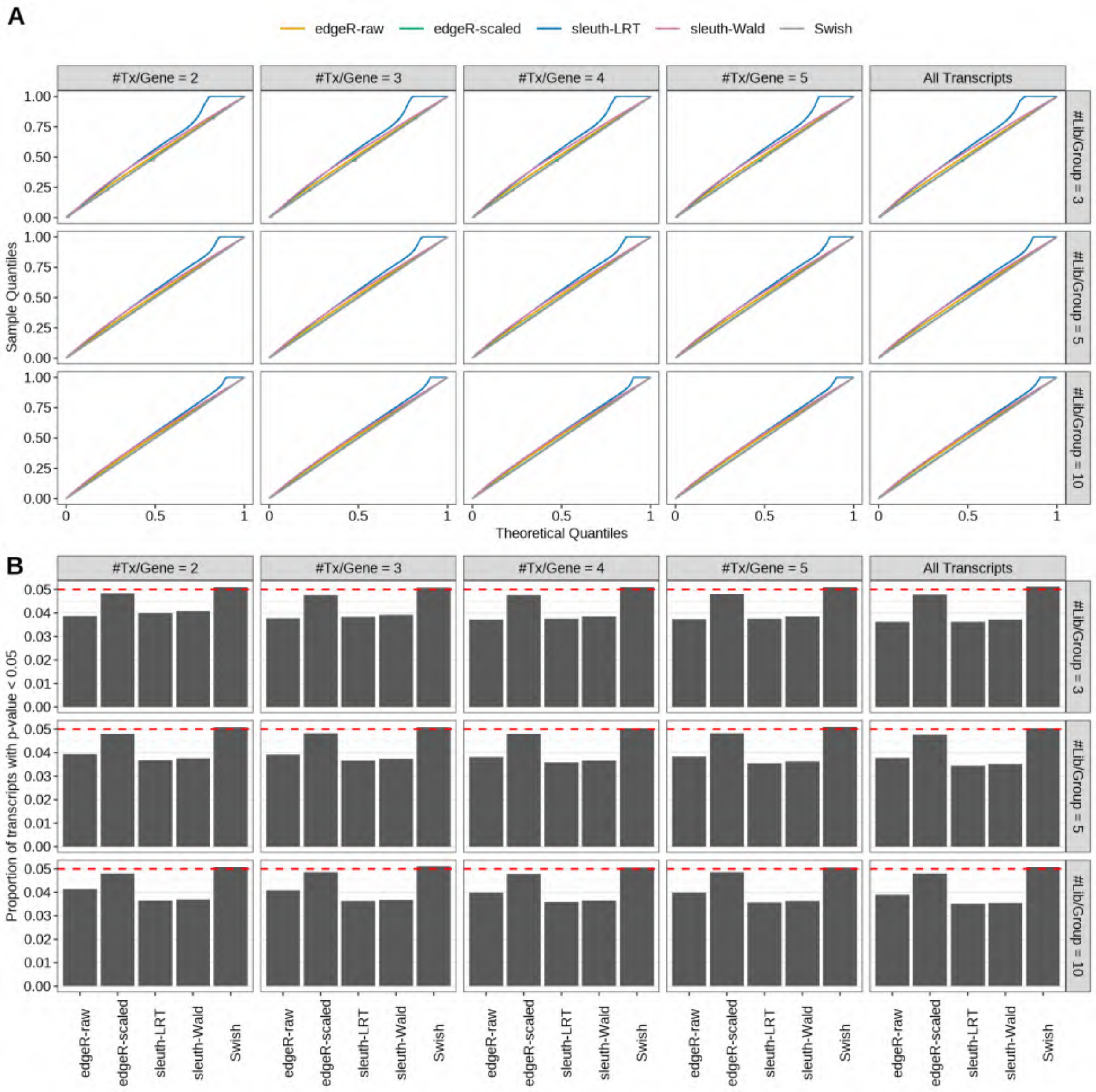

Figure S53: Simulation results. Scenario with mm39 genome, 100bp single-end reads quantified with kallisto, and balanced libraries. (A) QQ plots of p-values for simulations without any differential expression (averaged over 20 simulations). (B) Proportion of transcripts with unadjusted p-values less than 0.05 for simulations without any differential expression (averaged over 20 simulations)

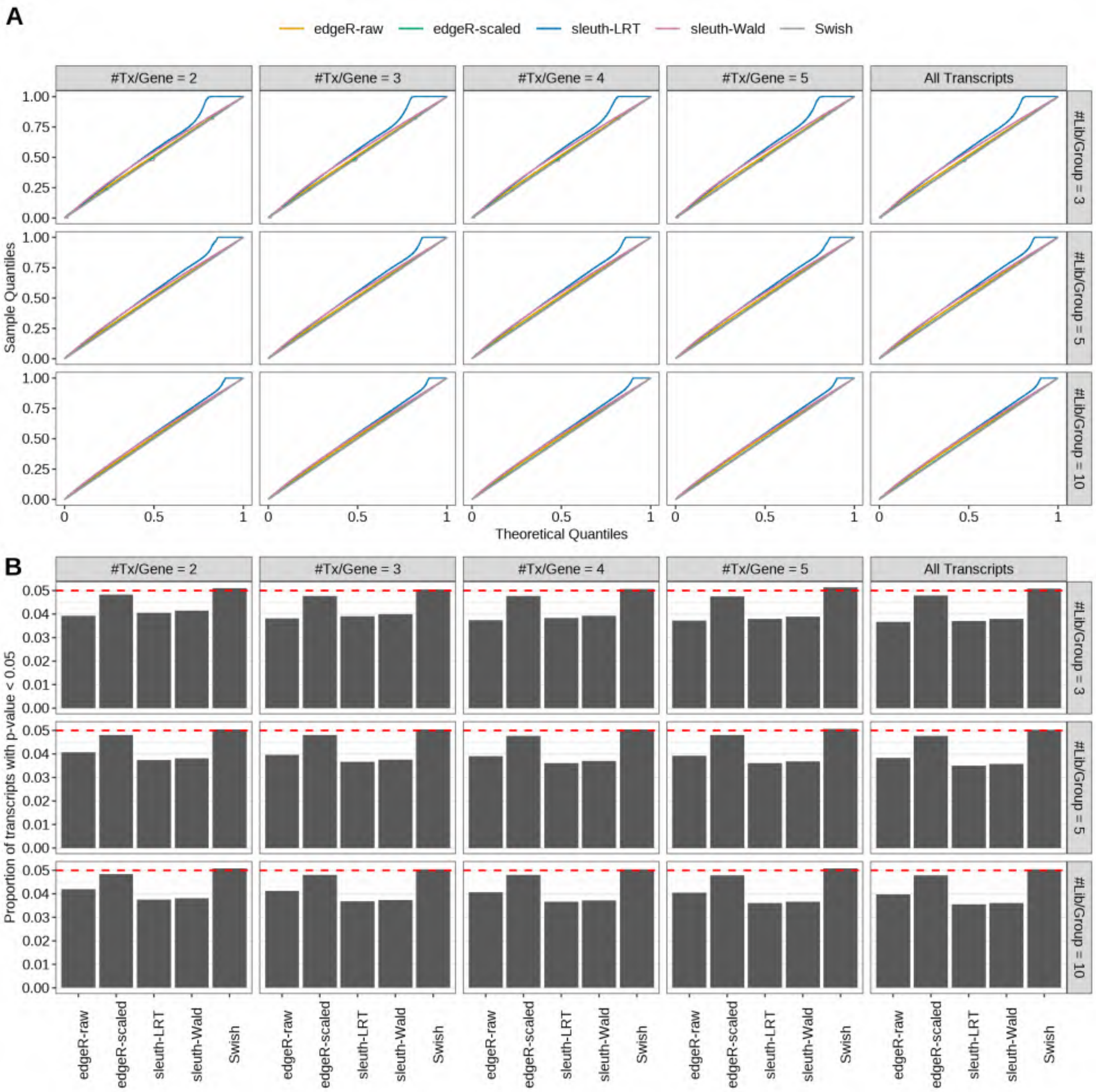

Figure S54: Simulation results. Scenario with mm39 genome, 125bp single-end reads quantified with kallisto, and balanced libraries. (A) QQ plots of p-values for simulations without any differential expression (averaged over 20 simulations). (B) Proportion of transcripts with unadjusted p-values less than 0.05 for simulations without any differential expression (averaged over 20 simulations)

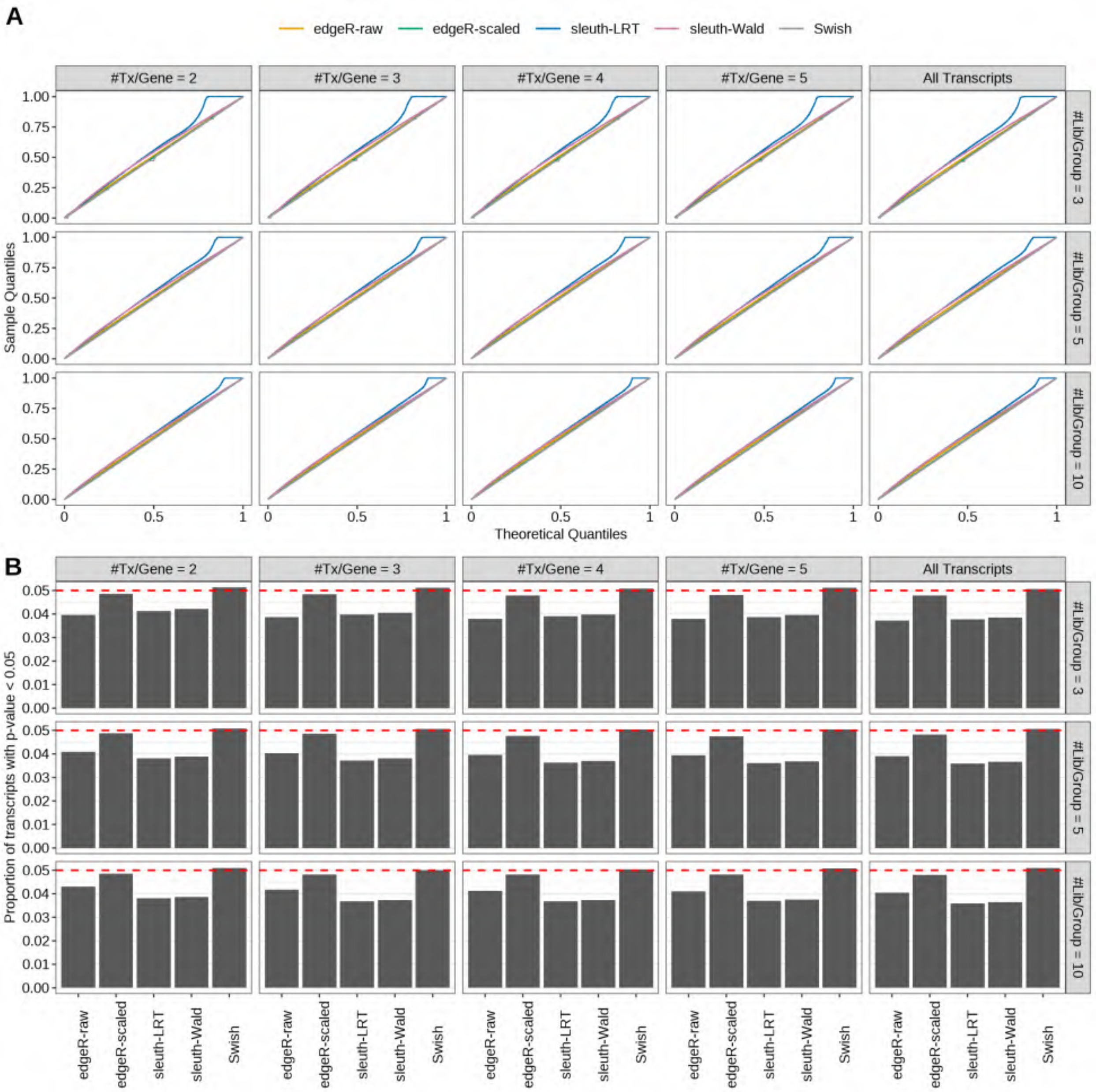

Figure S55: Simulation results. Scenario with mm39 genome, 150bp single-end reads quantified with kallisto, and balanced libraries. (A) QQ plots of p-values for simulations without any differential expression (averaged over 20 simulations). (B) Proportion of transcripts with unadjusted p-values less than 0.05 for simulations without any differential expression (averaged over 20 simulations)

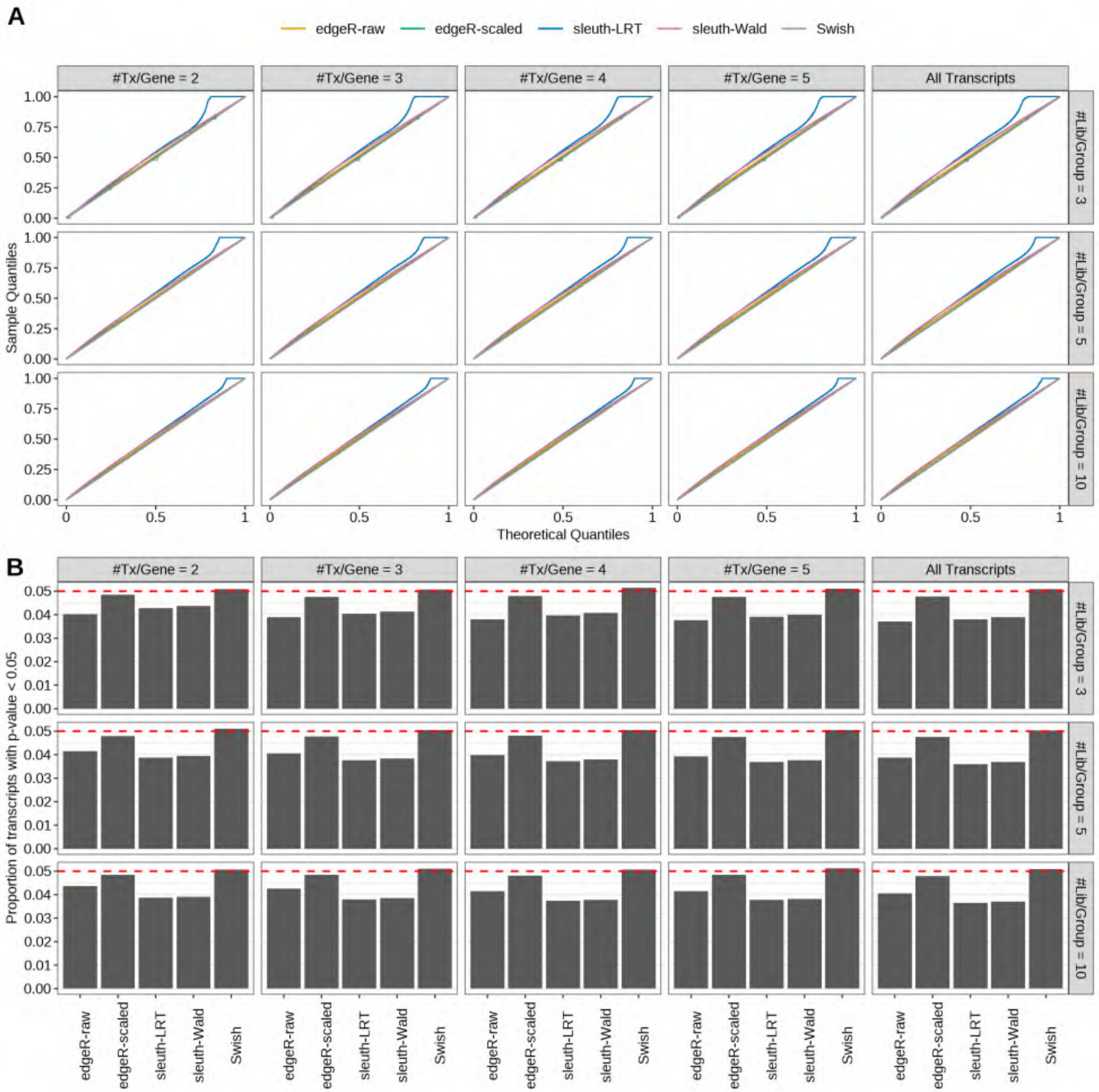

Figure S56: Simulation results. Scenario with mm39 genome, 50bp paired-end reads quantified with kallisto, and balanced libraries. (A) QQ plots of p-values for simulations without any differential expression (averaged over 20 simulations). (B) Proportion of transcripts with unadjusted p-values less than 0.05 for simulations without any differential expression (averaged over 20 simulations)

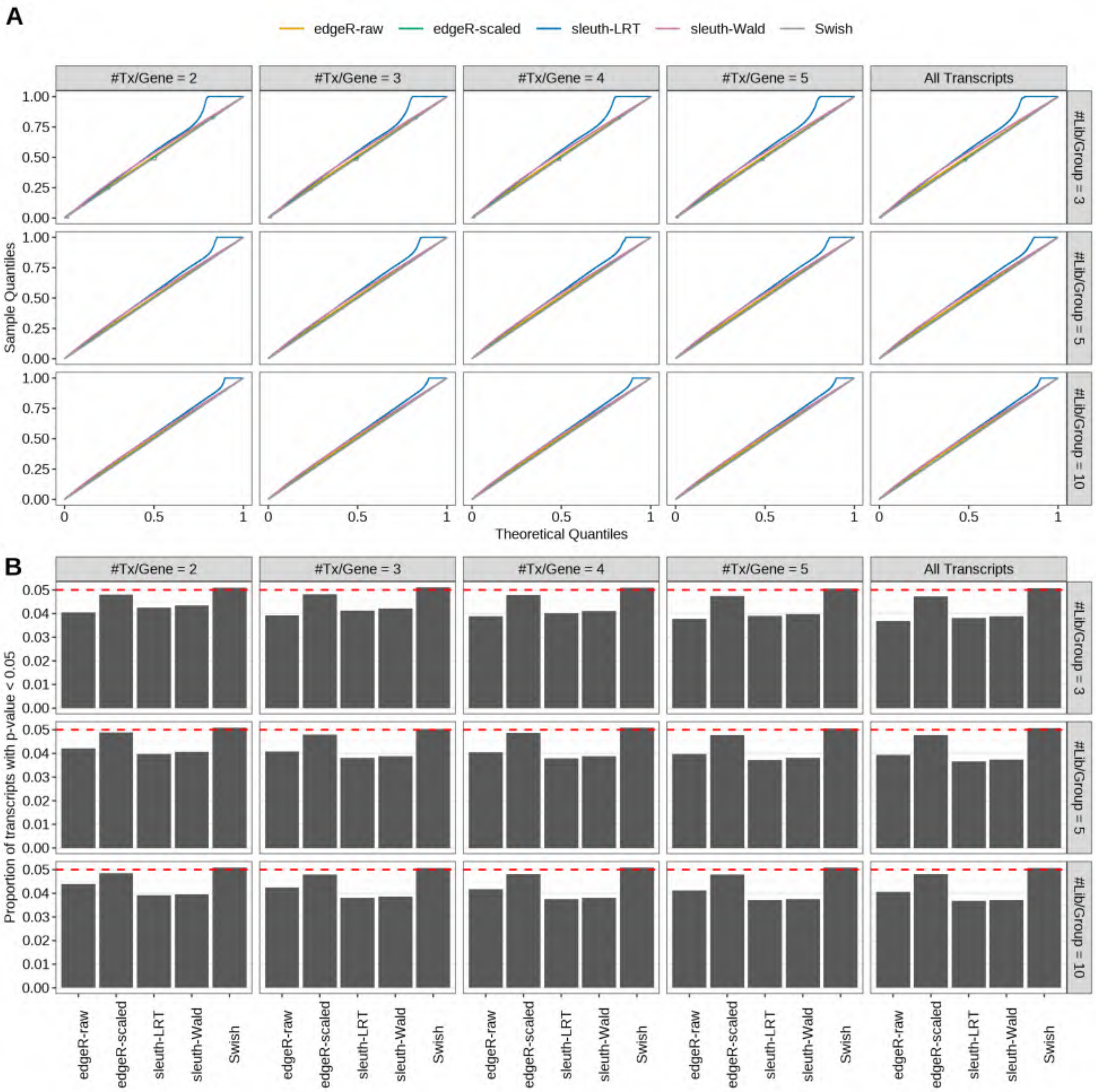

Figure S57: Simulation results. Scenario with mm39 genome, 75bp paired-end reads quantified with kallisto, and balanced libraries. (A) QQ plots of p-values for simulations without any differential expression (averaged over 20 simulations). (B) Proportion of transcripts with unadjusted p-values less than 0.05 for simulations without any differential expression (averaged over 20 simulations)

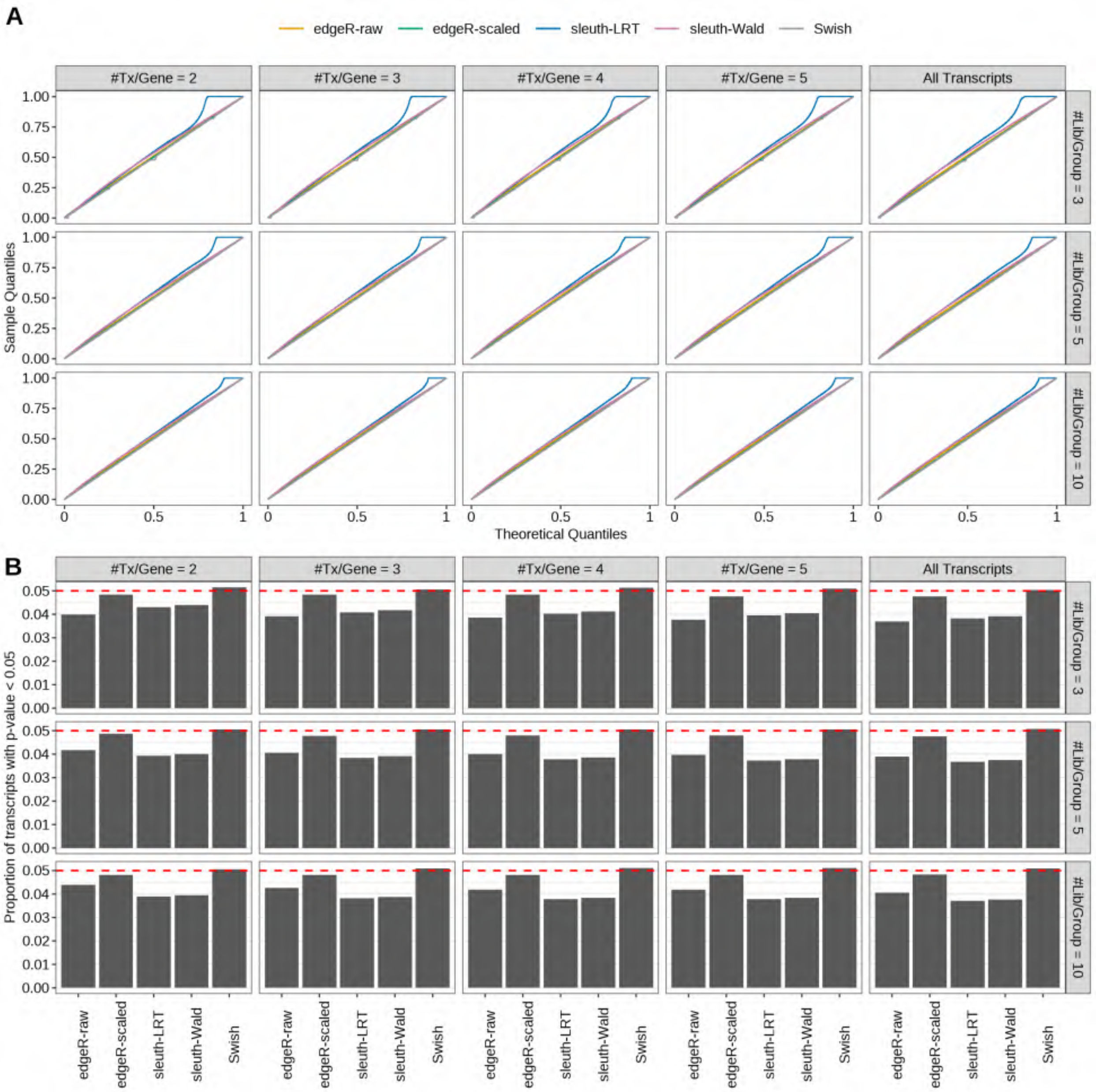

Figure S58: Simulation results. Scenario with mm39 genome, 100bp paired-end reads quantified with kallisto, and balanced libraries. (A) QQ plots of p-values for simulations without any differential expression (averaged over 20 simulations). (B) Proportion of transcripts with unadjusted p-values less than 0.05 for simulations without any differential expression (averaged over 20 simulations)

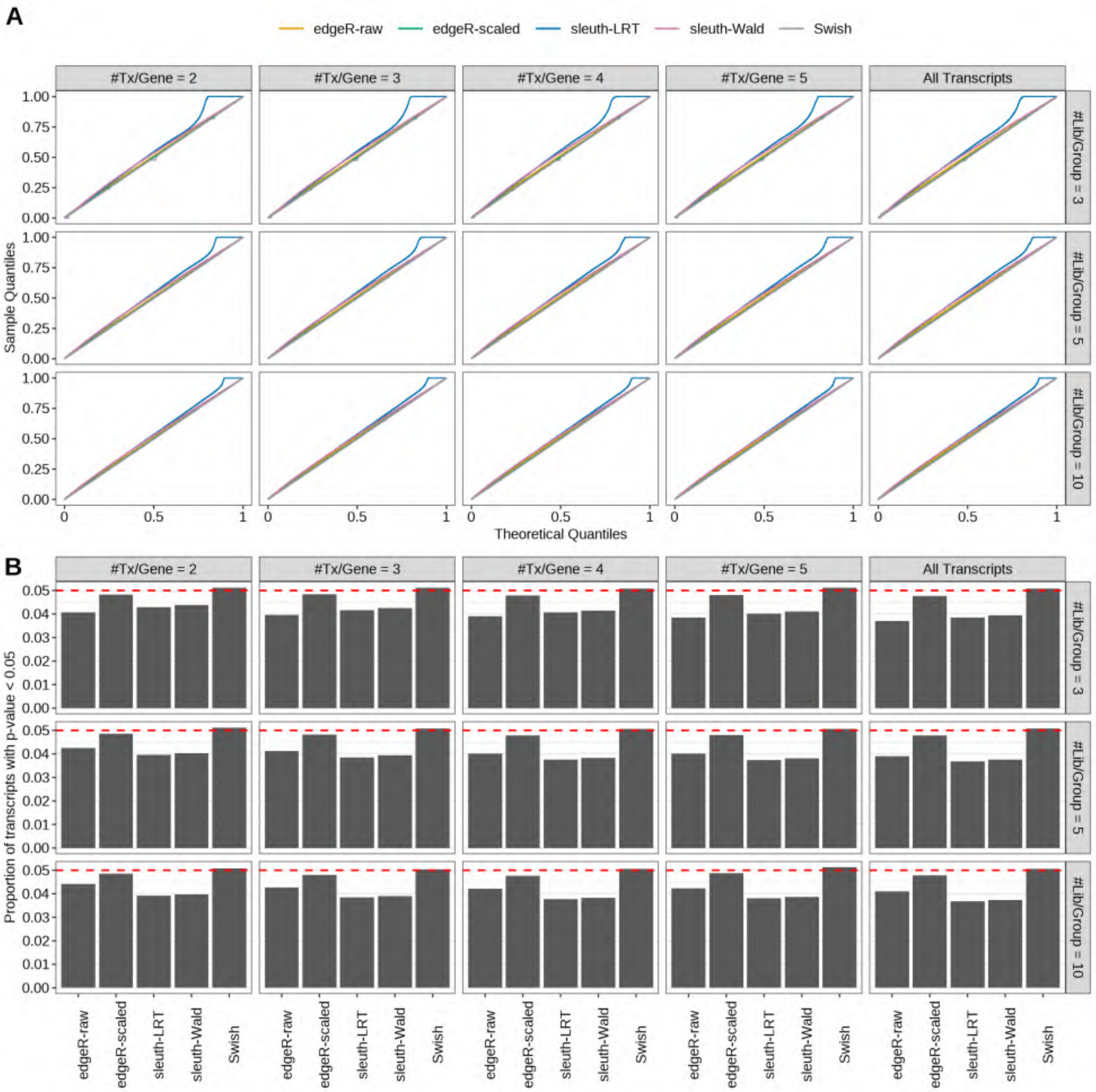

Figure S59: Simulation results. Scenario with mm39 genome, 125bp paired-end reads quantified with kallisto, and balanced libraries. (A) QQ plots of p-values for simulations without any differential expression (averaged over 20 simulations). (B) Proportion of transcripts with unadjusted p-values less than 0.05 for simulations without any differential expression (averaged over 20 simulations)

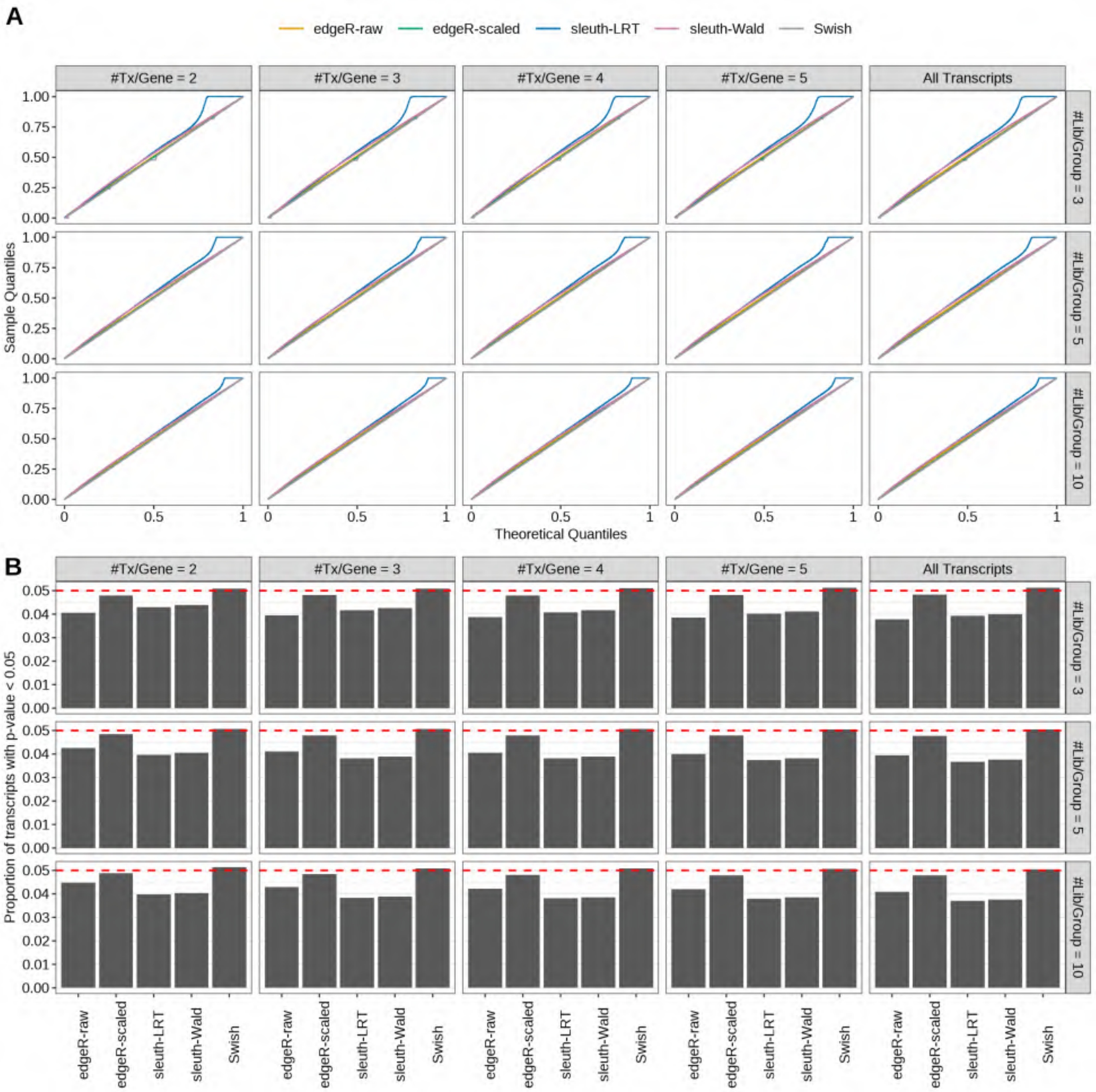

Figure S60: Simulation results. Scenario with mm39 genome, 150bp paired-end reads quantified with kallisto, and balanced libraries. (A) QQ plots of p-values for simulations without any differential expression (averaged over 20 simulations). (B) Proportion of transcripts with unadjusted p-values less than 0.05 for simulations without any differential expression (averaged over 20 simulations)

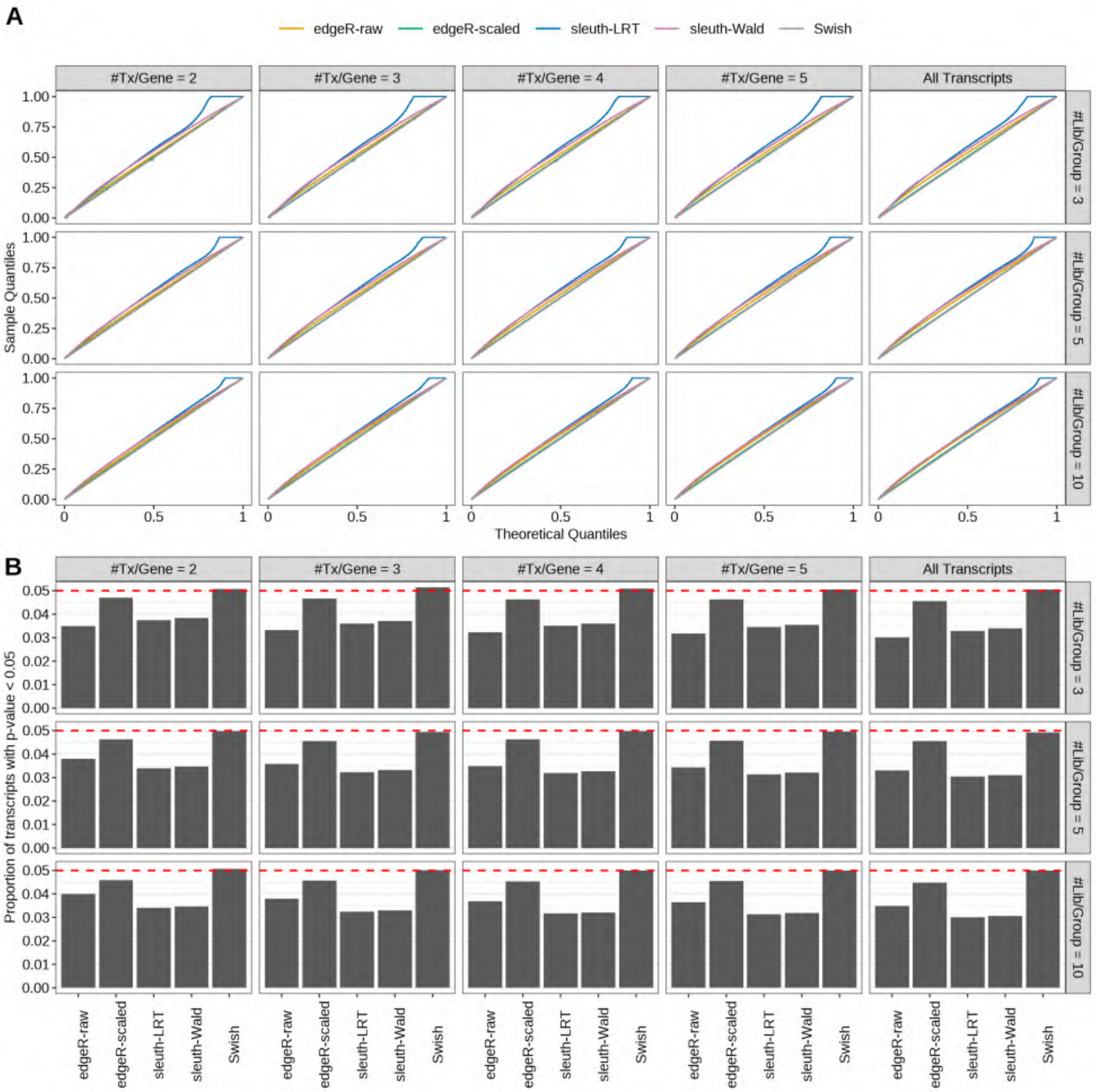

Figure S61: Simulation results. Scenario with mm39 genome, 50bp single-end reads quantified with Salmon, and unbalanced libraries. (A) QQ plots of p-values for simulations without any differential expression (averaged over 20 simulations). (B) Proportion of transcripts with unadjusted p-values less than 0.05 for simulations without any differential expression (averaged over 20 simulations)

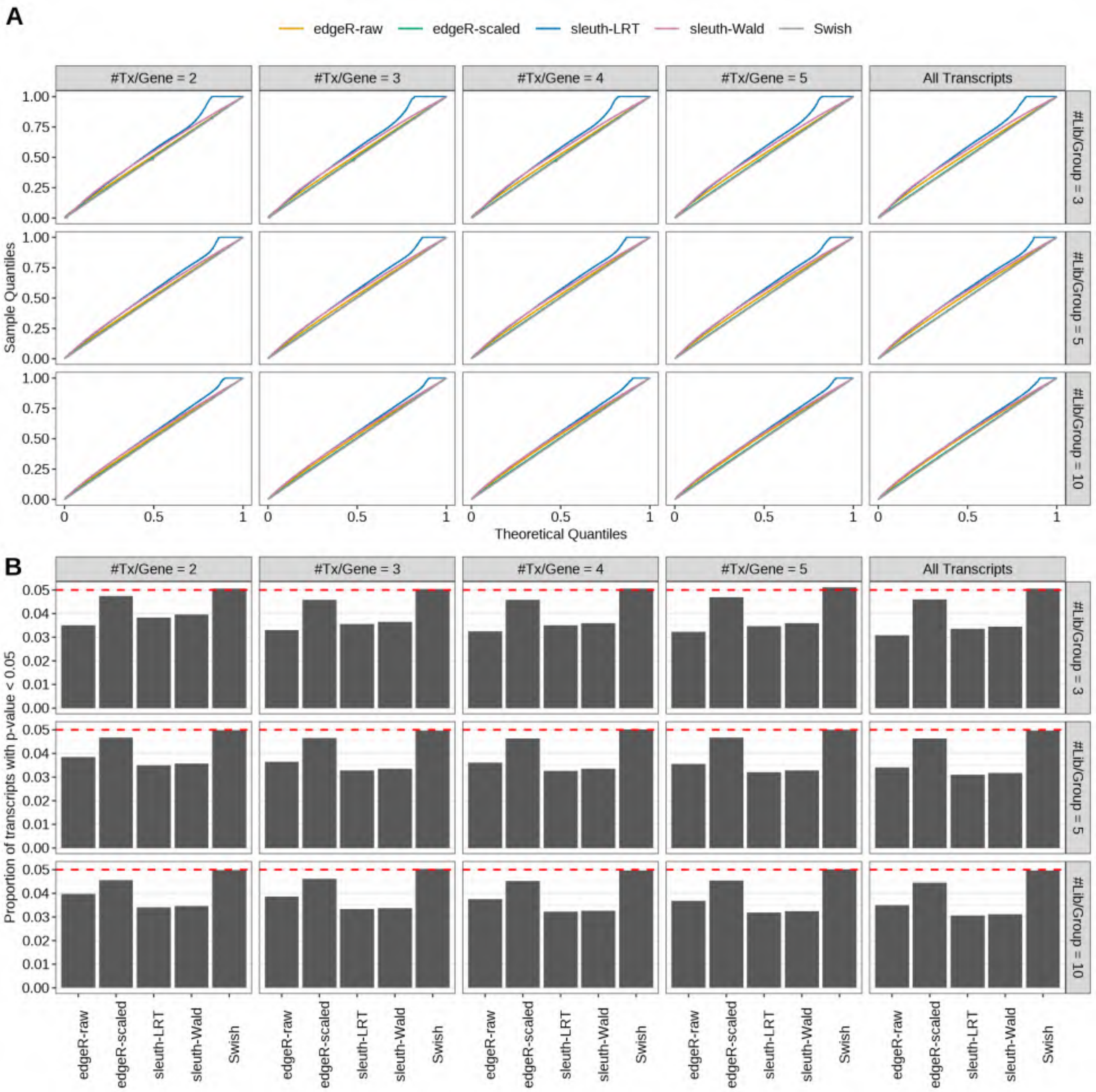

Figure S62: Simulation results. Scenario with mm39 genome, 75bp single-end reads quantified with Salmon, and unbalanced libraries. (A) QQ plots of p-values for simulations without any differential expression (averaged over 20 simulations). (B) Proportion of transcripts with unadjusted p-values less than 0.05 for simulations without any differential expression (averaged over 20 simulations)

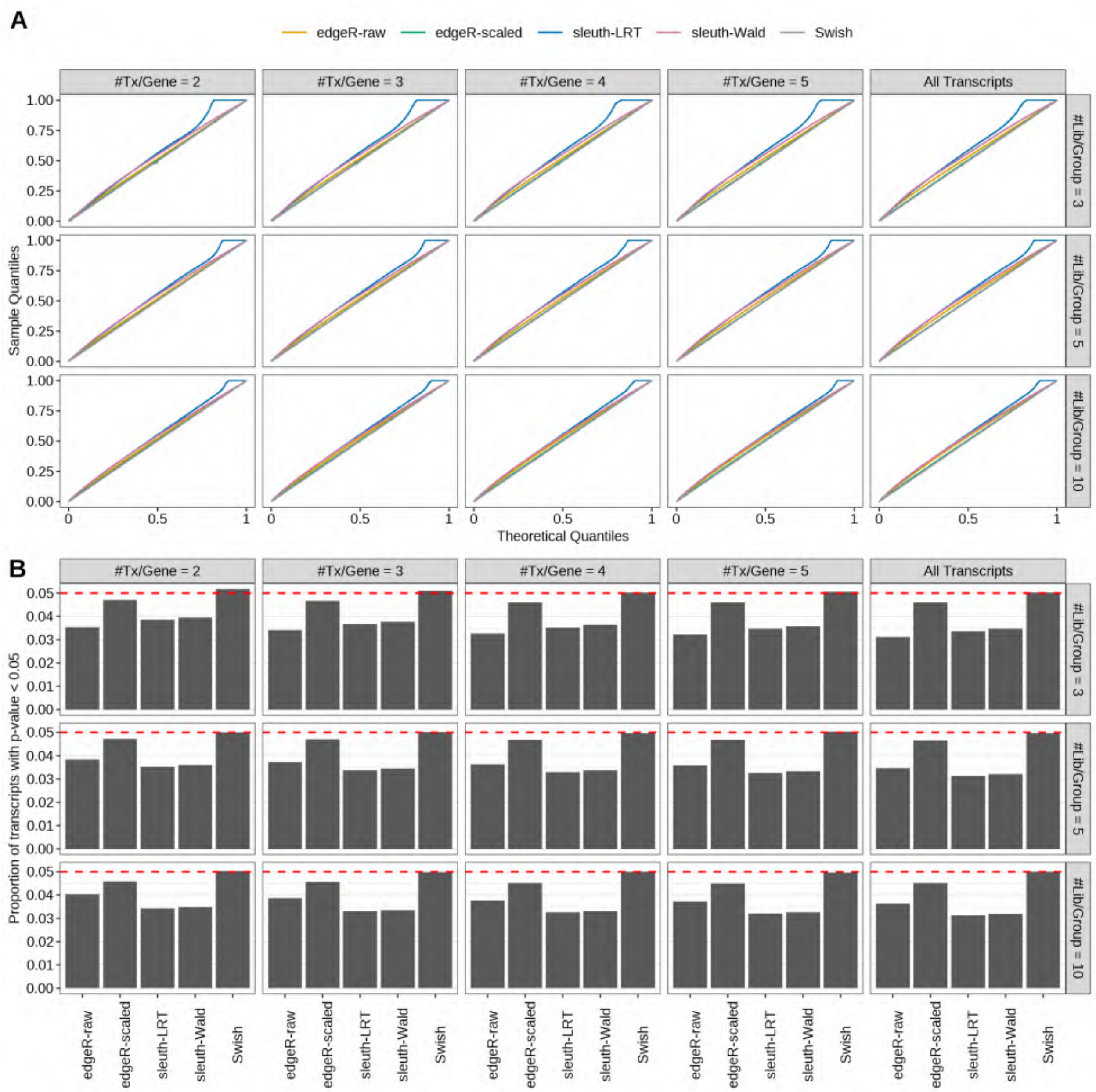

Figure S63: Simulation results. Scenario with mm39 genome, 100bp single-end reads quantified with Salmon, and unbalanced libraries. (A) QQ plots of p-values for simulations without any differential expression (averaged over 20 simulations). (B) Proportion of transcripts with unadjusted p-values less than 0.05 for simulations without any differential expression (averaged over 20 simulations)

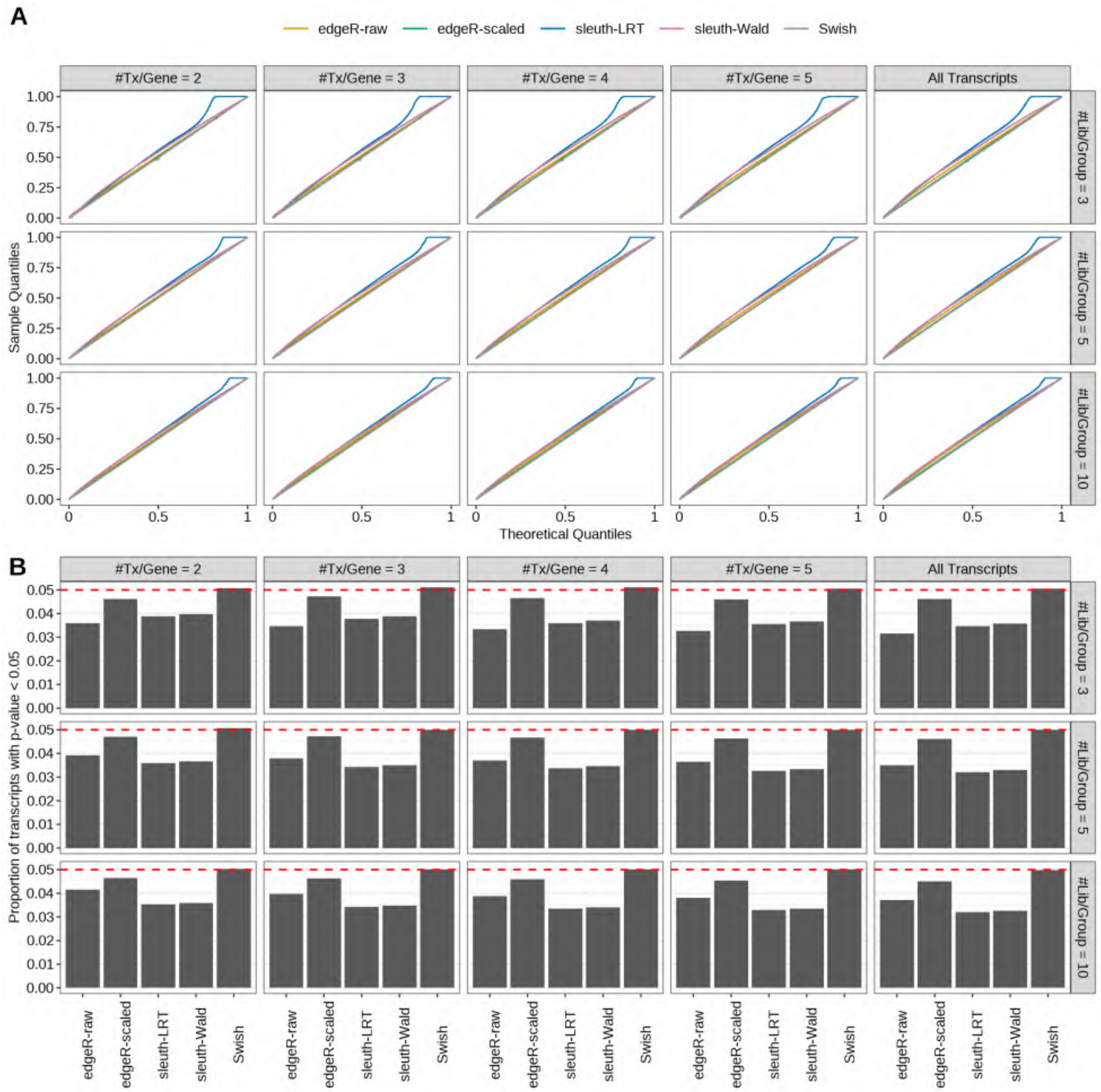

Figure S64: Simulation results. Scenario with mm39 genome, 125bp single-end reads quantified with Salmon, and unbalanced libraries. (A) QQ plots of p-values for simulations without any differential expression (averaged over 20 simulations). (B) Proportion of transcripts with unadjusted p-values less than 0.05 for simulations without any differential expression (averaged over 20 simulations)

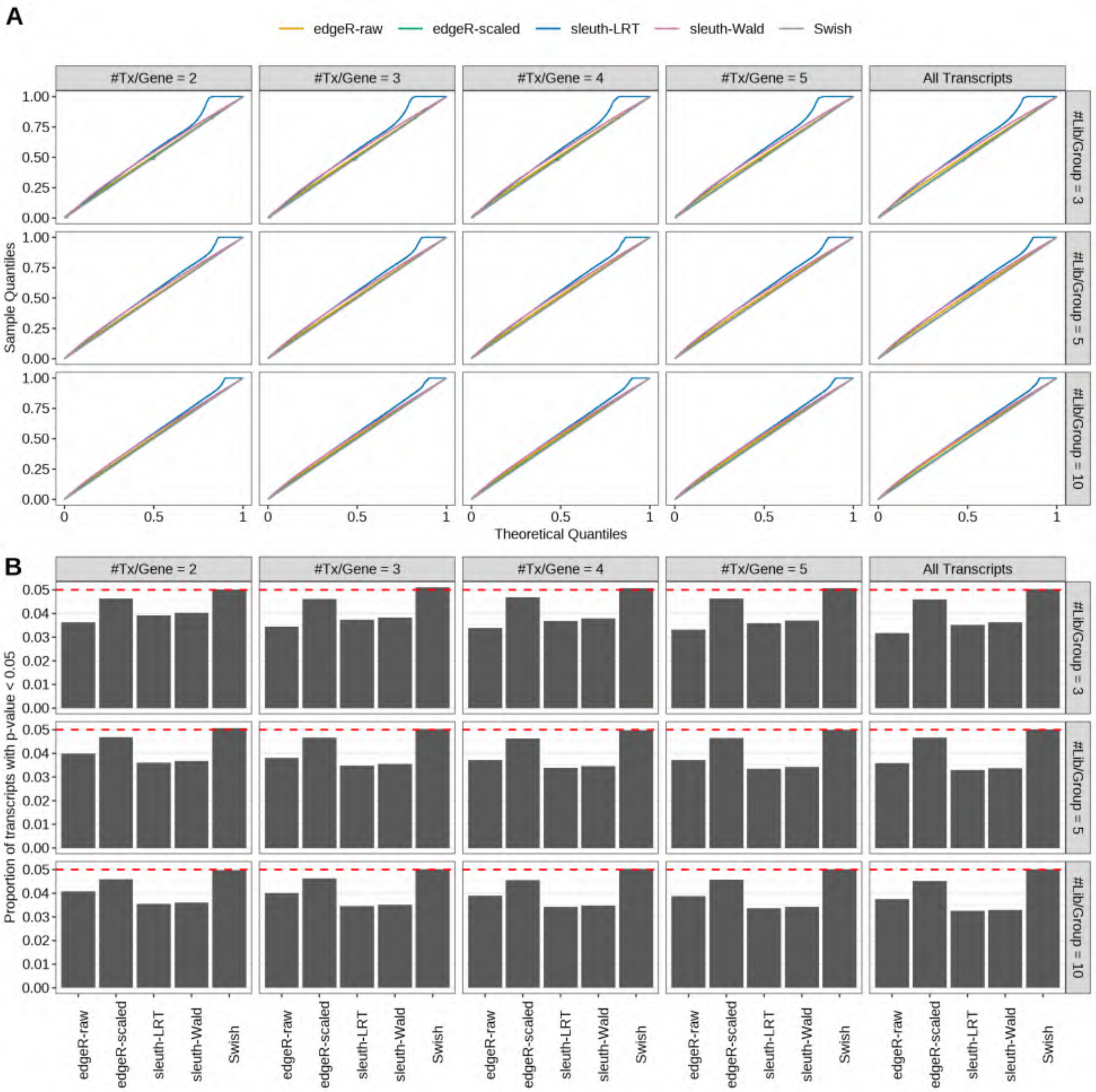

Figure S65: Simulation results. Scenario with mm39 genome, 150bp single-end reads quantified with Salmon, and unbalanced libraries. (A) QQ plots of p-values for simulations without any differential expression (averaged over 20 simulations). (B) Proportion of transcripts with unadjusted p-values less than 0.05 for simulations without any differential expression (averaged over 20 simulations)

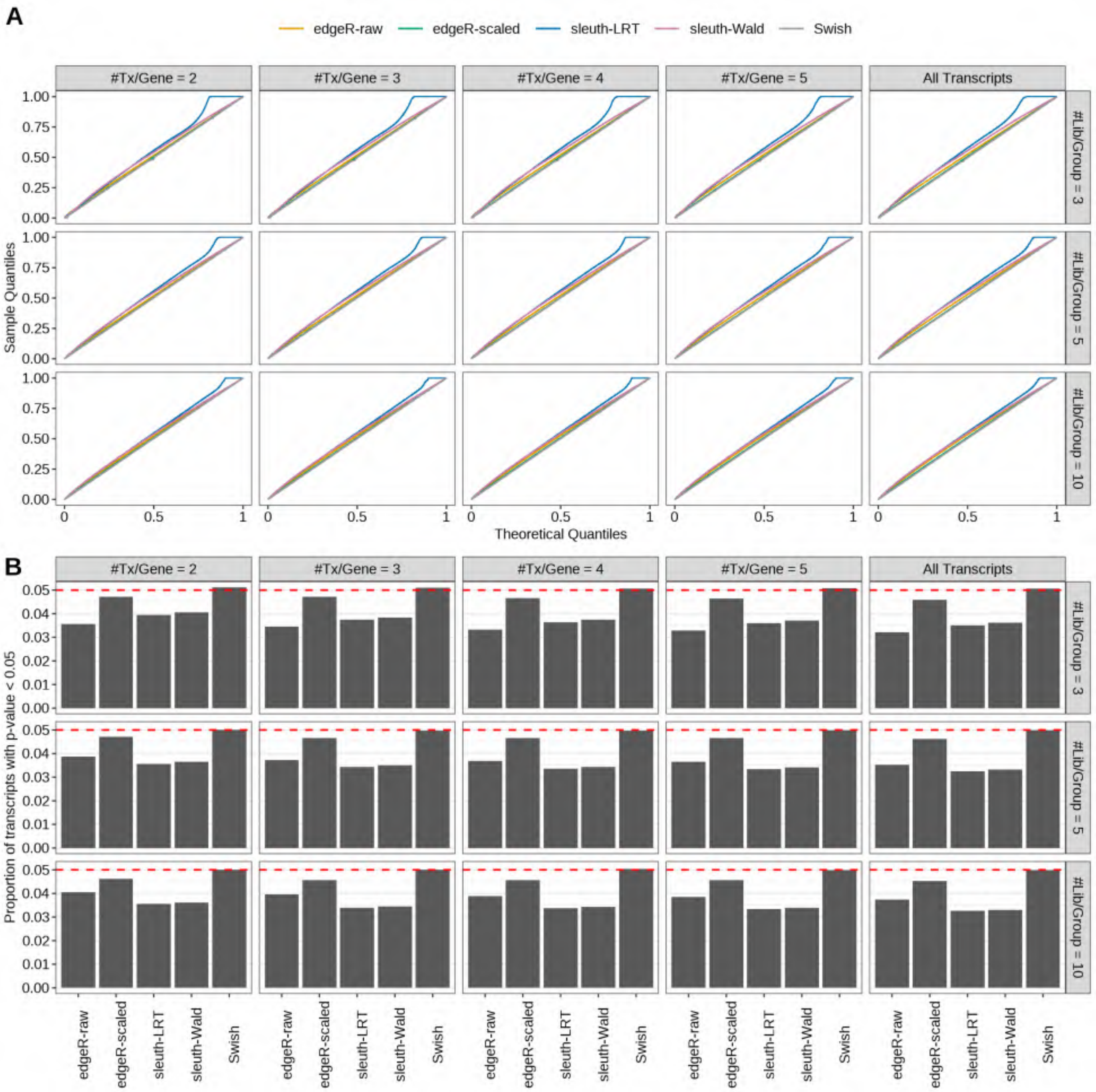

Figure S66: Simulation results. Scenario with mm39 genome, 50bp paired-end reads quantified with Salmon, and unbalanced libraries. (A) QQ plots of p-values for simulations without any differential expression (averaged over 20 simulations). (B) Proportion of transcripts with unadjusted p-values less than 0.05 for simulations without any differential expression (averaged over 20 simulations)

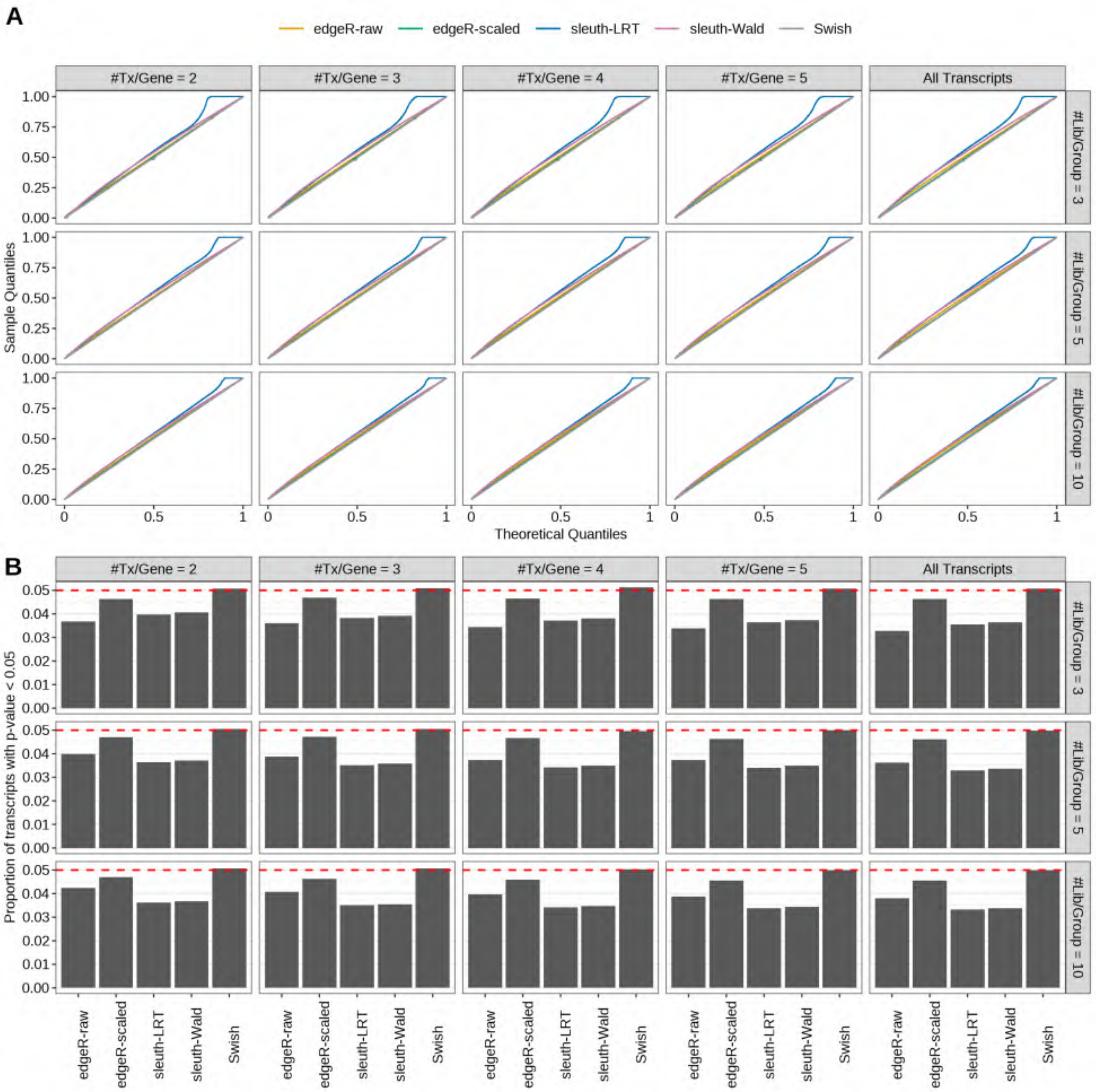

Figure S67: Simulation results. Scenario with mm39 genome, 75bp paired-end reads quantified with Salmon, and unbalanced libraries. (A) QQ plots of p-values for simulations without any differential expression (averaged over 20 simulations). (B) Proportion of transcripts with unadjusted p-values less than 0.05 for simulations without any differential expression (averaged over 20 simulations)

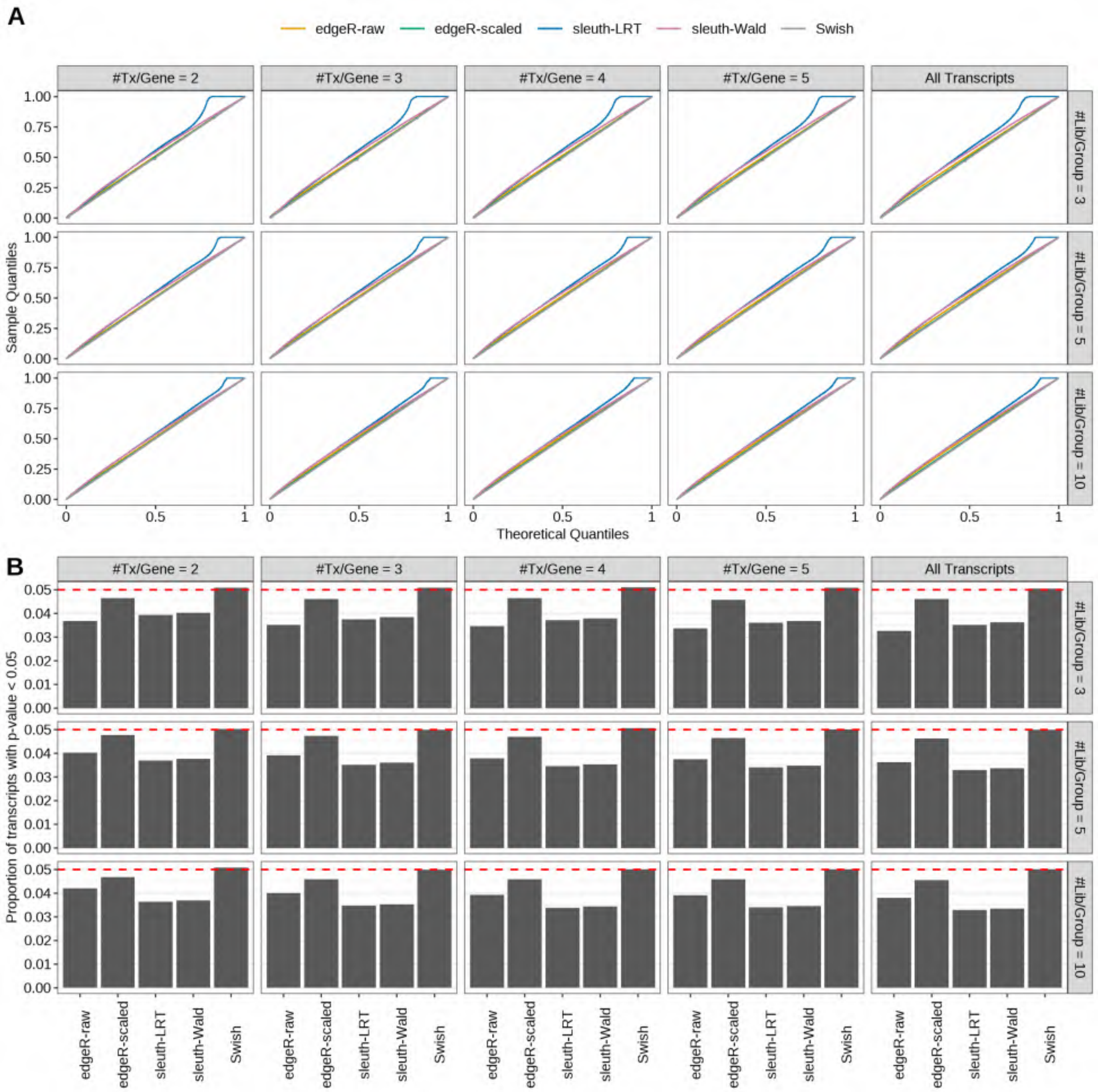

Figure S68: Simulation results. Scenario with mm39 genome, 100bp paired-end reads quantified with Salmon, and unbalanced libraries. (A) QQ plots of p-values for simulations without any differential expression (averaged over 20 simulations). (B) Proportion of transcripts with unadjusted p-values less than 0.05 for simulations without any differential expression (averaged over 20 simulations)

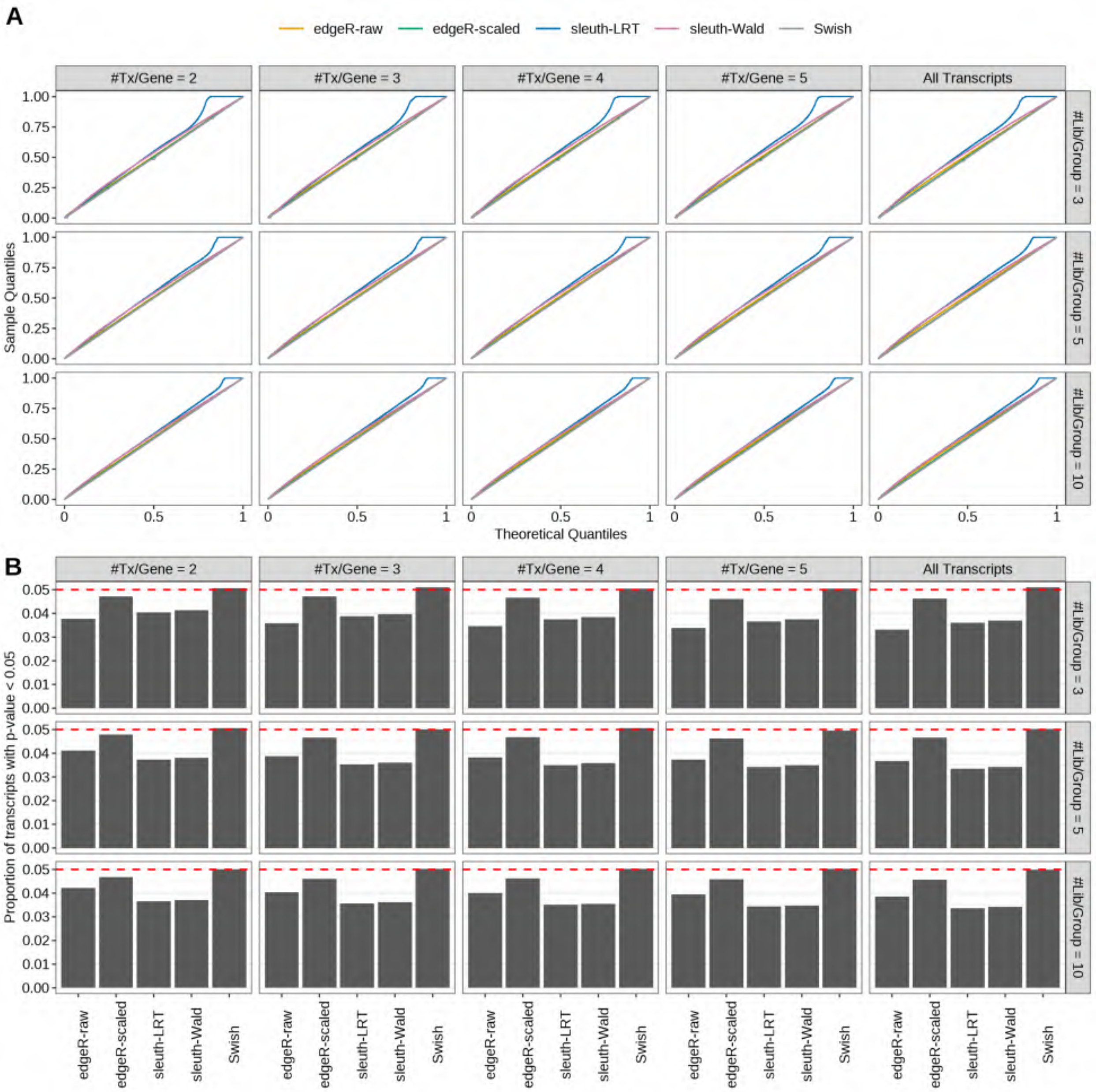

Figure S69: Simulation results. Scenario with mm39 genome, 125bp paired-end reads quantified with Salmon, and unbalanced libraries. (A) QQ plots of p-values for simulations without any differential expression (averaged over 20 simulations). (B) Proportion of transcripts with unadjusted p-values less than 0.05 for simulations without any differential expression (averaged over 20 simulations)

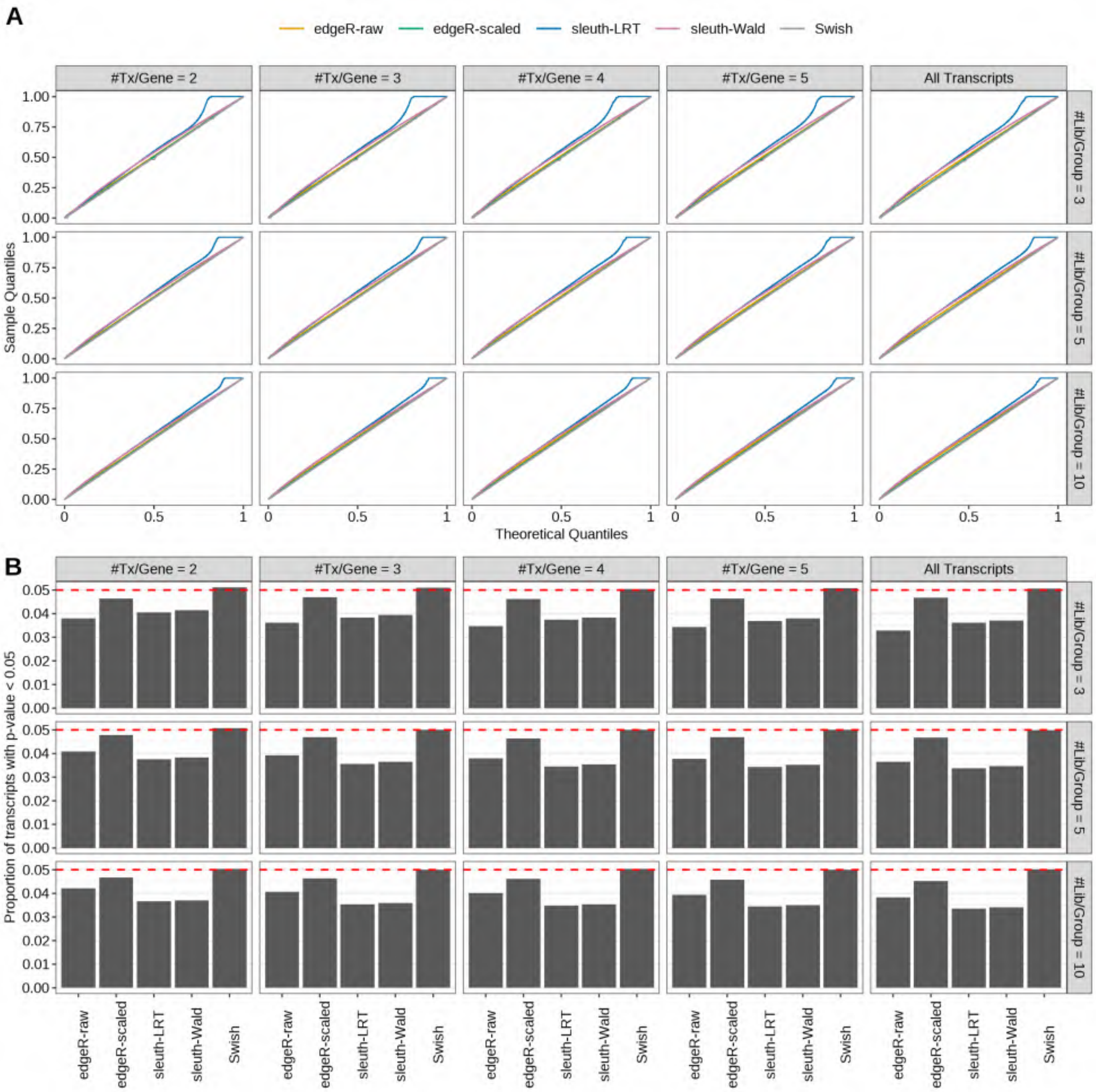

Figure S70: Simulation results. Scenario with mm39 genome, 150bp paired-end reads quantified with Salmon, and unbalanced libraries. (A) QQ plots of p-values for simulations without any differential expression (averaged over 20 simulations). (B) Proportion of transcripts with unadjusted p-values less than 0.05 for simulations without any differential expression (averaged over 20 simulations)

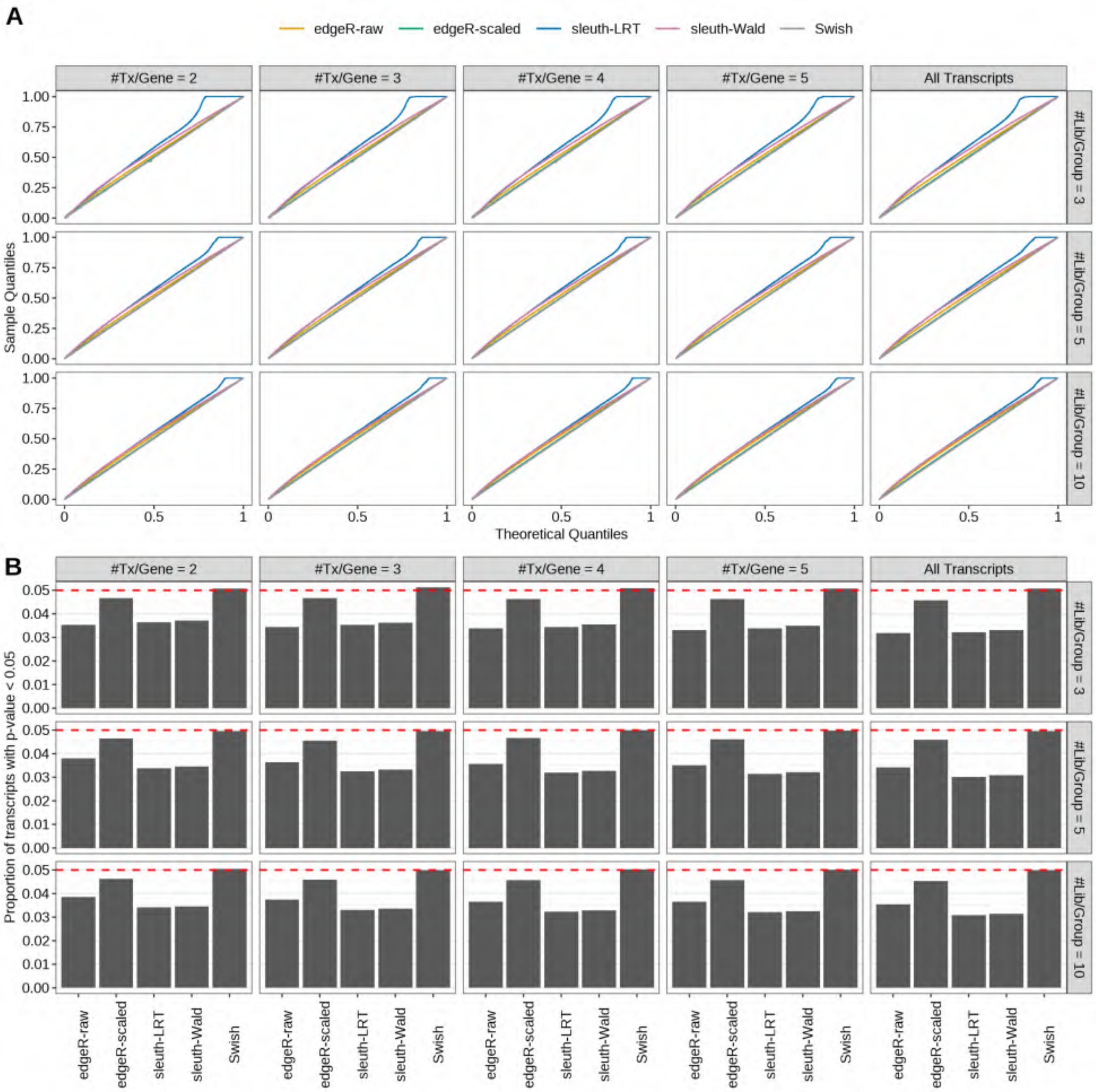

Figure S71: Simulation results. Scenario with mm39 genome, 50bp single-end reads quantified with kallisto, and unbalanced libraries. (A) QQ plots of p-values for simulations without any differential expression (averaged over 20 simulations). (B) Proportion of transcripts with unadjusted p-values less than 0.05 for simulations without any differential expression (averaged over 20 simulations)

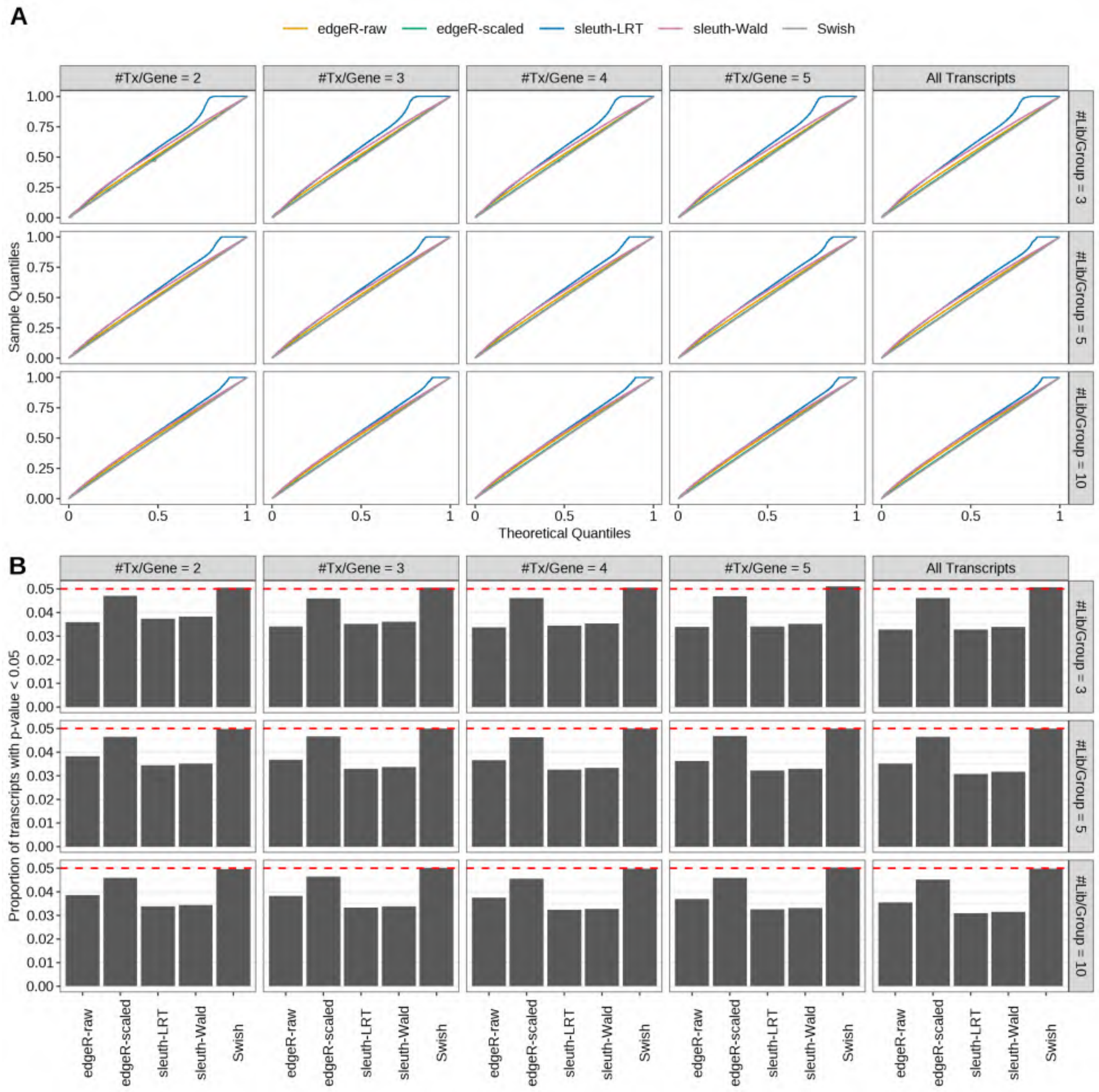

Figure S72: Simulation results. Scenario with mm39 genome, 75bp single-end reads quantified with kallisto, and unbalanced libraries. (A) QQ plots of p-values for simulations without any differential expression (averaged over 20 simulations). (B) Proportion of transcripts with unadjusted p-values less than 0.05 for simulations without any differential expression (averaged over 20 simulations)

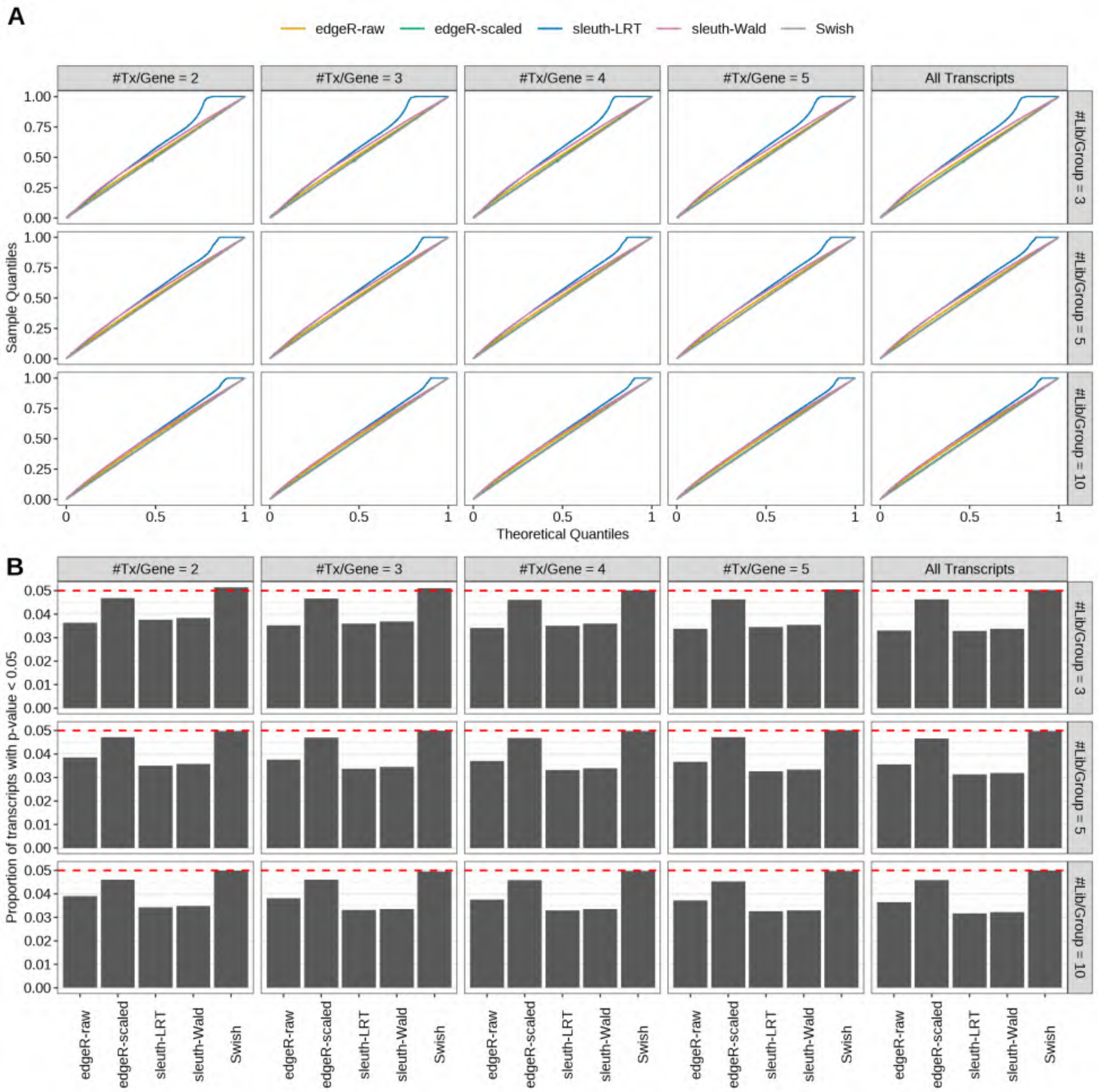

Figure S73: Simulation results. Scenario with mm39 genome, 100bp single-end reads quantified with kallisto, and unbalanced libraries. (A) QQ plots of p-values for simulations without any differential expression (averaged over 20 simulations). (B) Proportion of transcripts with unadjusted p-values less than 0.05 for simulations without any differential expression (averaged over 20 simulations)

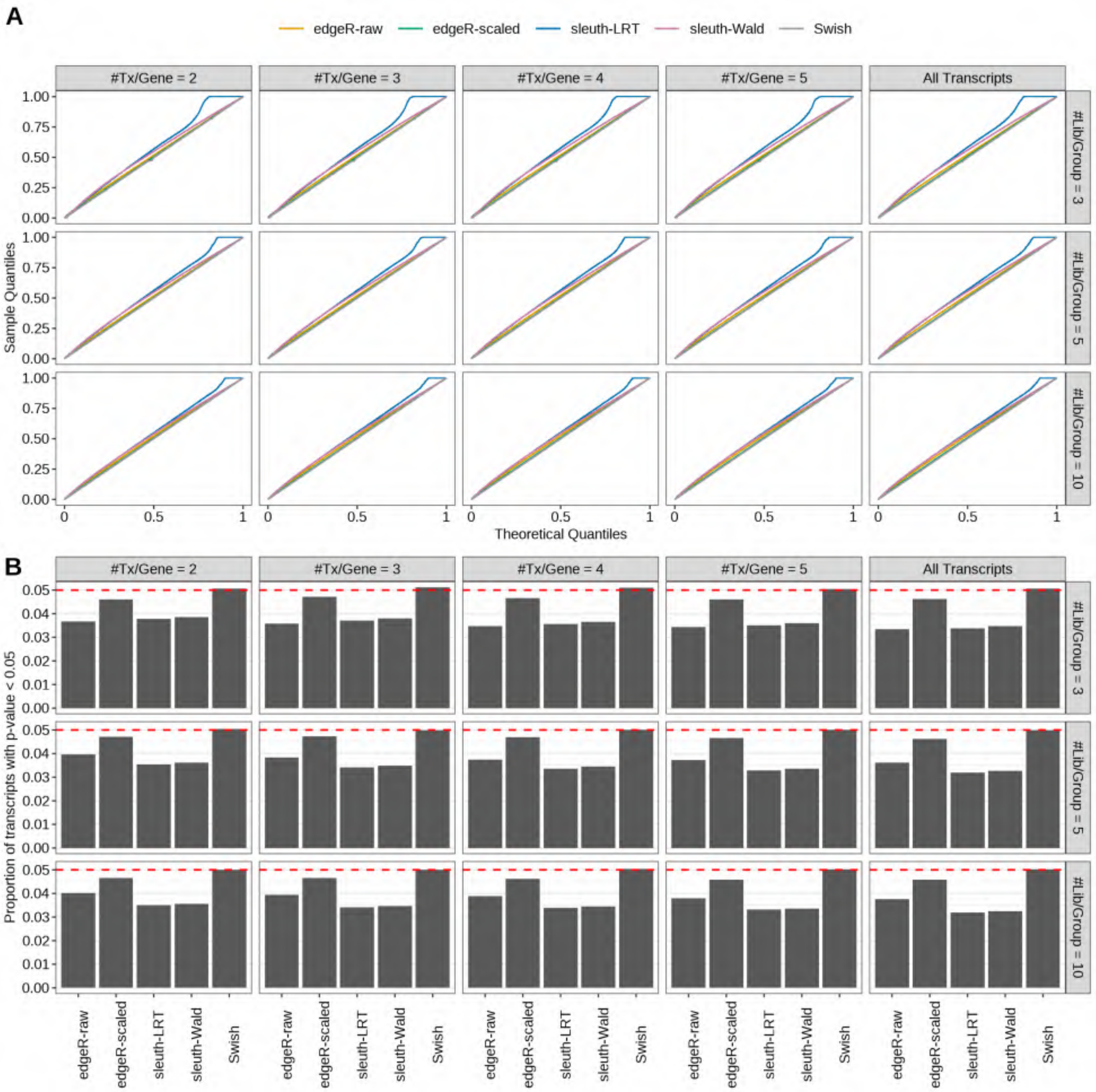

Figure S74: Simulation results. Scenario with mm39 genome, 125bp single-end reads quantified with kallisto, and unbalanced libraries. (A) QQ plots of p-values for simulations without any differential expression (averaged over 20 simulations). (B) Proportion of transcripts with unadjusted p-values less than 0.05 for simulations without any differential expression (averaged over 20 simulations)

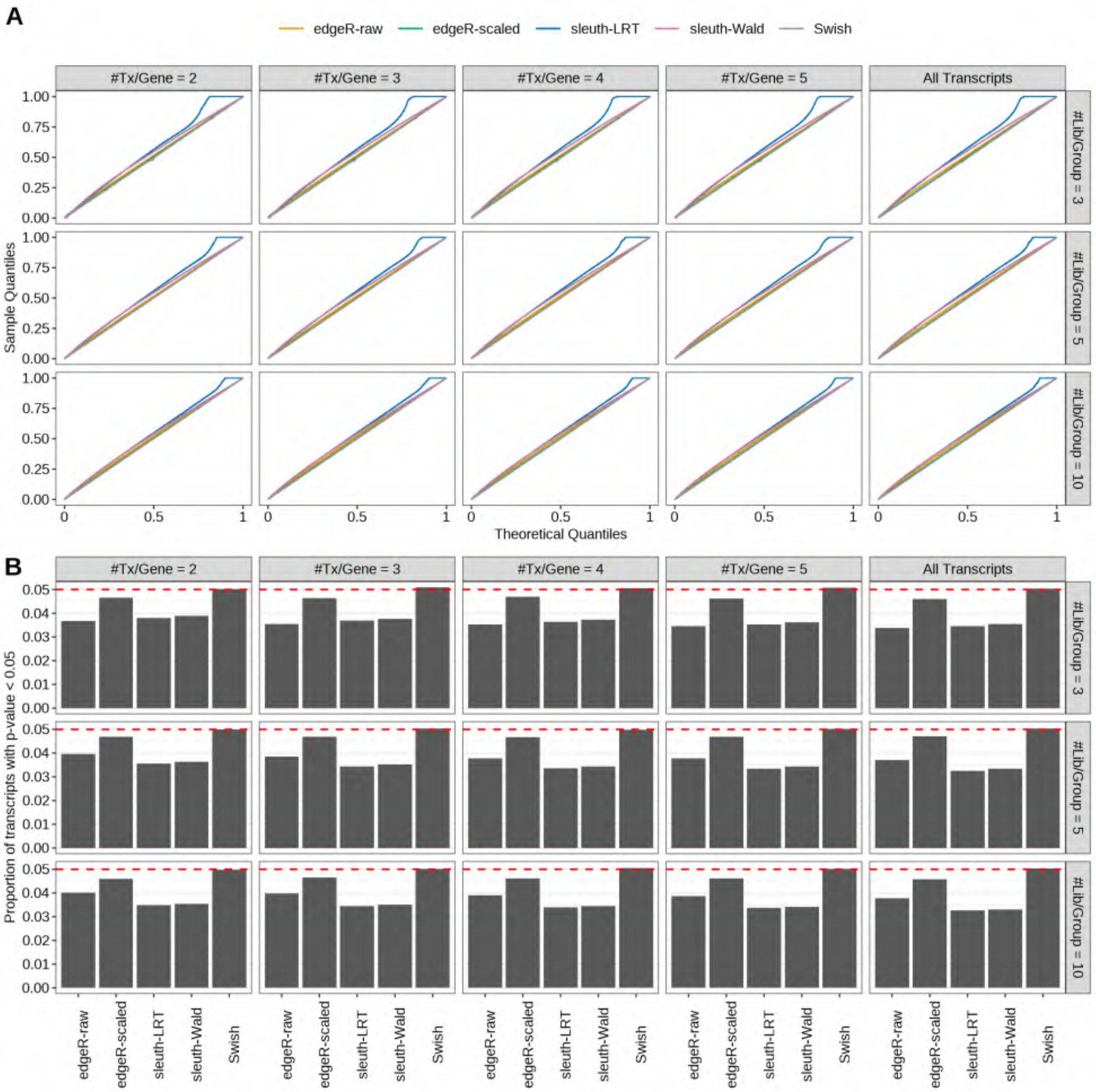

Figure S75: Simulation results. Scenario with mm39 genome, 150bp single-end reads quantified with kallisto, and unbalanced libraries. (A) QQ plots of p-values for simulations without any differential expression (averaged over 20 simulations). (B) Proportion of transcripts with unadjusted p-values less than 0.05 for simulations without any differential expression (averaged over 20 simulations)

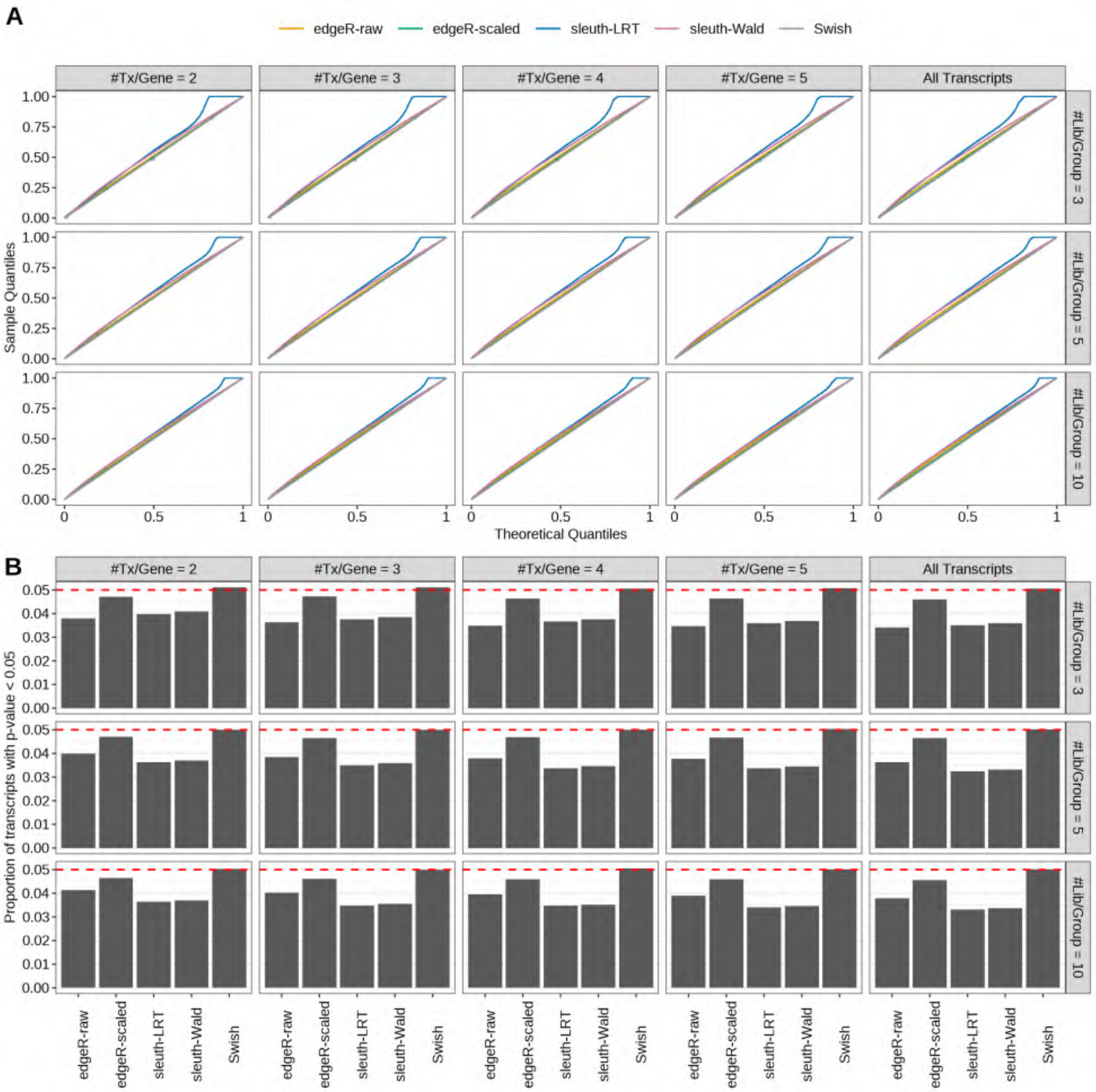

Figure S76: Simulation results. Scenario with mm39 genome, 50bp paired-end reads quantified with kallisto, and unbalanced libraries. (A) QQ plots of p-values for simulations without any differential expression (averaged over 20 simulations). (B) Proportion of transcripts with unadjusted p-values less than 0.05 for simulations without any differential expression (averaged over 20 simulations)

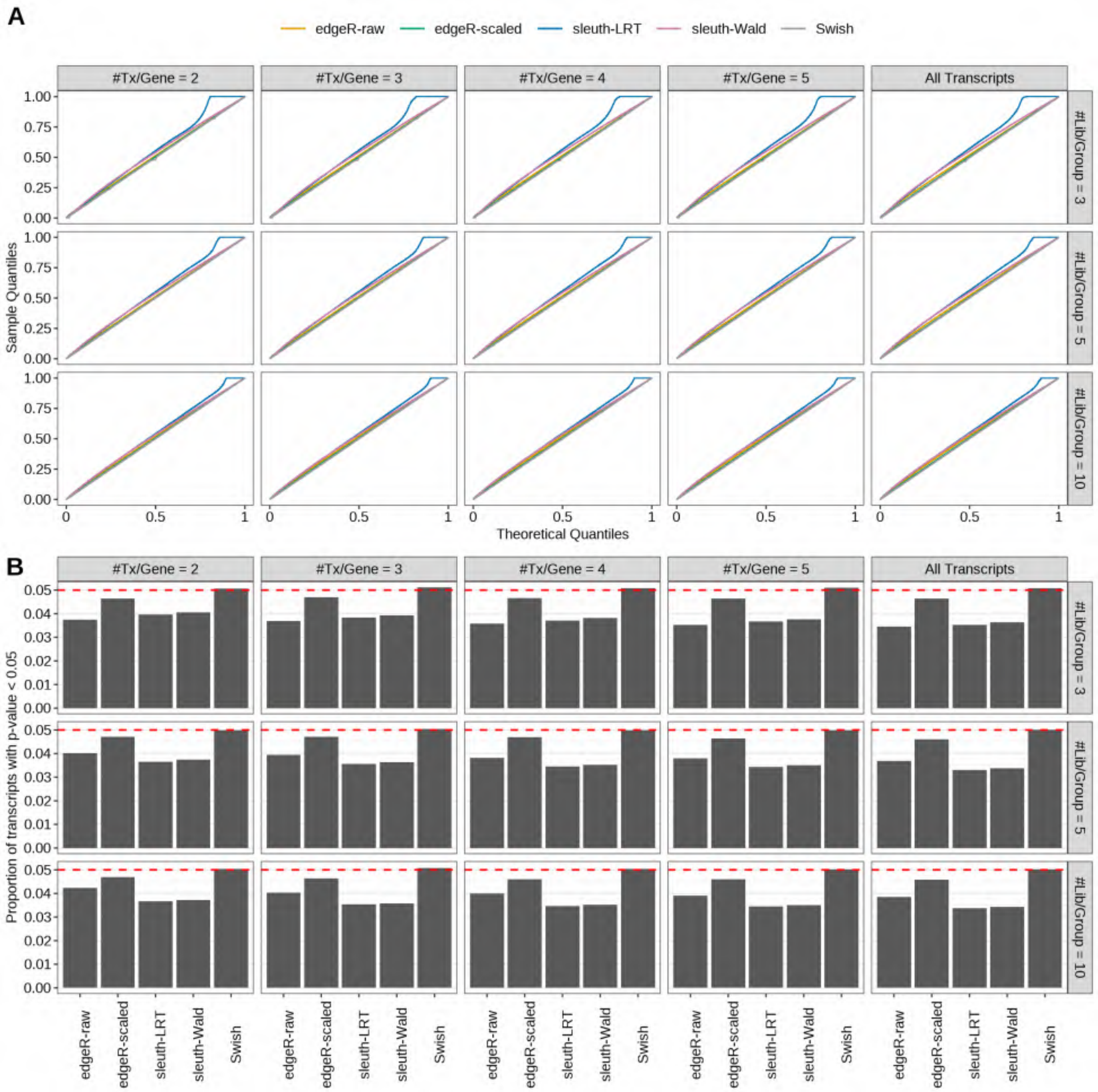

Figure S77: Simulation results. Scenario with mm39 genome, 75bp paired-end reads quantified with kallisto, and unbalanced libraries. (A) QQ plots of p-values for simulations without any differential expression (averaged over 20 simulations). (B) Proportion of transcripts with unadjusted p-values less than 0.05 for simulations without any differential expression (averaged over 20 simulations)

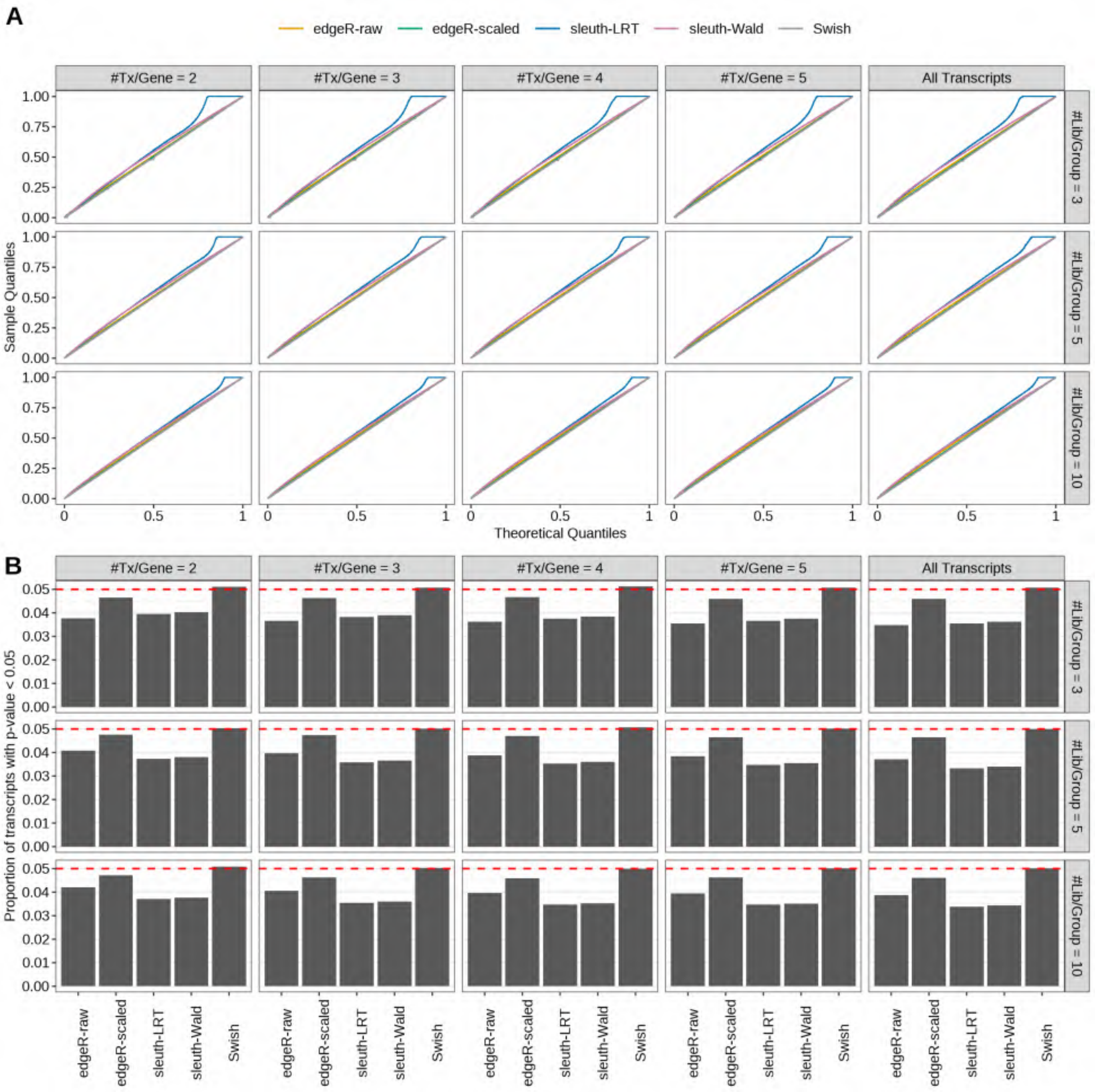

Figure S78: Simulation results. Scenario with mm39 genome, 100bp paired-end reads quantified with kallisto, and unbalanced libraries. (A) QQ plots of p-values for simulations without any differential expression (averaged over 20 simulations). (B) Proportion of transcripts with unadjusted p-values less than 0.05 for simulations without any differential expression (averaged over 20 simulations)

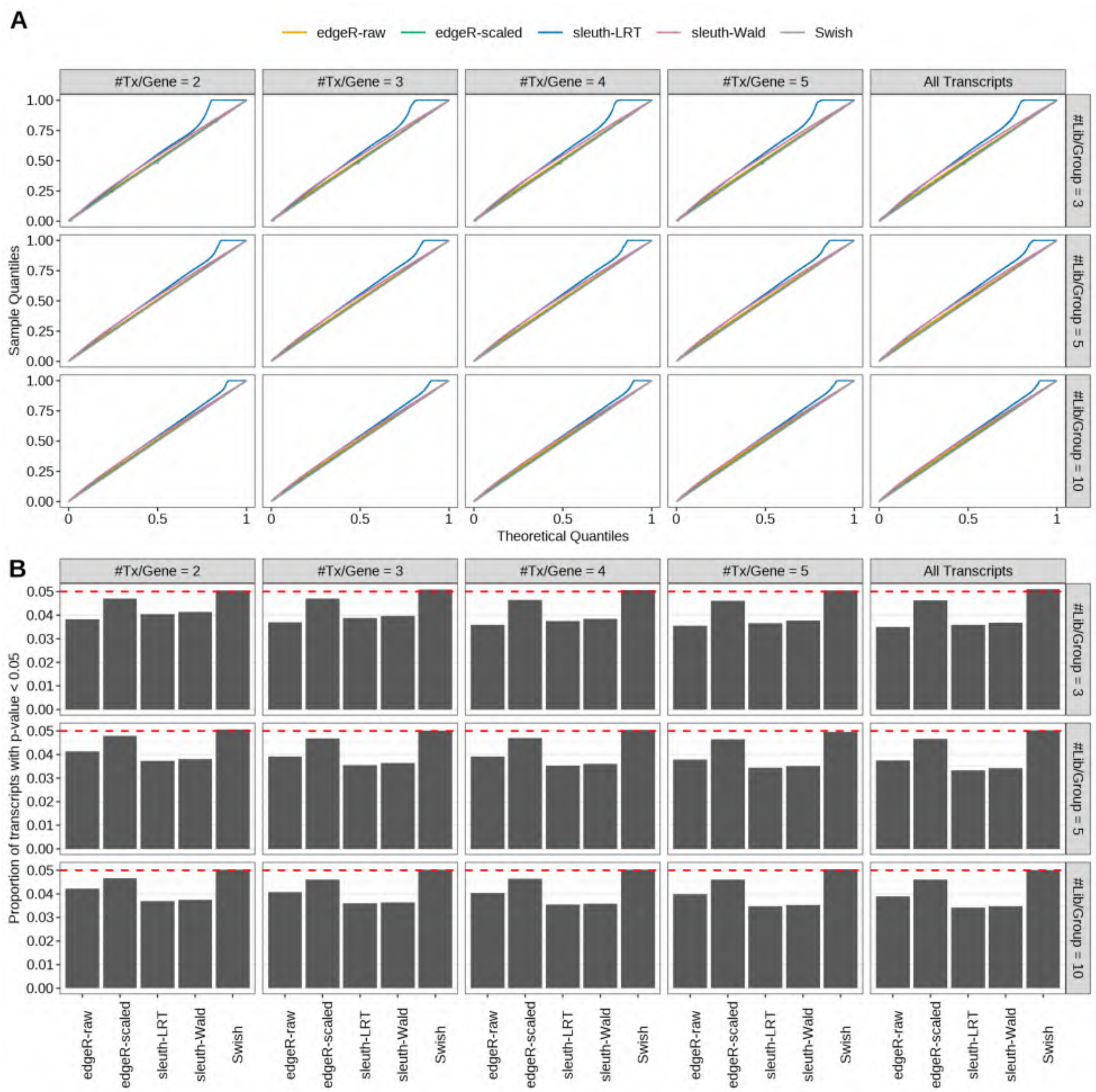

Figure S79: Simulation results. Scenario with mm39 genome, 125bp paired-end reads quantified with kallisto, and unbalanced libraries. (A) QQ plots of p-values for simulations without any differential expression (averaged over 20 simulations). (B) Proportion of transcripts with unadjusted p-values less than 0.05 for simulations without any differential expression (averaged over 20 simulations)

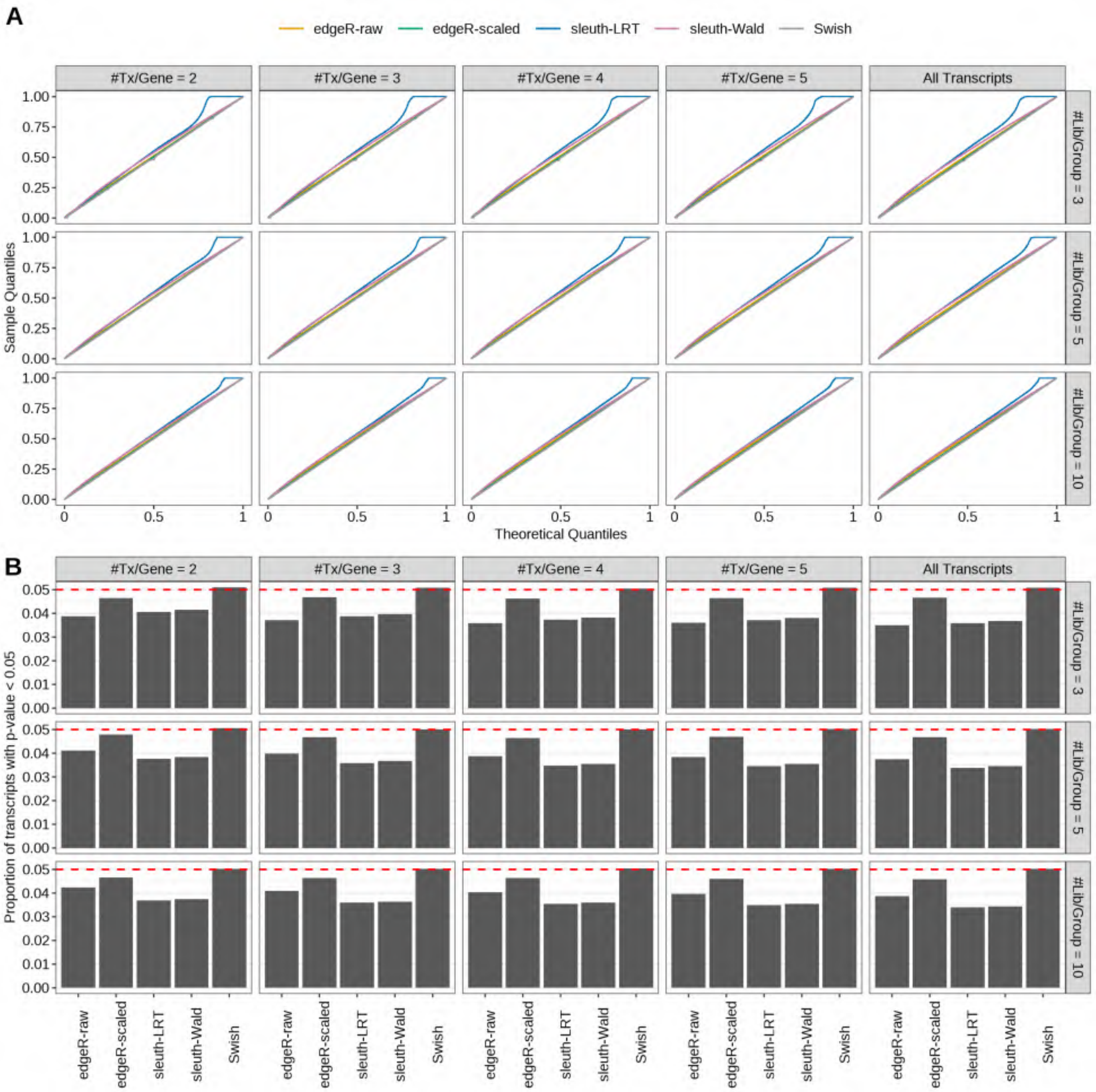

Figure S80: Simulation results. Scenario with mm39 genome, 150bp paired-end reads quantified with kallisto, and unbalanced libraries. (A) QQ plots of p-values for simulations without any differential expression (averaged over 20 simulations). (B) Proportion of transcripts with unadjusted p-values less than 0.05 for simulations without any differential expression (averaged over 20 simulations)

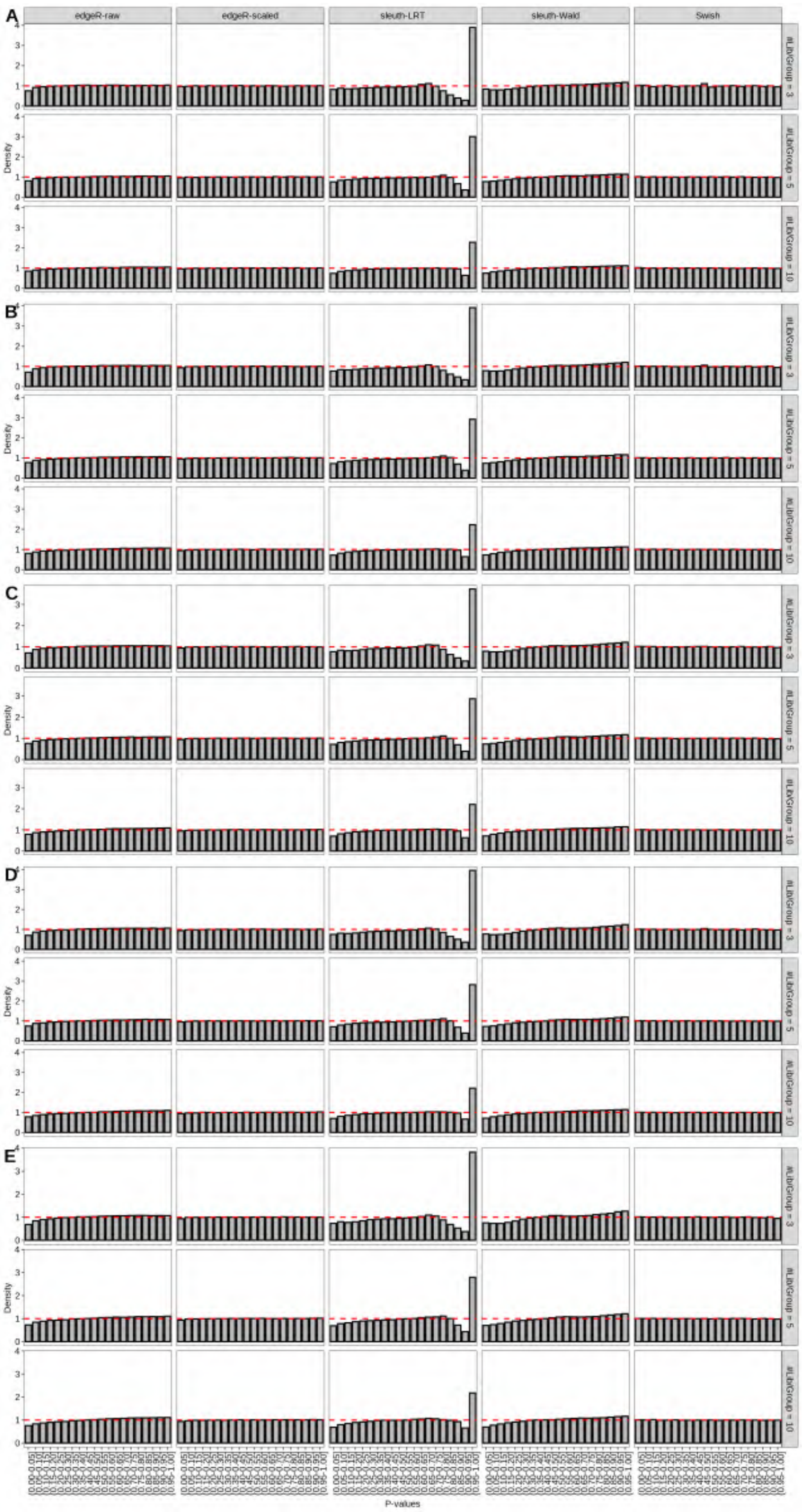

Figure S81: Simulation results. Scenario with mm39 genome, 50bp single-end reads quantified with Salmon, and balanced libraries. Density histograms of p-values from null simulations (averaged over 20 simulations). (A) Top 2 transcripts/gene expressed. (B) Top 3 transcripts/gene expressed. (C) Top 4 transcripts/gene expressed. (D) Top 5 transcripts/gene expressed. (E) All transcripts from the reference dataset are expressed.

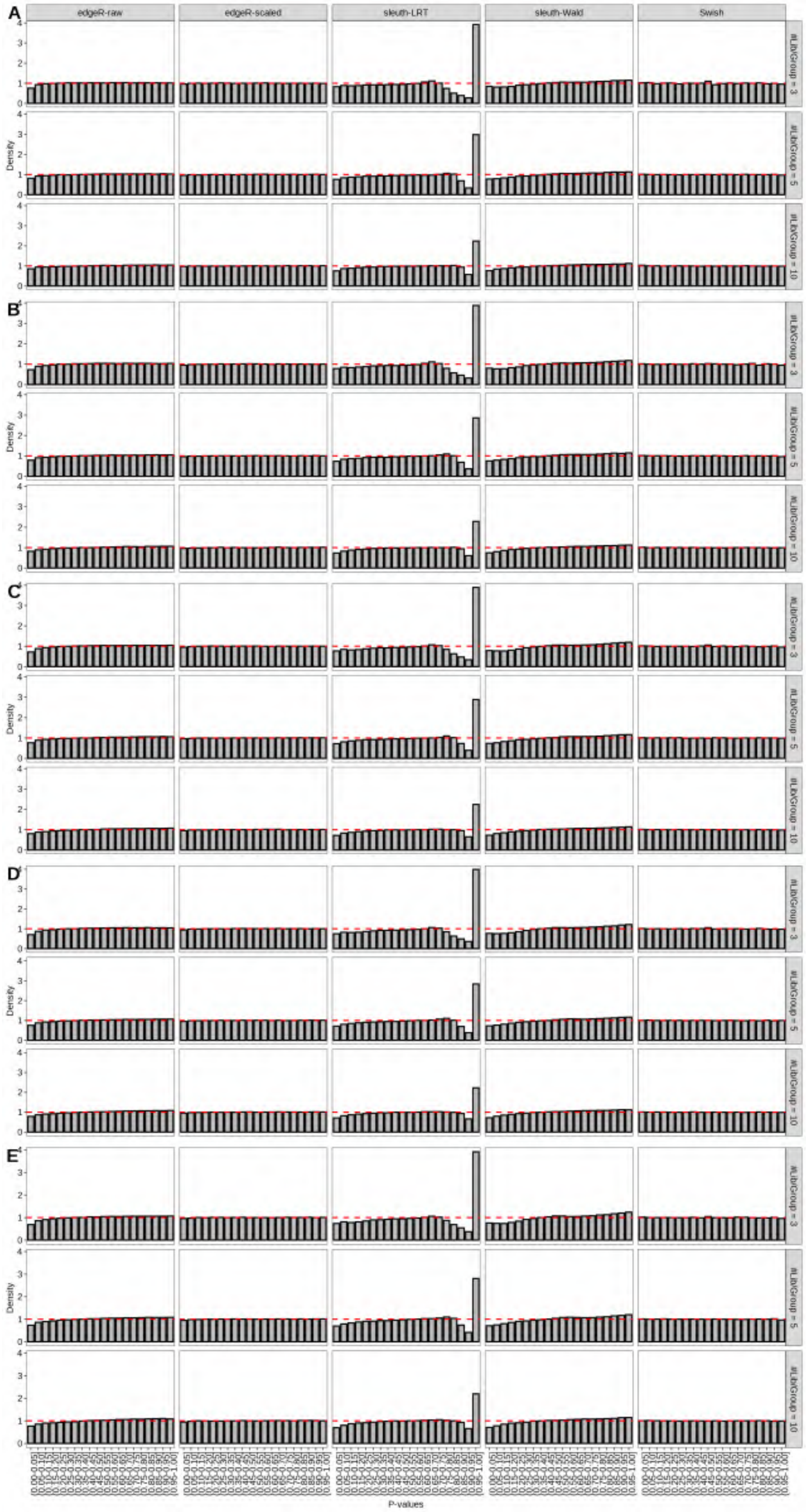

Figure S82: Simulation results. Scenario with mm39 genome, 75bp single-end reads quantified with Salmon, and balanced libraries. Density histograms of p-values from null simulations (averaged over 20 simulations). (A) Top 2 transcripts/gene expressed. (B) Top 3 transcripts/gene expressed. (C) Top 4 transcripts/gene expressed. (D) Top 5 transcripts/gene expressed. (E) All transcripts from the reference dataset are expressed.

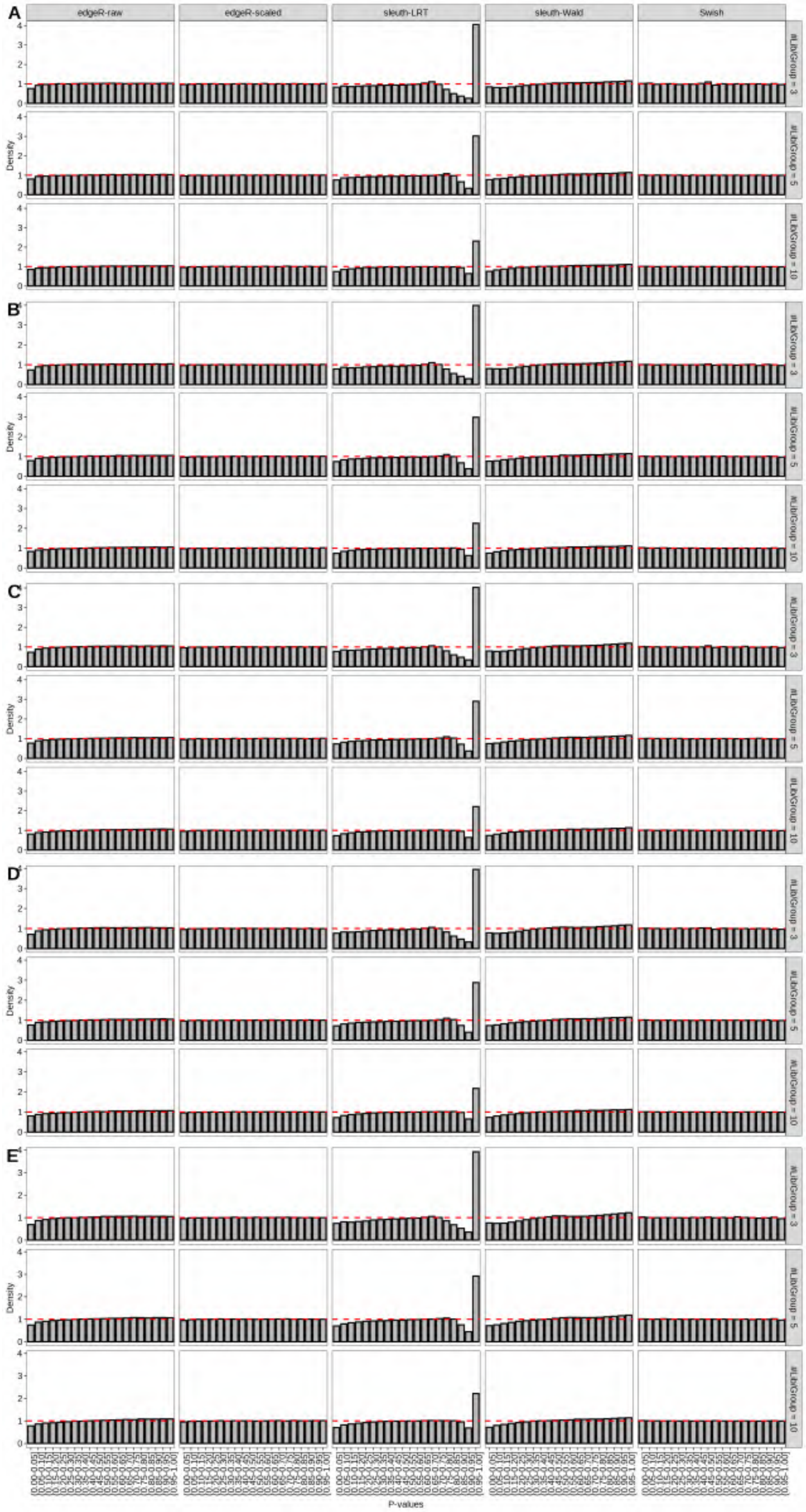

Figure S83: Simulation results. Scenario with mm39 genome, 100bp single-end reads quantified with Salmon, and balanced libraries. Density histograms of p-values from null simulations (averaged over 20 simulations). (A) Top 2 transcripts/gene expressed. (B) Top 3 transcripts/gene expressed. (C) Top 4 transcripts/gene expressed. (D) Top 5 transcripts/gene expressed. (E) All transcripts from the reference dataset are expressed.

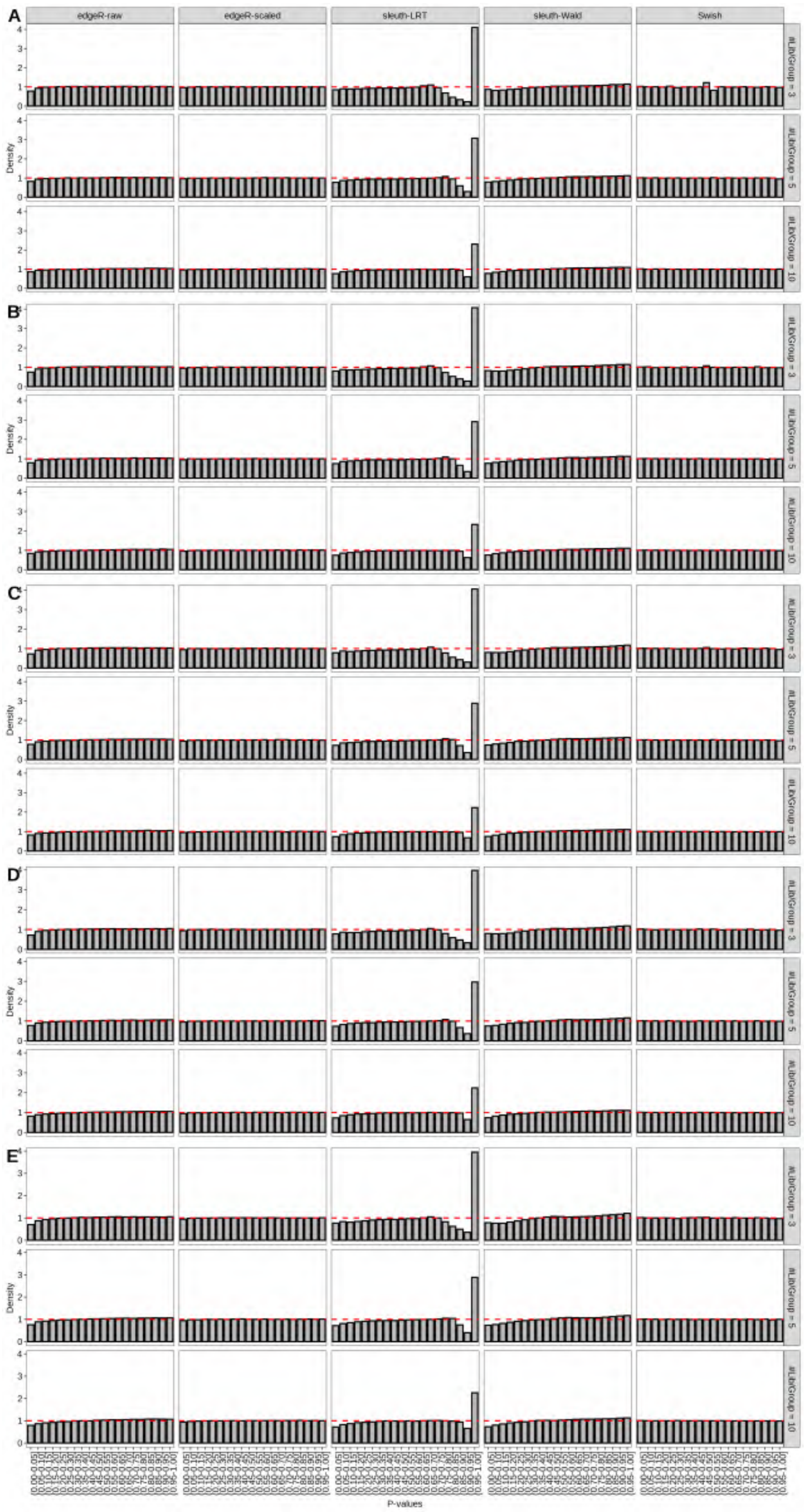

Figure S84: Simulation results. Scenario with mm39 genome, 125bp single-end reads quantified with Salmon, and balanced libraries. Density histograms of p-values from null simulations (averaged over 20 simulations). (A) Top 2 transcripts/gene expressed. (B) Top 3 transcripts/gene expressed. (C) Top 4 transcripts/gene expressed. (D) Top 5 transcripts/gene expressed. (E) All transcripts from the reference dataset are expressed.

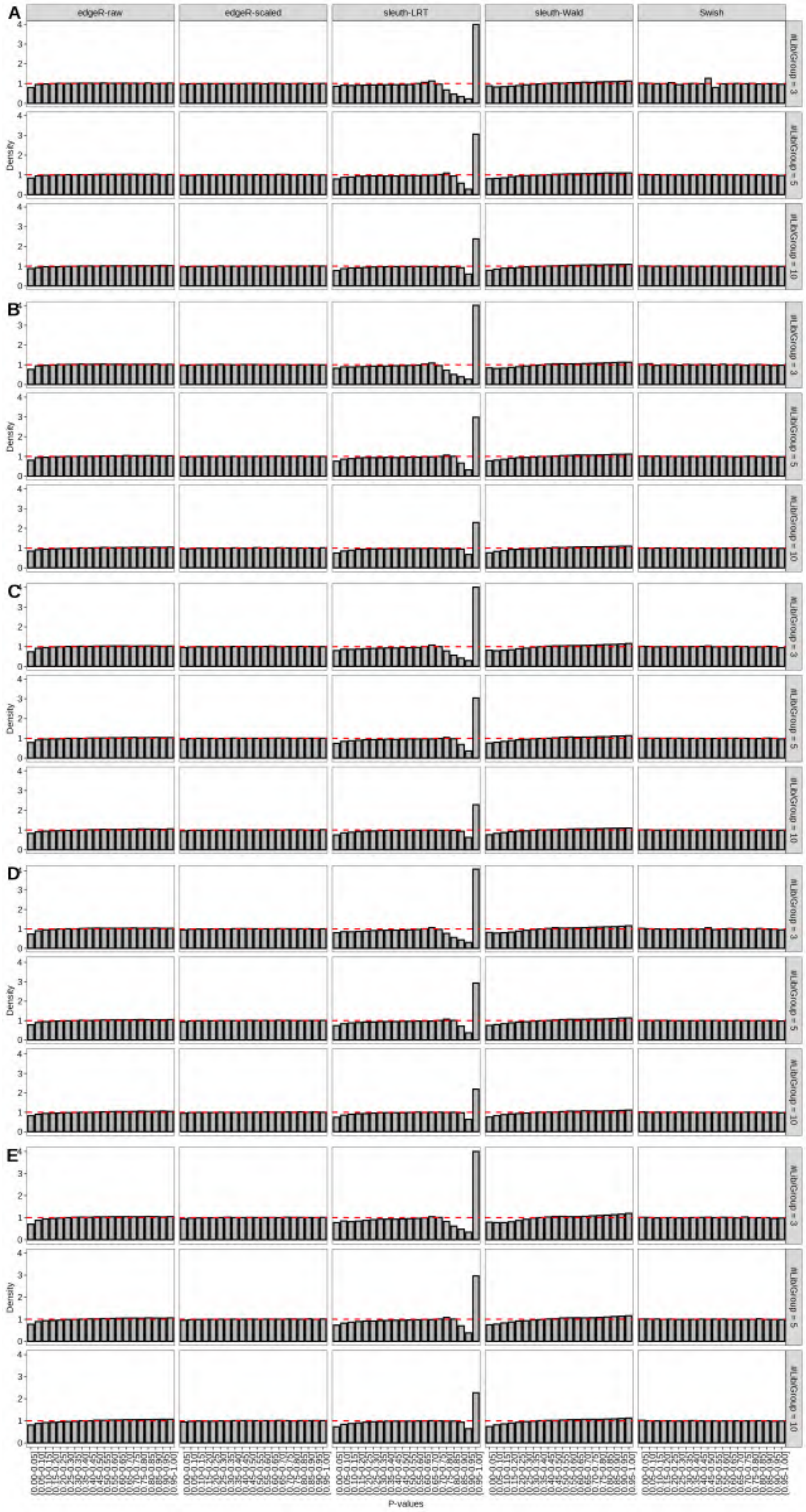

Figure S85: Simulation results. Scenario with mm39 genome, 150bp single-end reads quantified with Salmon, and balanced libraries. Density histograms of p-values from null simulations (averaged over 20 simulations). (A) Top 2 transcripts/gene expressed. (B) Top 3 transcripts/gene expressed. (C) Top 4 transcripts/gene expressed. (D) Top 5 transcripts/gene expressed. (E) All transcripts from the reference dataset are expressed.

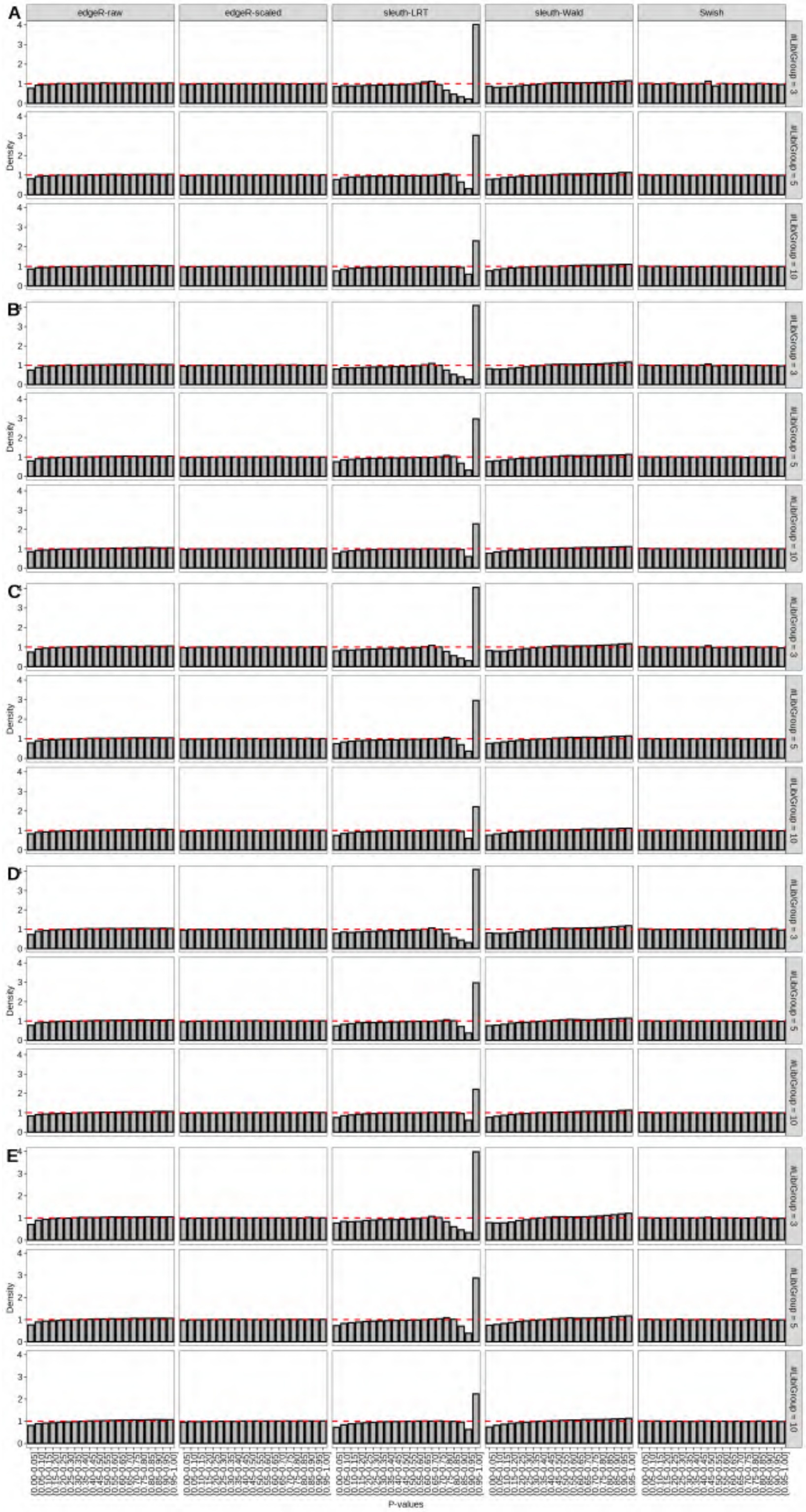

Figure S86: Simulation results. Scenario with mm39 genome, 50bp paired-end reads quantified with Salmon, and balanced libraries. Density histograms of p-values from null simulations (averaged over 20 simulations). (A) Top 2 transcripts/gene expressed. (B) Top 3 transcripts/gene expressed. (C) Top 4 transcripts/gene expressed. (D) Top 5 transcripts/gene expressed. (E) All transcripts from the reference dataset are expressed.

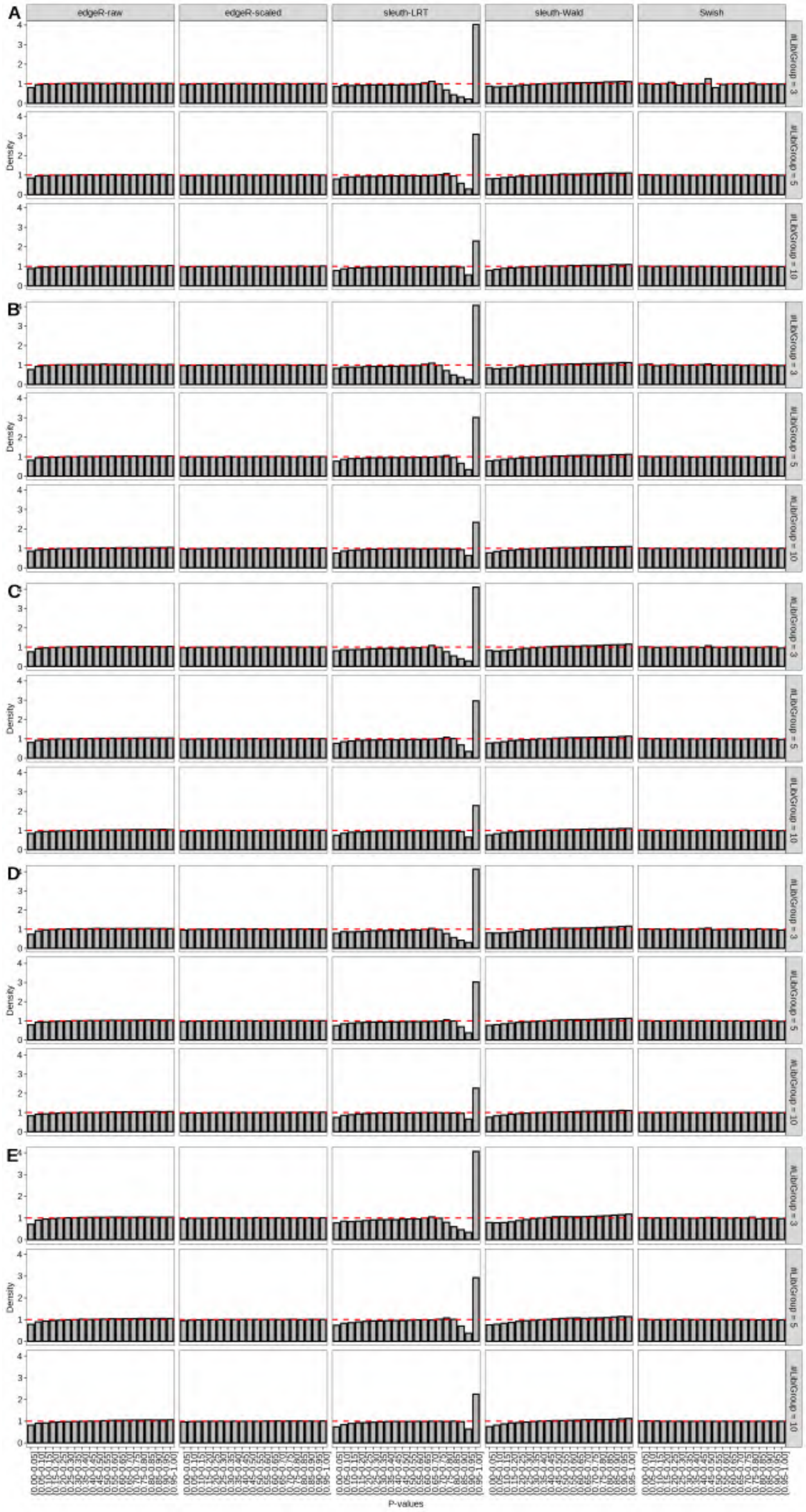

Figure S87: Simulation results. Scenario with mm39 genome, 75bp paired-end reads quantified with Salmon, and balanced libraries. Density histograms of p-values from null simulations (averaged over 20 simulations). (A) Top 2 transcripts/gene expressed. (B) Top 3 transcripts/gene expressed. (C) Top 4 transcripts/gene expressed. (D) Top 5 transcripts/gene expressed. (E) All transcripts from the reference dataset are expressed.

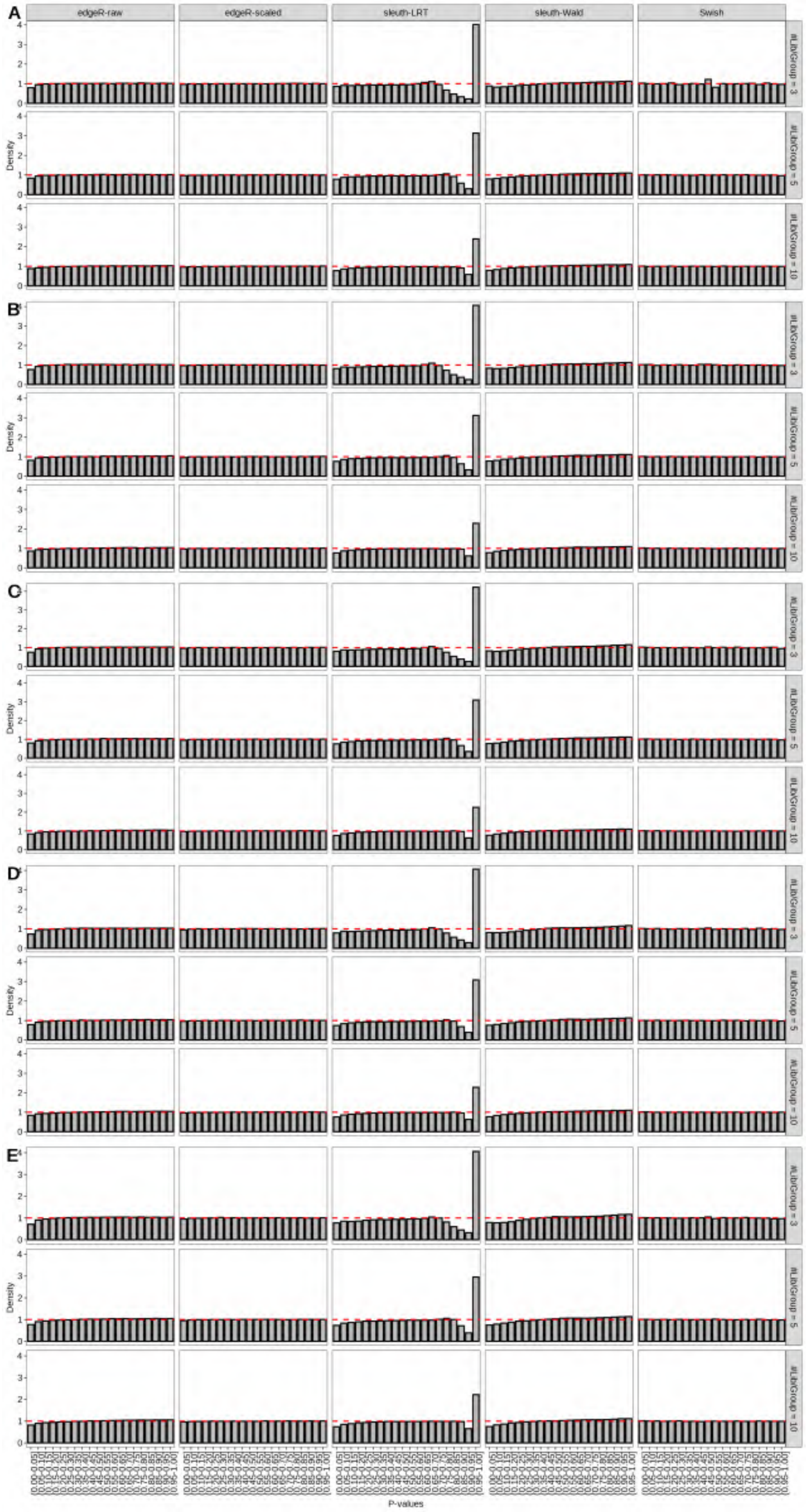

Figure S88: Simulation results. Scenario with mm39 genome, 100bp paired-end reads quantified with Salmon, and balanced libraries. Density histograms of p-values from null simulations (averaged over 20 simulations). (A) Top 2 transcripts/gene expressed. (B) Top 3 transcripts/gene expressed. (C) Top 4 transcripts/gene expressed. (D) Top 5 transcripts/gene expressed. (E) All transcripts from the reference dataset are expressed.

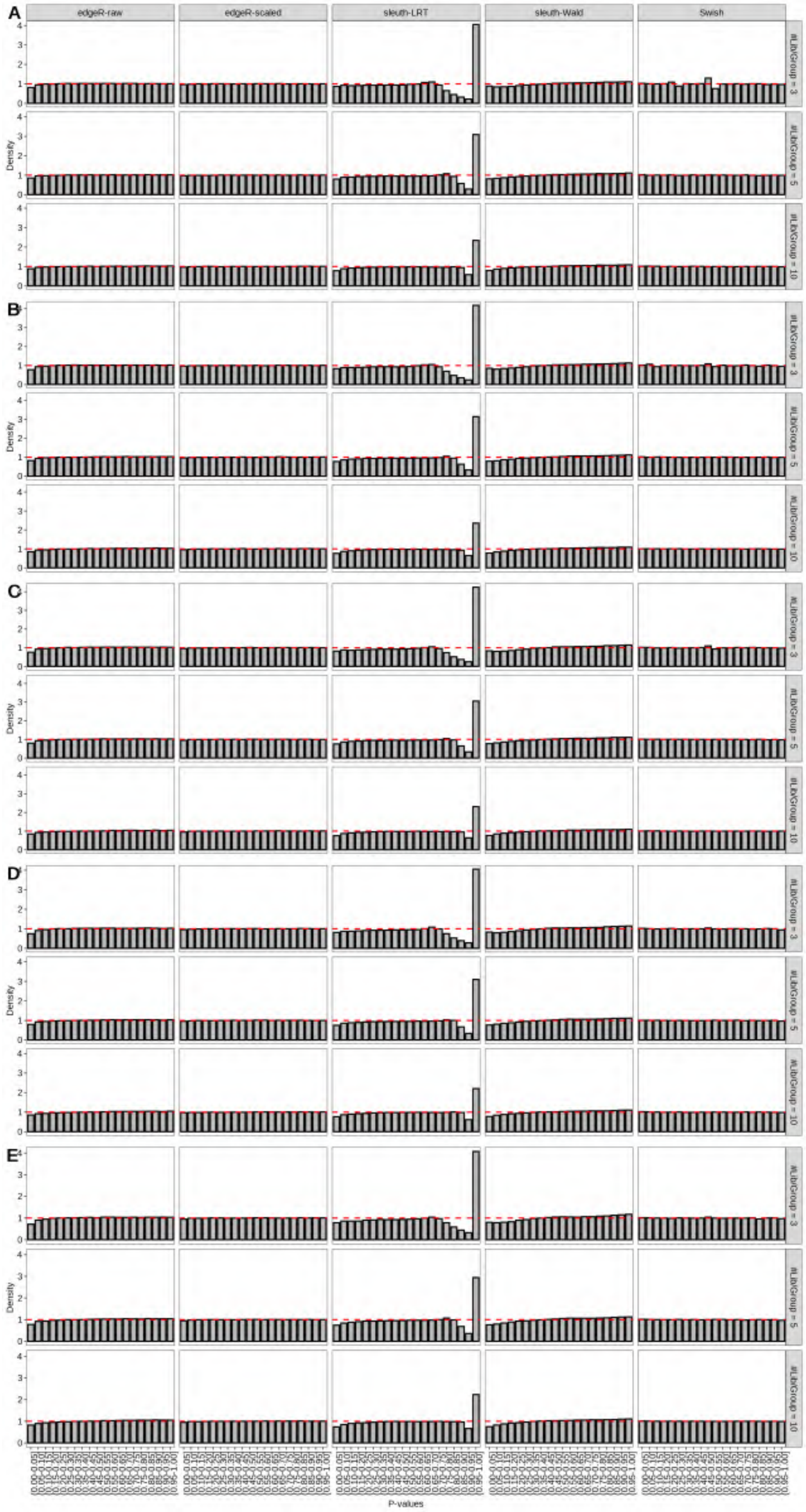

Figure S89: Simulation results. Scenario with mm39 genome, 125bp paired-end reads quantified with Salmon, and balanced libraries. Density histograms of p-values from null simulations (averaged over 20 simulations). (A) Top 2 transcripts/gene expressed. (B) Top 3 transcripts/gene expressed. (C) Top 4 transcripts/gene expressed. (D) Top 5 transcripts/gene expressed. (E) All transcripts from the reference dataset are expressed.

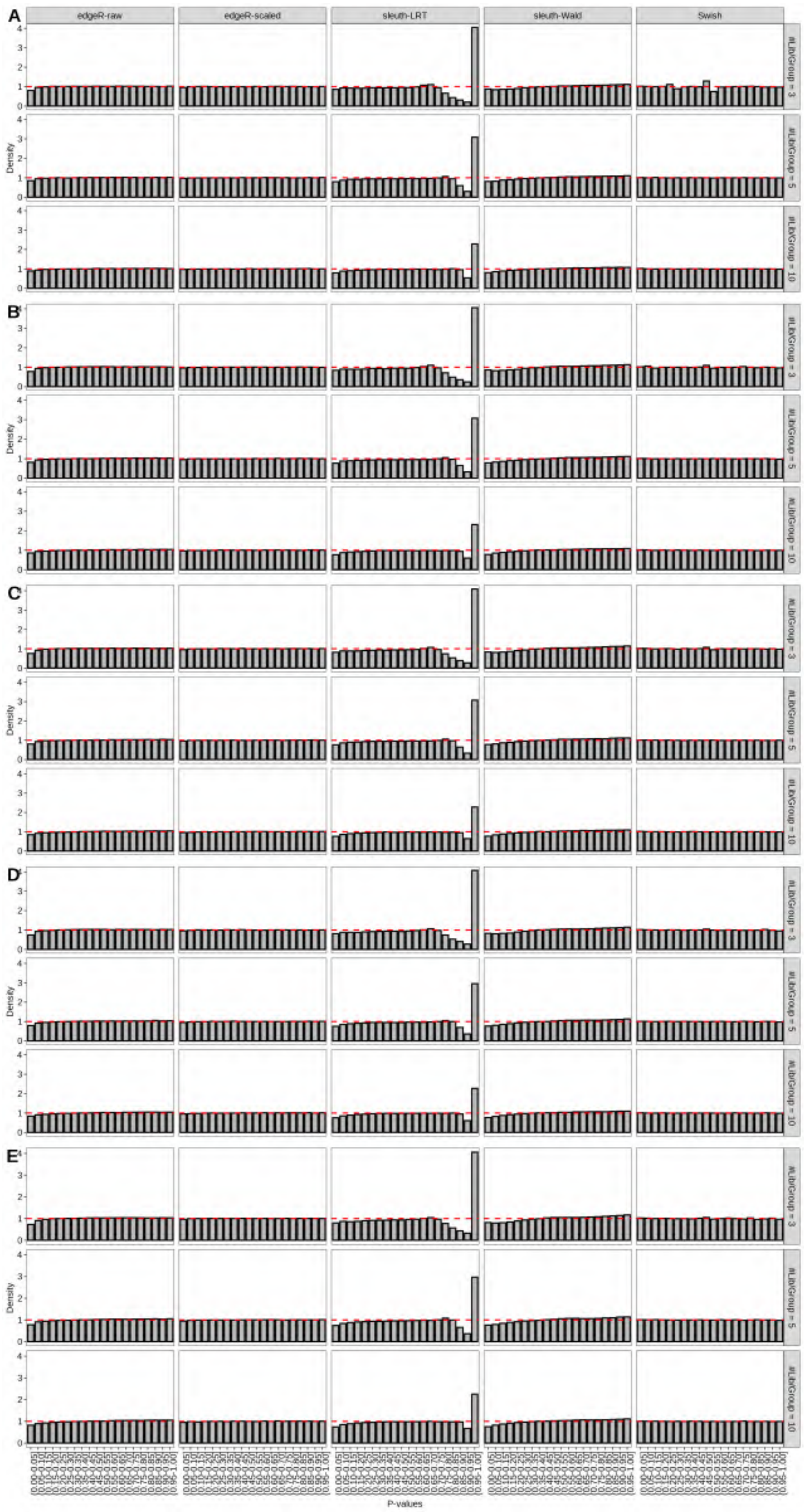

Figure S90: Simulation results. Scenario with mm39 genome, 150bp paired-end reads quantified with Salmon, and balanced libraries. Density histograms of p-values from null simulations (averaged over 20 simulations). (A) Top 2 transcripts/gene expressed. (B) Top 3 transcripts/gene expressed. (C) Top 4 transcripts/gene expressed. (D) Top 5 transcripts/gene expressed. (E) All transcripts from the reference dataset are expressed.

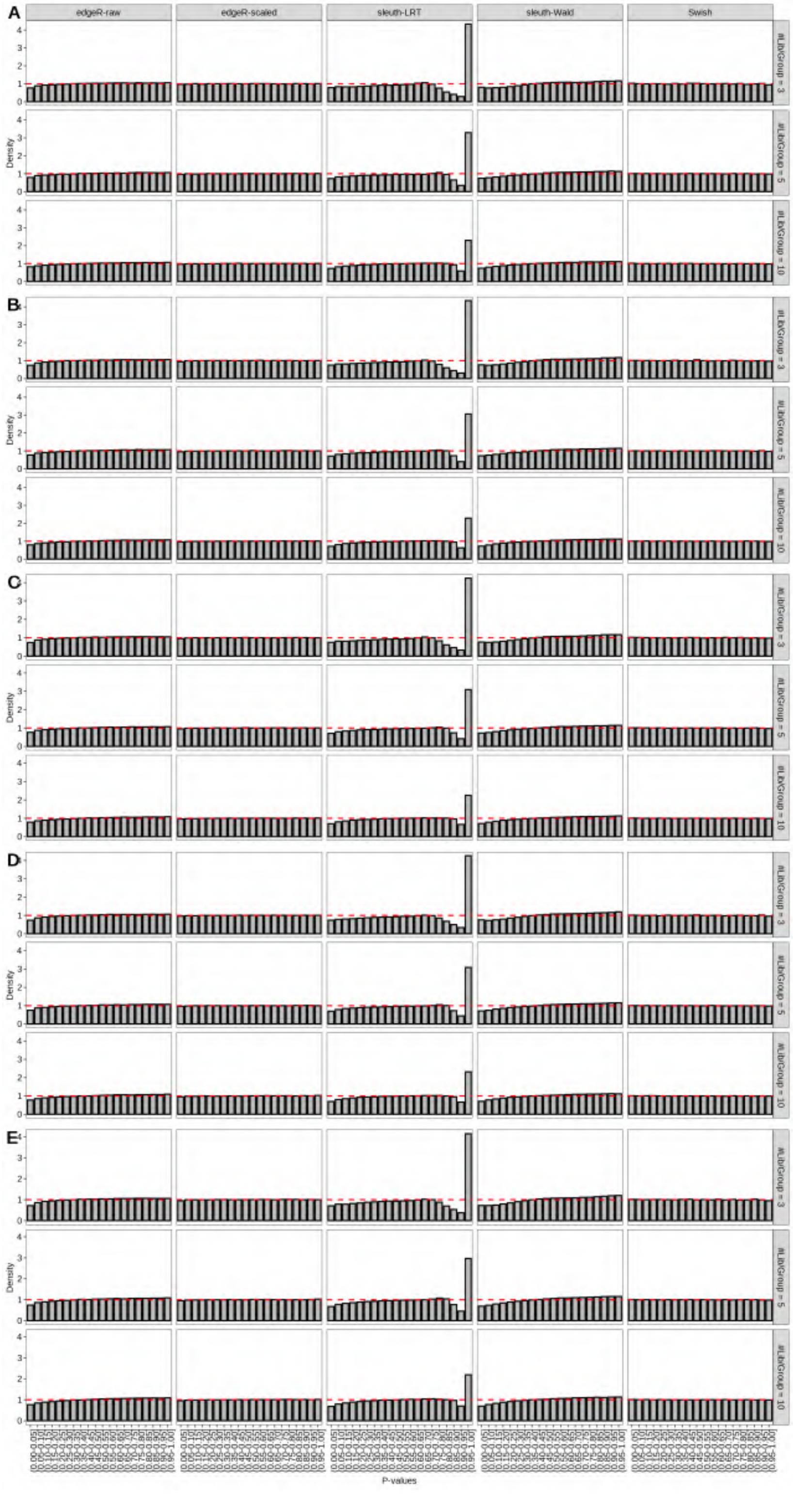

Figure S91: Simulation results. Scenario with mm39 genome, 50bp single-end reads quantified with kallisto, and balanced libraries. Density histograms of p-values from null simulations (averaged over 20 simulations). (A) Top 2 transcripts/gene expressed. (B) Top 3 transcripts/gene expressed. (C) Top 4 transcripts/gene expressed. (D) Top 5 transcripts/gene expressed. (E) All transcripts from the reference dataset are expressed.

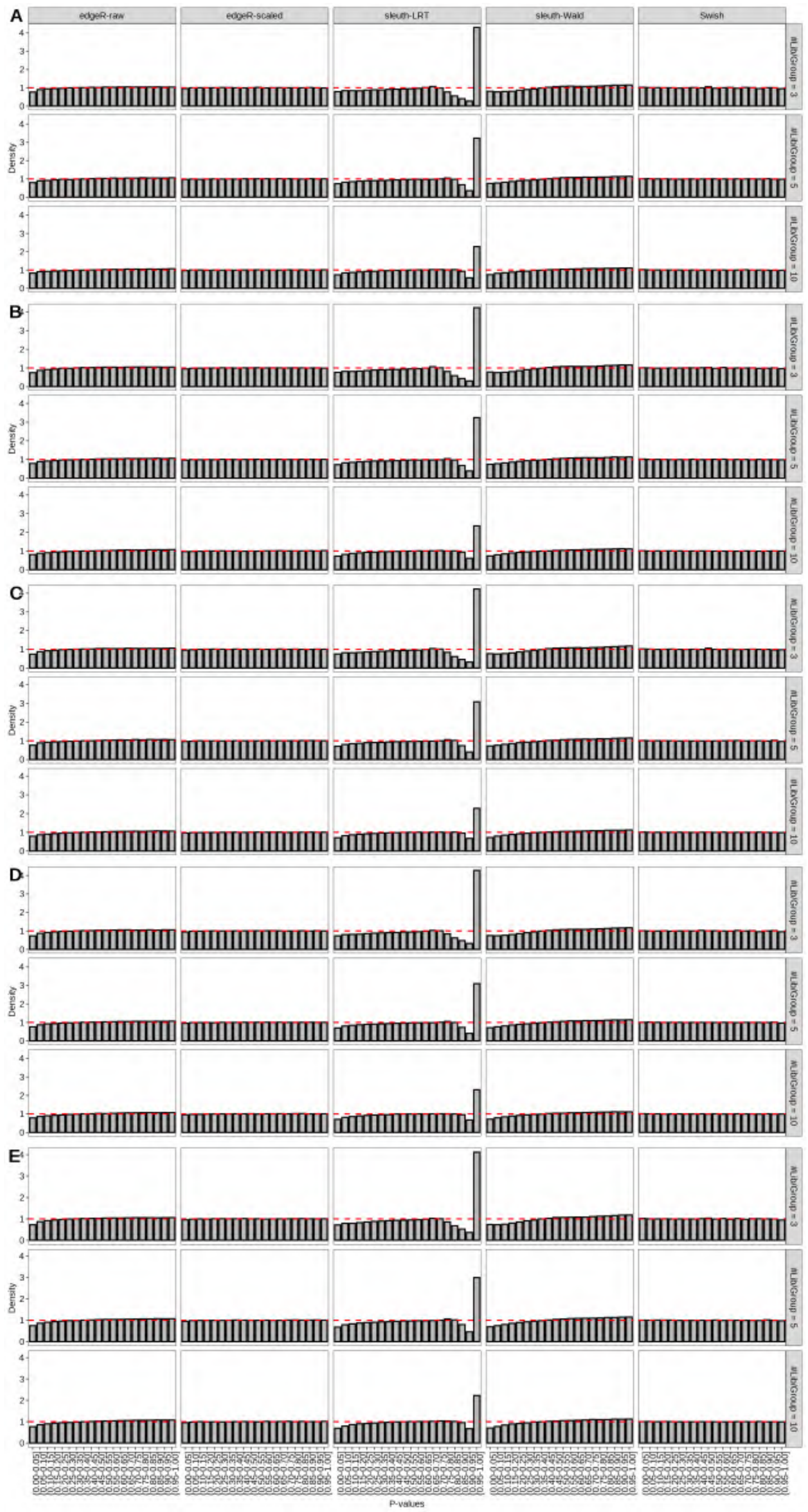

Figure S92: Simulation results. Scenario with mm39 genome, 75bp single-end reads quantified with kallisto, and balanced libraries. Density histograms of p-values from null simulations (averaged over 20 simulations). (A) Top 2 transcripts/gene expressed. (B) Top 3 transcripts/gene expressed. (C) Top 4 transcripts/gene expressed. (D) Top 5 transcripts/gene expressed. (E) All transcripts from the reference dataset are expressed.

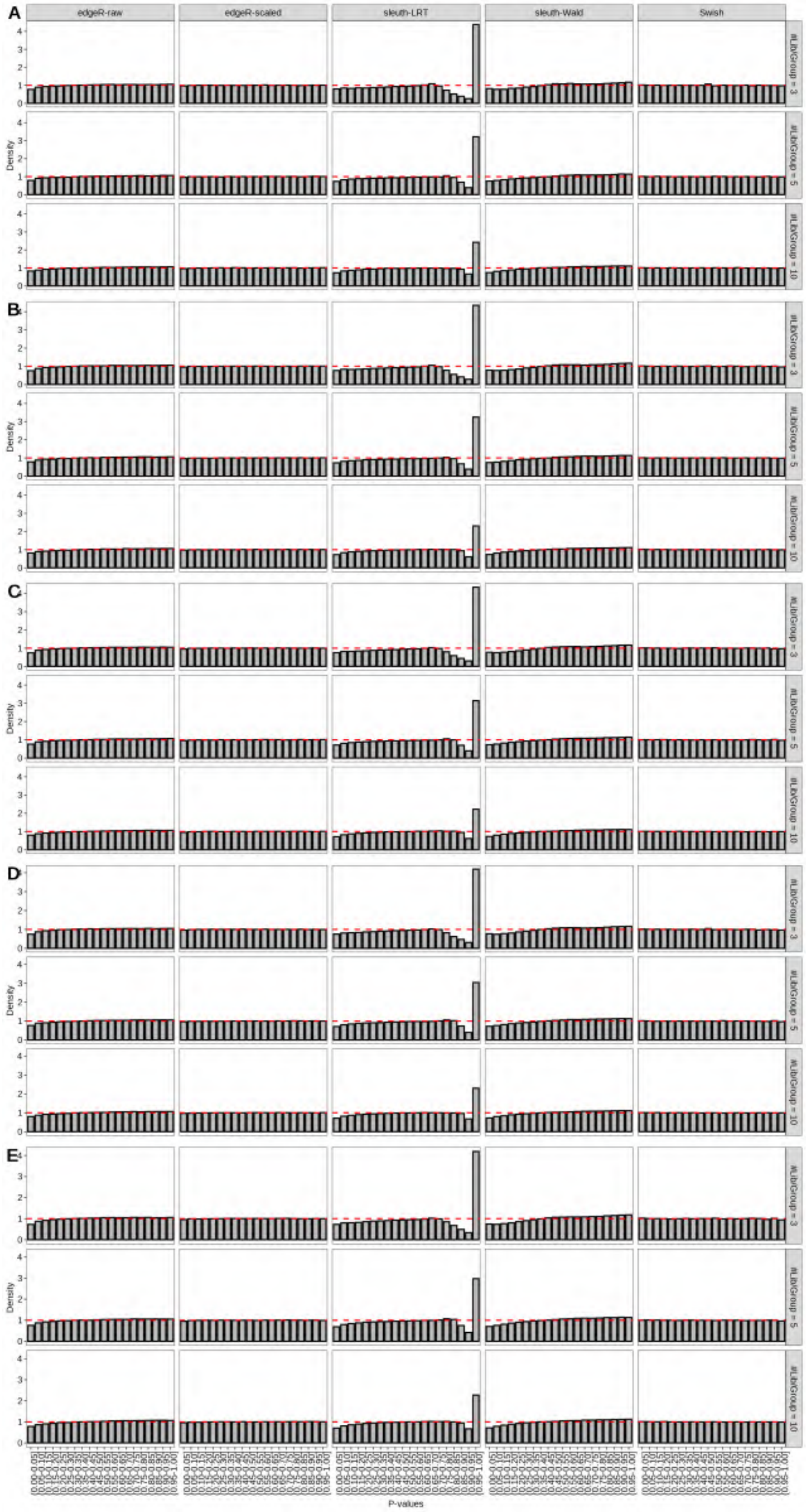

Figure S93: Simulation results. Scenario with mm39 genome, 100bp single-end reads quantified with kallisto, and balanced libraries. Density histograms of p-values from null simulations (averaged over 20 simulations). (A) Top 2 transcripts/gene expressed. (B) Top 3 transcripts/gene expressed. (C) Top 4 transcripts/gene expressed. (D) Top 5 transcripts/gene expressed. (E) All transcripts from the reference dataset are expressed.

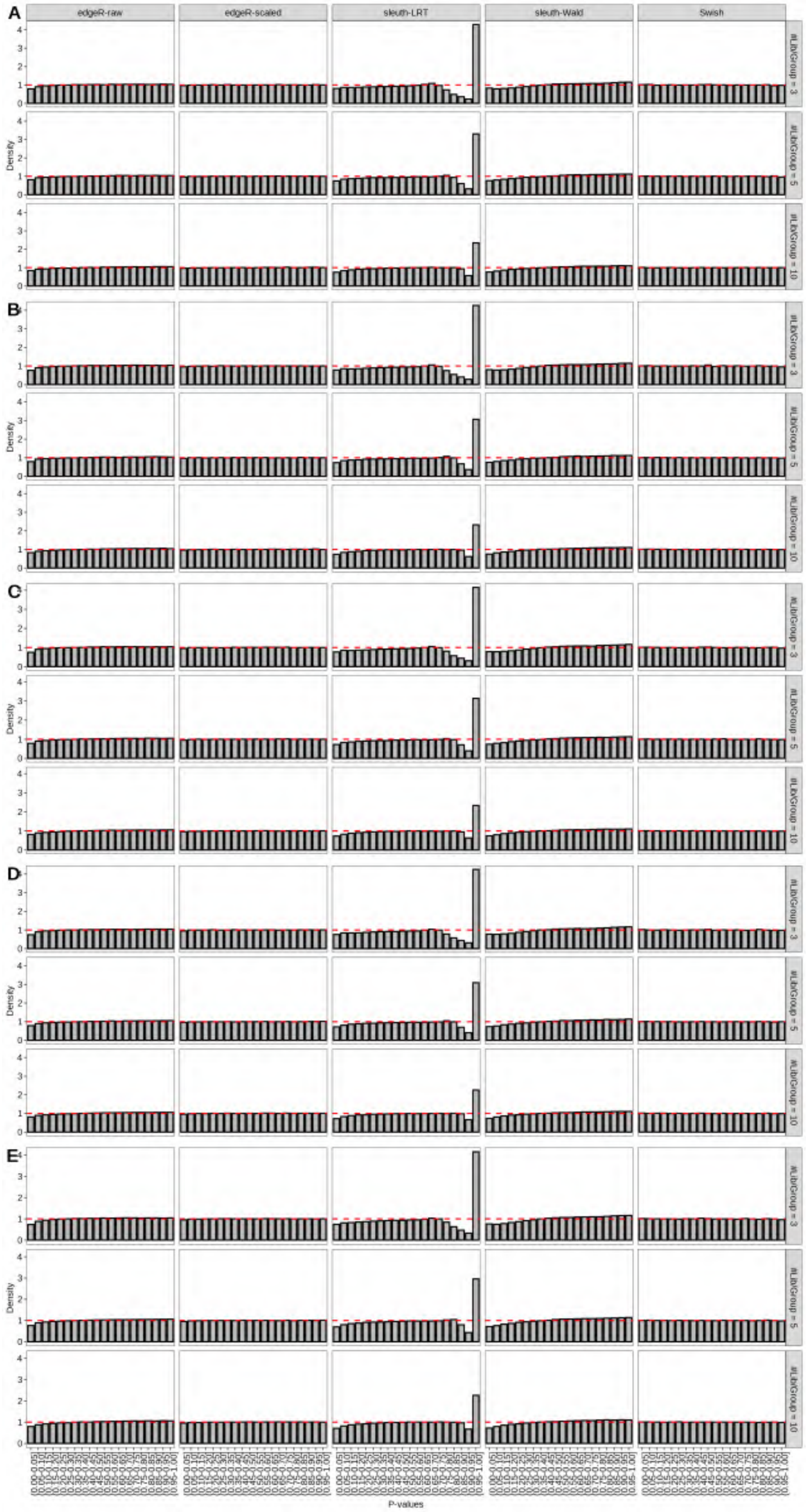

Figure S94: Simulation results. Scenario with mm39 genome, 125bp single-end reads quantified with kallisto, and balanced libraries. Density histograms of p-values from null simulations (averaged over 20 simulations). (A) Top 2 transcripts/gene expressed. (B) Top 3 transcripts/gene expressed. (C) Top 4 transcripts/gene expressed. (D) Top 5 transcripts/gene expressed. (E) All transcripts from the reference dataset are expressed.

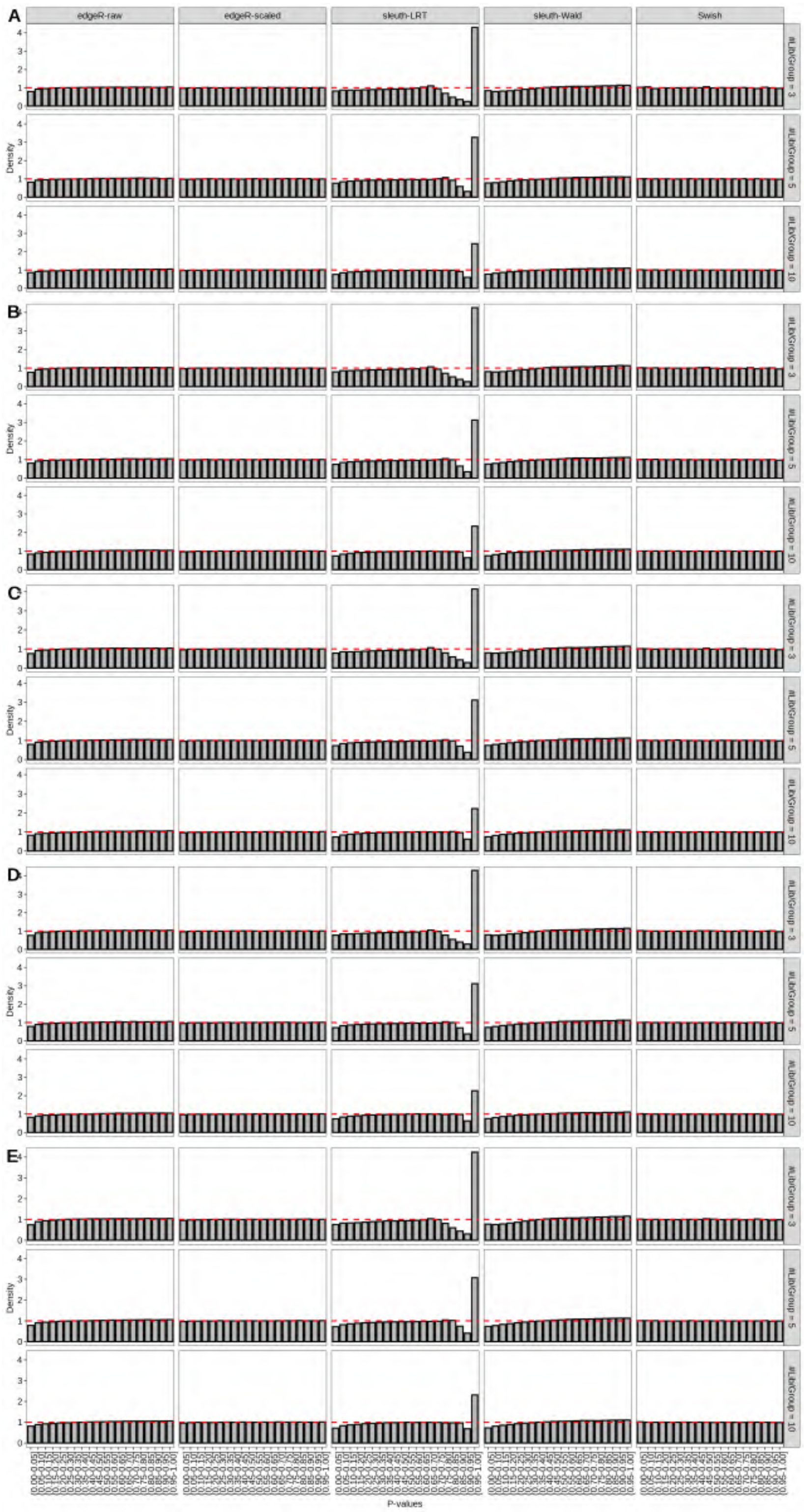

Figure S95: Simulation results. Scenario with mm39 genome, 150bp single-end reads quantified with kallisto, and balanced libraries. Density histograms of p-values from null simulations (averaged over 20 simulations). (A) Top 2 transcripts/gene expressed. (B) Top 3 transcripts/gene expressed. (C) Top 4 transcripts/gene expressed. (D) Top 5 transcripts/gene expressed. (E) All transcripts from the reference dataset are expressed.

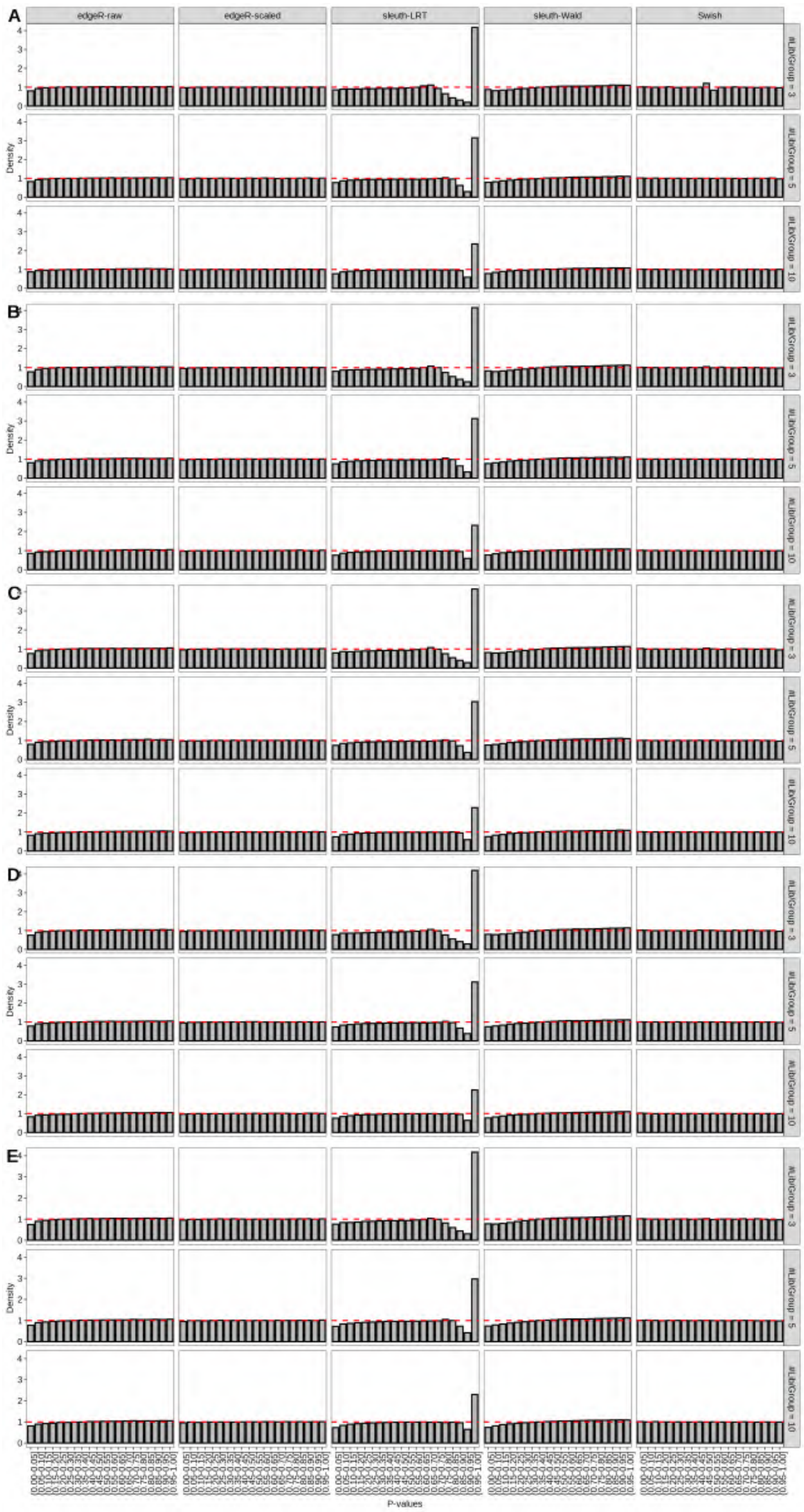

Figure S96: Simulation results. Scenario with mm39 genome, 50bp paired-end reads quantified with kallisto, and balanced libraries. Density histograms of p-values from null simulations (averaged over 20 simulations). (A) Top 2 transcripts/gene expressed. (B) Top 3 transcripts/gene expressed. (C) Top 4 transcripts/gene expressed. (D) Top 5 transcripts/gene expressed. (E) All transcripts from the reference dataset are expressed.

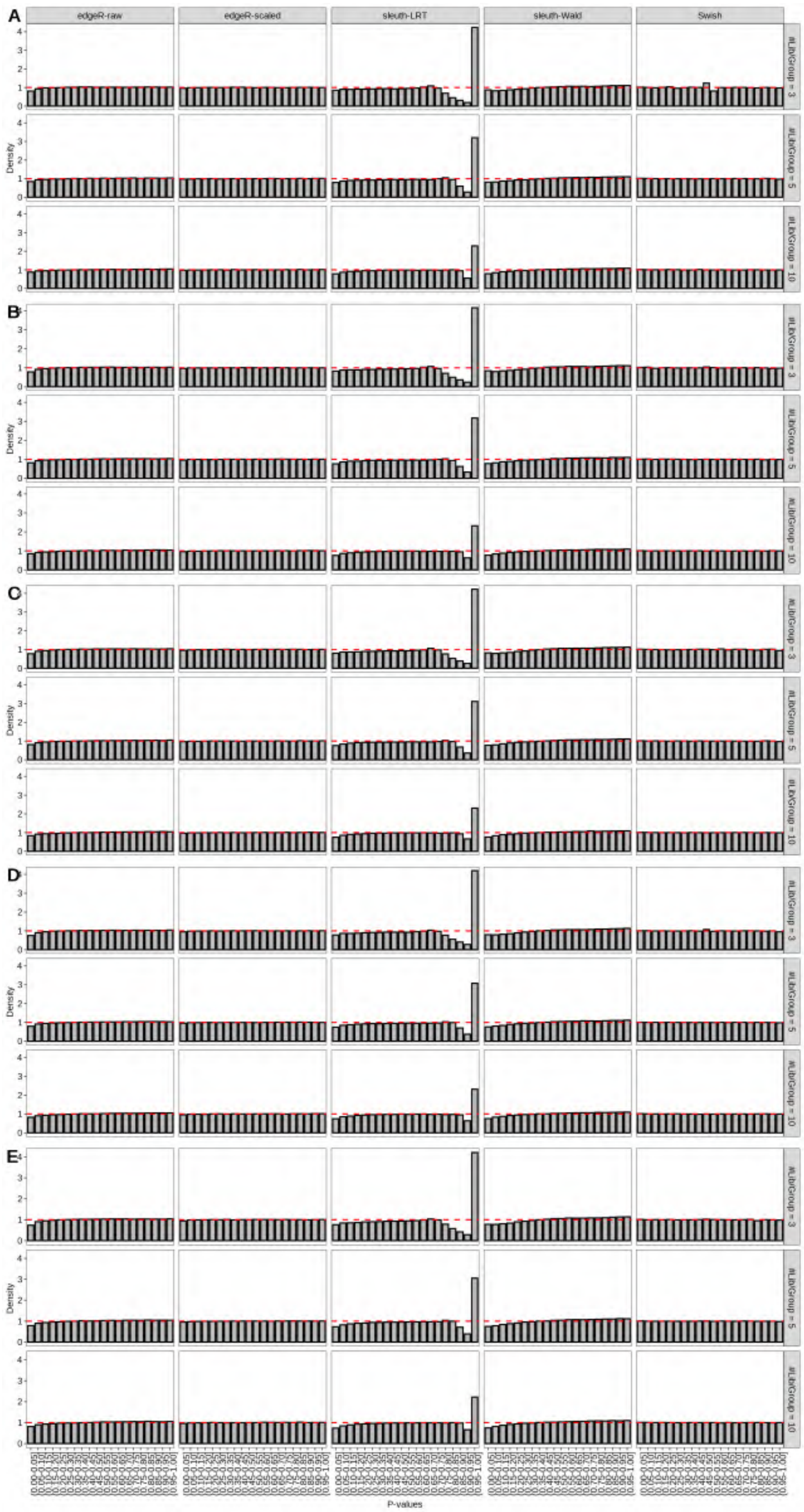

Figure S97: Simulation results. Scenario with mm39 genome, 75bp paired-end reads quantified with kallisto, and balanced libraries. Density histograms of p-values from null simulations (averaged over 20 simulations). (A) Top 2 transcripts/gene expressed. (B) Top 3 transcripts/gene expressed. (C) Top 4 transcripts/gene expressed. (D) Top 5 transcripts/gene expressed. (E) All transcripts from the reference dataset are expressed.

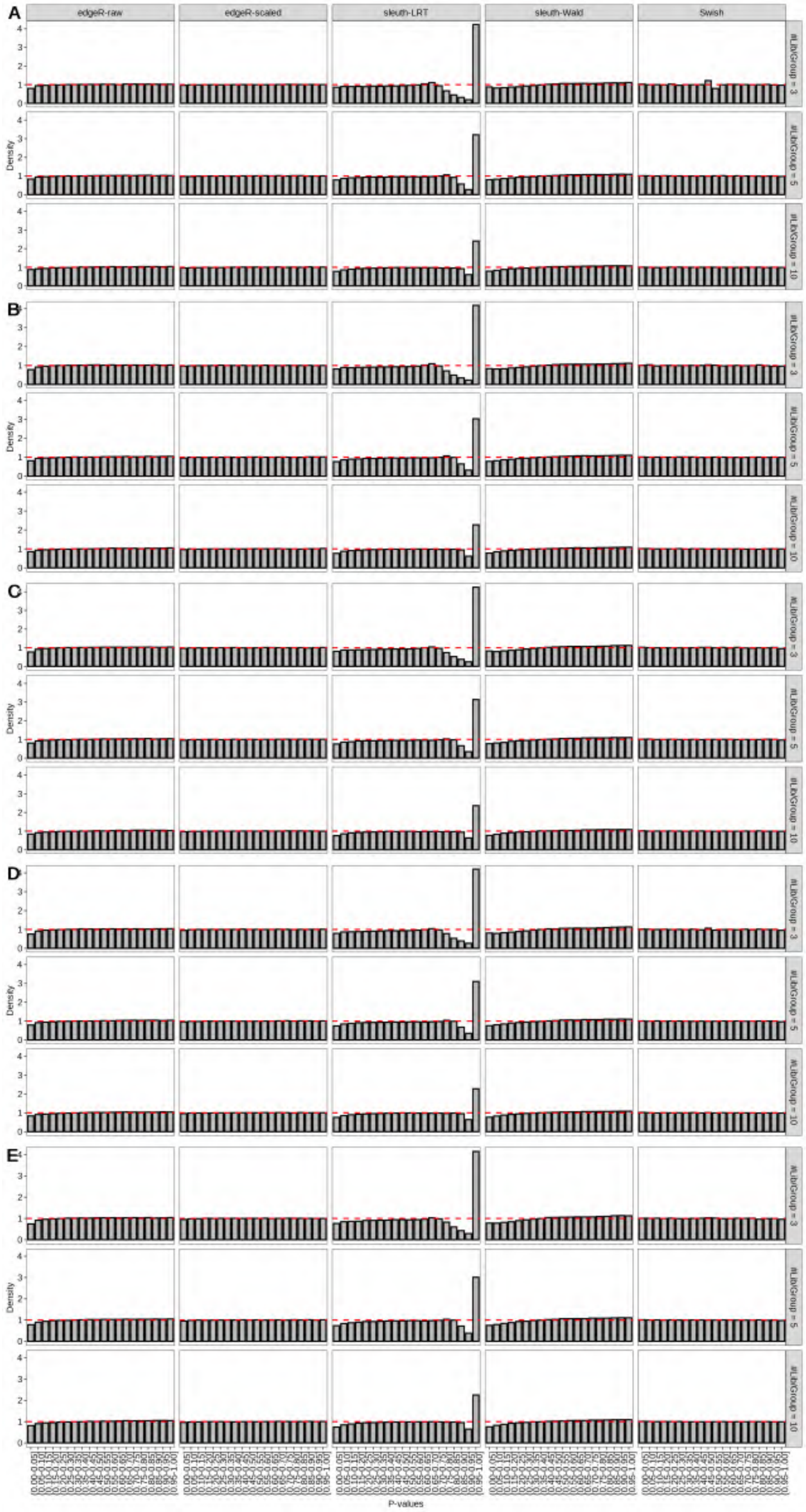

Figure S98: Simulation results. Scenario with mm39 genome, 100bp paired-end reads quantified with kallisto, and balanced libraries. Density histograms of p-values from null simulations (averaged over 20 simulations). (A) Top 2 transcripts/gene expressed. (B) Top 3 transcripts/gene expressed. (C) Top 4 transcripts/gene expressed. (D) Top 5 transcripts/gene expressed. (E) All transcripts from the reference dataset are expressed.

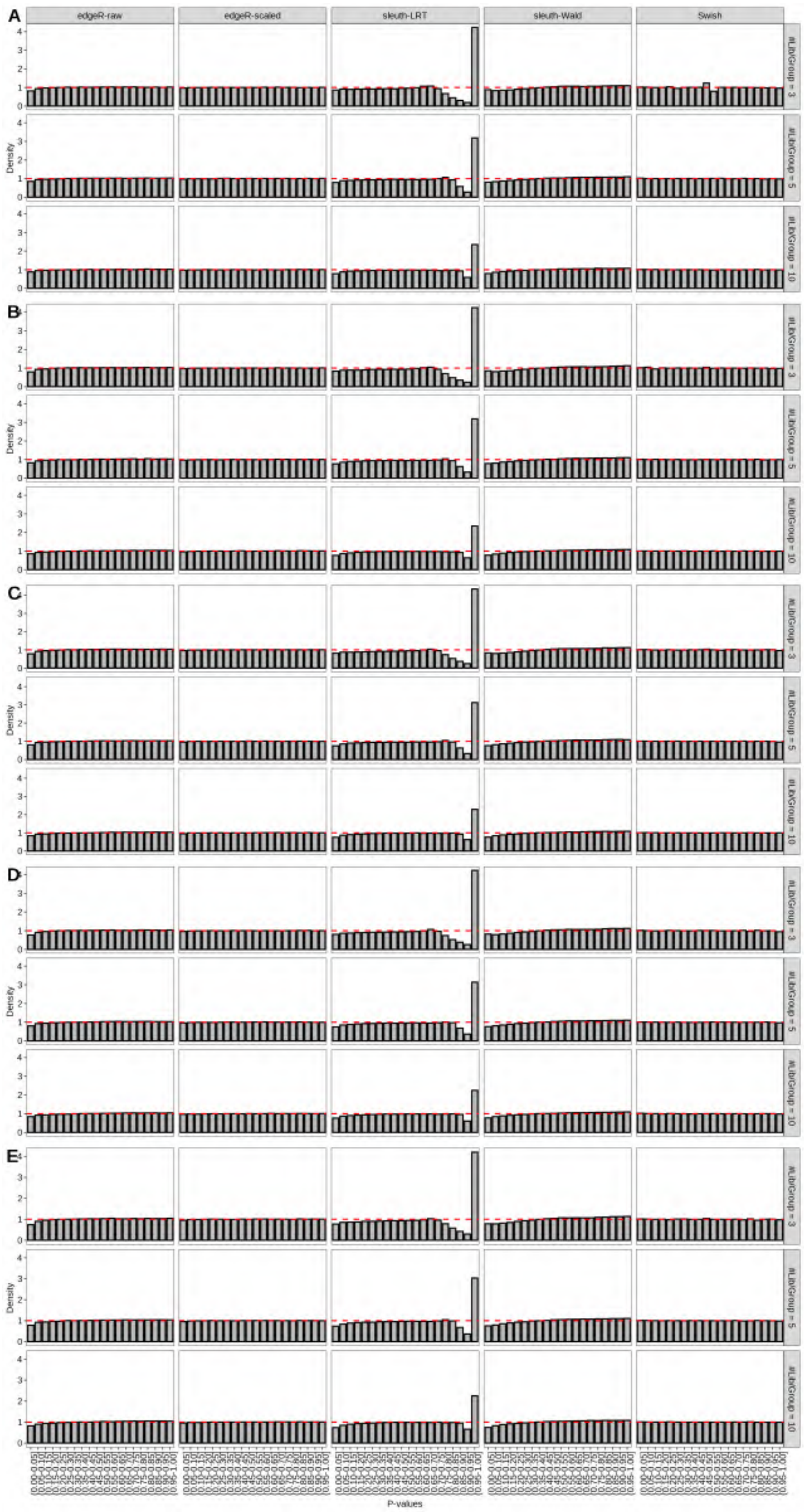

Figure S99: Simulation results. Scenario with mm39 genome, 125bp paired-end reads quantified with kallisto, and balanced libraries. Density histograms of p-values from null simulations (averaged over 20 simulations). (A) Top 2 transcripts/gene expressed. (B) Top 3 transcripts/gene expressed. (C) Top 4 transcripts/gene expressed. (D) Top 5 transcripts/gene expressed. (E) All transcripts from the reference dataset are expressed.

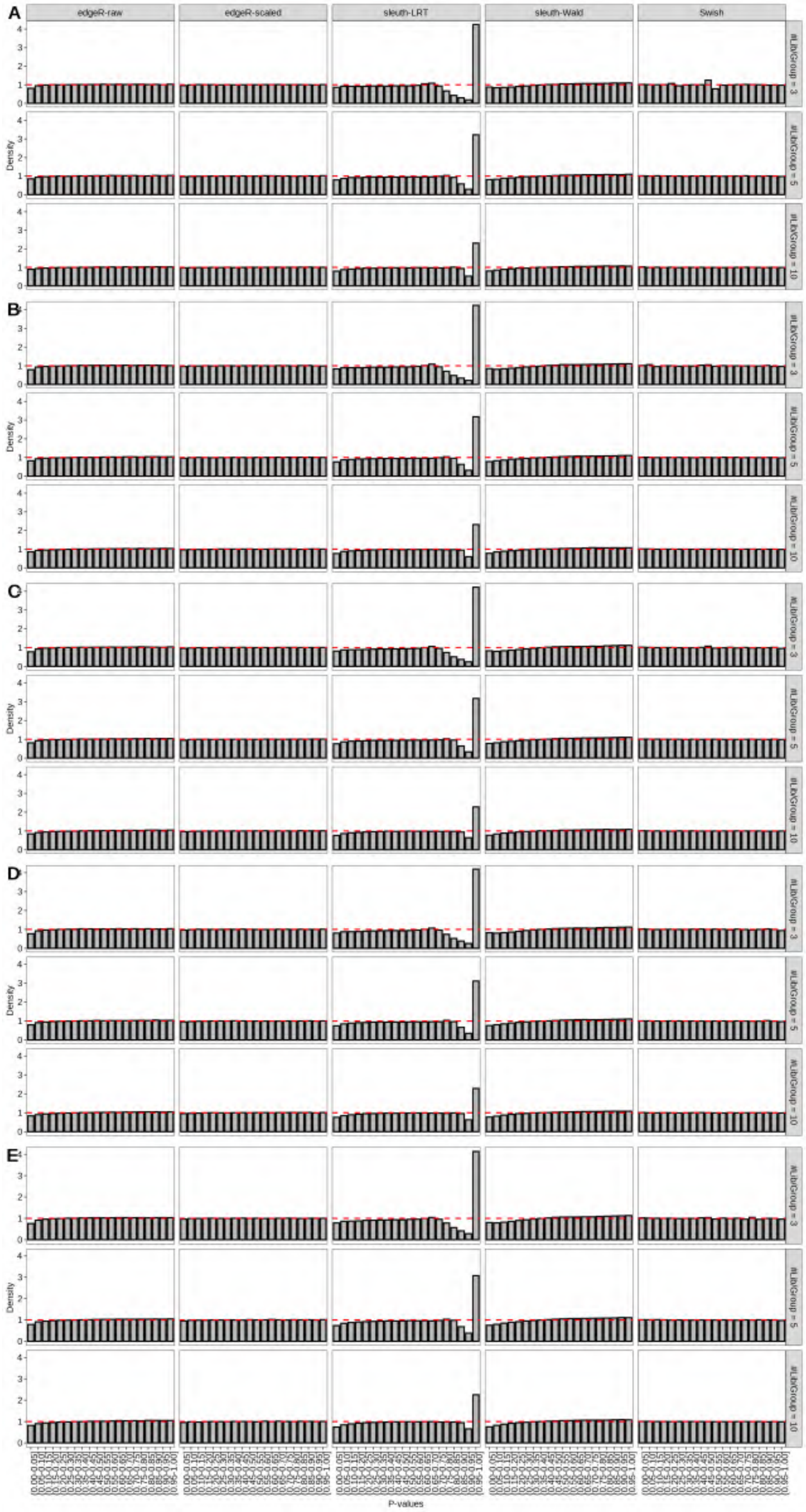

Figure S100: Simulation results. Scenario with mm39 genome, 150bp paired-end reads quantified with kallisto, and balanced libraries. Density histograms of p-values from null simulations (averaged over 20 simulations). (A) Top 2 transcripts/gene expressed. (B) Top 3 transcripts/gene expressed. (C) Top 4 transcripts/gene expressed. (D) Top 5 transcripts/gene expressed. (E) All transcripts from the reference dataset are expressed.

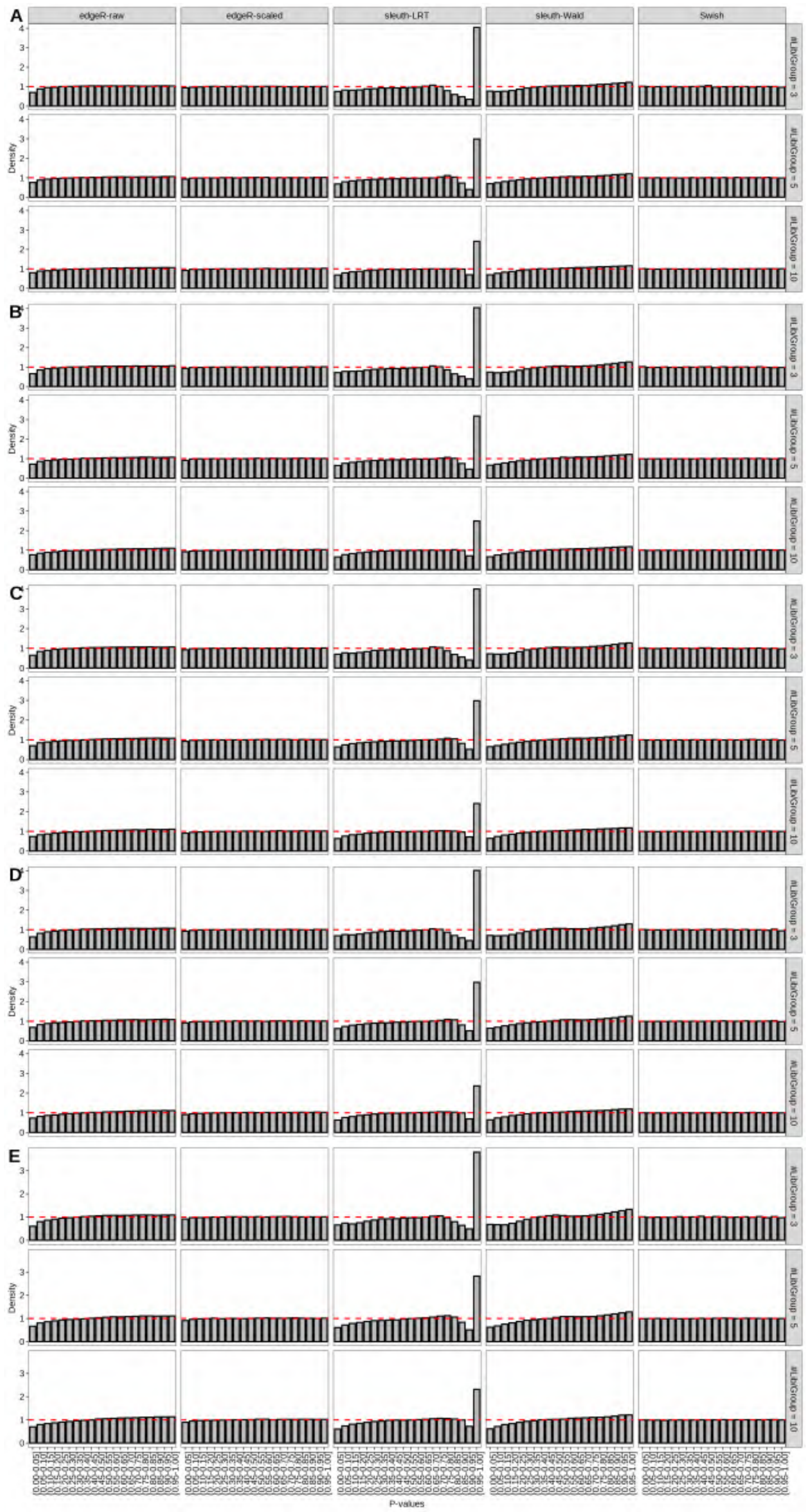

Figure S101: Simulation results. Scenario with mm39 genome, 50bp single-end reads quantified with Salmon, and unbalanced libraries. Density histograms of p-values from null simulations (averaged over 20 simulations). (A) Top 2 transcripts/gene expressed. (B) Top 3 transcripts/gene expressed. (C) Top 4 transcripts/gene expressed. (D) Top 5 transcripts/gene expressed. (E) All transcripts from the reference dataset are expressed.

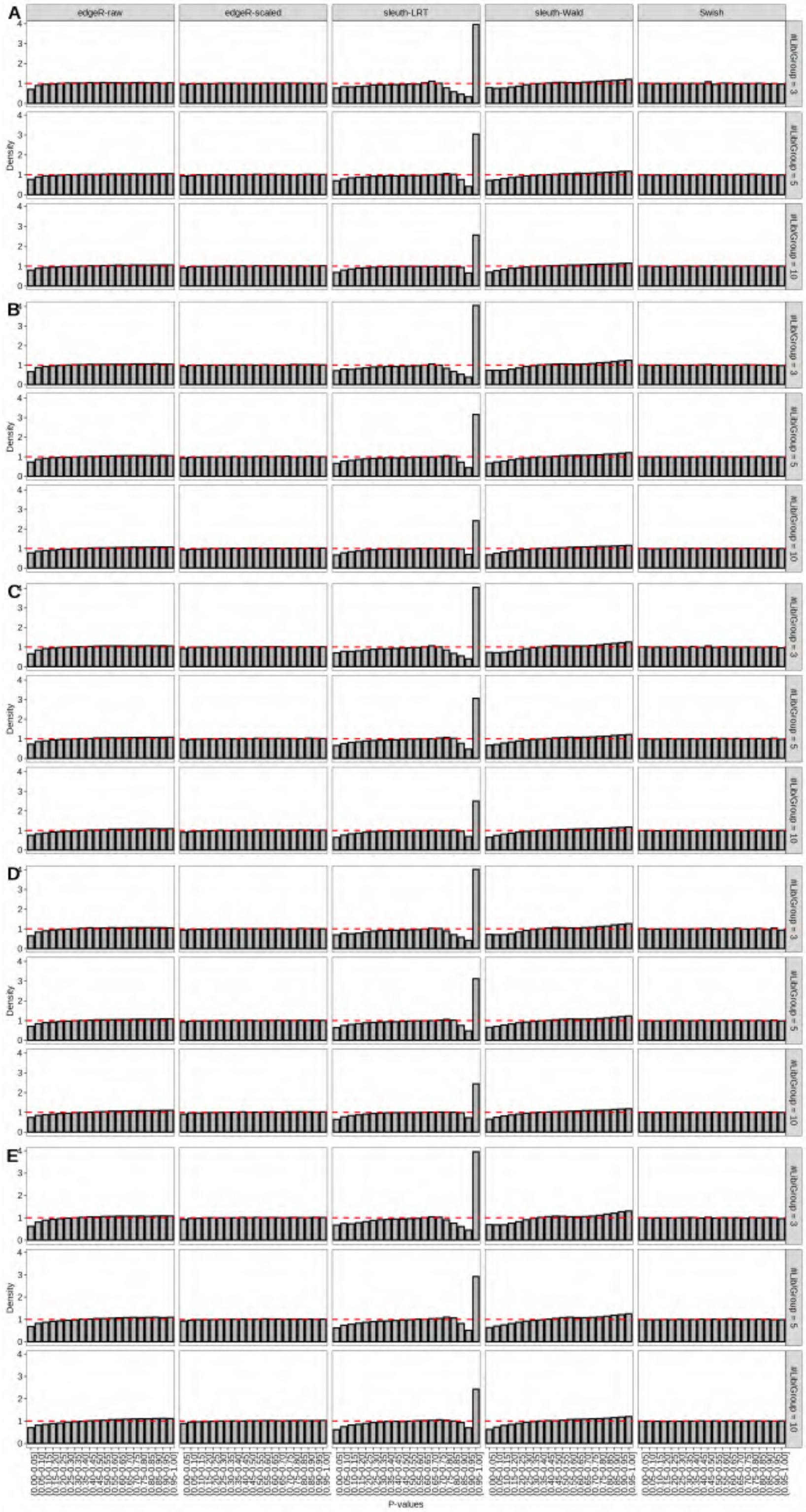

Figure S102: Simulation results. Scenario with mm39 genome, 75bp single-end reads quantified with Salmon, and unbalanced libraries. Density histograms of p-values from null simulations (averaged over 20 simulations). (A) Top 2 transcripts/gene expressed. (B) Top 3 transcripts/gene expressed. (C) Top 4 transcripts/gene expressed. (D) Top 5 transcripts/gene expressed. (E) All transcripts from the reference dataset are expressed.

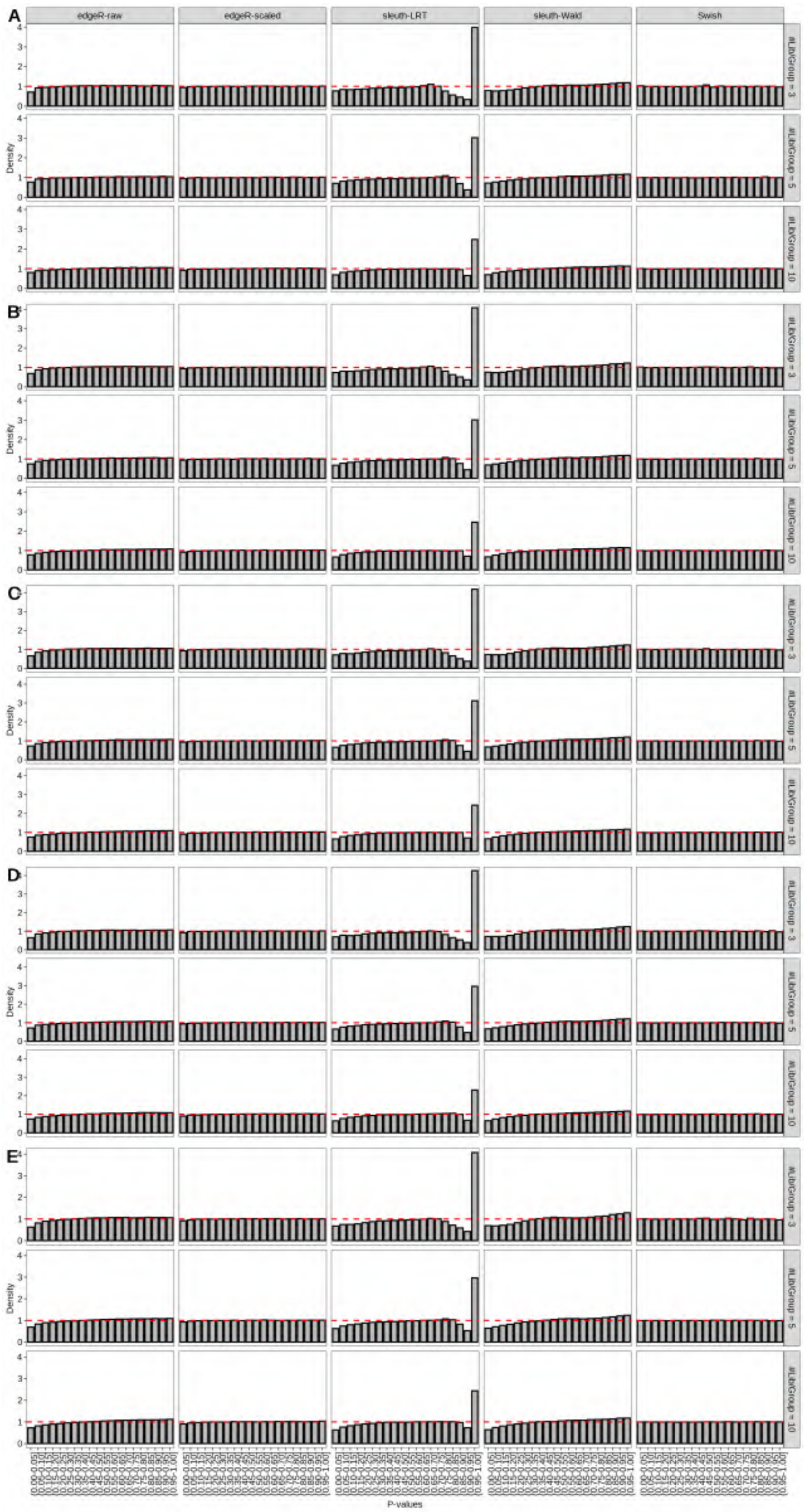

Figure S103: Simulation results. Scenario with mm39 genome, 100bp single-end reads quantified with Salmon, and unbalanced libraries. Density histograms of p-values from null simulations (averaged over 20 simulations). (A) Top 2 transcripts/gene expressed. (B) Top 3 transcripts/gene expressed. (C) Top 4 transcripts/gene expressed. (D) Top 5 transcripts/gene expressed. (E) All transcripts from the reference dataset are expressed.

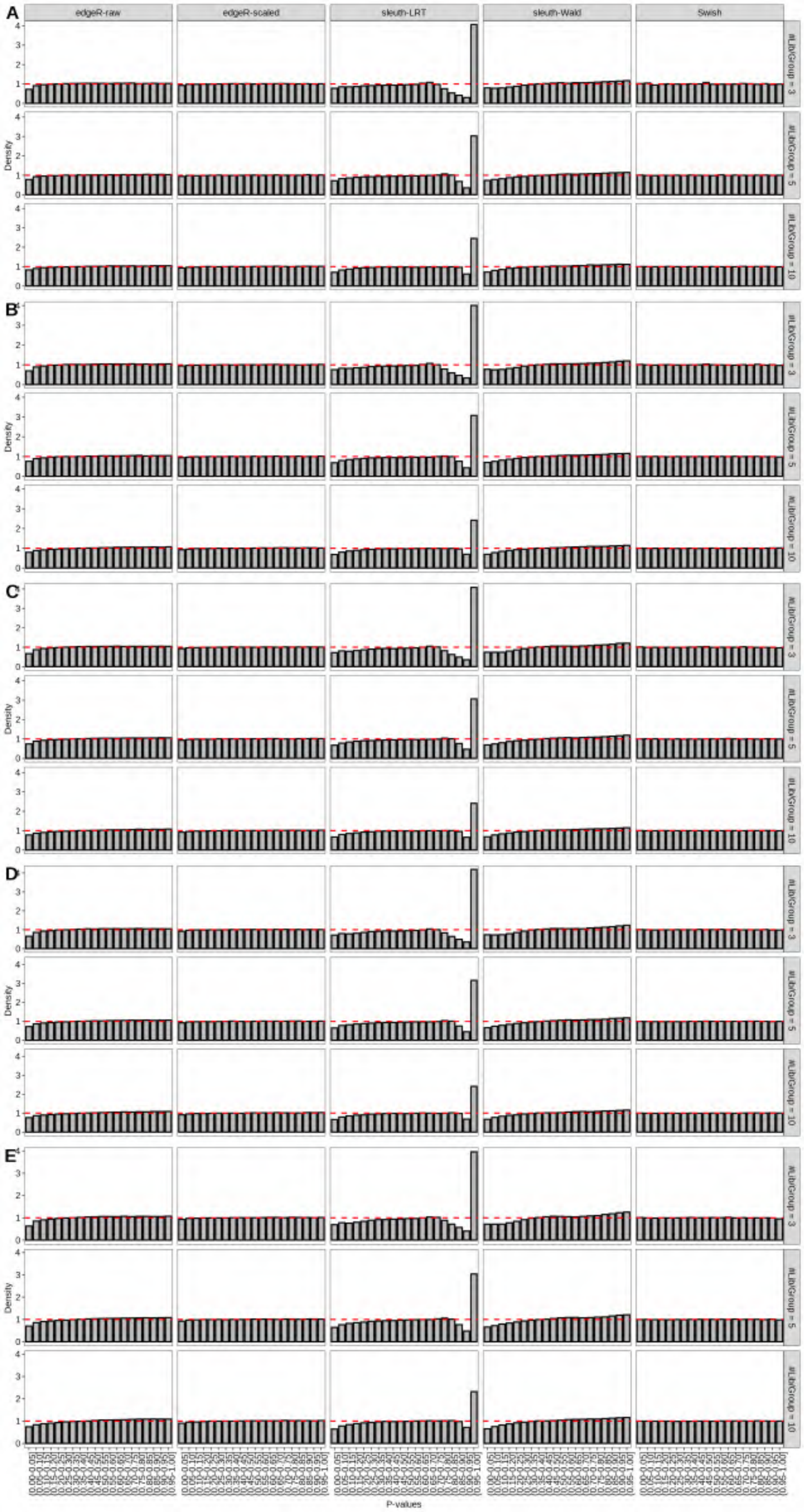

Figure S104: Simulation results. Scenario with mm39 genome, 125bp single-end reads quantified with Salmon, and unbalanced libraries. Density histograms of p-values from null simulations (averaged over 20 simulations). (A) Top 2 transcripts/gene expressed. (B) Top 3 transcripts/gene expressed. (C) Top 4 transcripts/gene expressed. (D) Top 5 transcripts/gene expressed. (E) All transcripts from the reference dataset are expressed.

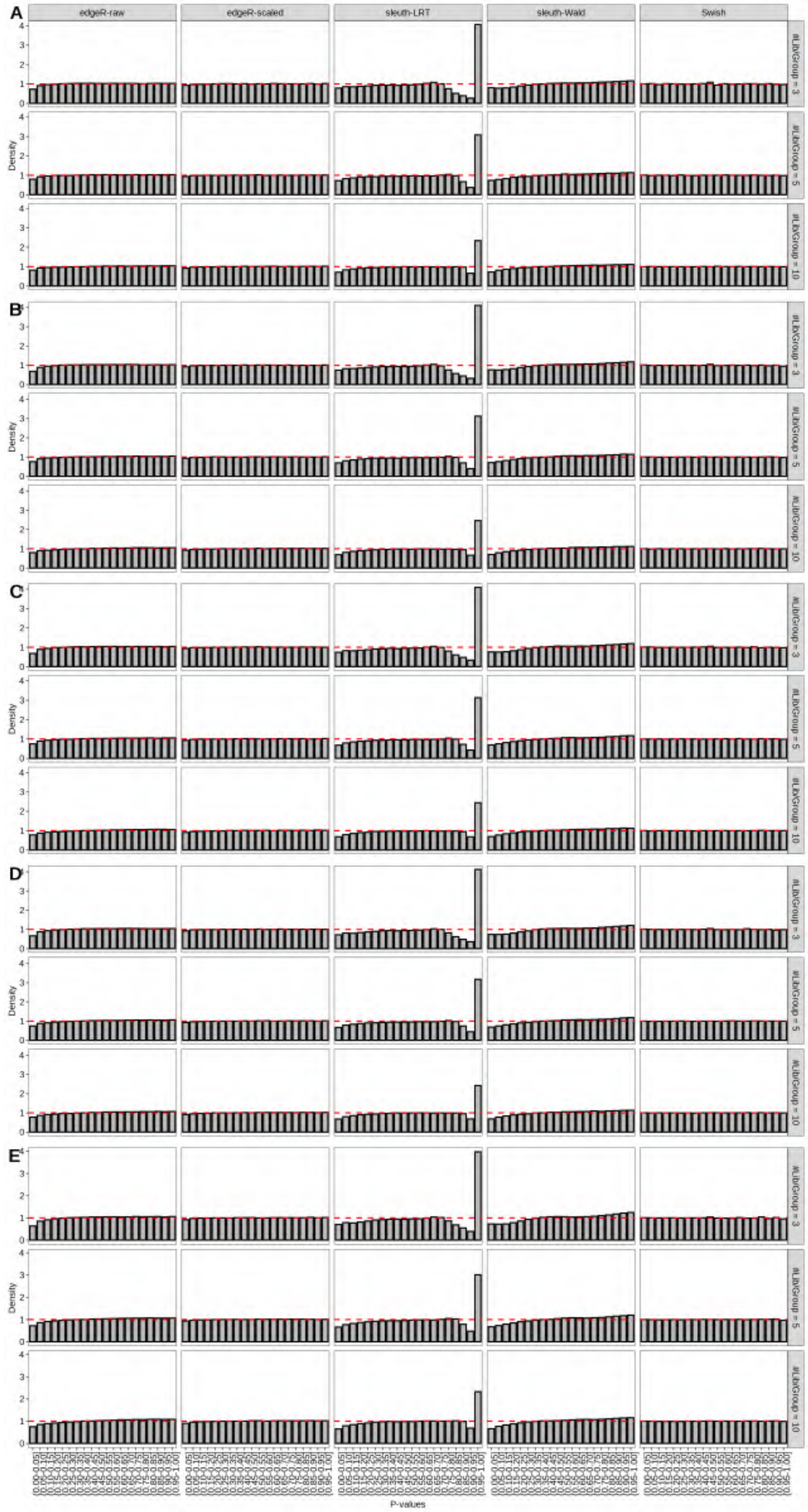

Figure S105: Simulation results. Scenario with mm39 genome, 150bp single-end reads quantified with Salmon, and unbalanced libraries. Density histograms of p-values from null simulations (averaged over 20 simulations). (A) Top 2 transcripts/gene expressed. (B) Top 3 transcripts/gene expressed. (C) Top 4 transcripts/gene expressed. (D) Top 5 transcripts/gene expressed. (E) All transcripts from the reference dataset are expressed.

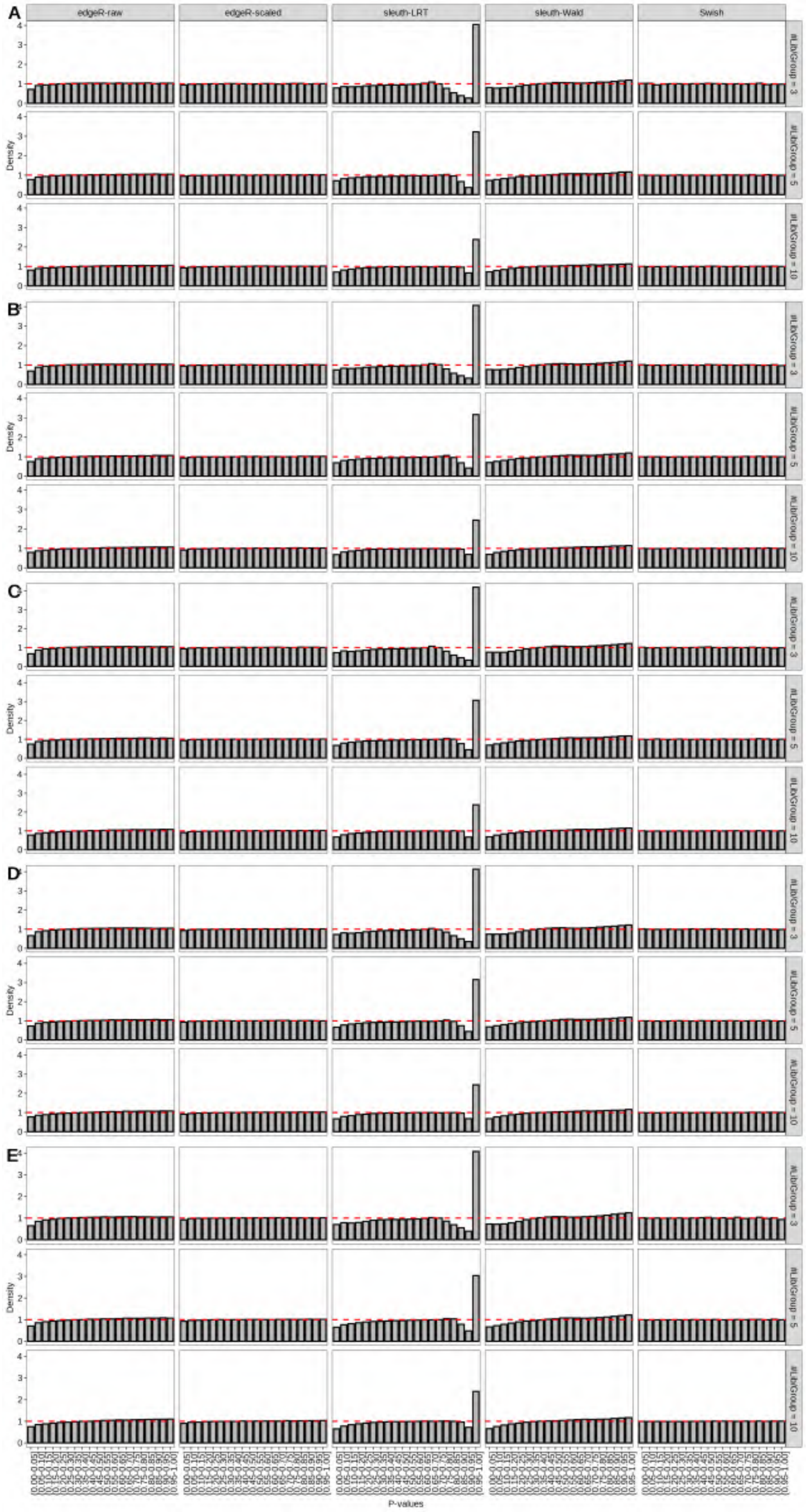

Figure S106: Simulation results. Scenario with mm39 genome, 50bp paired-end reads quantified with Salmon, and unbalanced libraries. Density histograms of p-values from null simulations (averaged over 20 simulations). (A) Top 2 transcripts/gene expressed. (B) Top 3 transcripts/gene expressed. (C) Top 4 transcripts/gene expressed. (D) Top 5 transcripts/gene expressed. (E) All transcripts from the reference dataset are expressed.

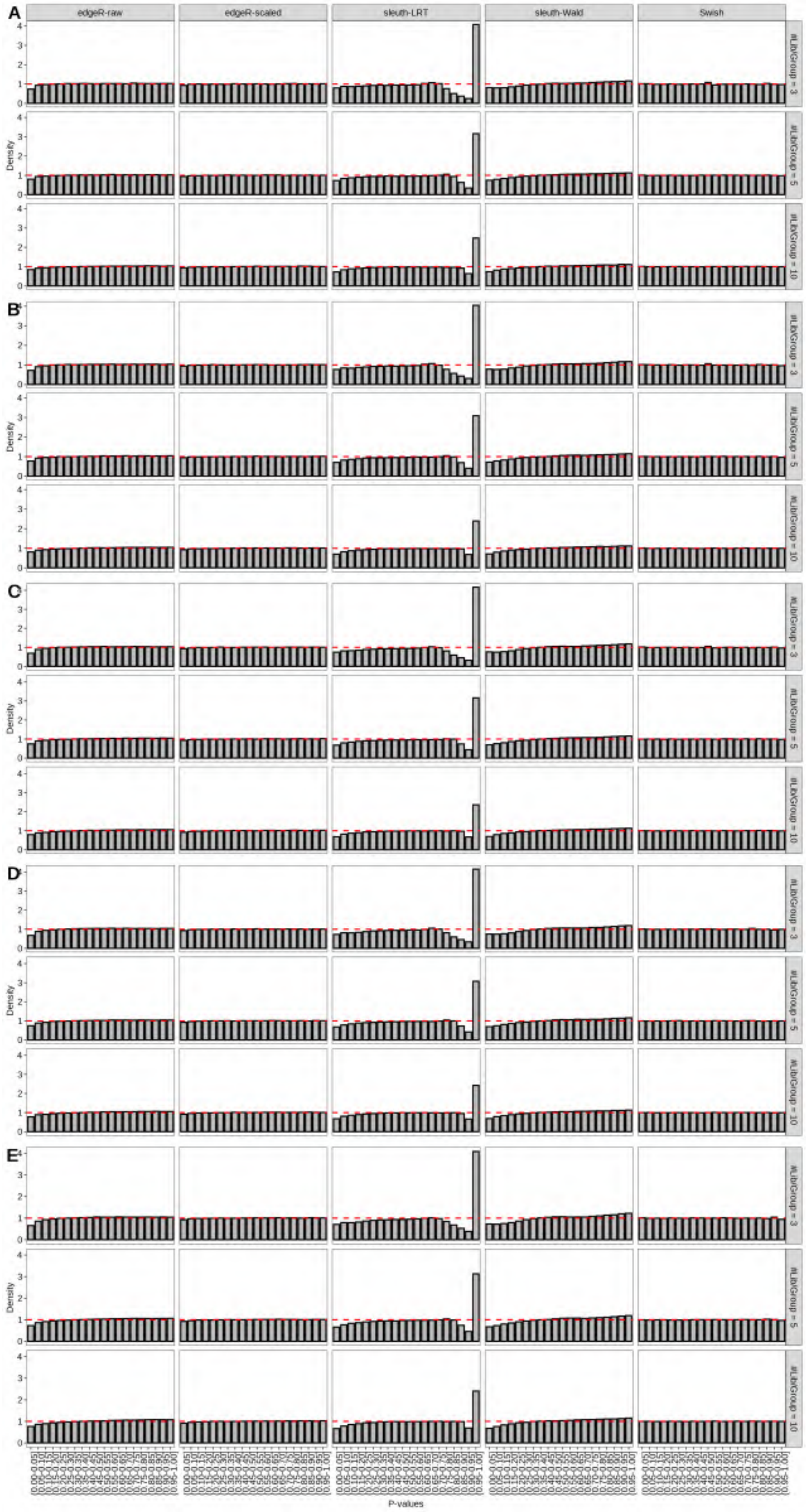

Figure S107: Simulation results. Scenario with mm39 genome, 75bp paired-end reads quantified with Salmon, and unbalanced libraries. Density histograms of p-values from null simulations (averaged over 20 simulations). (A) Top 2 transcripts/gene expressed. (B) Top 3 transcripts/gene expressed. (C) Top 4 transcripts/gene expressed. (D) Top 5 transcripts/gene expressed. (E) All transcripts from the reference dataset are expressed.

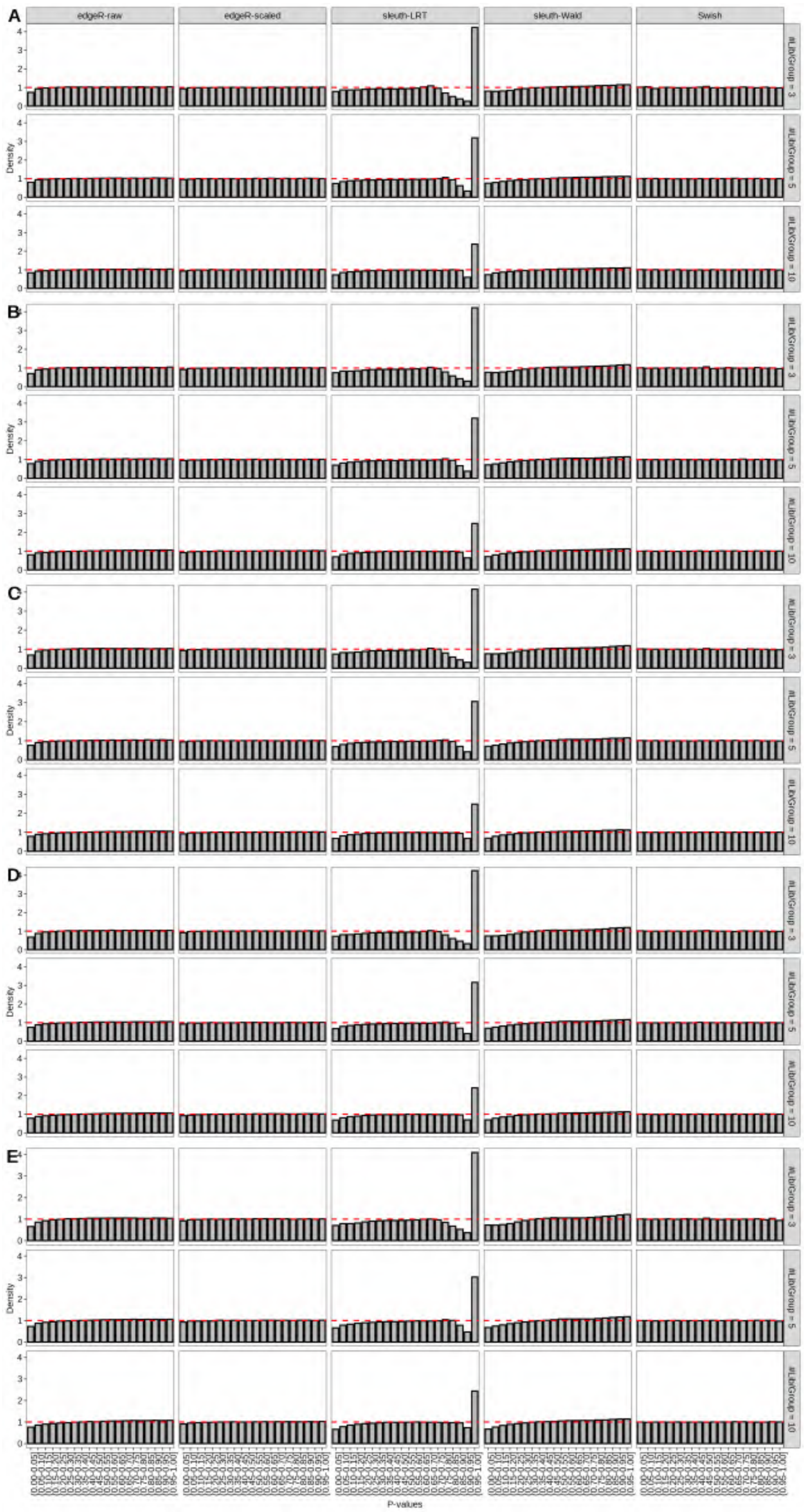

Figure S108: Simulation results. Scenario with mm39 genome, 100bp paired-end reads quantified with Salmon, and unbalanced libraries. Density histograms of p-values from null simulations (averaged over 20 simulations). (A) Top 2 transcripts/gene expressed. (B) Top 3 transcripts/gene expressed. (C) Top 4 transcripts/gene expressed. (D) Top 5 transcripts/gene expressed. (E) All transcripts from the reference dataset are expressed.

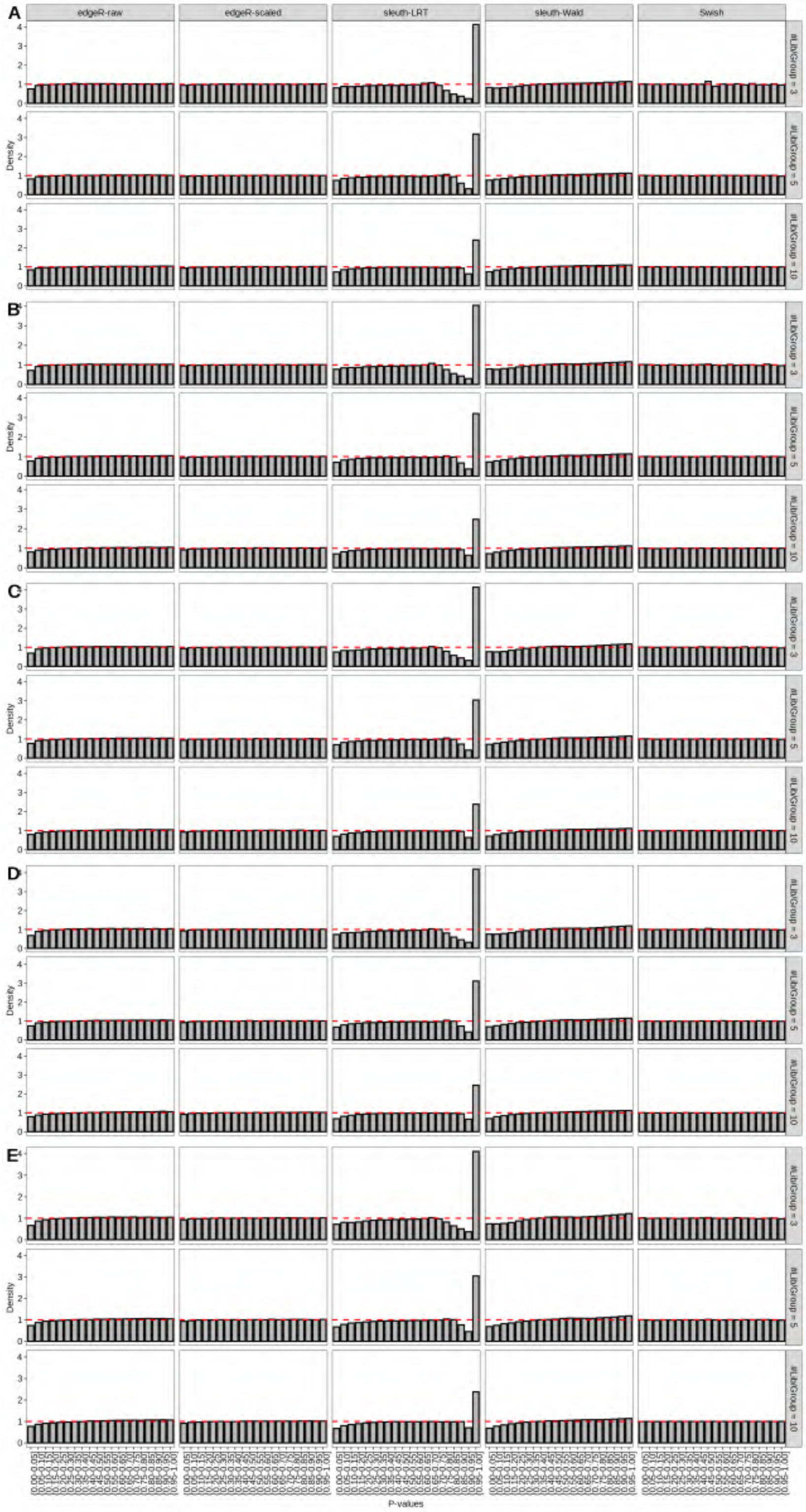

Figure S109: Simulation results. Scenario with mm39 genome, 125bp paired-end reads quantified with Salmon, and unbalanced libraries. Density histograms of p-values from null simulations (averaged over 20 simulations). (A) Top 2 transcripts/gene expressed. (B) Top 3 transcripts/gene expressed. (C) Top 4 transcripts/gene expressed. (D) Top 5 transcripts/gene expressed. (E) All transcripts from the reference dataset are expressed.

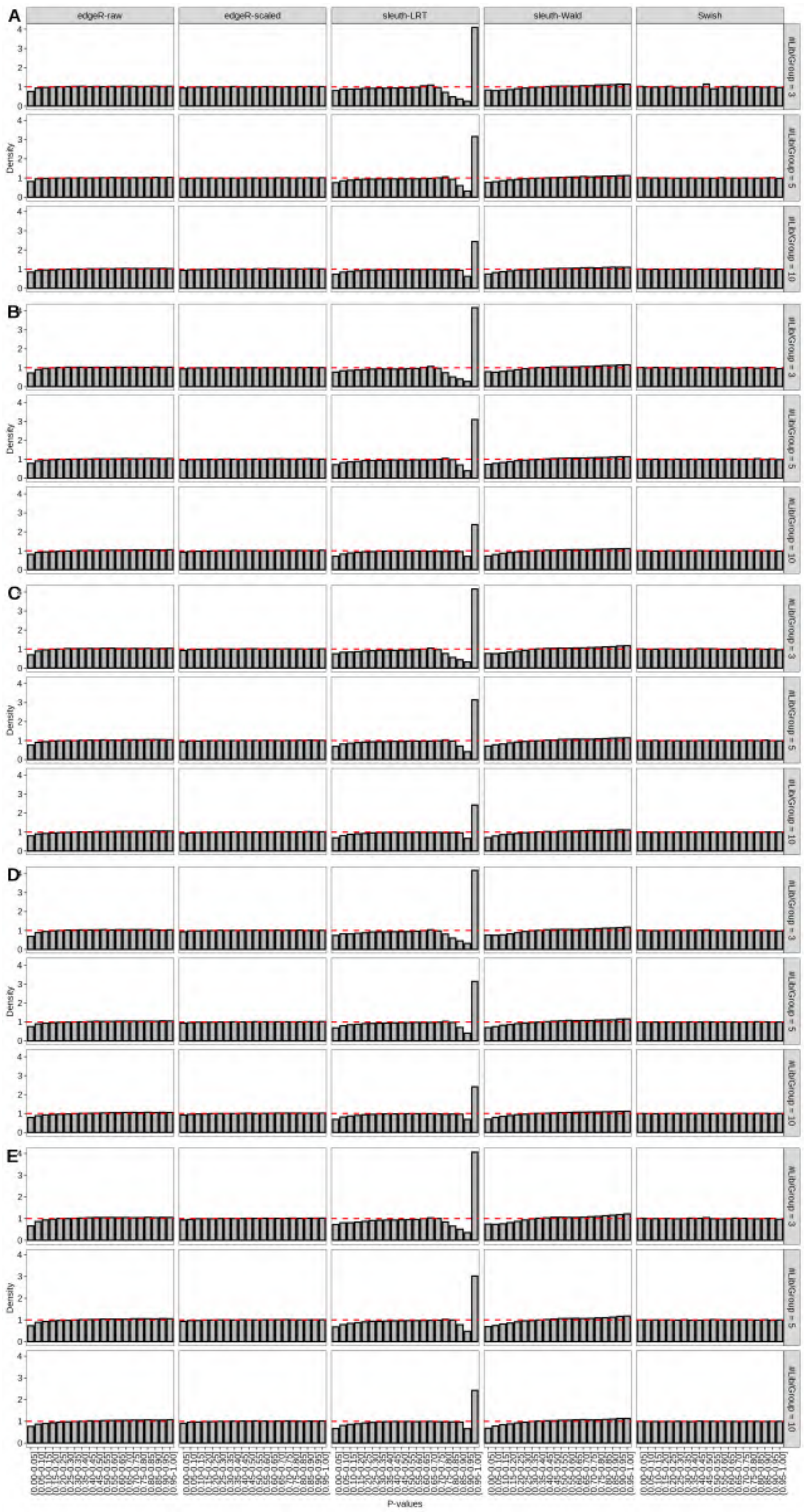

Figure S110: Simulation results. Scenario with mm39 genome, 150bp paired-end reads quantified with Salmon, and unbalanced libraries. Density histograms of p-values from null simulations (averaged over 20 simulations). (A) Top 2 transcripts/gene expressed. (B) Top 3 transcripts/gene expressed. (C) Top 4 transcripts/gene expressed. (D) Top 5 transcripts/gene expressed. (E) All transcripts from the reference dataset are expressed.

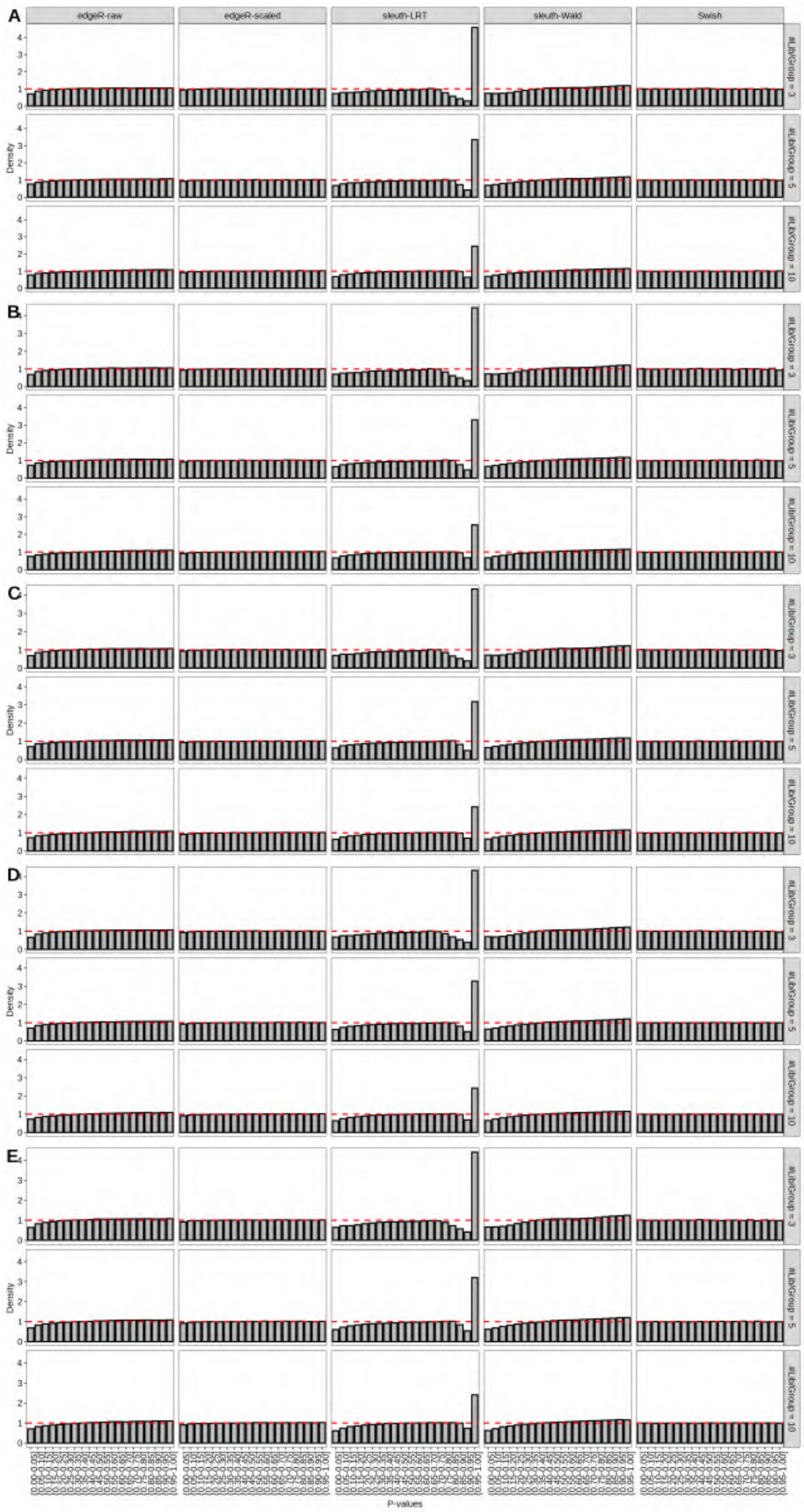

Figure S111: Simulation results. Scenario with mm39 genome, 50bp single-end reads quantified with kallisto, and unbalanced libraries. Density histograms of p-values from null simulations (averaged over 20 simulations). (A) Top 2 transcripts/gene expressed. (B) Top 3 transcripts/gene expressed. (C) Top 4 transcripts/gene expressed. (D) Top 5 transcripts/gene expressed. (E) All transcripts from the reference dataset are expressed.

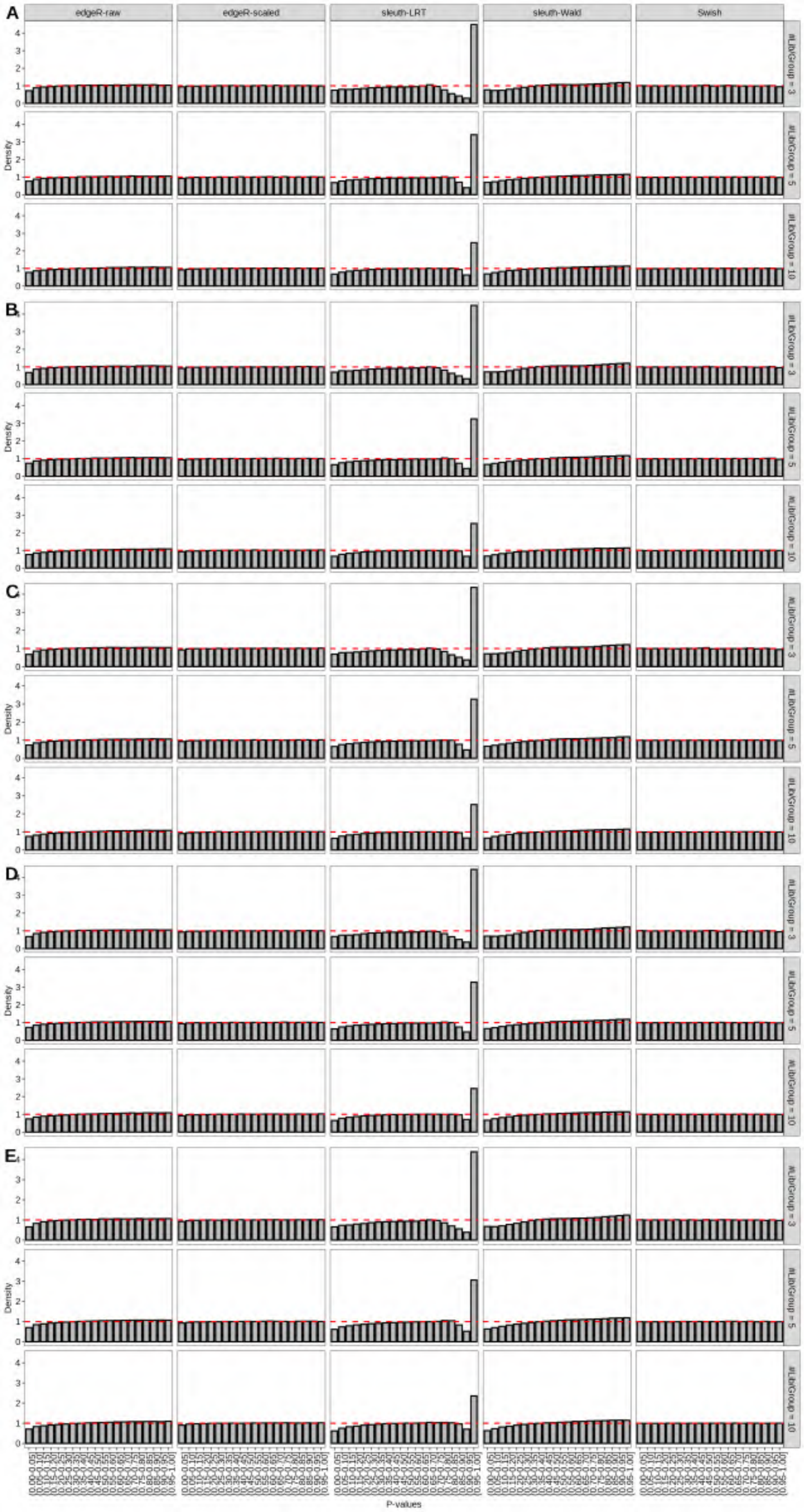

Figure S112: Simulation results. Scenario with mm39 genome, 75bp single-end reads quantified with kallisto, and unbalanced libraries. Density histograms of p-values from null simulations (averaged over 20 simulations). (A) Top 2 transcripts/gene expressed. (B) Top 3 transcripts/gene expressed. (C) Top 4 transcripts/gene expressed. (D) Top 5 transcripts/gene expressed. (E) All transcripts from the reference dataset are expressed.

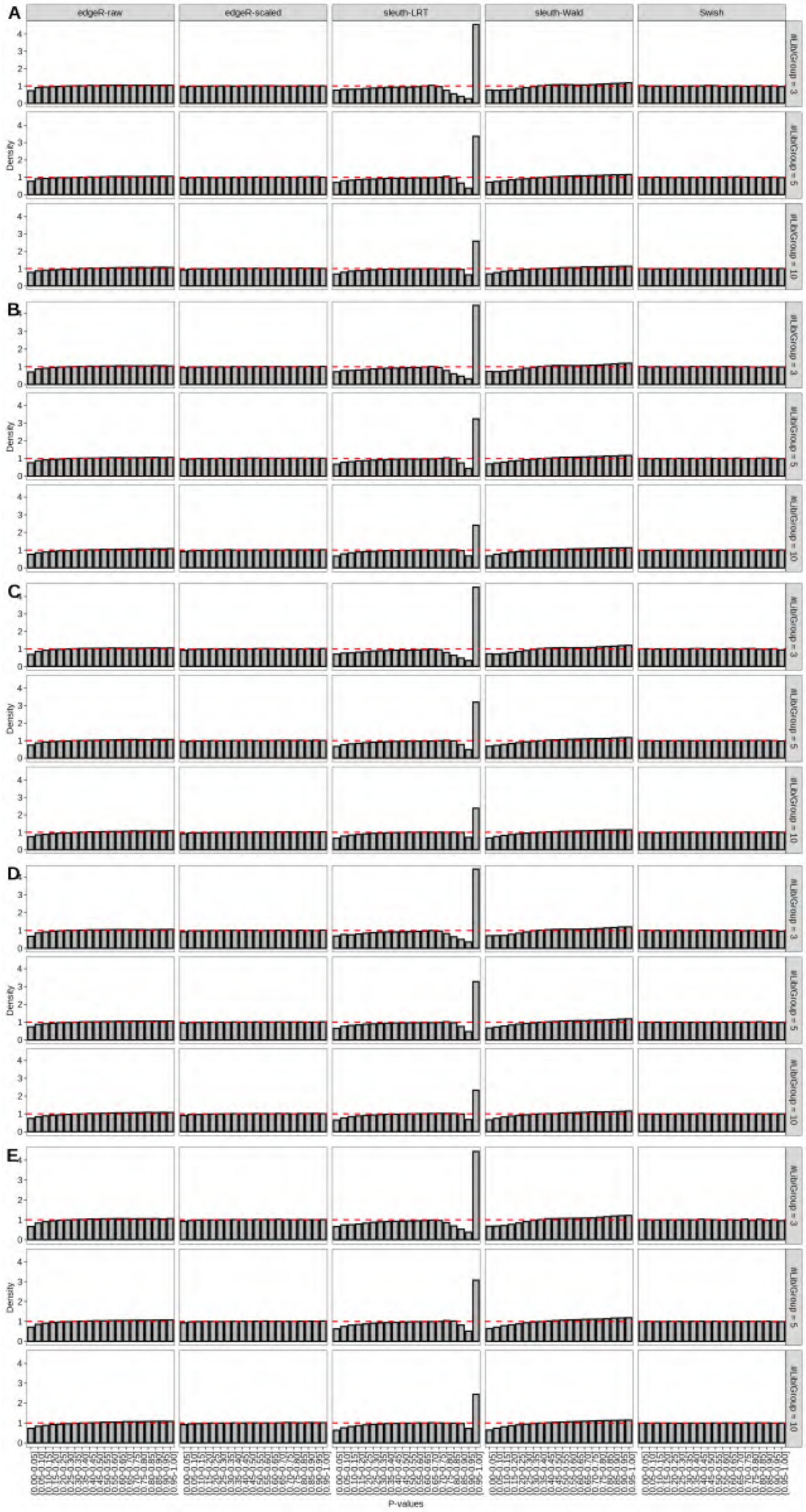

Figure S113: Simulation results. Scenario with mm39 genome, 100bp single-end reads quantified with kallisto, and unbalanced libraries. Density histograms of p-values from null simulations (averaged over 20 simulations). (A) Top 2 transcripts/gene expressed. (B) Top 3 transcripts/gene expressed. (C) Top 4 transcripts/gene expressed. (D) Top 5 transcripts/gene expressed. (E) All transcripts from the reference dataset are expressed.

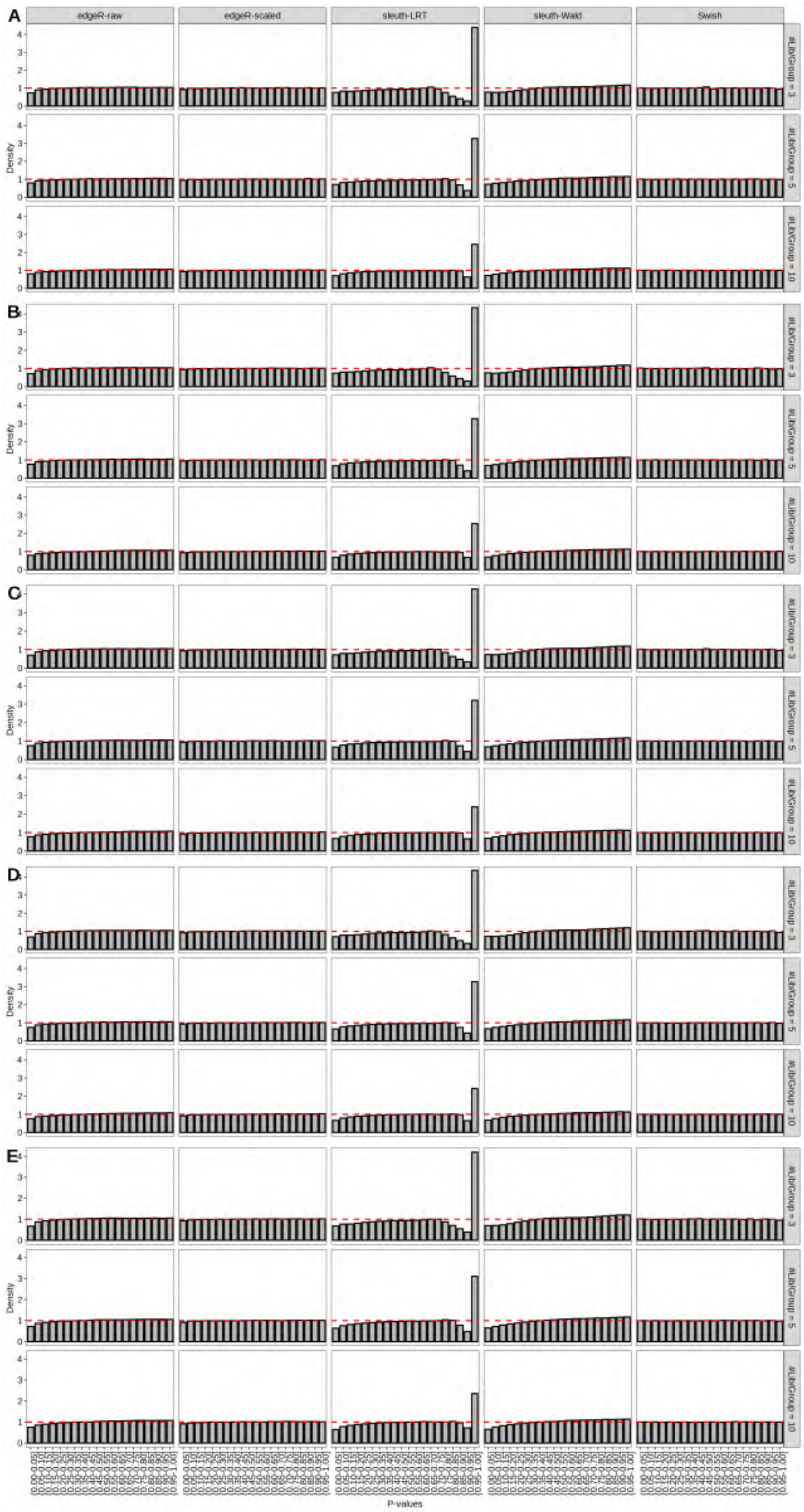

Figure S114: Simulation results. Scenario with mm39 genome, 125bp single-end reads quantified with kallisto, and unbalanced libraries. Density histograms of p-values from null simulations (averaged over 20 simulations). (A) Top 2 transcripts/gene expressed. (B) Top 3 transcripts/gene expressed. (C) Top 4 transcripts/gene expressed. (D) Top 5 transcripts/gene expressed. (E) All transcripts from the reference dataset are expressed.

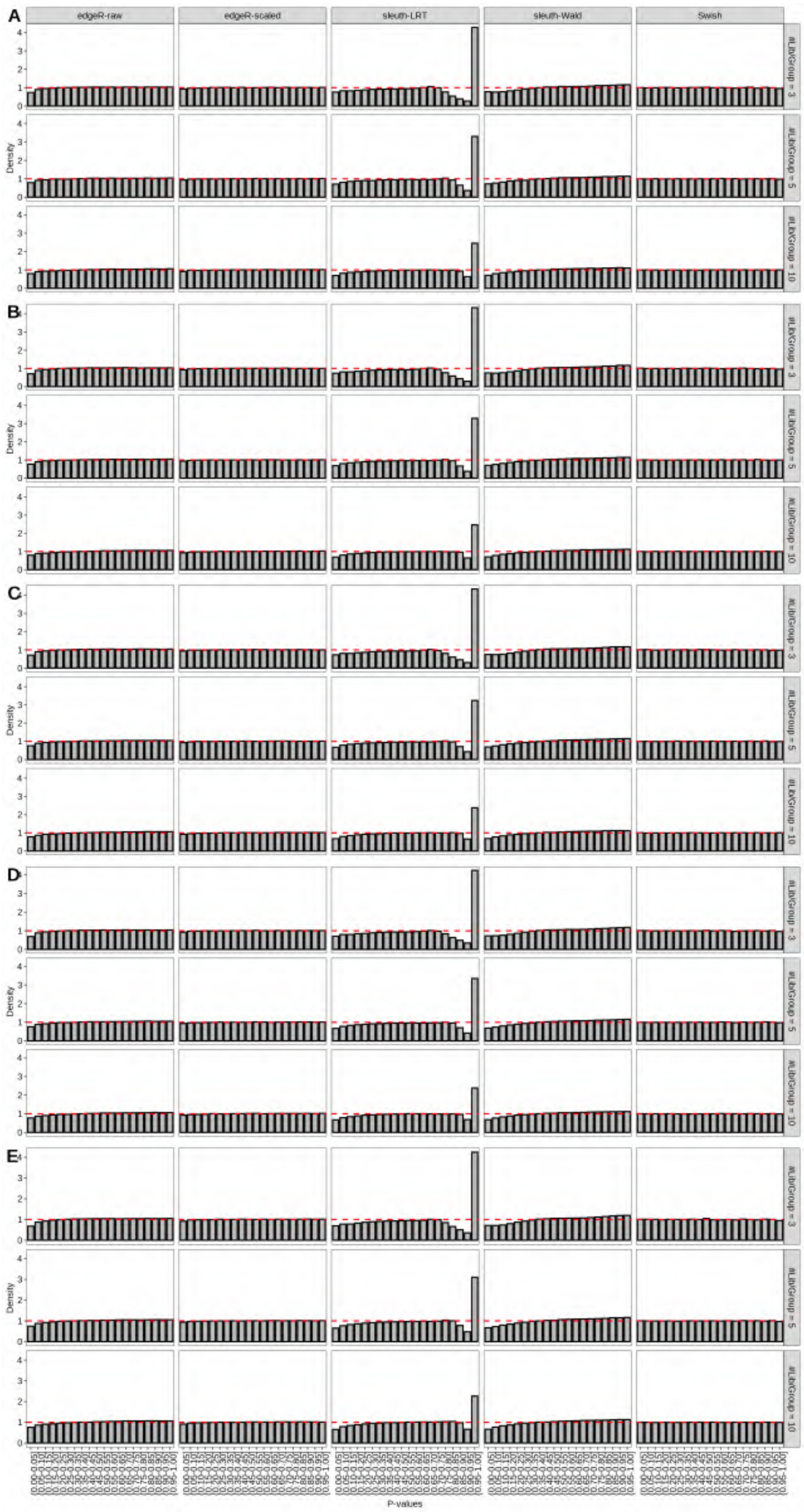

Figure S115: Simulation results. Scenario with mm39 genome, 150bp single-end reads quantified with kallisto, and unbalanced libraries. Density histograms of p-values from null simulations (averaged over 20 simulations). (A) Top 2 transcripts/gene expressed. (B) Top 3 transcripts/gene expressed. (C) Top 4 transcripts/gene expressed. (D) Top 5 transcripts/gene expressed. (E) All transcripts from the reference dataset are expressed.

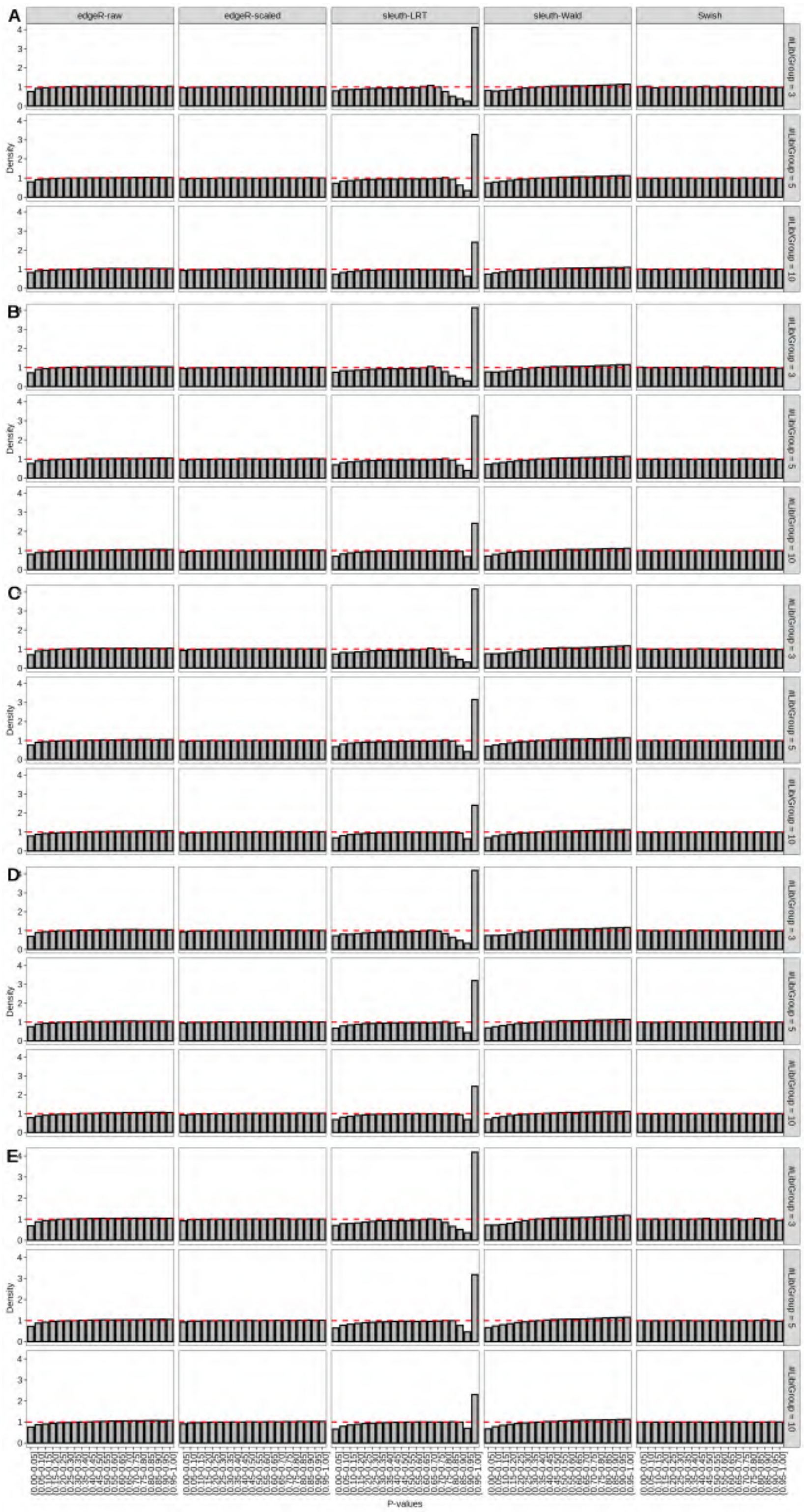

Figure S116: Simulation results. Scenario with mm39 genome, 50bp paired-end reads quantified with kallisto, and unbalanced libraries. Density histograms of p-values from null simulations (averaged over 20 simulations). (A) Top 2 transcripts/gene expressed. (B) Top 3 transcripts/gene expressed. (C) Top 4 transcripts/gene expressed. (D) Top 5 transcripts/gene expressed. (E) All transcripts from the reference dataset are expressed.

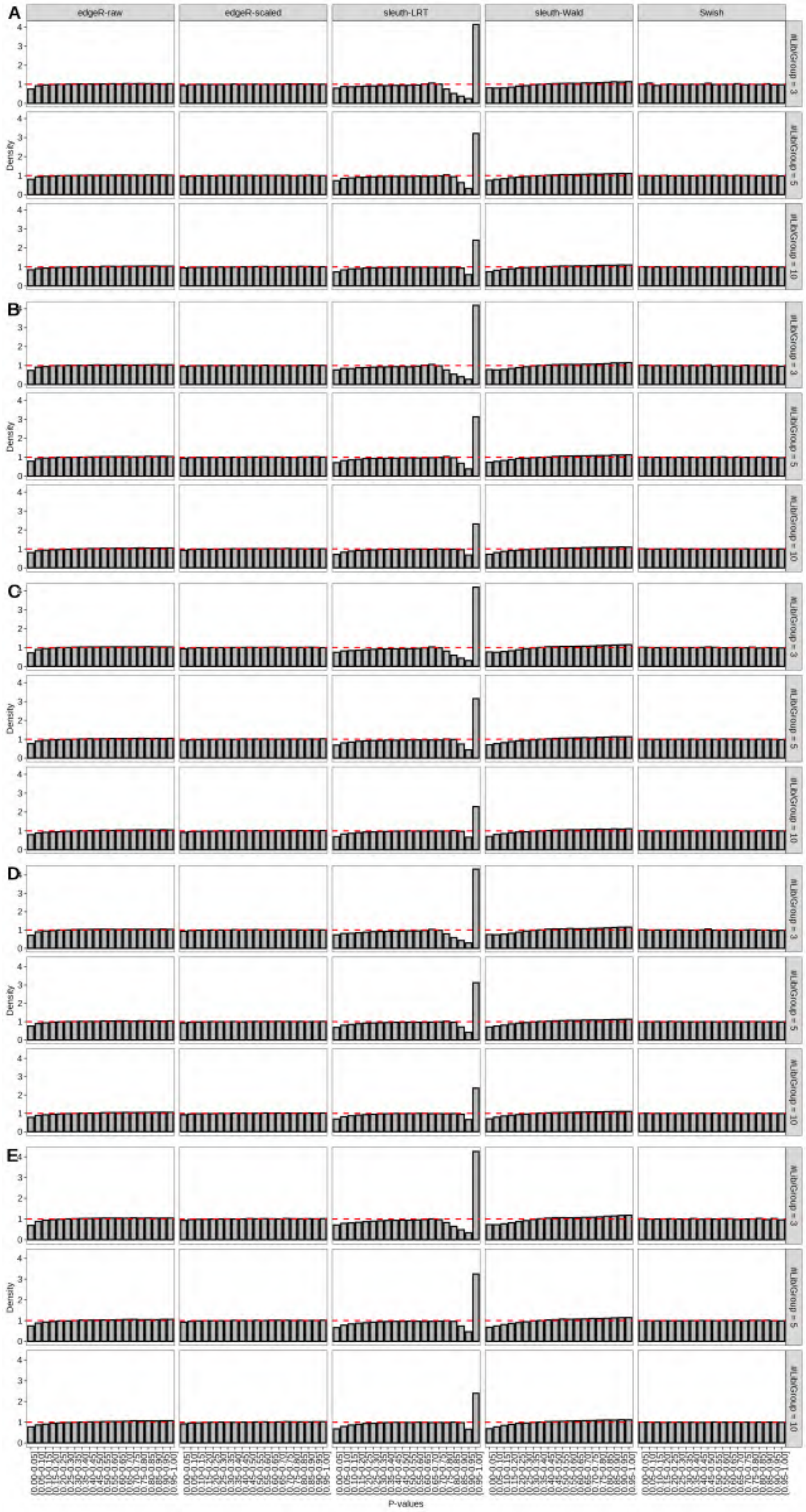

Figure S117: Simulation results. Scenario with mm39 genome, 75bp paired-end reads quantified with kallisto, and unbalanced libraries. Density histograms of p-values from null simulations (averaged over 20 simulations). (A) Top 2 transcripts/gene expressed. (B) Top 3 transcripts/gene expressed. (C) Top 4 transcripts/gene expressed. (D) Top 5 transcripts/gene expressed. (E) All transcripts from the reference dataset are expressed.

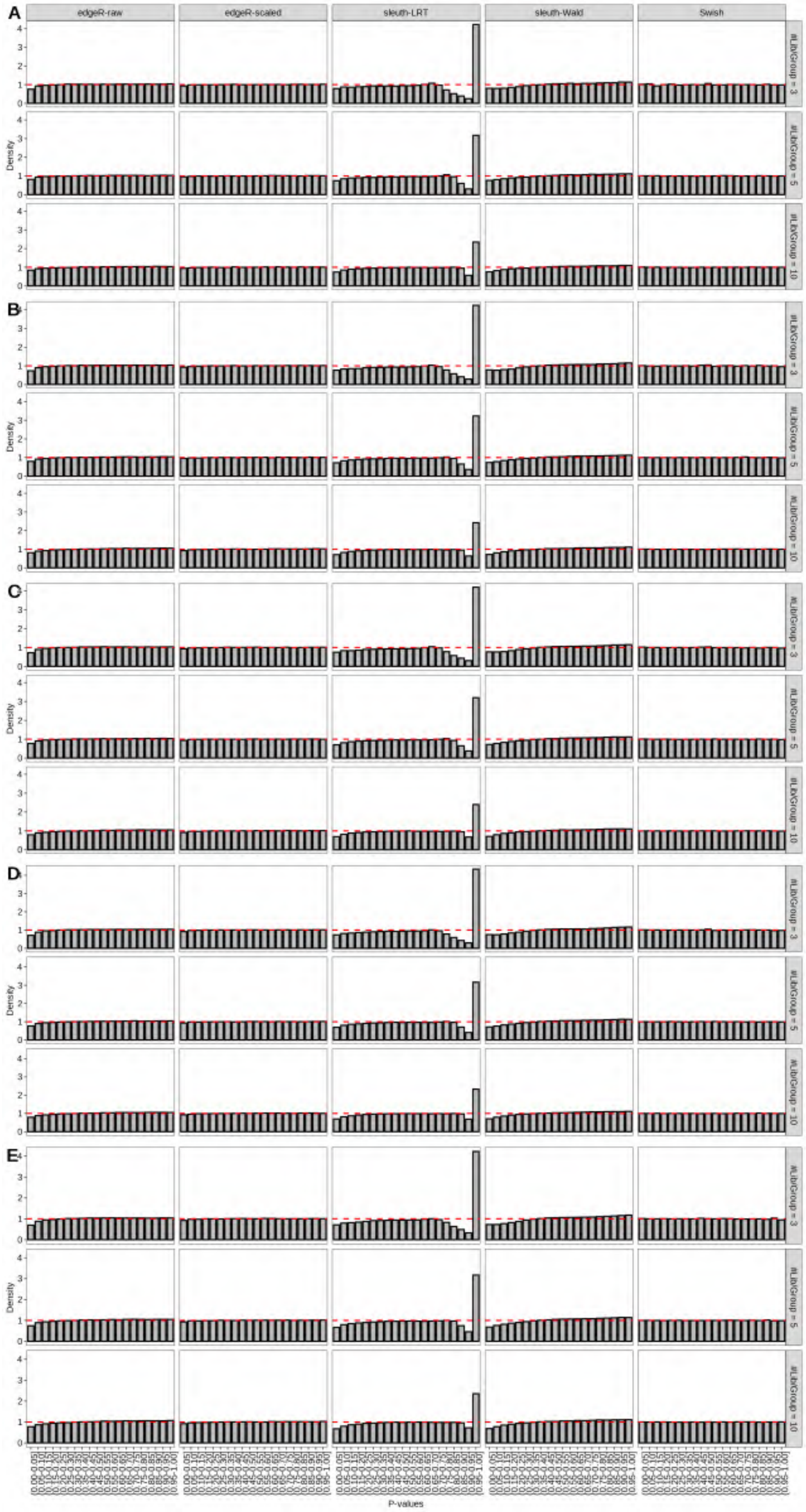

Figure S118: Simulation results. Scenario with mm39 genome, 100bp paired-end reads quantified with kallisto, and unbalanced libraries. Density histograms of p-values from null simulations (averaged over 20 simulations). (A) Top 2 transcripts/gene expressed. (B) Top 3 transcripts/gene expressed. (C) Top 4 transcripts/gene expressed. (D) Top 5 transcripts/gene expressed. (E) All transcripts from the reference dataset are expressed.

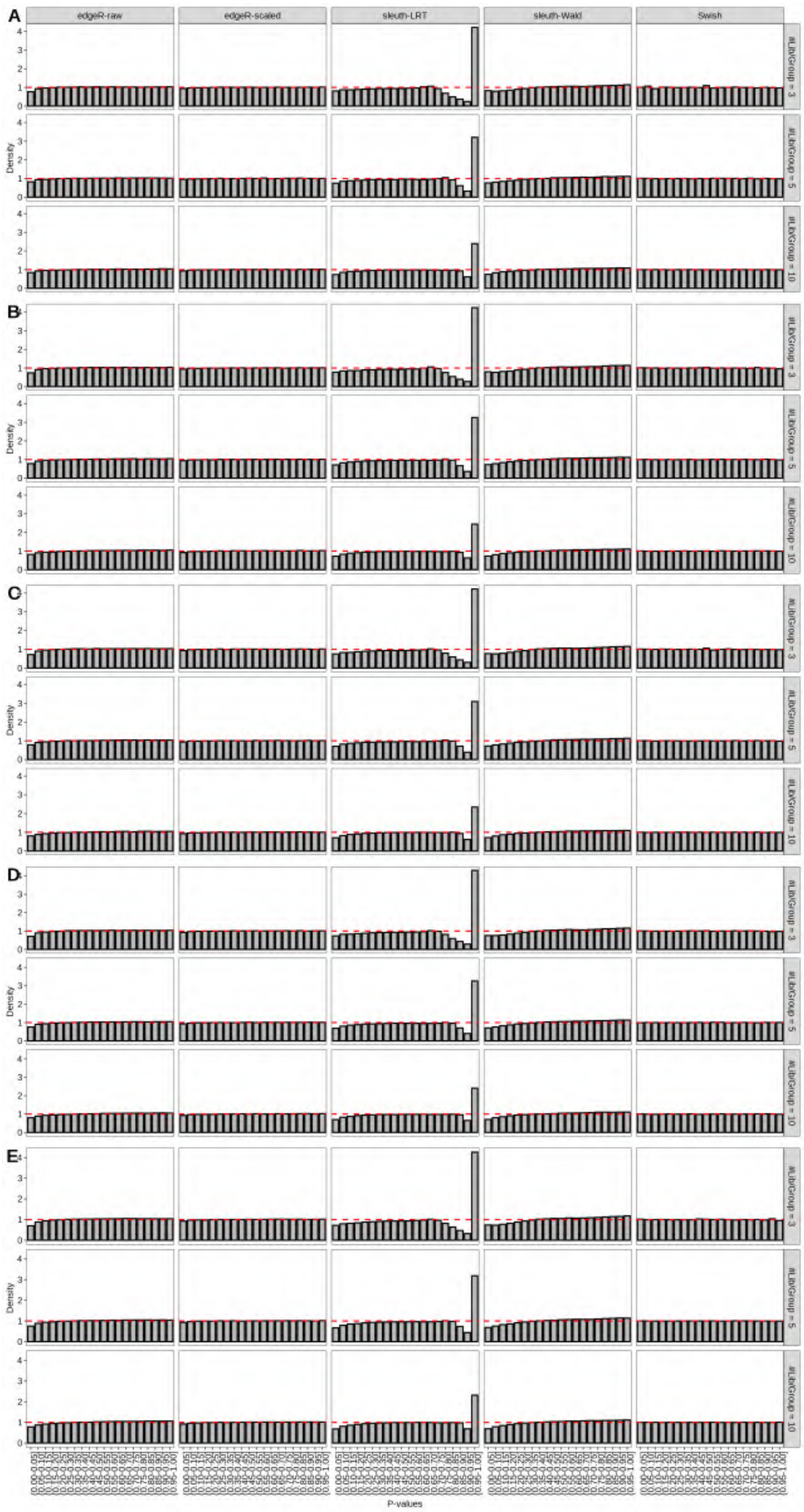

Figure S119: Simulation results. Scenario with mm39 genome, 125bp paired-end reads quantified with kallisto, and unbalanced libraries. Density histograms of p-values from null simulations (averaged over 20 simulations). (A) Top 2 transcripts/gene expressed. (B) Top 3 transcripts/gene expressed. (C) Top 4 transcripts/gene expressed. (D) Top 5 transcripts/gene expressed. (E) All transcripts from the reference dataset are expressed.

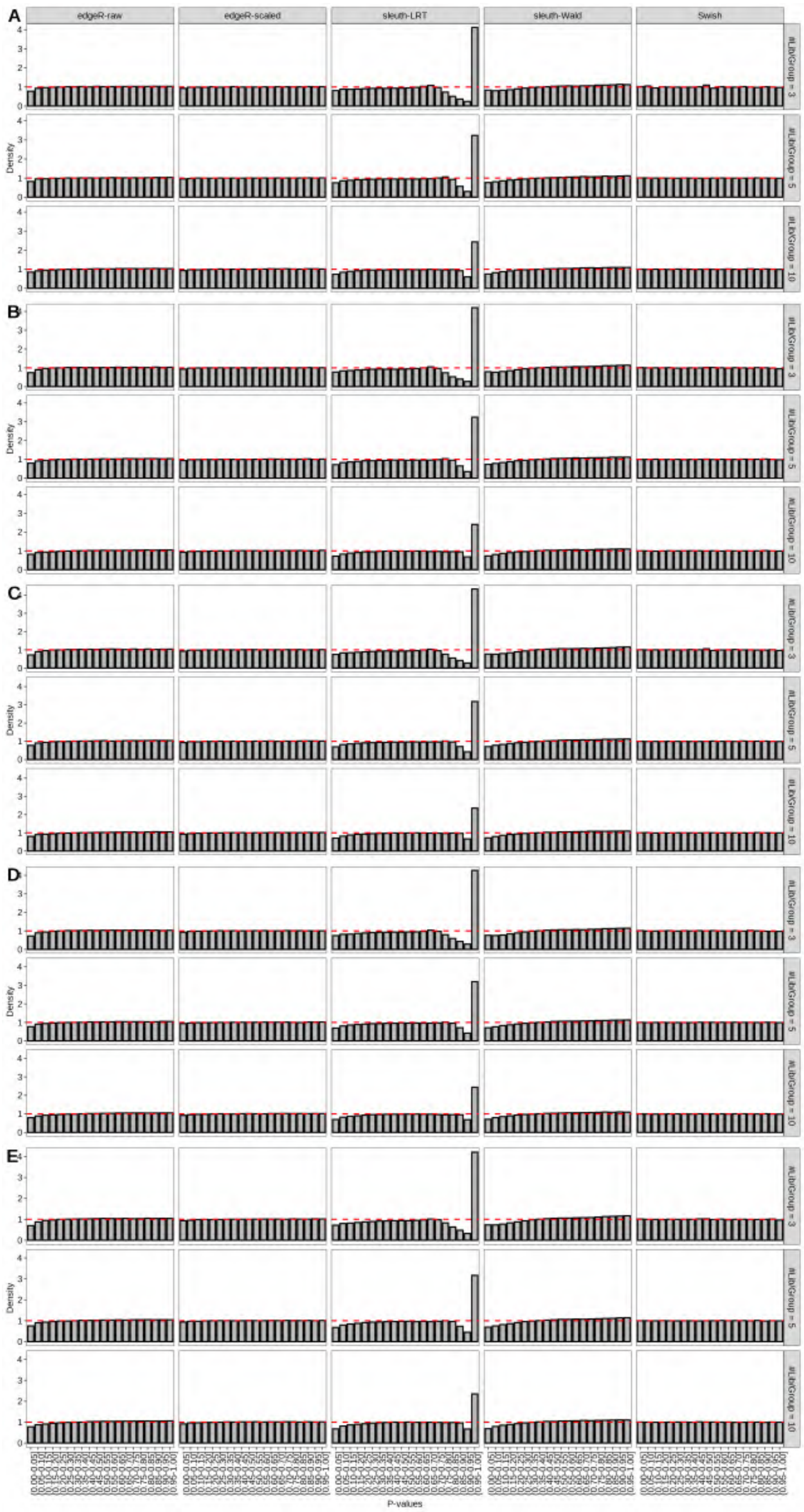

Figure S120: Simulation results. Scenario with mm39 genome, 150bp paired-end reads quantified with kallisto, and unbalanced libraries. Density histograms of p-values from null simulations (averaged over 20 simulations). (A) Top 2 transcripts/gene expressed. (B) Top 3 transcripts/gene expressed. (C) Top 4 transcripts/gene expressed. (D) Top 5 transcripts/gene expressed. (E) All transcripts from the reference dataset are expressed.

### 2 Variance model for transcript counts

Following the same strategy presented in our simulation study, we simulated data with the Rsubread package and the function `simReads` (Liao, Smyth, and Shi (2019)) from 10 technical replicates by assuming no variation over samples for the underlying transcript-specific expression level. Next, we quantified the expression of transcripts with kallisto (Bray et al. (2016)) for the simulated technical replicate samples. We then estimated transcript-specific means and variances for both true counts and estimated counts and compared the mean-variance relationship between both simulated and quantified counts (Figure S121).

As expected, the estimated mean-variance relationship of true counts resembled the one from a Poisson distribution (i.e. mean = variance). In contrast, for quantified counts, we observed that the estimated variance for certain transcripts was much larger than the estimated mean. Specifically, for transcripts associated with multi-transcripts genes, the observed variance of transcript-level counts was higher than the mean, whereas for transcripts associated with single-transcript genes their variance function still resembled the one from a Poisson model.

The resulting variance of transcript-level counts appears to be a linear function of the mean. The extra-to-Poisson variation is transcript-specific, as it depends on the degree of overlap of each transcript to other annotated transcripts; for certain transcripts such an extra variation is nearly non-existent (as for transcripts that do not overlap any other annotated transcripts), and for other transcripts the extra variation is likely to be influenced by the amount of sequence overlap with other annotated transcripts. These results motivated the use of a quasi-Poisson model for transcript quantification.

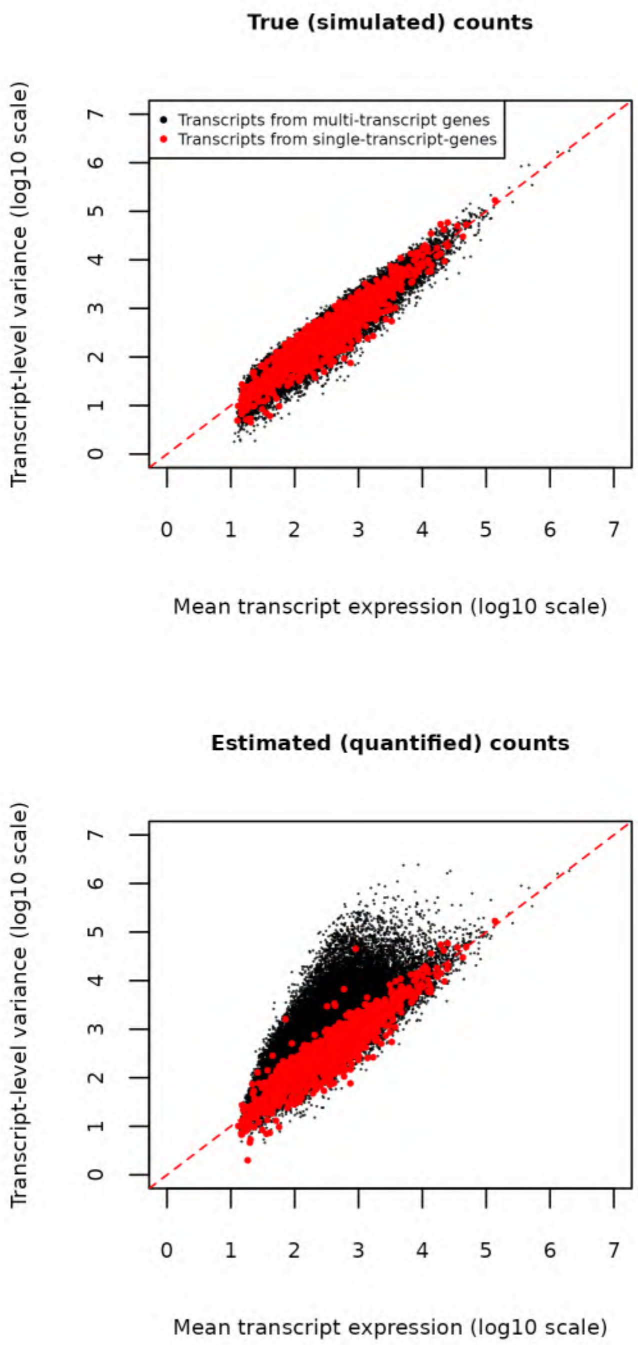

Figure S121: Quantification step introduces quasi-Poisson variation to transcript-level counts. Data from 10 simulated technical replicate samples by assuming no variation over samples for the underlying transcript-specific expression level. At the top, true (simulated) counts from Rsubread's simReads output for data generated as technical replicates (variance = mean). At the bottom, estimated (quantified) counts as output by kallisto from the same set of technical replicates.

### 3 Differential transcript expression in human adenocarcinoma cell lines

#### 3.1 Mapping ambiguity overdispersion increases in annotated transcript-rich loci

We applied the proposed mapping ambiguity overdispersion estimator at the gene-level to quantify the ambiguity associated with gene-level counts obtained post transcript quantification. To this end, we summarized transcript-level quantifications from Salmon to the gene-level with the R/Bioconductor package `tximport` (Soneson, Love, and Robinson (2015); Figure S122) and made use of the presented empirical Bayes mapping ambiguity overdispersion estimator.

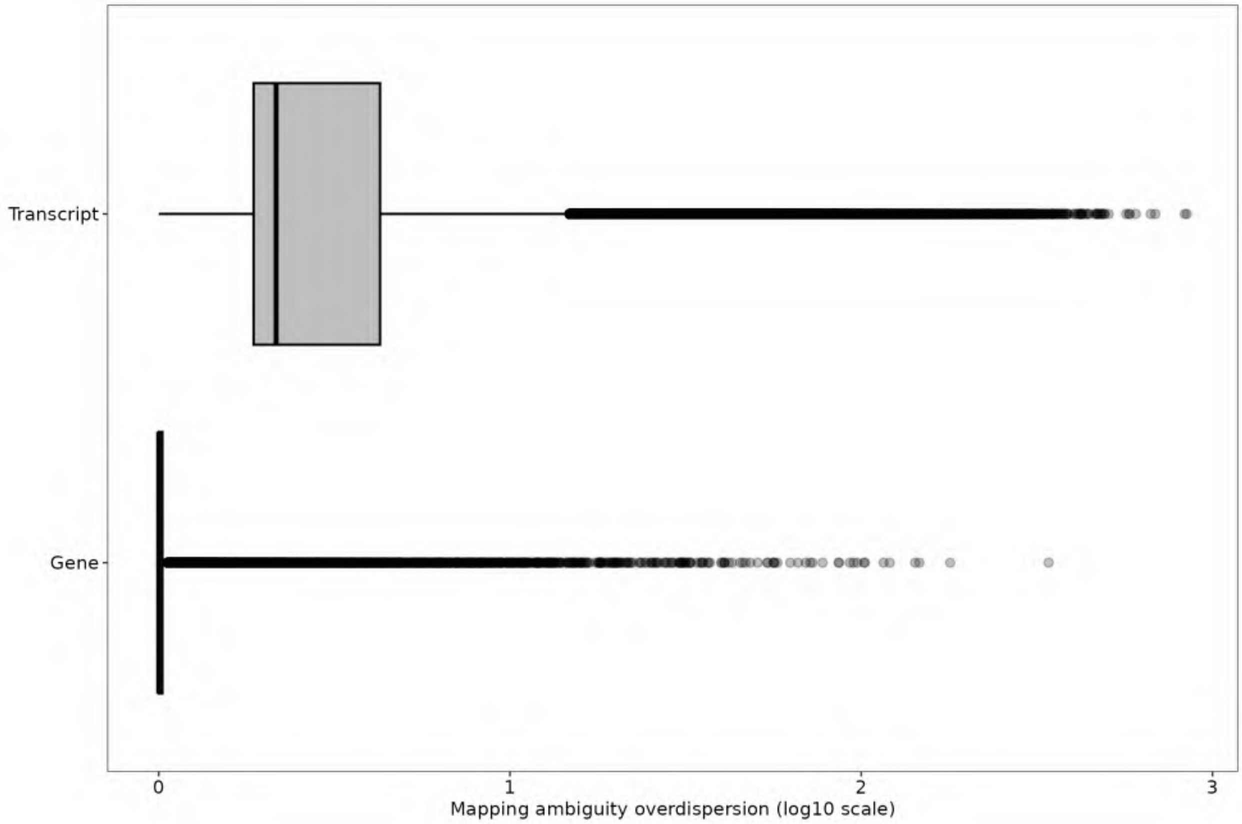

Figure S122: Distribution of mapping amiguity overdispersion computed at the transcript-level and at the gene-level. The mapping ambiguity overdispersion at the gene-level was estimated with bootstrap counts summarized to the gene-wise counts computed with the function summarizeToGene from the tximport package.

We observed that the mapping ambiguity at the transcript-level, here represented by the mapping ambiguity overdispersion estimates, was much higher than those computed at the gene-level. Yet, we did observe a significant amount of genes that had an associated mapping ambiguity overdispersion estimate substantially high, 2-3 orders of magnitude higher than transcripts associated with single-transcript genes. Upon further inspection, we observed that the majority of such genes did not have an associated Entrez ID identifier and were either gene models, not fully validated genes, or protein-coding genes that overlap such less curated genes.

Table S11: Top 100 genes with largest mapping ambiguity overdispersion. Data from the RNA-seq experiment of the human cell lines generated with paired-end reads (GSE172421).

| Ensembl ID | Symbol | Biotype | Entrez ID | Overdispersion |
|---|---|---|---|---|
| ENSG00000205609.12 | EIF3CL | protein coding | 728689 | 341.55 |
| ENSG00000278996.1 | FP671120.5 | lncRNA | - | 179.20 |
| ENSG00000268861.7 | AC008878.3 | protein coding | 23370 | 146.41 |
| ENSG00000285238.2 | AC006064.6 | protein coding | - | 142.46 |
| ENSG00000015568.13 | RGPD5 | protein coding | 84220 | 120.51 |
| ENSG00000178397.13 | FAM220A | protein coding | 84792 | 115.78 |
| ENSG00000253797.2 | UTP14C | protein coding | 9724 | 102.42 |
| ENSG00000277067.4 | CU634019.2 | lncRNA | 102724843 | 101.82 |
| ENSG00000259040.5 | BLOC1S5-TXNDC5 | protein coding | - | 97.40 |
| ENSG00000254692.1 | AL136295.1 | protein coding | - | 95.07 |
| ENSG00000283782.2 | AC116366.3 | protein coding | - | 92.76 |
| ENSG00000285447.1 | ZNF883 | protein coding | 169834 | 86.45 |
| ENSG00000279809.1 | AC005538.2 | TEC | - | 86.21 |
| ENSG00000259132.1 | AL132780.3 | protein coding | - | 77.63 |
| ENSG00000269547.1 | AC011455.2 | protein coding | - | 72.68 |
| ENSG00000284057.1 | AP001273.2 | protein coding | - | 71.60 |
| ENSG00000154545.16 | MAGED4 | protein coding | 728239 | 69.53 |
| ENSG00000288534.1 | AP001931.2 | protein coding | - | 66.13 |
| ENSG00000257411.2 | AC034102.2 | protein coding | - | 62.78 |
| ENSG00000261771.5 | DNAAF4-CCPG1 | lncRNA | - | 57.70 |
| ENSG00000256861.1 | AC048338.1 | protein coding | - | 56.66 |
| ENSG00000260537.2 | AC012184.2 | protein coding | - | 56.47 |
| ENSG00000202198.1 | AL162581.1 | misc RNA | - | 56.15 |
| ENSG00000273590.4 | SMIM11B | protein coding | 102723553 | 54.19 |
| ENSG00000234289.6 | H2BS1 | protein coding | 54145 | 54.02 |
| ENSG00000268738.3 | HSFX2 | protein coding | 100130086 | 50.76 |
| ENSG00000270276.2 | H4C15 | protein coding | 554313 | 48.35 |
| ENSG00000276077.4 | CU633904.2 | lncRNA | 102724951 | 46.84 |
| ENSG00000286600.1 | AC119427.2 | lncRNA | - | 46.00 |
| ENSG00000235655.3 | H3P6 | processed pseudogene | - | 45.17 |
| ENSG00000249624.9 | AP000295.1 | protein coding | - | 42.82 |
| ENSG00000249590.7 | AC004832.3 | protein coding | - | 41.95 |
| ENSG00000132207.17 | SLX1A | protein coding | 548593 | 40.81 |
| ENSG00000261915.6 | AC026954.2 | protein coding | - | 40.81 |
| ENSG00000283239.1 | AC019257.8 | protein coding | - | 40.72 |
| ENSG00000286075.1 | AC009412.1 | protein coding | - | 40.08 |
| ENSG00000272921.1 | AC005832.4 | protein coding | - | 39.93 |
| ENSG00000270181.3 | BIVM-ERCC5 | protein coding | - | 37.08 |
| ENSG00000183054.11 | RGPD6 | protein coding | 729540 | 36.75 |
| ENSG00000180658.4 | OR2A4 | protein coding | 79541 | 36.52 |
| ENSG00000249839.1 | AC011330.1 | unprocessed pseudogene | - | 35.48 |
| ENSG00000196826.7 | AC008758.1 | protein coding | - | 35.27 |
| ENSG00000269900.3 | RMRP | lncRNA | - | 35.18 |
| ENSG00000270316.1 | BORCS7-ASMT | protein coding | - | 34.80 |
| ENSG00000130283.9 | GDF1 | protein coding | 2657 | 34.08 |

Table S11: Top 100 genes with largest mapping ambiguity overdispersion. Data from the RNA-seq experiment of the human cell lines generated with paired-end reads (GSE172421). *(continued)*

| Ensembl ID | Symbol | Biotype | Entrez ID | Overdispersion |
|---|---|---|---|---|
| ENSG00000260371.1 | AC026464.3 | protein coding | - | 32.72 |
| ENSG00000285258.1 | ATXN7 | protein coding | 6314 | 32.57 |
| ENSG00000188223.9 | AD000671.1 | protein coding | - | 32.55 |
| ENSG00000254673.1 | AC110275.1 | protein coding | - | 32.18 |
| ENSG00000271672.1 | DUXAP8 | transcribed processed pseudogene | - | 31.58 |
| ENSG00000167131.17 | CCDC103 | protein coding | 388389 | 31.58 |
| ENSG00000231259.5 | AC125232.1 | unprocessed pseudogene | - | 31.44 |
| ENSG00000256349.1 | AP002748.5 | protein coding | - | 31.33 |
| ENSG00000160201.11 | U2AF1 | protein coding | 7307 | 30.92 |
| ENSG00000234964.4 | FABP5P7 | processed pseudogene | - | 30.73 |
| ENSG00000284554.2 | AL022318.4 | protein coding | - | 30.63 |
| ENSG00000258465.8 | AL139011.2 | protein coding | - | 30.63 |
| ENSG00000124208.16 | TMEM189-UBE2V1 | protein coding | 387522 | 30.40 |
| ENSG00000184110.14 | EIF3C | protein coding | 8663 | 30.32 |
| ENSG00000285901.1 | AC008012.1 | protein coding | - | 29.90 |
| ENSG00000284989.1 | AL451062.4 | protein coding | - | 29.07 |
| ENSG00000257315.2 | ZBED6 | protein coding | 100381270 | 28.83 |
| ENSG00000285404.1 | Z82190.2 | protein coding | - | 28.80 |
| ENSG00000287856.1 | AL445524.2 | protein coding | - | 28.14 |
| ENSG00000205670.11 | SMIM11A | protein coding | 54065 | 27.82 |
| ENSG00000180581.7 | SRP9P1 | processed pseudogene | - | 27.75 |
| ENSG00000235288.3 | AC099329.1 | lncRNA | - | 27.42 |
| ENSG00000262526.2 | AC120057.2 | protein coding | - | 26.60 |
| ENSG00000285304.1 | Z83844.3 | protein coding | - | 26.17 |
| ENSG00000224831.3 | TMEM183B | processed pseudogene | - | 26.17 |
| ENSG00000285130.2 | AL358113.1 | protein coding | - | 25.92 |
| ENSG00000214076.3 | CPSF1P1 | transcribed processed pseudogene | - | 25.53 |
| ENSG00000267645.5 | AC105052.3 | protein coding | - | 25.42 |
| ENSG00000279423.1 | AL445363.3 | TEC | - | 25.18 |
| ENSG00000268173.3 | AC007192.1 | protein coding | - | 25.18 |
| ENSG00000280987.4 | MATR3 | protein coding | 9782 | 24.44 |
| ENSG00000285585.1 | AC069444.2 | protein coding | - | 24.25 |
| ENSG00000281453.1 | TGFB2-OT1 | lncRNA | - | 23.69 |
| ENSG00000235884.4 | LINC00941 | lncRNA | - | 23.51 |
| ENSG00000271153.1 | RPL23AP88 | processed pseudogene | - | 23.33 |
| ENSG00000285920.2 | AC087721.2 | protein coding | - | 22.44 |
| ENSG00000285053.1 | TBCE | protein coding | 6905 | 22.12 |
| ENSG00000277125.1 | AC211476.10 | unprocessed pseudogene | - | 22.02 |
| ENSG00000283201.1 | AC092329.3 | protein coding | 440519 | 21.68 |
| ENSG00000257767.3 | AC002996.1 | protein coding | - | 21.66 |
| ENSG00000285953.1 | AC000120.4 | protein coding | - | 21.54 |
| ENSG00000270066.3 | AL356488.2 | lncRNA | - | 21.32 |
| ENSG00000285565.1 | AL671762.1 | lncRNA | - | 21.12 |
| ENSG00000270882.2 | H4C14 | protein coding | 8370 | 20.78 |
| ENSG00000198406.7 | BZW1P2 | processed pseudogene | - | 20.66 |

Table S11: Top 100 genes with largest mapping ambiguity overdispersion. Data from the RNA-seq experiment of the human cell lines generated with paired-end reads (GSE172421). *(continued)*

| Ensembl ID | Symbol | Biotype | Entrez ID | Overdispersion |
|---|---|---|---|---|
| ENSG00000269711.1 | AC008763.3 | protein coding | - | 20.47 |
| ENSG00000286098.1 | AC008770.4 | protein coding | 284391 | 20.30 |
| ENSG00000283765.1 | AC131160.1 | protein coding | 55486 | 20.23 |
| ENSG00000264545.2 | AL359922.1 | protein coding | - | 20.11 |
| ENSG00000288380.1 | AC118281.1 | protein coding | - | 20.09 |
| ENSG00000243708.10 | PLA2G4B | protein coding | 100137049 | 20.06 |
| ENSG00000232882.1 | PHKA1P1 | processed pseudogene | - | 19.98 |
| ENSG00000163635.18 | ATXN7 | protein coding | 6314 | 19.82 |
| ENSG00000264668.2 | AC138696.1 | protein coding | - | 19.63 |
| ENSG00000172780.16 | RAB43 | protein coding | 339122 | 19.61 |

#### 3.2 Gene-level analysis

Table S12: edgeR results from a DE analysis at the gene-level for a set of cancer-related genes comparing cell lines H1975 and HCC827. Data from the paired-end RNA-seq experiment of the the human cell lines (GSE172421). Such genes have at least one its transcripts differentially expressed between cell lines (nominal FDR 0.05 at the transcript-level).

| Ensembl ID | Symbol | Entrez ID | logFC | logCPM | F | PValue | FDR |
|---|---|---|---|---|---|---|---|
| ENSG00000132155.12 | RAF1 | 5894 | -0.623 | 5.942 | 39.959 | 1.465e-05 | 3.090e-05 |
| ENSG00000171552.13 | BCL2L1 | 598 | 0.723 | 8.187 | 28.776 | 8.277e-05 | 1.511e-04 |
| ENSG00000105976.15 | MET | 4233 | 0.552 | 9.092 | 24.635 | 1.773e-04 | 3.064e-04 |
| ENSG00000129521.14 | EGLN3 | 112399 | -0.272 | 5.197 | 7.648 | 1.458e-02 | 1.923e-02 |
| ENSG00000142208.16 | AKT1 | 207 | 0.268 | 7.431 | 5.828 | 2.922e-02 | 3.714e-02 |

##### 3.3 Results from single-end read experiment

In Figure S123 we present the mean-difference plot highlighting differentially expressed transcripts between NCI-H1975 and HCC827 cell lines from a DTE analysis of the Illumina short-read RNA-seq experiment generated with single-end read sequencing protocol.

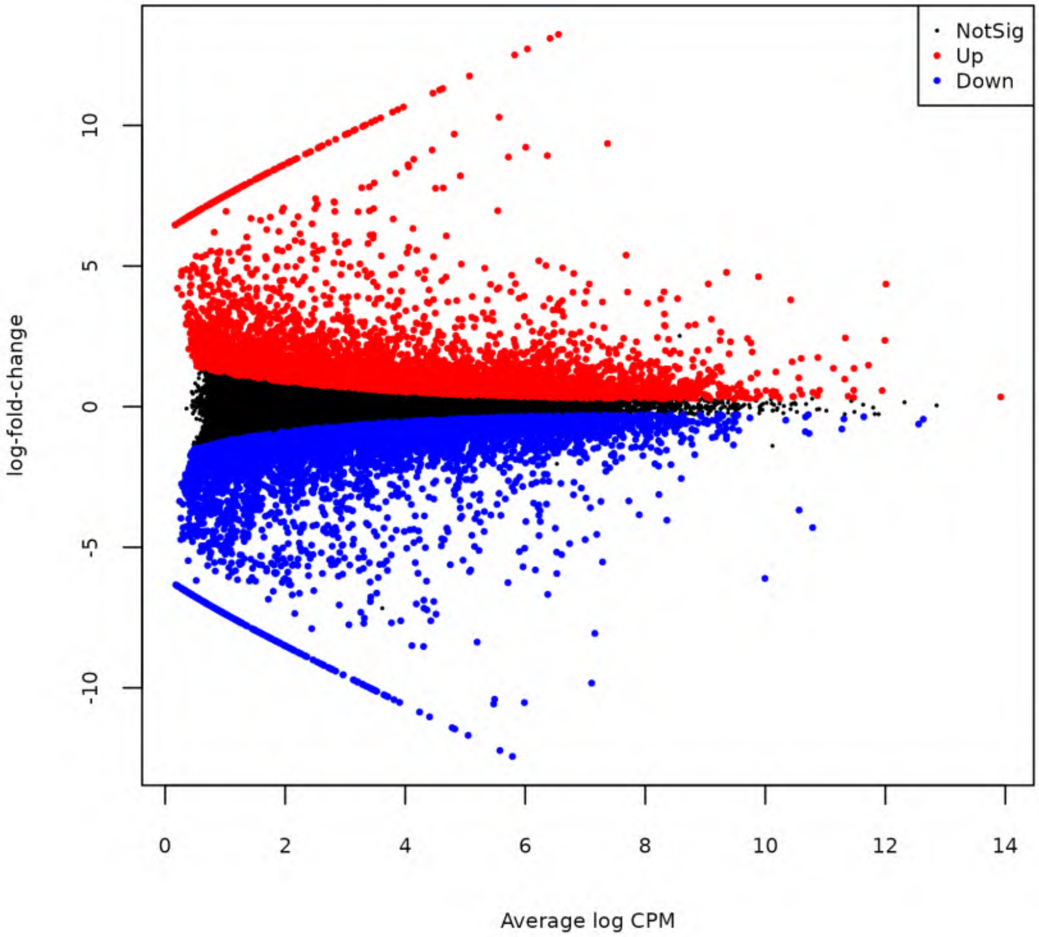

Figure S123: Mean-difference plot highlighting differentially expressed transcripts between NCI-H1975 and HCC827 cell lines from Illumina short single-end read RNA-seq experiment of the human adenocarcinoma cell lines (NCBI Gene Expression Omnibus accession number GSE86337)

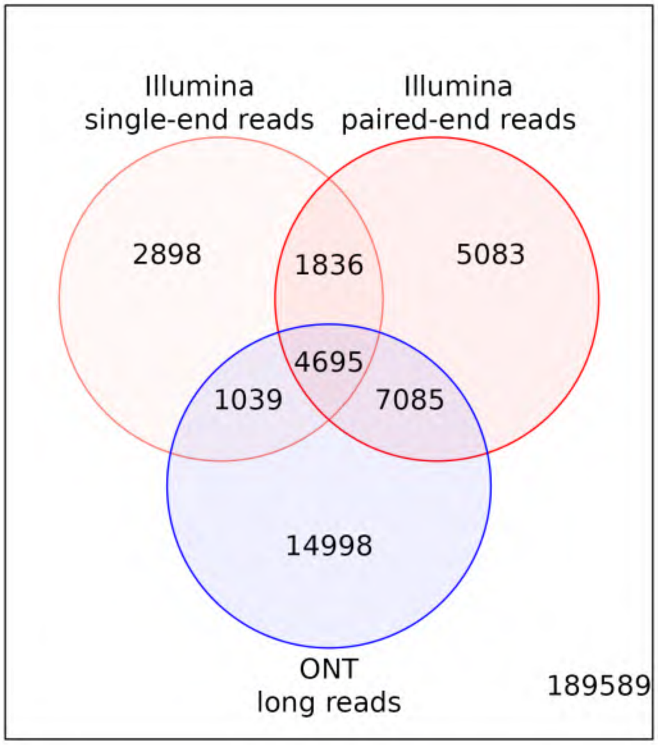

Figure S124: Venn diagram comparing DTE results from edgeR with count scaling using both single- and paired-end read data, as well as with edgeR with raw counts using ONT long-read data.
